# Discovery and chemical optimisation of a Potent, Bi-cyclic (Bicycle^®^) Antimicrobial Inhibitor of *Escherichia coli* PBP3

**DOI:** 10.1101/2024.03.20.581580

**Authors:** Catherine E. Rowland, Hector Newman, Tazmin T. Martin, Rachel Dods, Nikolaos Bournakas, James M. Wagstaff, Nick Lewis, Steven J. Stanway, Matthew Balmforth, Celia Kessler, Katerine van Rietschoten, Dom Bellini, David I. Roper, Adrian J. Lloyd, Christopher G. Dowson, Michael J. Skynner, Paul Beswick, Michael J. Dawson

## Abstract

Penicillin binding proteins (PBPs) are well validated antimicrobial targets, but the prevalence of β-lactamase driven resistance and, more rarely, target-based mutations, necessitates new classes of PBP-targeting drugs. Here we describe the discovery and optimisation of novel, bicyclic peptide (Bicycle^®^) inhibitors of *E. coli* PBP3 (*Ec*PBP3) using a proprietary phage display platform, and their conjugation to linear antimicrobial peptides to confer outer membrane permeation. These molecules exhibited high-affinity binding to *E. coli* PBP3 and a viable spectrum of killing activity against clinically relevant species of the Enterobacterales. X-ray crystallography was used to explore the mode of binding to PBP3, enabling increased target affinity and improvement of *in vitro* stability. These compounds bind to the transpeptidase active site cleft of PBP3 and represent a novel non-β-lactam chemical class of high affinity, non-covalent penicillin binding protein inhibitors.

## Introduction

Penicillin binding proteins (PBPs) function within the peptidoglycan biosynthesis pathway in coordinated multi-protein complexes.^1^ They have distinct functions in the synthesis and remodelling of this core component of the bacterial cell wall.^2^ In the rod-shaped Enterobacterales, peptidoglycan biosynthesis at the site of future cell division (i.e. the septum) is performed by proteins of the divisome complex.^3^ Within the family of genes that encode peptidoglycan production the gene product of *ftsI*, named PBP3 in *E. coli,* is an essential monofunctional, Class B PBP transpeptidase. PBP3 catalyses the cross-linking of peptide stems from adjacent glycan strands in peptidoglycan to create the strong, mesh-like structure of the final polymer (Fig. 1a).^2,4^

**Figure 1:**
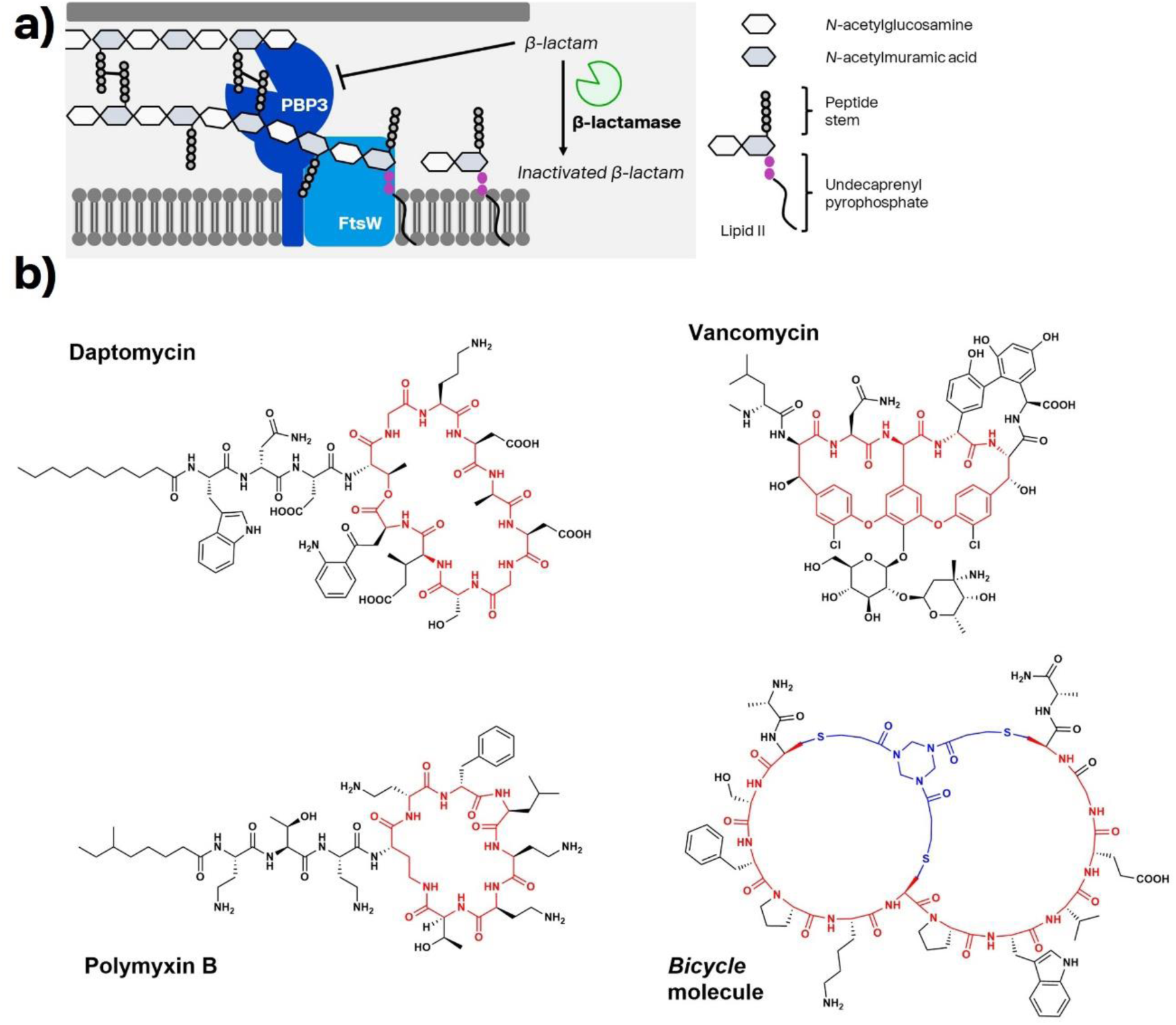
Schematic of *Ec*PBP3 function in cell wall biosynthesis and comparison of a Bicycle molecule to existing cyclic peptide antimicrobials. a) *E. coli* penicillin binding protein 3 (PBP3) works as a functional complex with the SEDS protein and lipid II glycosyltransferase FtsW to build peptidoglycan at the septum, the future site of cell division. β-lactam antibiotics inhibit the transpeptidase activity of PBP3, exerting an antimicrobial effect. One of the major resistance mechanisms is expression of β-lactamase enzymes which cleave the pharmacophore of this antibiotic class. b) Cyclic peptides are a precedented chemotype in existing antimicrobials.

The β-lactam class of antibiotics, which includes penicillins, cephalosporins, monobactams and carbapenems, exhibit efficacy against both Gram-negative and Gram-positive bacteria and are well tolerated, efficacious and broadly prescribed.^5^ Their mechanism of action is through covalently modifying the active site serine in the transpeptidase site of PBPs, leading to the inhibition of cell well biosynthesis and cell death.^5–7^ Many β-lactams show partial selectivity for individual PBPs within a bacterial species, and many clinically used drugs including piperacillin and aztreonam inhibit cell division by preferential binding to PBP3 of *E. coli*.^8^

Resistance to β-lactams can occur through a variety of mechanisms. In Gram-negative bacteria, including *E. coli*, decreased susceptibility to β-lactams results primarily from expression of β-lactamase enzymes, which hydrolyse the pharmacophore of these compounds rendering them inactive.^5^ In response, a number of β-lactamase inhibitors have been developed and introduced into medicine but new classes of β-lactamase enzymes which are recalcitrant to these inhibitors have evolved as a result of this selection pressure.^9^ Novel inhibitors of PBPs which do not suffer from deactivation by hydrolysis of the β-lactam pharmacophore would be highly desirable, to offer a new route to antimicrobial chemotherapy on an established target.

Bicycle molecules are bicyclic peptides formed through the structural constraint of peptides around a trimeric scaffold via the formation of thioether cysteine linkages.^10,11^ Cyclic peptides are common among marketed antimicrobials. The identification of compounds such as daptomycin and vancomycin (Fig. 1b) illustrate that these occupy a privileged chemical space, as compared to typical molecules screened in high-content screening libraries.^12–15^ In addition to the precedence for cyclic peptides as antimicrobials, Bicycle molecules offer additional benefits, namely their ability to be conjugated as well as a larger surface area for binding, when compared to a Lipinski-compliant small molecule. Identification of target-binding bicyclic peptides via phage display (Fig. 2a) confers in-built tolerance to conjugation at the C-terminus. This modularity of Bicycle molecules has been exploited to generate conjugates in fluorescent or radio-labelled formats and with pharmacologically active moieties such as cytotoxins.^16–19^

**Figure 2:**
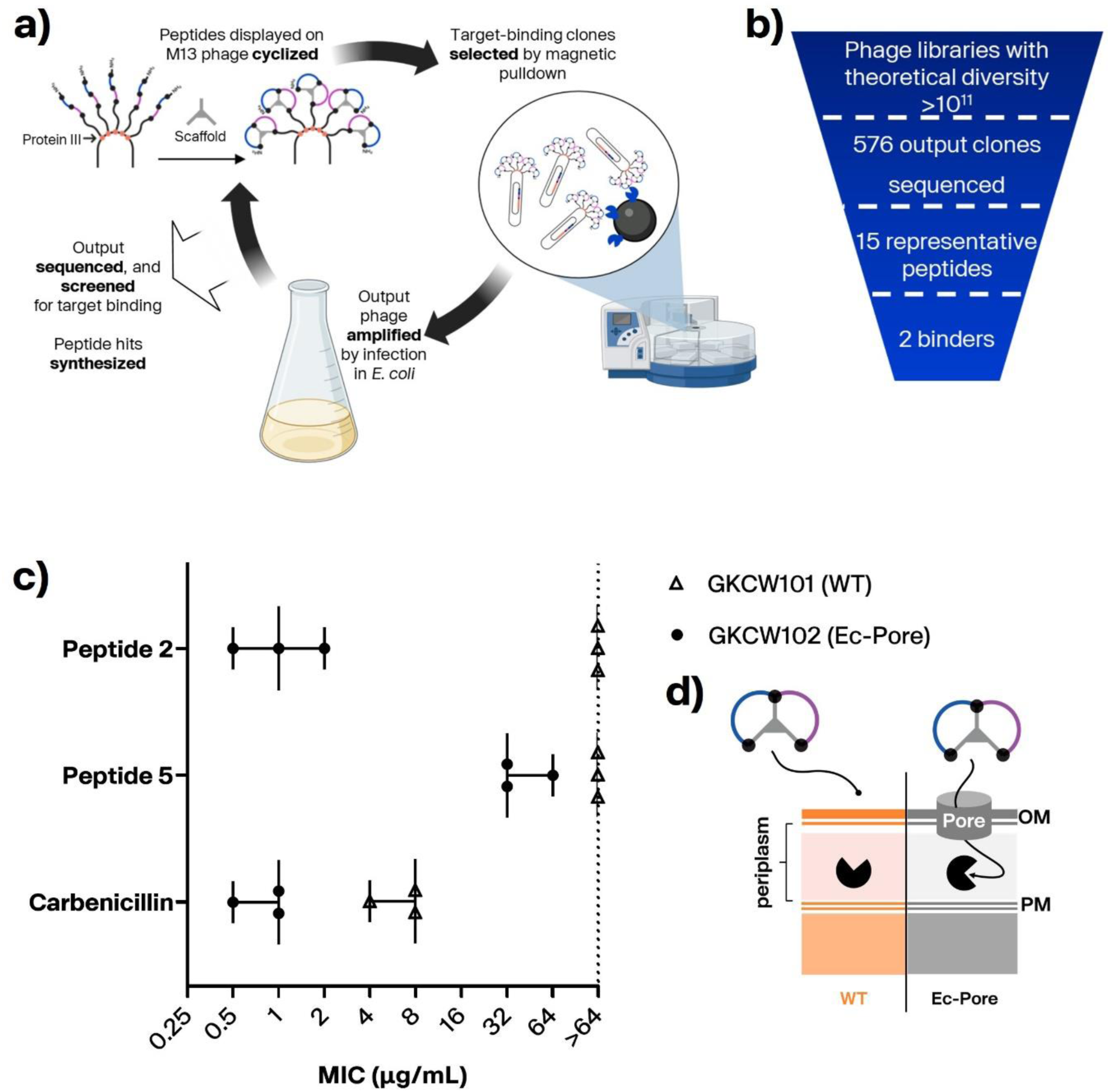
Identification and whole cell screening of *Ec*PBP3-binding bicyclic peptides. a) A modified phage display process to identify Bicycle binders of target proteins.^10^ Created with BioRender.com b) The output of phage selections was sampled by Sanger sequencing and representative peptides were chosen for chemical synthesis based on binding in a bead-based luminescence assay. From these synthetic peptides, 2 were confirmed as binders by SPR. c) An *E. coli* strain expressing a constitutively open FhuA pore (FhuAΔC/Δ4L) (d) was used to screen for whole cell activity of peptides, which are expected to required addition of a moiety driving permeation. Points represent biological replicate data; data are plotted with median and range. d) Schematic of hyperporinated strain used to screen whole cell activity of peptides.^28^

The large surface area for binding aligns well with prokaryotic target biology, where known inhibitors are frequently derived from natural products which are often larger, more complex structures compared to traditional small molecules.^20^ This large surface area also allows Bicycle molecules to be developed with exquisite target selectivity.^21^ Finally, Bicycle molecules are fully synthetic and tuneable using simple peptide chemistry. Chemical optimisation of binding and pharmacokinetic properties is simplified compared to natural product antimicrobials, which are synthetically challenging and usually only modifiable at a few positions.^22,23^ In many cases, the structural complexity of natural products limits their chemical tractability and may therefore require biosynthetic engineering or semi-synthesis, thus limiting the extent to which comprehensive structure-activity relationship (SAR) information can be determined.^24^

One disadvantage of the chemical nature of bicyclic peptides is that they are excluded by the outer membrane barrier (with an average size of ∼2000 Da compared to the ∼600 Da exclusion limit of outer membrane (OM) pores in Gram-negative bacteria).^25^ Therefore, we have explored strategies to deliver the peptide into the periplasm and have previously reported an approach for screening antimicrobial peptides as ‘vectors’ to confer OM penetrance to bicyclic peptides.^26^

Here, we describe the use of the proprietary Bicycle phage display discovery platform to identify and optimise novel, bicyclic peptide inhibitors of *E. coli* PBP3 (*Ec*PBP3), and their conjugation to linear antimicrobial peptides to confer outer membrane permeation.

## Results

### High-affinity bicyclic peptide inhibitors of *Ec*PBP3 were rapidly identified using a proprietary phage display platform

Bicyclic peptide phage libraries express a diverse array of linear peptides which are then covalently cyclised with a three-fold symmetrical small molecule scaffold via 3 cysteine residues, whose variable placement defines the loop sizes (i.e. the peptide ‘format’).^10^ The variable amino acid loop lengths, scaffold moieties and amino acid residues all contribute to the large diversity within Bicycle molecule libraries.

N-terminally truncated *Ec*PBP3 was panned against 28 peptide format libraries on the 1,3,5-triacryoyl-1,3,5-triazinane (TATA) scaffold, in 6 mixes (Table S-1).^10^ After 4 rounds of selection, 576 output clones were triaged by sequencing and screening to identify 15 representative hit sequences for chemical synthesis by solid phase peptide synthesis (SPPS) and further characterisation (Table S-2).

Binding of output peptides to *Ec*PBP3 was evaluated using a fluorescence polarisation (FP) assay. Initial screening was performed using BOCILLIN-FL, a fluorescently modified penicillin analogue, as a “tracer”, to triage peptides based on binding at the transpeptidase active site in a competition format assay.^27^ Of the 15 cyclised peptides tested, 3 (peptides **2**, **5** and **13**) showed binding to *Ec*PBP3 as indicated by competition of BOCILLIN-FL binding (Table S-2).

Staining of PBPs extracted from bacterial membranes with Bocillin FL was used to determine the binding of peptide **2** (and its all-*D* analogue) to other PBPs in *E. coli*.^8^ This method showed that peptide **2** bound only to the active site of *E. coli* PBP3 (Fig. S-2).

Peptides **2** and **5** were assayed by SPR in an EcPBP3 binding assay, with their K_d_ reported as 5.23 and 369 nM, respectively (Table S-2).

Peptide **2** was synthesised with fluorescein at the N or C terminus for use as a tracer in the FP competition assay.^16,17,19^ In direct FP binding experiments using fluorescent derivatives of peptide **2**, the binding affinity was dependent upon the position of the fluorophore, with micromolar binding from the C-terminally fluoresceinated clone compared to low nanomolar binding by the N-terminal fluoresceinated peptide (tracer **1**, K_d_: 6.86nM –Table S-3, Fig. S-1).

### Peptide 2 shows antimicrobial activity in *E. coli* strains with permeabilised outer membranes

To evaluate the engagement of *Ec*PBP3 by a bicyclic peptide in whole cells lacking the outer membrane permeability barrier, peptides were screened in ‘hyperporinated’ *E. coli* strains in which a constitutively open FhuA pore (FhuAΔC/Δ4L, lacking N-terminal cork domain and external loops) was chromosomally expressed under control of an arabinose-inducible promoter.^28^ No bacterial growth inhibition was observed in the control strain (GKCW101, transformed with an empty expression cassette), but peptide **2** had an MIC of 0.5-2 μg/ml against GKCW102 (expressing FhuAΔC/Δ4L), demonstrating that the peptide was able to engage *Ec*PBP3 in the whole cell when access to the periplasm was achieved (Fig. 2c). The weaker binder, peptide **5**, showed limited activity in the hyperporinated strain.

### Peptide 2 forms interactions with both precedented and previously unengaged pockets of the transpeptidase catalytic cleft

A crystal structure of peptide **2** in complex with the transpeptidase domain of *Ec*PBP3 (Table S-4 and Fig. 3; Protein Data Bank (PDB) code: 8RTZ) was solved at high resolution (1.5 Å). The Bicycle molecule binds in the active site cleft, in a region previously demonstrated as the binding site for β-lactams such as piperacillin and AIC499, a novel monobactam.^29,30^ The buried surface area of the peptide bound to the protein is approximately double that of the buried surface area of the protein reacted with piperacillin (1832 Å^2^ and 951 Å^2^ respectively). The bound Bicycle molecule extends into a region of the active site unreached by the sidechains of β-lactams (Fig. 3a, b). Compared to *apo* structures of *Ec*PBP3, the backbone conformation of the bound *Ec*PBP3 is largely unchanged despite the presence of the large ligand (RMSD of 0.357 Å; Fig. S-3).^29^ The peptide adopts a rough “Figure-of-8” conformation within the active site, with the N- and C-terminus of the peptide linked via two arms of the scaffold and completing the “8” (Fig. 3). Using surface filling models to examine the interactions between the protein and ligand shows that several of the residues (i.e. Phe2 and Pro5-Trp6-Val7-Glu8 and TATA scaffold) are in tight complex with the protein, close to the minimal distance between the protein and ligand (Fig. S-4).

**Figure 3:**
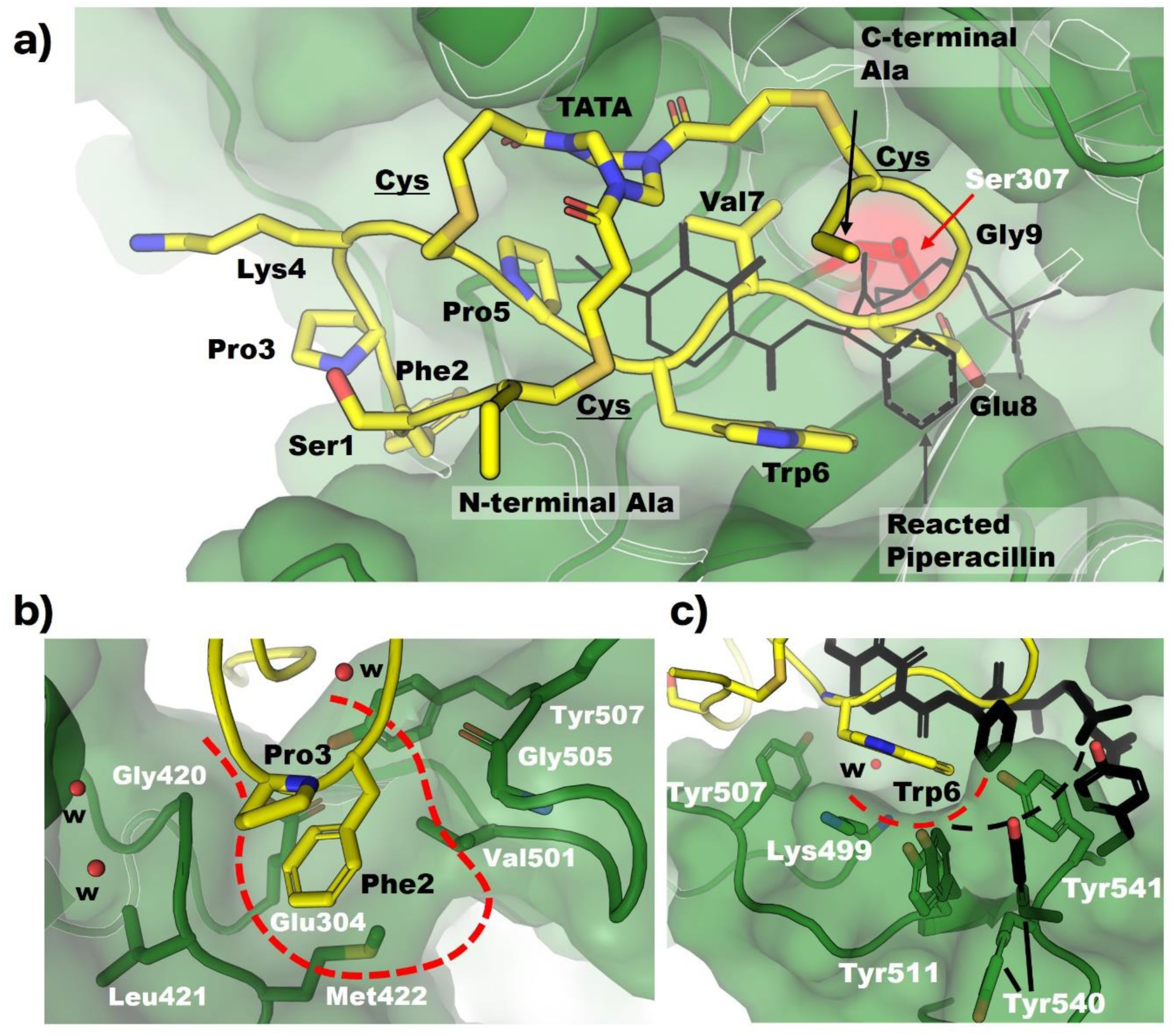
Overview of the interaction of EcPBP3 and peptide 2 (PDB code: 8RTZ). *Ec*PBP3 (green) is in surface and cartoon representation; Peptide 2 is shown in yellow in cartoon representation, with residue side chains in stick. The variable residues of the peptide are numbered from N to C terminus whilst the invariant cysteines are underlined. The scaffold molecule: TATA, conjugated to each of the cysteines is labelled. The “catalytic” serine is highlighted in red. A molecule of PBP3-reacted piperacillin (PDB code: 6I1I) is shown in a) and c)).^29^ The Bicycle occupies many of the same active site regions as piperacillin and extends further into a pocket not previously observed to be occupied with ligand (b). Residues forming the “hydrophobic wall” (dashed lines) are shown in (c). The residues’ positions in the piperacillin-reacted structure (black sticks) and the Bicycle complex (green sticks and surface) are shown.

The electron density for the peptide residues and the scaffold moiety is largely complete, with the exception of the lysine sidechain and N-terminal residues (Fig. S-5). Similarly, electron density for important catalytic residues of the active site, including Ser307 (the catalytic serine) is complete. Ser307 is observed to adopt two conformations: with the sidechain in two different rotamers (Fig. S-6). It is unclear why Ser307 adopts these two conformations. The peptide is 5.3 Å away and only interacts with the conserved active site catalytic residues (i.e. Ser307, Lys 310, Ser359, Asn361, Lys494 and Thr495) via bridging waters (Fig. S-6).

Some elements of the peptide interaction with *Ec*PBP3 are similar to interactions observed in the structure of piperacillin reacted with *Ec*PBP3. Three essentially equivalent hydrogen bonds are seen in the two structures (Fig. S-7). The backbone of the peptide overlaps with the position of the central peptidic portion (particularly the diketopiperazine and phenylglycine groups) of reacted piperacillin. Two of the conserved hydrogen bonds (labelled **A** and **C** in Fig. S-7) are formed from this central region. The third hydrogen bond is formed between Thr497 and an acidic group (Glu8 in the Bicycle and carboxylic acid from piperacillin). Whilst this acidic group is well conserved amongst β-lactams, this hydrogen bond does not seem essential for the interaction as modification of Glu8 in the peptide to alanine did not affect the binding affinity (Fig. 4). A “hydrophobic wall” (comprising Tyr541, Tyr511 and Lys499 of *Ec*PBP3) was also observed to form around Trp6 of the peptide, as described in the interaction of reacted aztreonam with *P. aeruginosa* PBP3 (Fig. 3c).^31^ *Ec*PBP3 reacted with piperacillin also forms a hydrophobic wall (composed of Tyr511, Tyr540 and Tyr541), but the centre of this formation is shifted compared to the one observed in the complex with the Bicycle molecule (Fig. 3).^29^ Key interactions of the amino acid side chains of the peptide with *Ec*PBP3 are summarised in Fig. 4d.

**Figure 4:**
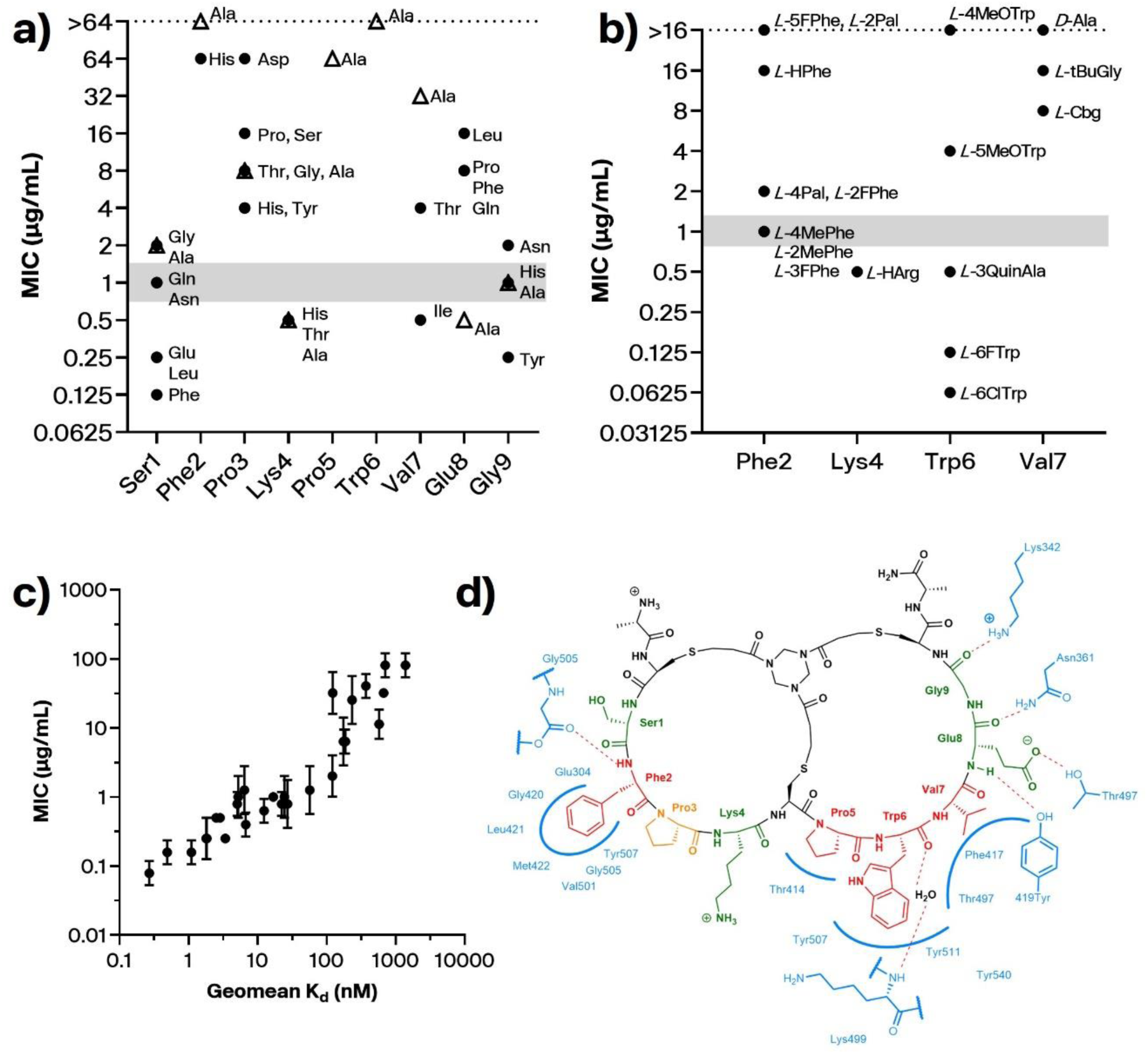
Summary of amino acid tolerance and residue essentiality for Peptide 2, with key peptide-protein interactions from the crystal structure. Single residue library selections, alanine scanning and medicinal chemistry were used to evaluate the tolerance for substitution at each position, and rationalised using structural biology. Whole-cell potency of peptides was determined in MIC against *E. coli* GKCW102 for peptides with a) natural amino acid substitutions or b) non-natural substitutions in peptide 2. Points represent the median of 3 biological replicates. Grey bars indicate median MIC for peptide 2. Binding data are provided in Extended data; full data are provided in the Supplementary materials. Triangles, alanine scan peptides; circles, all other substitutions. c) Binding affinity (SPR) against MIC for peptide series derived from peptide 2. Kd values presented as geometric mean of at least 2 biological replicates. MICs were determined against E. coli GKCW102. Values are presented as median and range and were derived from 3 biological replicates. Full data provided in Table S-7. d) Representation of the binding interaction with EcPBP3, as per the crystal structure. Residues essential for binding are shown in red, those of moderate importance in orange and those which can be mostly freely substituted, green.

### Structure-rationalised modifications of peptide 2 resulted in improved affinity and whole cell activity in porinated strains

Concurrent with generating the co-crystal structure, a variety of approaches were used to reveal the structure-activity relationship (SAR) of the interaction of peptide **2** with *Ec*PBP3. We first used the affinity-based selection of the phage platform in SAR selections (Fig. 4a). Here, each residue in the peptide was individually randomised to probe the tolerance (based on the sequences recovered in the selection output) for natural amino acid substitutions at each position in the Bicycle molecule. The use of the phage display platform to reveal the amino acid tolerance at each residue allowed us to rapidly triage the 171 different sequences covering this set, therefore, guiding the synthesis of a reduced panel of peptides for affinity determination and generating comprehensive SAR without the need for iterative rounds of peptide synthesis. Similarly, we performed an alanine scan of peptide **2**, in which each amino acid in turn was replaced by an alanine to define the core binding motif (Fig. 4a). The core binding motif and tractable residues revealed by SAR selections and alanine scanning was subsequently rationalised by the co-crystal structure (Fig. 4d).

Certain key residues in the binding interaction showed little tolerance for amino acid substitution (Fig. 4a). There was good agreement between the SAR library and alanine scan techniques. Pro5 and Trp6 were essentially invariant when considering substitution by natural amino acids, as those identified in the SAR selection output were associated with at least a 100-fold reduction in binding affinity in FP competition (Table S-7). Identification of these substitutions in the SAR selection was likely due to detection of weak binding interactions due to the avidity introduced by multivalent display at the phage surface.^32,33^

Similarly, Phe2 did not tolerate substitution by other natural amino acids. These data were in agreement with structural data showing that the engagement of Trp6 and Phe2 are key for the interaction of peptide **2** with *Ec*PBP3. By contrast the residues Ser1, Lys4, Glu8 and Gly9, which do not interact with *Ec*PBP3, showed tolerance for substitution by amino acids with distinct properties from the original residue. When screened in the FP competition assay, both the parent peptide (**2**) and multiple derivatives at Ser1, Lys4, Glu8 and Gly9 were reported with low nanomolar affinities that were below the sensitivity limit of the assay (as imposed by the protein concentration, 9.4 nM). We therefore transferred to SPR for screening of further peptides – Extended Data Fig. 1b includes SPR data for example peptides tested by FP, such as Ser1Leu (**27**, 3.36 nM – Table S-7) and Ser1Phe (**28**, 1.12 nM)

Having defined the core binding motif via SAR selections and alanine scanning and subsequently rationalised these data via structural biology, we next extended the series by incorporating non-natural amino acids at key positions in the peptide (Fig. 4b). Phe2, Trp6 and Val7 were modified to pursue increased binding affinity, whilst Lys4 was modified to increase proteolytic stability.

Trp6 of the peptide interacts with a hydrophobic wall comprising Tyr541, Tyr511 and Lys499 of *Ec*PBP3 (Fig. 3c), in a similar fashion to the interactions of the *gem*-dimethyl group of aztreonam with Tyr503, Tyr532 and Phe533 of *P. aeruginosa* PBP3.^31^ More than 10 fold improvement in binding affinity was achieved by further hydrophobic extension of Trp6 by incorporation of 6ClTrp (**65**, 0.269 nM) or 6FTrp (**66**, 0.489 nM) – these substitutions were also associated with the best potency in hyperporinated strain MICs (Fig. 4b).

Loop 1 of the peptide extends beyond the binding footprint of β-lactams within the transpeptidase catalytic site of *Ec*PBP3, with Phe2 binding a previously unengaged pocket of *Ec*PBP3 created by Glu304, Met422 and Val501 of the protein (Fig. 3b). Modifications to the electronics of the phenyl ring were variably tolerated, with potency maintained for the switch to 4MePhe but lost with 5FPhe (Fig. 4b, Fig. S-7).

Val7 of the peptide tolerated substitution by isoleucine and threonine only in the natural amino acid substitutions (Fig. 4a). Val7 participates in intramolecular interactions with the TATA scaffold and C-terminal cysteine, making a significant contribution to the positioning of the C-terminal half of the peptide. The included non-natural substitutions to Val7 were poorly tolerated, likely as a consequence of the small pocket occupied by this residue which allowed limited space for further modification. The solvent-facing Lys4 residue was replaced with HArg (peptide **25**) to offer improvement in the proteolytic stability of the peptide – this substitution was well tolerated with no loss in binding or MIC (Fig. 4b, Fig. S-7).

To evaluate whether *in vitro* binding affinity of peptides in the series was a predictor of target engagement in the whole cell context, binding affinity was plotted against hyperporinated strain MIC (Fig. 4c). This analysis demonstrated that increases in affinity for *Ec*PBP3 were associated with greater potency against the hyperporinated strain, indicating that binding affinity was a good predictor of target engagement *in vivo*.

### Whole cell activity against wild-type *E. coli* conferred by a vector conjugation strategy

With a potent binder of recombinant *Ec*PBP3 protein in hand and having demonstrated that bicyclic peptides could engage *Ec*PBP3 in the whole cell context, we required a mechanism to confer outer membrane permeation of the peptides for cell killing in wild type bacteria. A split luciferase assay for outer membrane uptake (SLALOM) was used to identify antimicrobial peptides from the literature which could act as a vector for periplasm access of a luciferase fragment.^26^ A promising peptide from this work, DRAMP18563 (named for the entry in the Data Repository for AntiMicrobial Peptides), was shown to confer potent wild type activity against *E. coli* strains when conjugated to peptide **2** (conjugate **1**) or a lipid-II targeting peptide.^26,34^

Activity against wild type bacteria was maintained in a conjugate in which the DRAMP18563 ‘vector’ peptide was switched to the retroinverso format (i.e. all-*D* amino acids, in the inverse sequence – conjugate **2**). The modification of the vector also conferred improved plasma stability to the conjugate molecule (Table S-9). To demonstrate the contribution of PBP3 inhibition to whole-cell activity of the conjugate molecules, we synthesised a conjugate in which the bicyclic peptide ‘warhead’ comprised all-*D* amino acids, which showed no binding to *Ec*PBP3 (conjugate **3**). Conjugate **3** was 32-fold weaker in MIC, indicating that the activity of the peptide-vector conjugates was driven by inhibition of *Ec*PBP3, and relied upon vector-conferred periplasm access. Representative conjugate molecules were not associated with toxicity to mammalian cells when screened in haemolysis and mammalian cell cytotoxicity assays up to 50 mM (Table 1 and S-9).

**Table 1:**
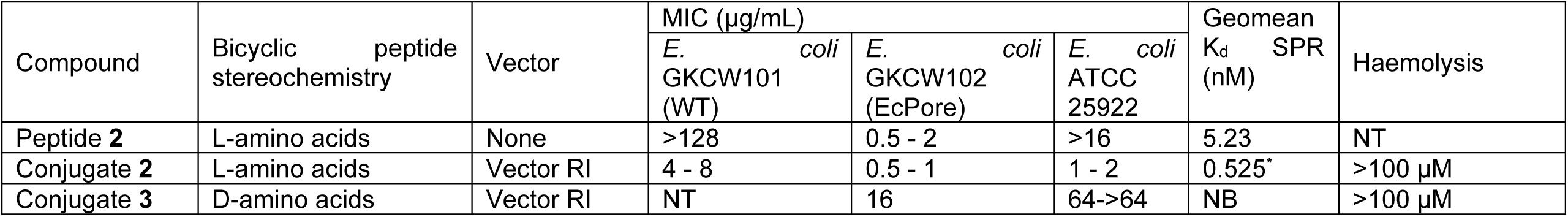
MIC and haemolysis of peptides and conjugates against *E. coli*. MIC values are representative of the range in n≥3 assays. MICs were determined in caMHB for hyperporinated strains, and in MHB for ATCC 25922. NB, no binding (top concentration tested 5000 nM); NT, not tested; RI, retro-inverso. * n=1. Poor fit observed in SPR, likely due to conjugation to charged peptide.

| Compound | Bicyclic peptide stereochemistry | Vector | MIC (µg/mL) |  |  | Geomean K<sub>d</sub> SPR (nM) | Haemolysis |
| --- | --- | --- | --- | --- | --- | --- | --- |
|  |  |  | *E. coli* GKCW101 (WT) | *E. coli* GKCW102 (EcPore) | *E. coli* ATCC 25922 |  |  |
| Peptide **2** | L-amino acids | None | >128 | 0.5 - 2 | >16 | 5.23 | NT |
| Conjugate **2** | L-amino acids | Vector RI | 4 - 8 | 0.5 - 1 | 1 - 2 | 0.525* | >100 µM |
| Conjugate **3** | D-amino acids | Vector RI | NT | 16 | 64->64 | NB | >100 µM |

### Capping and amino acid substitution improved *in vitro* stability without loss of potency

We next looked to evaluate the *in vitro* stability of the series. To improve blood stability of the bicyclic peptide, certain precedented substitutions and modifications were made to the sequence. We have described substitution of the protease-labile Lys4 above; the exocyclic alanines were removed; and the amino terminus of the peptide was capped (Fig. 5a).

**Figure 5:**
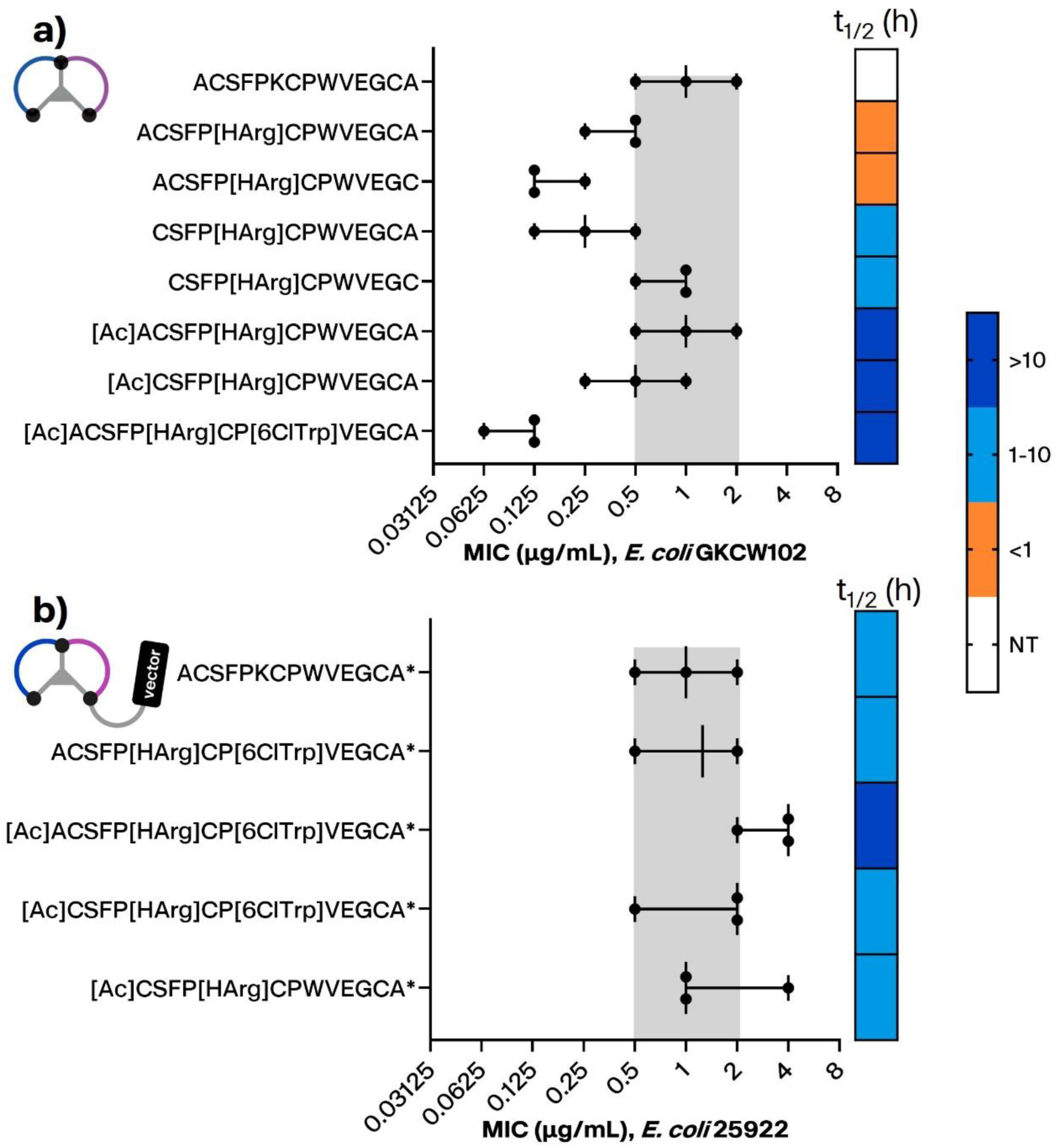
MIC and *in vitro* stability of modified *Bicycles* (a) and conjugates (b). (a) Potency of *Bicycles* against the hyperporinated *E. coli* strain GKCW102 was evaluated alongside stability in mouse blood. The parent peptide (2) is presented for comparison in MIC, but was not tested in stability assays. Corresponding peptide numbers are (top to bottom) **2**, **25**, **70**, **71**, **72**, **73**, **74**, **75**. (b) Potency of conjugates against *E. coli* ATCC25922 was evaluated alongside stability in mouse blood. Corresponding conjugate numbers are (top to bottom) **2**, **4**, **5**, **6**, **7**. Grey shaded area indicates the potency range of the parent compound (Peptide **2** in (a), Conjugate **2** in (b)). Each point represents a biological replicate measure, error bars show the range and median.

A range of substitutions were tolerated for activity in the hyperporinated strain, and certain key substitutions amongst these were associated with improved stability in mouse blood. Capping at the N-terminus appeared to be a key modification conferring improved stability, as peptides incorporating an N-terminal cap exhibited half-lives >10 h in mouse blood (peptides **73**-**75**), compared to <1 h when the N-terminal alanine was unmodified (peptides **25** and **70**; Fig. 5a). A promising combination of potency and stability was observed with peptide **75**, in which the affinity-boosting 6ClTrp was introduced to the stabilized peptide coupled with N-terminal acetylation and HArg4. A subset of peptides with the tolerated substitutions were taken forward for synthesis as DRAMP conjugates.

Stabilised peptide conjugates with the DRAMP-RI vector were tested in MIC assays against a wild-type *E. coli* strain, and evaluated for mouse blood stability (Fig. 5b). Surprisingly, the unmodified bicyclic peptide as a conjugate showed *in vitro* stability in range with the capped and substituted compounds, suggesting that attachment of the DRAMP-RI vector improved stability regardless of the stability of the peptide alone. N-terminal capping of the peptide appeared to have minimal impact on the *in vitro* stability of the conjugates.

### DRAMP conjugates show an encouraging spectrum of activity against strains of the Enterobacterales

To evaluate the spectrum of activity of the conjugate series, we further tested exemplar conjugates in a representative strain set of Enterobacterales, along with additional Gram negative pathogens. Encouragingly, the activities of conjugates in this series were maintained in accordance with the sequence conservation of the targeted gene product of *ftsI*, with activity in a similar range to that of *E. coli* across members of the closely related Enterobacterales and lost when sequence homology of *ftsI* dropped below 94% (Extended Data Table 1). The correspondence of conjugate **2** activity with *ftsI* sequence conservation further supported the mode-of-action of the anti-infective conjugate via PBP inhibition.

## Discussion

Efforts in the development of PBP inhibitors in recent years have encompassed further exploration of the β-lactam class, particularly with respect to β-lactamase resistance; structure-guided design of PBP inhibitors from the diazabicyclo-octane (DBO) β-lactamase inhibitors; and screening of protease-targeted compound libraries – readers are directed to Bertonha *et al.* for a review of recently developed PBP inhibitors.^35^ Notably, a key aspect of many of these efforts has been to address OM uptake. An example β-lactam where OM uptake has been optimized is cefiderocol, a cephalosporin-class β-lactam whose OM permeation is enhanced by a catechol siderophore moiety.^36–38^ Structure-guided optimization has been used to generate a DBO compound (ETX0462) whose activity includes inhibition of both multiple classes of β-lactamases and PBP1a/PBP3.^39^ A key stage in the development of this compound was the enhancement of access through porins based on structure-porin permeation relationships, thus improving upon the microbiological profile of an earlier compound.^39^ The pyrrolidine-2,3-diones described by Lopez-Perez *et al.* showed binding and inhibition of *P. aeruginosa* PBP3, but required outer membrane permeabilization (by screening in the presence of polymyxin B nonapeptide) for access of the compounds to the target.^40^ Altogether, these compounds illustrate both the promise of new chemical matter against an established target, and the known challenge of access to the Gram negative bacterial cell.

Here we have used bicyclic peptide phage display to identify novel, highly selective inhibitors of *Ec*PBP3. The compounds described here are functional inhibitors of septal peptidoglycan biosynthesis and were optimised to low nanomolar affinities; showed potent activity in MICs; and represent a novel non β-lactam based PBP chemotype. Phage display offers a rapid and high-throughput approach to antimicrobial discovery, allowing significant flexibility in exploring SAR whilst screening a target precedented by marketed antimicrobials. This addresses a key challenge in prior application of high-throughput screening approaches to antimicrobial discovery, namely the need for new approaches exploiting appropriate chemical space for potential antimicrobials.

We previously described a split luciferase assay technique (SLALOM) for identifying peptide sequences that could act as ‘vectors’ for bicyclic peptides.^26^ This technique was used to identify DRAMP18563, which when conjugated to peptide **2** showed whole cell activity against *E. coli* and related organisms of the Enterobacterales. The antimicrobial activity of this conjugate was through inhibition of PBP3, as demonstrated by the lack of activity from a conjugate containing a non-PBP3 binding peptide. Certain challenges are known to be associated with antimicrobial peptides, as in some instances these compounds exert non-specific toxic effects upon both bacterial and mammalian cells.^41^ However, we have not observed *in vitro* toxicity associated with our molecules at concentrations greater than 80-fold the active concentrations in MIC (Table 1, S-9).

In structural studies, peptide **2** was shown to bind across the catalytic cleft of *Ec*PBP3 where it makes key interactions with both precedented binding pockets (i.e., the hydrophobic wall engaged by Trp6 and the hydrogen bond network near the active site) (Fig. 3c), and previously unengaged sites in the cleft such as the binding pocket of Phe2 (Fig. 3b). The importance of these interactions for binding was further supported by alanine scanning and SAR selections, where mutation of Trp6 or Phe2 resulted in a loss of binding. The large surface area of peptide **2** allows a broad engagement across the span of the transpeptidase catalytic cleft of *Ec*PBP3. As novel, non-covalent inhibitors of *Ec*PBP3 the series offers a unique tool to probe the function of this enzyme, and to reveal new tractable sites for the inhibition of *Ec*PBP3.

These data provide the basis for the development of anti-infective *Bicycle* molecule conjugates as novel antimicrobial drugs. Critical further work to support this ambition will be to optimize pharmacokinetics properties and investigate activity in *in vivo* models of infection as well as determining their propensity to induce resistance. Mutations in *Ec*PBP3, in clinical strains, typically arise at a loop distant from the transpeptidase catalytic site and are associated with decreased susceptibility to β-lactams including aztreonam.^42,43^

In summary, the novel antimicrobials described here represent the first in a new class of non-covalent penicillin binding protein inhibitors and exemplify the potential of the Bicycle phage display platform for discovery of new antimicrobial compounds.

## Online methods

### Protein production and crystallography

Two *Ec*PBP3 protein constructs were used. For assays, selections and SPR a construct with residues 60−588 which has the N-terminal domain, but not the transmembrane helix, was used as previously described.^44^

Owing to the difficulty of crystallising this N-terminal domain containing construct, an expression construct consisting of *E. coli* PBP3, transpeptidase (TP) only domain, residues 234-572, Δ280-294 was cloned by Vector Builder into a pET-47b(+) vector that contained an N-terminal 6-His tag followed by a human rhinovirus (HRV) 3C protease tag. The protein was expressed and purified as previously described by Bellini *et al.* (2019) into a final buffer: 10 mM Tris-HCl, pH8, 500 mM NaCl.^45^

The following non-native residues: ‘GPGYQDP’ remain at the NTD of the final expressed protein due to the restriction cloning used.

For co-crystallisation with peptide, *Ec*PBP3 protein (20 mg/ml) in 10mM Tris HCL pH 8.0, 500 nM NaCl and peptide was incubated at a molar ratio of 1:1.5 protein to peptide on ice for three hours. The complex was spun at 15,000 xg to remove any precipitate and the supernatant used in subsequent crystallisation experiments. The best crystals were obtained from drops consisting of 2 µl protein and 1 µl of well solution. Crystals from the following condition were harvested for data collection: 0.1M Tris pH 9.0 and 20 % (w/v) PEG 6K supplemented with an additive, 30 % (w/v) dextran sulphate sodium salt, Mr 5K. Crystals were frozen for data collection with the addition of 20 % glycerol.

Buried surface area was calculated using a Pymol (v 2.4.1,Schrodinger LLC) script. The script finds the solvent-accessible surface area of the individual components (i.e. protein with Bicycle molecule removed (P), Bicycle molecule with protein removed (B) and the Bicycle molecule-protein complex (C)), then calculated the buried surface area as: P+B-C, and equivalent for the piperacillin-reacted complex.

Figures generated in Pymol (v 2.4.1,Schrodinger LLC) unless stated.

### Antimicrobial assays

Minimal inhibition concentration (MIC) assays were performed in accordance with CLSI standards, using cation-adjusted Mueller Hinton broth (caMHB; Sigma Aldrich) unless otherwise specified. Induction of pore expression in hyperporinated strains *E. coli* GKCW101 and GKCW102 was performed as previously described, with caMHB as the culture medium.^28^ MIC data are presented from at least two biological replicates unless otherwise indicated in figure legends.

### SPR assays

Surface plasmon resonance (SPR) assays were performed using *Ec*PBP3 immobilized on CM5 sensor chips (GE Healthcare) via amine-coupling chemistry in assay buffer (10 mM HEPES pH 8.0, 200 mM NaCl, 0.05 % (v/v) surfactant P20). The chip surface was activated by injection (10 μL/min) of 0.4 M EDC and 0.1 M NHS (in a 1:1 ratio) with a 7 min contact time. Protein was diluted to 25 μg/mL in 10mM MOPS (pH 7.0) and injected over the activated sensor chip surface to reach immobilization of 1500-2500 response units. Excess hydroxysuccinimidyl groups on the chip surface were deactivated by injection (10 μL/min) of 1M ethanolamine hydrochloride pH 8.0 with a 7 min contact time. Reference flow cells were prepared as described, but without ligand (*Ec*PBP3 protein).

Binding analysis runs were performed at 37°C in running buffer (10 mM HEPES pH 8.0, 200 mM NaCl, 0.05 % (w/v) Tween-20, 2 % (v/v) DMSO). Test compounds (and control runs of buffer only) were injected in running buffer over the *Ec*PBP3 chip and reference flow cell, at flow rate of 50 μL/min for 150 second contact time and 2400 second dissociation time. Each injection was followed by a wash with 50 % (v/v) DMSO. All compounds were tested in at least two independent biological replicates unless otherwise stated.

#### Data Analysis

Raw sensorgrams data were solvent-corrected, reference-subtracted and blank-buffer-subtracted (double-referencing) before kinetic and affinity analysis to account for non-specific binding and injection artifacts. Equilibrium dissociation constant (K_d_) values was determined using the Biacore™ Insight Evaluation Software (version 4.0.8 or version 5.0.17) using kinetic fitting. K_d_ values presented are the geometric mean of at least two biological replicates.

### Mouse blood and plasma stability

Stability assays were performed using freshly collected matrices from CD-1 mice, with anticoagulant as stated per dataset. Compound stocks in DMSO were tested at 5μM final concentration (0.5% (v/v) DMSO). Compounds were incubated in blood or plasma at 37°C for 24h, with sampling at 0, 1, 2, 4, 6 and 24h. Samples were prepared via protein precipitation using organic solvent containing internal standard, and analyzed by LC-MS/MS. Compound remaining was quantified based on the ratio of analyte to internal standard, and the half-life (t_1/2_) was calculated from a log linear plot of % remaining versus time.

### Chemistry

Compounds were isolated at >95 % purity, as assessed by high performance liquid chromatography (HPLC) (apart from screening peptides which were greater than 80% pure) – analytical traces can be found in the supporting information.

#### General Peptide Synthesis

Bicycle peptides were synthesized as previously described.^17^ Briefly, Bicycle peptides were synthesized on Rink amide resin using standard 9-fluorenylmethyloxycarbonyl (Fmoc) solid-phase peptide synthesis, either by manual coupling (for large scale) or using a Biotage SyroII automated peptide synthesizer (for small scale). Following trifluoroacetic acid based cleavage from the resin, peptides were precipitated with diethyl ether and dissolved in 50:50 acetonitrile/water. The crude peptides (at ∼1 mM concentration) were then cyclized with 1.3 equiv of TATA scaffold, using ammonium bicarbonate (100 mM) as a base. Completion of cyclization was determined by matrix-assisted laser desorption ionization time-of-flight (MALDI-TOF) or liquid chromatography-mass spectrometry (LC–MS). Once complete, the cyclization reaction was quenched using *N*-acetyl cysteine (10 equiv with respect to TATA), and the solutions were lyophilized. The residue was dissolved in an appropriate solvent and purified by reversed-phase (RP) HPLC. Peptide fractions of sufficient purity and the correct molecular weight (verified by either MALDI-TOF and HPLC or LC–MS) were pooled and lyophilized. Concentrations were determined by UV absorption using the extinction coefficient at 280 nm, which was based on Trp/Tyr content. Standard Fmoc amino acids, as well as nonproteinogenic Fmoc amino acids, were obtained from Sigma-Aldrich, Iris Biotech GmbH, Apollo Scientific, ChemImpex, and Fluorochem.

Fluorescent peptide synthesis and copper catalysed click conjugate were performed as previously described.^19,26^

## Supporting Information

General procedures; compound analytics; crystallography statistics; supporting crystallographic images; cytotoxicity data.

## Data availability

Crystallography structural data are deposited in the Protein Data Bank (PDB code: 8RTZ).

## Author contributions

The manuscript was written through contributions of all authors.

## Acknowledgements

These studies were funded by an SBRI award (Project no. 971626Q1), a Sustainability Fund Award (Project no. 77556) and a Biomedical Catalyst Award (Project no. 82975) from Innovate UK. H.N. is a UKRI-funded Innovation Scholars secondee between the University of Warwick and Bicycle Therapeutics (project reference: MR/W003554/1). SPR assays were established at Evotec (Verona) and subsequently performed at Bicycle Tx. X-ray crystallography was performed by Charles River Laboratories (Cambridge). Haemolysis studies were carried out at Evotec SE (Verona). Plasma stability studies were carried out at WuXi Apptec Co. Ltd. (Shanghai). Blood stability and broad panel MIC studies were carried out at Selvita d.o.o. (Zagreb).

We thank Prof. Helen Zgurskaya of University of Oklahoma for hyperporinated strains used in this work.

## Abbreviations

caMHB, cation-adjusted Mueller Hinton broth; Cbg, cyclobutylglycine; EcPBP3, *E. coli* PBP3; EDC, 1-ethyl-3-(3-dimethylaminopropyl) carbodiimide hydrochloride; FP, fluorescence polarisation; HArg, homoarginine; HPhe, homophenylalanine; MHB, Mueller Hinton broth; MIC, minimal inhibitory concentration; MOPS, 3-morpholin-4-ylpropane-1-sulfonic acid; NB, no binding; NHS, N-hydroxy succinimide; NT, not tested; SPR, surface plasmon resonance; TATA, 1,3,5-triacryoyl-1,3,5-triazinane; tBuGly, tert-butyl-leucine; 2FPhe, 2-fluoro-phenylalanine; 2MePhe, 2-methyl-phenylalanine; 2Pal, 2-pyridylalanine; 3FPhe, 3-fluoro-phenylalanine; 3QuinAla, 3-Quinoyl-L-alanine; 4MeOTrp, 4-methoxytryptophan; 4MePhe, 4-methyl-phenylalanine; 4Pal, 4-pyridylalanine; 5FPhe, pentafluoro-phenylalanine; 5MeOTrp, 5-methoxytryptophan; 6ClTrp, 6-chlorotryptophan; 6FlTrp, 6-fluorotryptophan.

## Extended Data

**Extended Data Figure 1:**
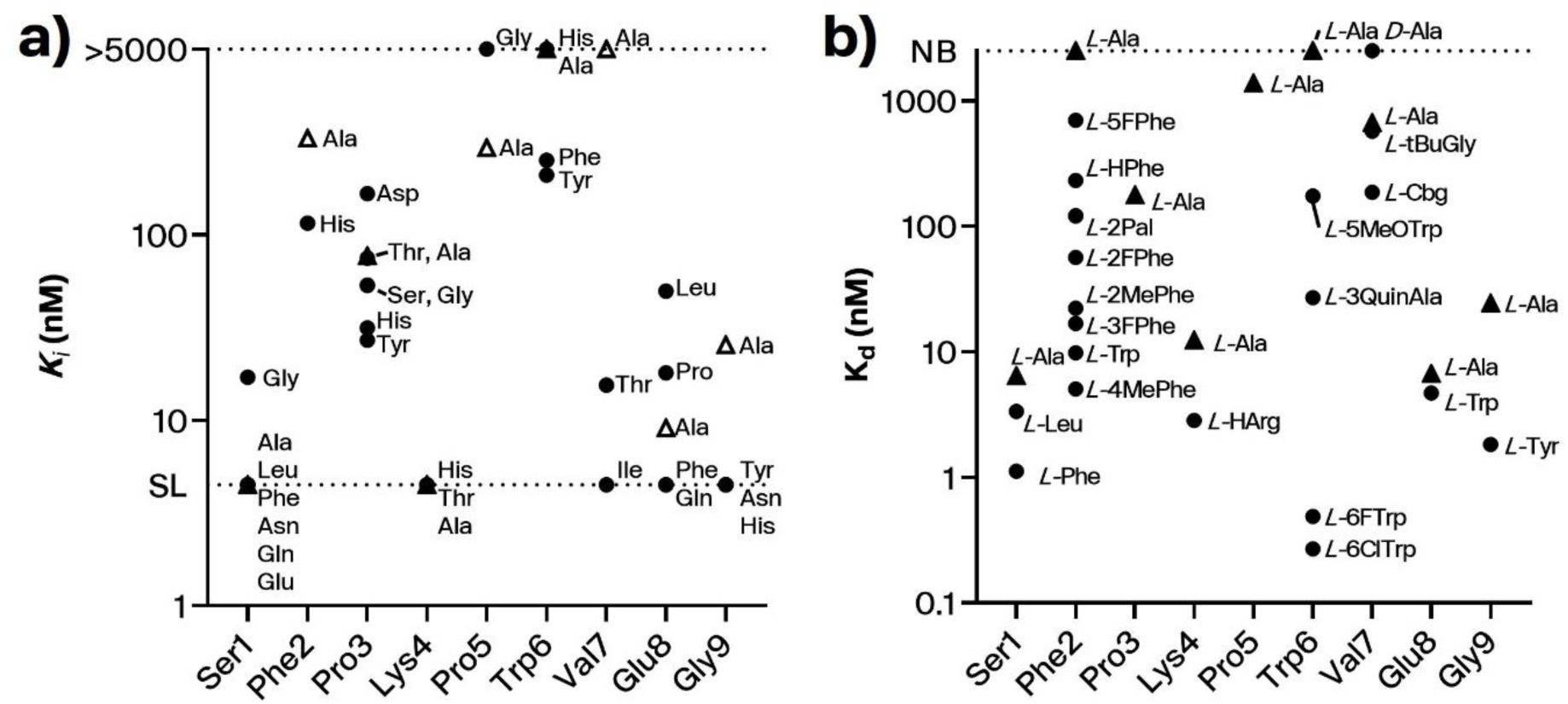
Tolerance for amino acid substitution in Peptide 2 as characterized by binding of *Ec*PBP3. Single residue library selections, alanine scanning and medicinal chemistry were used to evaluate the tolerance for substitution at each position, and rationalised using structural biology. Binding of peptides to *Ec*PBP3 was evaluated in a) FP competition assays using tracer **1** and b) SPR assays. Points represent the geometric mean of at least 2 replicates. Triangles, alanine scan peptides; circles, all other substitutions. SL, sensitivity limit of assay.

**Extended Data Table 1:** *Ec*PBP3-targeted bicyclic peptide-vector conjugates had promising spectrum of activity against related strains of the Enterobacterales. Antimicrobial activity of Conjugate 1 (**a**) and Conjugate 2 (**b**) in MIC assays against strains of the Enterobacterales order (*E. coli*, *C. freundii, K. pneumoniae, E. cloacae, P. mirabilis*) as well as two additional high-priority Gram negative pathogens, *A. baumannii* and *P. aeruginosa*. MICs were generated in caMHB medium, and number of replicates are given in brackets.

| Organism | % identity of *ftsI* gene product with *E. coli* | Strain | MIC (µg/ml) |  |
| --- | --- | --- | --- | --- |
|  |  |  | Conjugate 1 | Conjugate 2 |
| *Escherichia coli* | 100 | *E. coli* ATCC 25922 | 2,4 (2) | 2,8 (2) |
|  |  | *E. coli* ATCC BAA-2469 | 32,32 (2) | 8,8 (2) |
|  |  | *E. coli* NCTC 13441 | 4,4 (2) | 4,8 (2) |
|  |  | *E. coli* UTI89 | 4,4 (2) | 8 (1) |
| *Citrobacter freundii* | 96 | *C. freundii* AR0116 | 8,8 (2) | 8 (1) |
|  |  | *C. freundii* B121 | 4,4 (2) | 8 (1) |
| *Klebsiella pneumoniae* | 94 | *K. pneumoniae* ATCC 43816 | 4,4 (2) | 8,16 (2) |
|  |  | *K. pneumoniae* ATCC 700603 | 4,4 (2) | 16 (1) |
|  |  | *K. pneumoniae* B454 | 4,8 (2) | 16,32 (2) |
|  |  | *K. pneumoniae* KPNIH1 | 8,8 (2) | 32,>64 (2) |
| *Enterobacter cloacae* | 94 | *E. cloacae* AR0032 | 2,4 (2) | 4 (1) |
|  |  | *E. cloacae* AR0038 | 2,2 (2) | 8 (1) |
|  |  | *E. cloacae* B173 | 4,4 (2) | 16 (1) |
|  |  | *E. cloacae* KPC114 | 4,4 (2) | 8 (1) |
| *Proteus mirabilis* | 75 | *P. mirabilis* AR159 | >64, >128 (2) | >64 (1) |
|  |  | *P. mirabilis* ATCC 43071 | >64, >128 (2) | >64 (1) |
| *Pseudomonas aeruginosa* | 45 | *P. aeruginosa* ATCC 27853 | >64, >128 (2) | >64, >64 (2) |
|  |  | *P. aeruginosa* NCTC 13437 | >64, >128 (2) | >64 (1) |
| *Acinetobacter baumannii* | 40 | *A. baumannii* AR0275 | >64, >128 (2) | >64 (1) |
|  |  | *A. baumannii* NCTC 12156 | >64, >128 (2) | >64 (1) |

