## Supplementary Information for "Discovery and chemical optimisation of a Potent, Bi-cyclic (Bicycle^®^) Antimicrobial Inhibitor of *Escherichia coli* PBP3"

Table S-1: Characterization of hits from phage display selections against *Ec*PBP3………… S3

Table S-2: Binding affinity to *Ec*PBP3 for peptide 2 when fluorescently labelled at the C (Tracer 1) or N (Tracer 2) terminus………… S4

Figure S-1: Schematic of fluorescently-labelled peptides………….. S4

Figure S-2: Bocillin FL- and coomassie-stained gel of peptides binding to PBPs extracted from the bacterial inner membrane………….. S5

Table S-3 Statistics of crystalised complex of *Ec*PBP3 and peptide 2……….. S6

Figure S-3: Comparison of *Ec*PBP3 in complex with peptide 2 as compared to various previously reported apo structures of *Ec*PBP3………….. S7

Figure S-4. Demonstration of the tight-fitting nature of the interaction of the Bicycle molecule and protein….………… S8

Figure S-5: Two stereo views of the electron density on the Bicycle molecule………… S9

Figure S-6 View of the active site of *Ec*PBP3 complexed with peptide 2……. S10

Figure S-7 Two views comparing the hydrogen bonds formed in the complex of reacted piperacillin and PBP3 and those formed in the complex of peptide 2 with *Ec*PBP3………….. S11

Table S-4: Sequence homology (%) of the gene product of *ftsI* from relevant organisms of the Enterobacterales, plus *A. baumannii* and *P. aeruginosa*………… S12

Table S-5: MIC and binding evaluation of ala scan peptides…………. S12

Figure S-8: Non-natural substitutions screened in natural peptide 2……… S13

Table S-6: Binding affinity for EcPBP3 and activity against hyperporinated *E. coli* for peptides in the peptide 2 series………….. S14

Table S-7: MIC evaluation of peptides and conjugates against *E. coli*……….. S16

Figure S-9: Cytotoxicity of peptide 2 and conjugate 1 against human cells…………. S17

Table S-8: Plasma stability (mouse) of conjugates with DRAMP18563 variants………… S18

Table S-9: Blood stability (mouse) for SAR peptides…………. S18

Table S-10: Blood stability (mouse) for SAR conjugates…………………………… S18

Peptide sequences…………………………………………………………………………….. S29
Supplementary methods…………………………………………………………………… S23
Peptide QC data………………………………………………………………………............. S28

**Table S-1: Characterization of hits from phage display selections against EcPBP3.** Sequence motifs are highlighted in bold. NB, non-binder (compounds tested at 100 μM top concentration); NT, not tested.

| Peptide | Sequence | Copies in sequencing output | IC50 (nM) by fluorescence polarisation with Bocillin FL as tracer, Geometric mean (Lower 95% CI of geo. Mean; Upper 95% CI of geo. Mean, n=number of values) | *Ki* (nM) by fluorescence polarisation with tracer **1**, Geometric mean (Lower 95% CI of geo. Mean; Upper 95% CI of geo. Mean, n=number of values) | $K_d$ (nM) by SPR |
|---|---|---|---|---|---|
| 1 | ACAADRLWCLAKNDWCA | 43 | NB (n=2) | NT | NT |
| 2 | ACSFPKCPWVEGCA | 7 | **602 (587;618, n=2)** | **<9 (n=46)** | 5.23 |
| 3 | ACAKTPEWLCGFNAYCA | 6 | NB (n=2) | NT | NT |
| 4 | ACRGESSFCLMFPELCA | 5 | NB (n=2) | NT | NT |
| 5 | ACRTFGCWWEGCA | 2 | **1550 (469;2630, n=2)** | NT | 369 |
| 6 | ACADTIYSLTCVPYPCA | 2 | NB (n=2) | NT | NT |
| 7 | ACATEFL**YPL**CW**W**DNCA | 6 | NB (n=2) | NT | NT |
| 8 | ACEVV**YPL**CY**W**SDCA | 1 | NB (n=2) | NT | NT |
| 9 | ACYK**YPGL**CLEFTNMCA | 1 | NB (n=2) | NT | NT |
| 10 | AC**YPGL**PPELCSPSFCA | 1 | NB (n=2) | NT | NT |
| 11 | ACPYA**YPGL**CLEFKSCA | 1 | NB (n=2) | NT | NT |
| 12 | ACPERLCA**LGLP**T**L**RCA | 1 | NB (n=2) | NT | NT |
| 13 | ACLENCYYVPYYGYACA | 1 | **3800 (3000;4600, n=2)** | NT | NT |
| 14 | ACIRERCWDLKENDWCA | 1 | NB (n=2) | NT | NT |
| 15 | ACARPVVLCYWPEDCA | 1 | NB (n=2) | NT | NT |

13

**Table S-2: Binding affinity to EcPBP3 for peptide 2 when fluorescently labelled at the C (Tracer 1) or N (Tracer 2) terminus**.

| Compound | N-terminus | C-terminus | $K_d$ by fluorescence direct binding, Geomean (Lower 95% CI of geo. Mean; Upper 95% CI of geo. Mean, n=number of values) (nM) | $K_d$ by SPR (nM) |
|---|---|---|---|---|
| Tracer 1 | H | Sar$_6$K[FI] | 2110 (1830;2380, n=2) | NT |
| Tracer 2 | [FI]GSar$_5$ | $NH_2$ | 6.86 (5.12;8.60, n=47) | 28.1 |

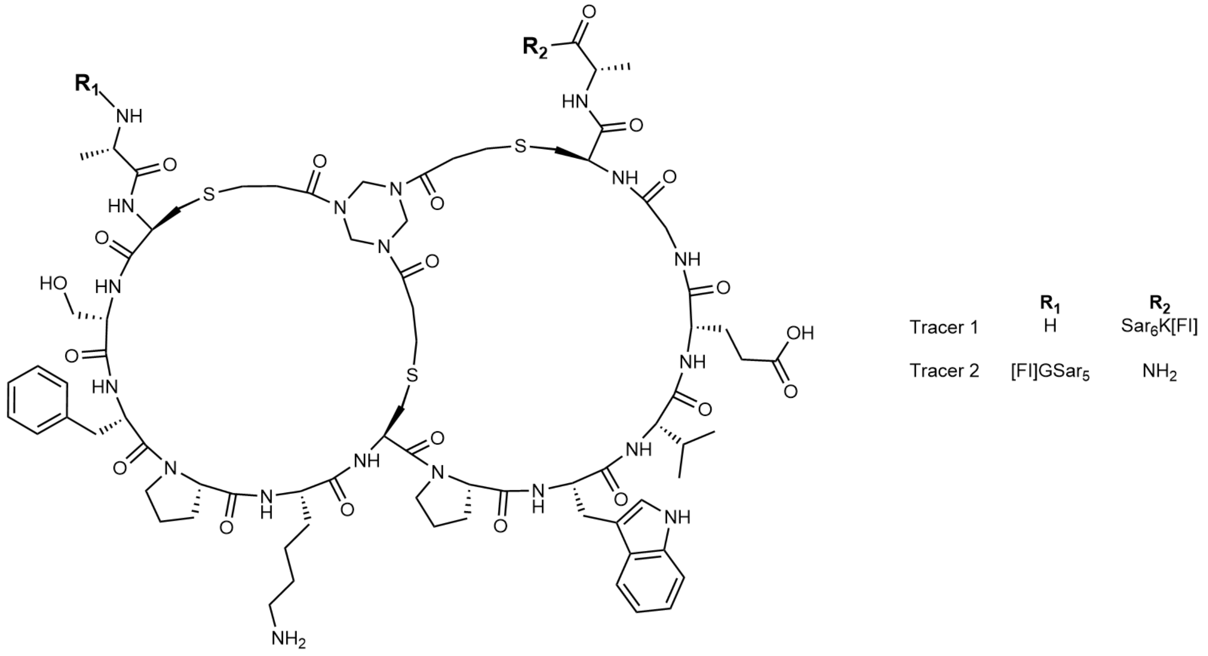

**Figure S-1: Schematic of fluorescently-labelled peptides.** Labelled peptides were used as tracers in fluorescence polarisation assays. Sar, sarcosine; FI, fluorescein.

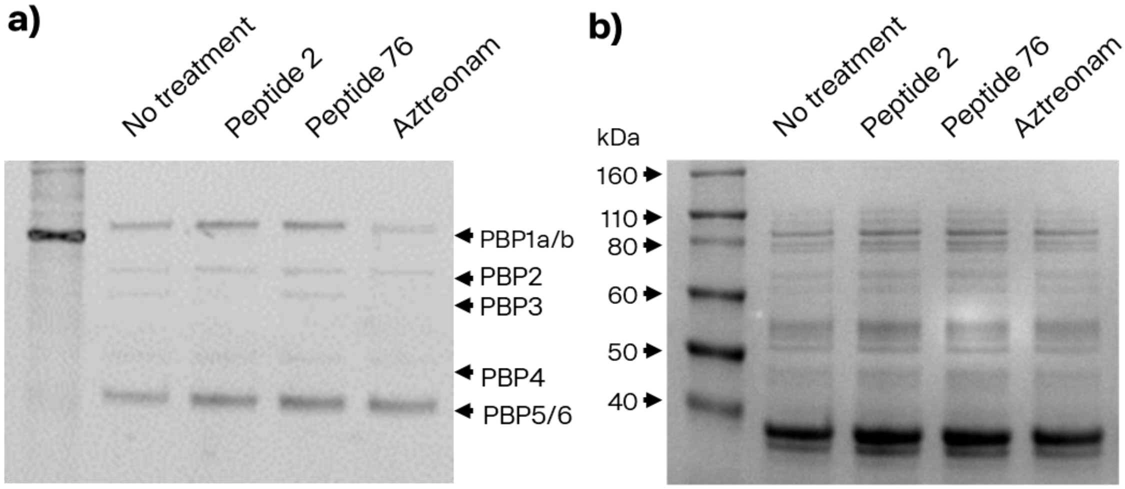

**Figure S-2: Bocillin FL- and coomassie-stained gel of peptides binding to PBPs extracted from the bacterial inner membrane.** PBPs were extracted from the bacterial inner membrane, incubated with the indicated compounds (Peptides at 3 µM, aztreonam at 10 µM) for 15 minutes then stained with Bocillin FL (10 µg/ml) for 10 minutes before being washed and imaged (a) for fluorescence when excited at 455-485 nM and (b) then stained with Coomassie and imaged in brightfield. Aztreonam was used as a positive control. Ladder is Novex Sharp Pre-Stained Protein Standard (Invitrogen), the 80 kDa band of which exhibits fluorescence when excited at 455-485 nM.

**Table S-3 Statistics of crystalised complex of EcPBP3 and peptide 2.**

| ***Data collection and processing statistics*** | |
|---|---|
| Synchrotron, Beam line | SLS, PXIII |
| Date of data collection | 23rd Apr 2020 |
| Wavelength (Å) | 1.000 |
| Detector type | Pilatus 2M |
| Transmission (%) | 100 |
| Temperature (K) | 100 |
| Exposure time (s) | 0.1 |
| Oscillation range per frame (°) | 0.2 |
| Overall rotation (°) | 360 |
| Resolution range (Å) | 45.04 – 1.52 |
| Number of observed reflections | 618393 |
| Number of unique reflections | 50525 |
| Multiplicity (overall and last shell) | 12.2 (7.1) |
| Completeness (%) (overall and last shell) | 99.6 (93.0) |
| $R_{pim}$ (%) (overall and last shell) | 0.6 (42.5) |
| Mean I/sigma (overall and last shell) | 23.8 (1.7) |
| Space group | $C\ 222_1$ |
| Unit cell parameters (Å); (°) | 97.89, 152.18, 43.67; 90, 90, 90 |

| ***Refinement statistics*** | |
|---|---|
| Refinement program | Refmac5 |
| Resolution range (Å) | 45.08 – 1.52 |
| Number of reflections (working; test) | 50510; 2525 |
| $R_{factor}$ (%) | 18.2 |
| $R_{free}$ (%) | 21.6 |
| Number of protein atoms modeled | 2838 |
| Number of water atoms modeled | 261 |
| Number of ligand atoms modeled | 24 |
| RMSD Bond lengths (Å) | 0.015 |
| RMSD Bond angles (°) | 1.88 |
| Mean overall protein B value ($Å^2$) | 26.38 |
| Mean water B value ($Å^2$) | 35.91 |
| Mean ligand B value ($Å^2$) | 30.40 |
| Ramachandran plot favored (%) | 98.49 |
| Ramachandran plot allowed (%) | 1.21 |
| Ramachandran plot outlier region (%) | 0.3 |

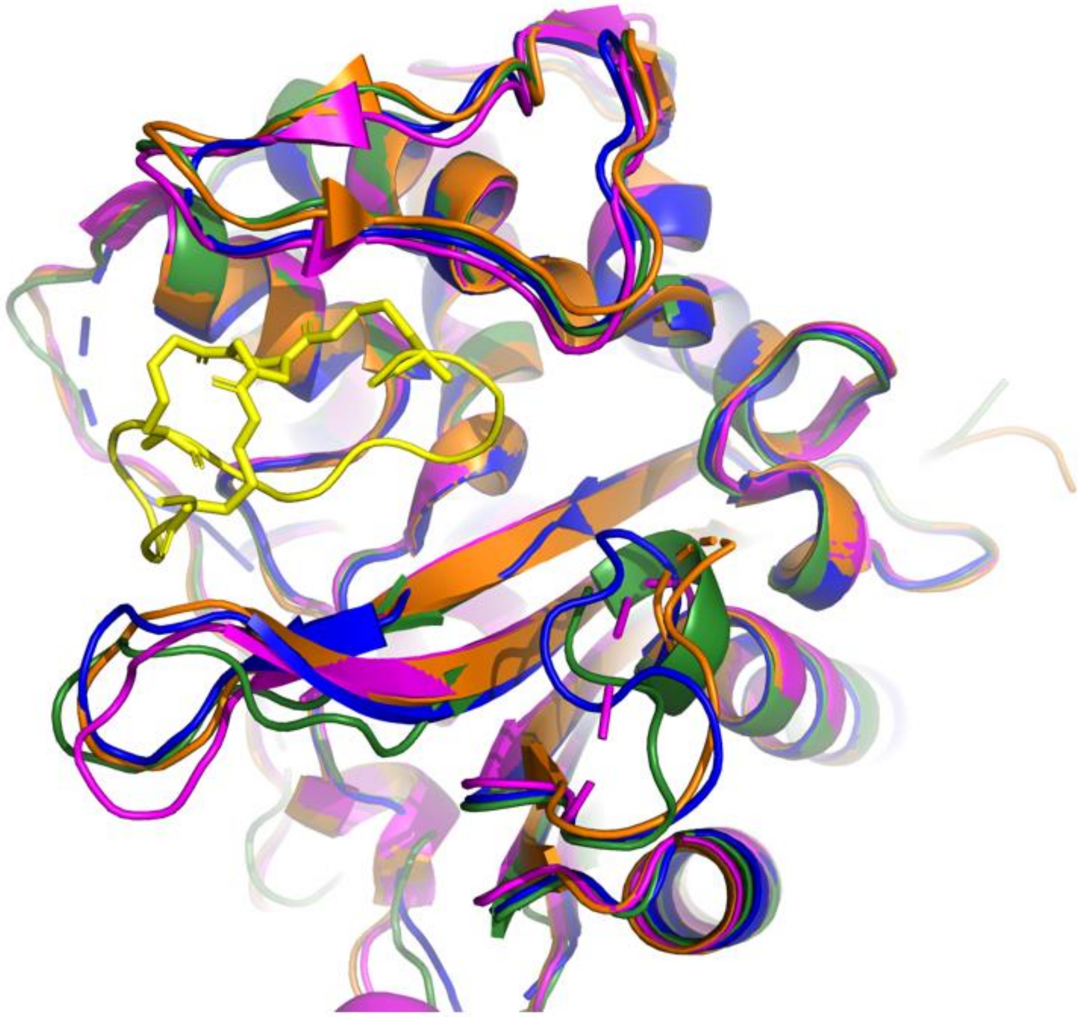

**Figure S-3: Comparison of EcPBP3 in complex with peptide 2 (PDB code: 8RTZ) as compared to various previously reported apo structures of EcPBP3.** PDB codes: 4BJP (Pink), 7ONO (Orange), 6HZQ (Blue). [1–3] Despite the presence of the large ligand (yellow), few changes in the backbone structure of the protein were observed, with only flexible loop regions such as the β3-β4 loop and the β5-α11 varying significantly between the structures. [3] Root mean squared deviation (RMSD) of 0.357 Å (calculated by Pymol, aligning *apo* EcPBP3 (PDB code: 6HZQ) and the EcPBP3:peptide complex).

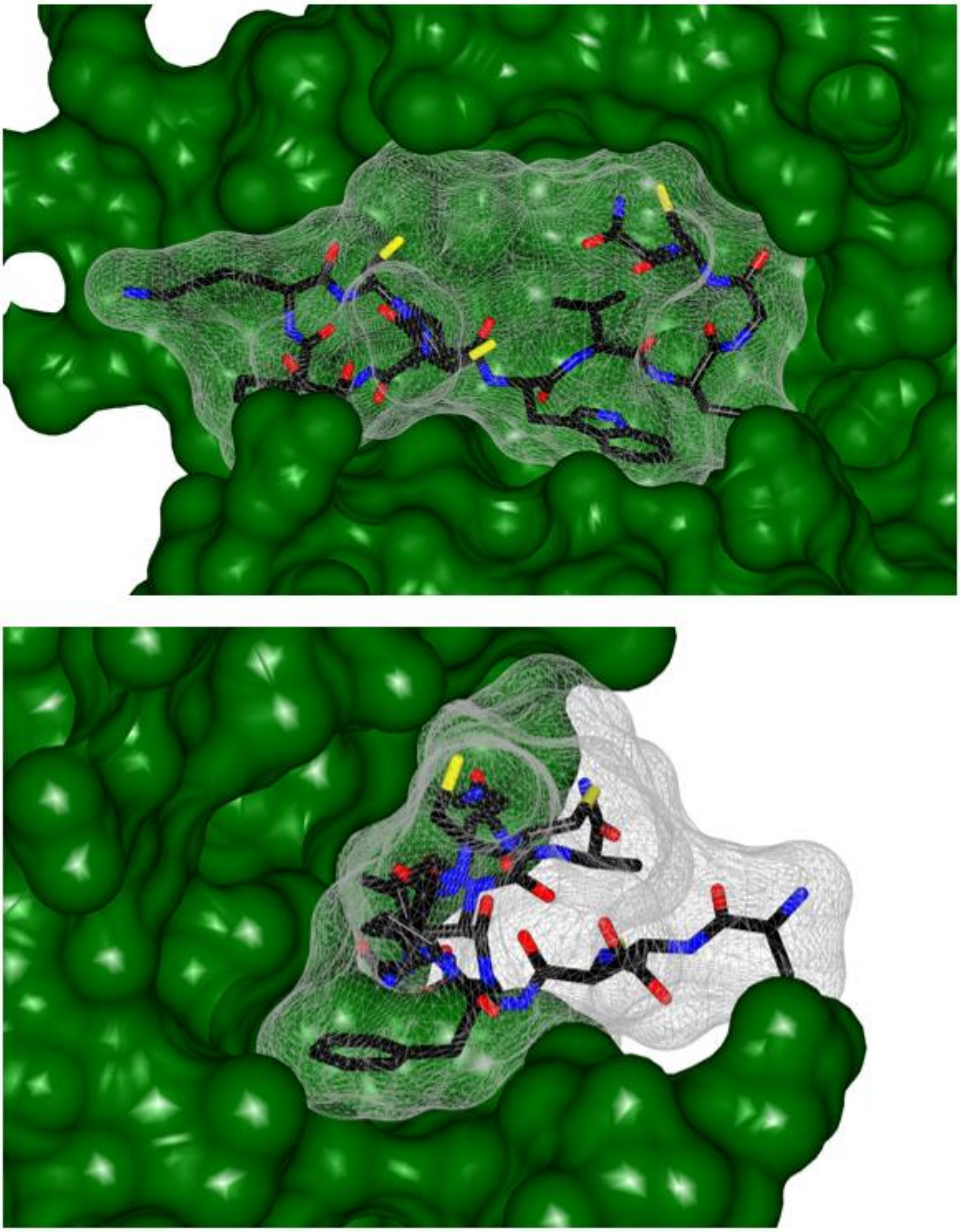

45 **Figure S-4. Demonstration of the tight-fitting nature of the interaction of the Bicycle molecule and protein.** Two views of the surface of the peptide:protein interaction. The peptide (peptide **2**) is shown in black and its space filling representation shown in grey mesh. The protein is represented in

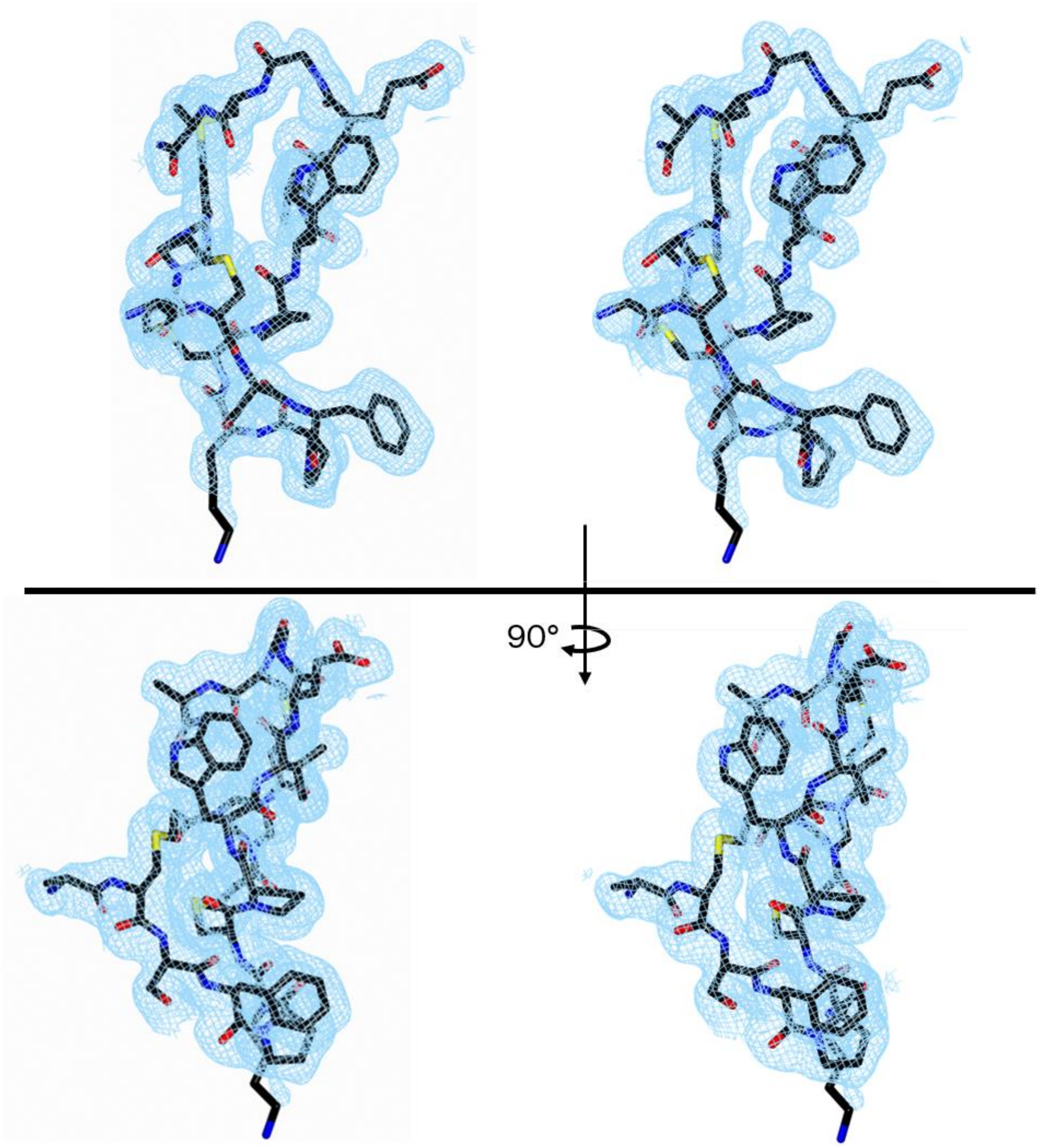

**Figure S-5: Two stereo views of the electron density on the Bicycle molecule.** Electron density is contoured at 1 σ. Composite Omit maps generated with PHENIX. [4] Figure generated in CCP4mg (v2.10.11).

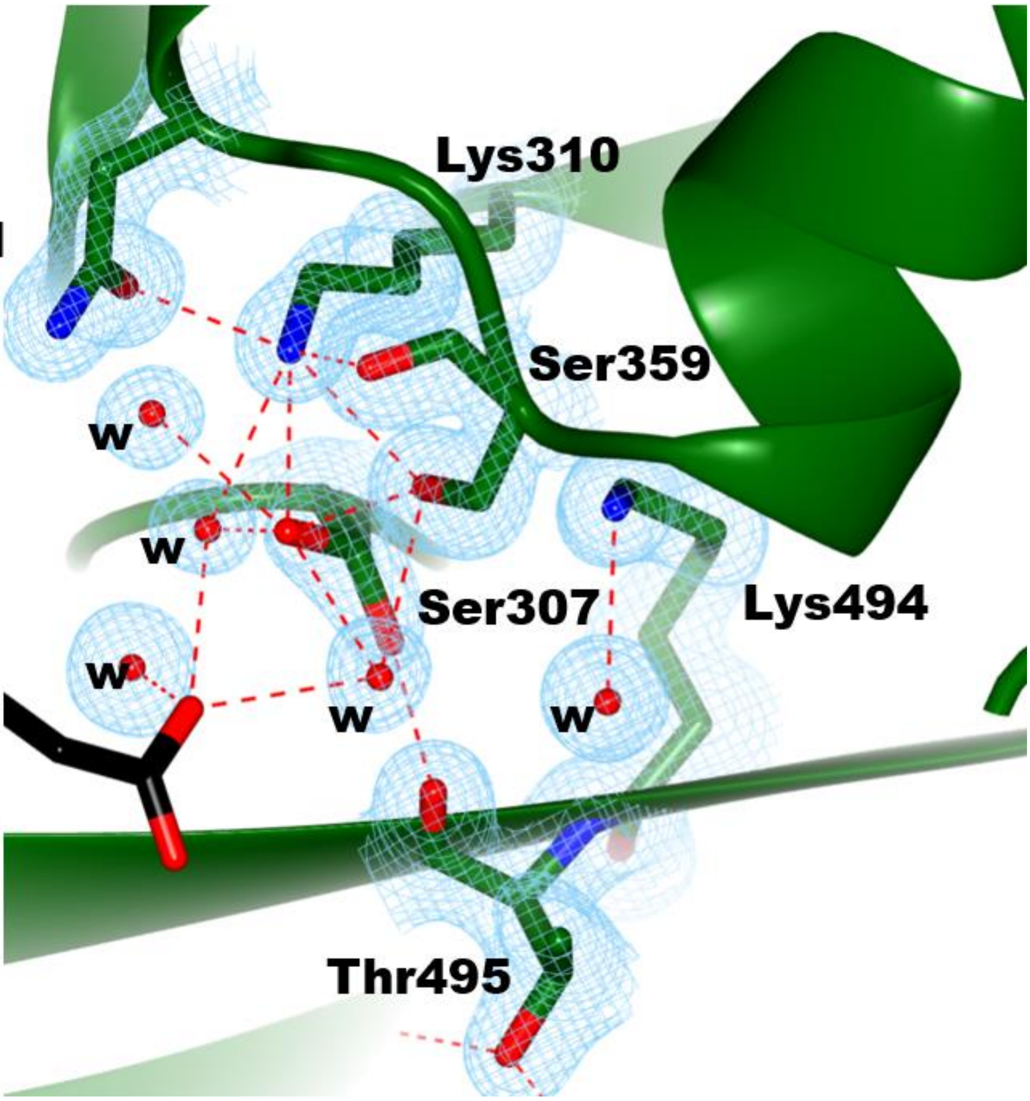

**Figure S-6 View of the active site of EcPBP3 complexed with peptide 2.** Side chains and electron densities are shown for conserved residues of the active site (green): Ser307 and Lys 310 (from the **S**xx**K** motif); Ser 359 and Asn361 (from the SxN motif); Lys494 and Thr495 from the **K**(S/**T**)G motif as well as several waters in the active site (indicated with a **w**). The glutamate residue (Glu8) from peptide 2 (black) is found adjacent (5.3 Å from the Glutamate Oε1 atom to Ser307 Cα atom) to the active site, but only interacts with active site residues via bridging waters. Density (light blue mesh, Composite Omit maps generated with PHENIX) for all active site residues is well defined. The catalytic serine (Ser307) exists in two alternate conformations (each with refined with 50% occupancy). Hydrogen bonds (red dashed lines) are shown between residues and key water molecules in the active site. Figure generated in CCP4mg (v2.10.11).

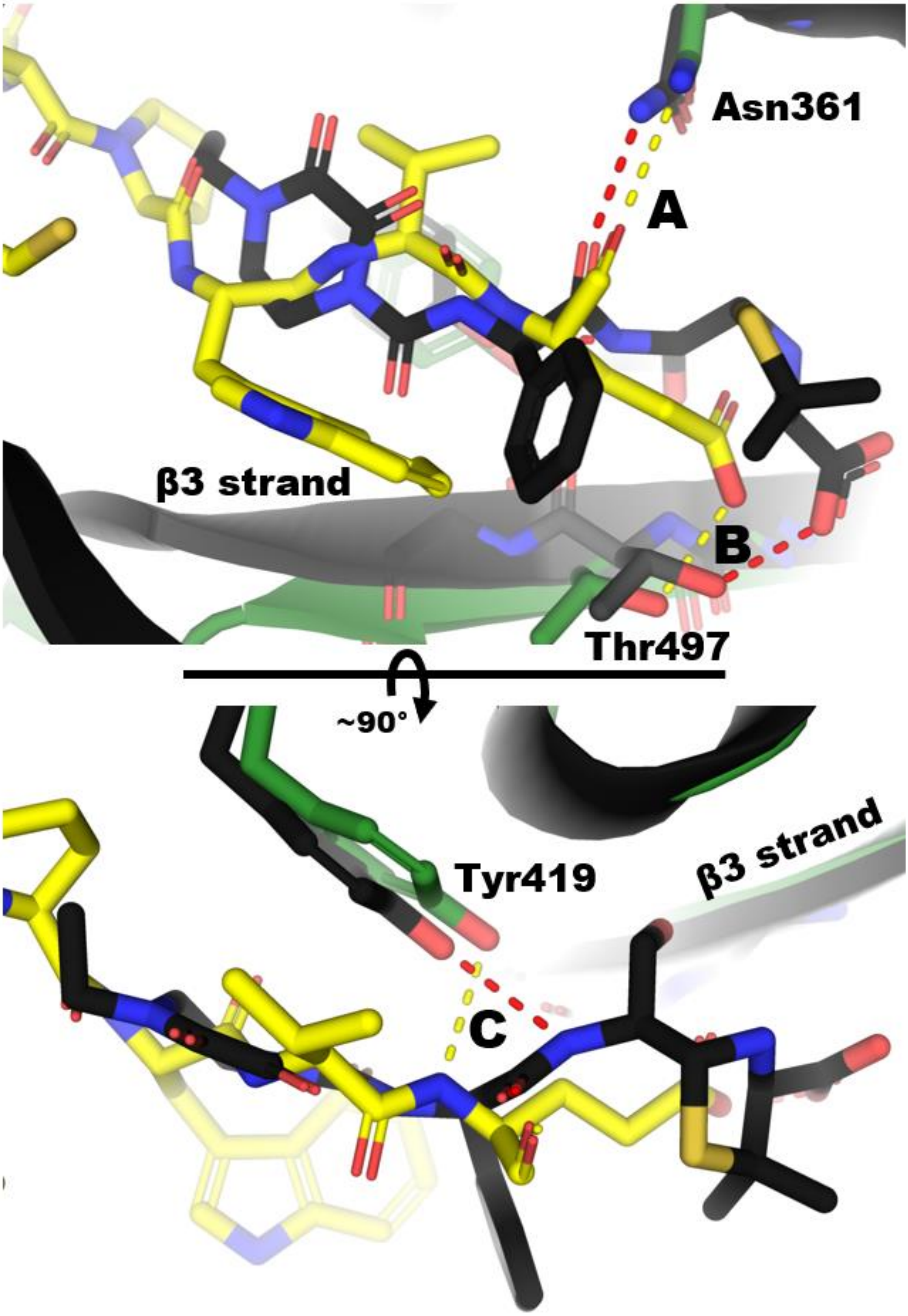

**Figure S-7 Two views comparing the hydrogen bonds formed in the complex of reacted piperacillin and PBP3 (red dashes, from PDB code: 6I1I) and those formed in the complex of peptide 2 with EcPBP3 (yellow dashes).** Three hydrogen bonds are conserved in the two binding interactions, labelled '**A**', '**B**' and '**C**'. Hydrogen bond '**A**' is formed between Asn361 and a peptide carboxyl in both molecules. Asn361 is in one of the conserved active site motifs of the PBPs (the Sx**N** motif) and this bond is seen in many other β-lactam:PBP structures. [3] Hydrogen bond '**B**' is formed between acidic groups on both molecules and Thr497. In β-lactams, this acidic group is highly conserved. Hydrogen bond '**C**' is formed between Tyr419's OH and a backbone amide nitrogen. The conservation of these hydrogen bonds in the two ligands is evidence of a shared binding mode.

**Table S-4: Sequence homology (%) of the gene product of *ftsI* from relevant organisms of the Enterobacterales, plus *A. baumannii* and *P. aeruginosa*.** Sequences are labelled with gene, organism and the NCBI accession number for the protein sequence.

| | FTSI_ECOLI | FTSI_CFREUNDII | FTSI_KPNEU | FTSI_ECLOACAE | FTSI_PMIRABILIS | FTSI_PAERU | FTSI_ABAUM |
|---|---|---|---|---|---|---|---|
| FTSI_ECOLI _WP_000642196.1 | 100 | 96 | 94 | 94 | 75 | 45 | 40 |
| FTSI_CFREUNDII _WP_088902745.1 | 96 | 100 | 93 | 95 | 74 | 46 | 39 |
| FTSI_KPNEUMONIAE _WP_002888559.1 | 94 | 93 | 100 | 96 | 75 | 46 | 40 |
| FTSI_ECLOACAE _CP033466.1 | 94 | 95 | 96 | 100 | 75 | 45 | 40 |
| FTSI_PMIRABILIS _EEI48461.1 | 75 | 74 | 75 | 75 | 100 | 43 | 39 |
| FTSI_PAERUGINOSA _UTQ31465.1 | 45 | 46 | 46 | 45 | 43 | 100 | 39 |
| FTSI_ABAUMANNII _AEK25671.1 | 40 | 39 | 40 | 40 | 39 | 39 | 100 |

**Table S-5: MIC and binding evaluation of ala scan peptides.** MICs were generated in caMHB medium. MIC data are given for n-3 biological replicates - values for peptide 2 are representative of range across >8 replicates in separate assays.

| Peptide | Sequence | *Ki* (nM) by fluorescence polarisation with BCY12820 as tracer, Geometric mean (Lower 95% CI of geo. Mean; Upper 95% CI of geo. Mean, n=number of values) | $K_d$ (nM) by SPR | MIC (µg/mL) *E. coli* GKCW102 (EcPore) |
|---|---|---|---|---|
| 2 | ACSFPKCPWVEGCA | <9 (n=46) | 5.23 | 0.5, 1, 2 |
| 16 | ACAFPKCPWVEGCA | <9 (n=3) | 6.44 | 0.5, 2, 2 |
| 17 | ACSAPKCPWVEGCA | 331, >5000 (n=2) | NB[1] | >64, >64, 256 |
| 18 | ACSFAKCPWVEGCA | 76.9 (68.9;85.0, n=3) | 178 | 4, 8, 8 |
| 19 | ACSFPACPWVEGCA | <9 (n=2) | 12.4 | 0.5, 0.5, 1 |
| 20 | ACSFPKCAWVEGCA | 294, >5000 (n=2) | 1380 | 64, 64, 128 |
| 21 | ACSFPKCPAVEGCA | >5000 (n=2) | NB[1] | >64, >64, >256 |
| 22 | ACSFPKCPWAEGCA | >5000 (n=2) | 669 | 32, 32, 32 |
| 23 | ACSFPKCPWVAGCA | 9.08 (4.64;13.5, n=3) | 6.71 | 0.25, 0.5, 0.5 |
| 24 | ACSFPKCPWVEACA | 25.5 (18.2;32.7, n=3) | 24.4 | 0.5, 1, 2 |

[1] No binding observed at 2500 nM

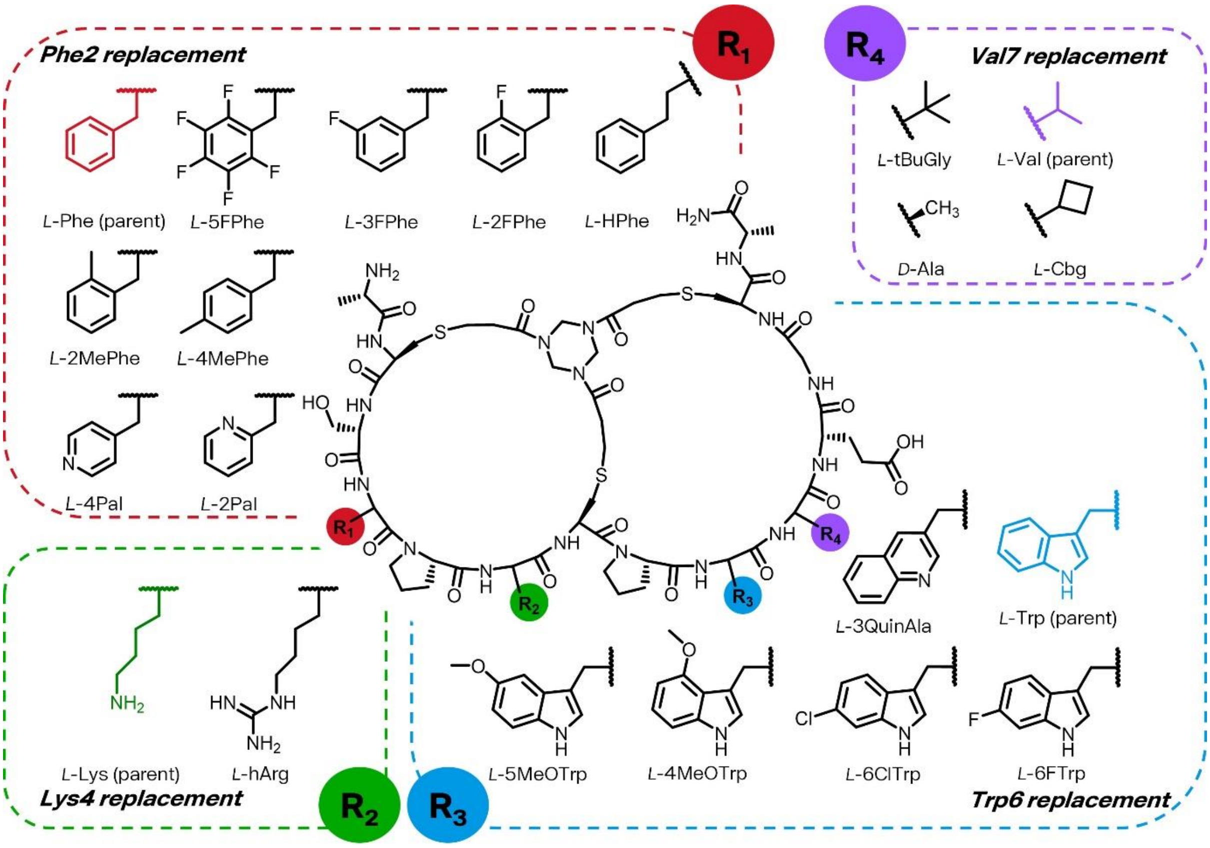

**Figure S-8: Non-natural substitutions screened in natural peptide 2.** 5FPhe, pentafluoro-phenylalanine; 3FPhe, 3-fluoro-phenylalanine; 2FPhe, 2-fluoro-phenylalanine; HPhe, homophenylalanine; 2MePhe, 2-methyl-phenylalanine; 4MePhe, 4-methyl-phenylalanine; 4Pal, 4-pyridylalanine; 2Pal, 2-pyridylalanine; hArg, homoarginine; 3QuinAla, 3-Quinoyl-L-alanine; 5MeOTrp, 5-methoxytryptophan; 4MeOTrp, 4-methoxytryptophan; 6ClTrp, 6-chlorotryptophan; 6FlTrp, 6-fluorotryptophan; tBuGly, tert-butyl-leucine; Cbg, cyclobutylglycine.

**Table S-6: Binding affinity for EcPBP3 and activity against hyperporinated *E. coli* for peptides in the peptide 2 series.** Substituted amino acids are marked in bold.

| Peptide no. | Sequence | *Ki* (nM) with BCY12820 as tracer, Geometric mean (Lower 95% CI of geo. Mean; Upper 95% CI of geo. Mean, n=number of values) | $K_d$ (nM) by SPR | MIC (µg/mL) *E. coli* GKCW102 (EcPore) |
|---|---|---|---|---|
| 25 | ACSFP**[hArg]**CPWVEGCA | <9 (n=7) | 2.85 | 0.5, 0.5, 0.5 |
| 26 | AC**G**FP**[HArg]**CPWVEGCA[CONH2] | 17.1 (1.44;32.7; n=2) | NT | 2, 2, 4 |
| 27 | AC**L**FP**[HArg]**CPWVEGCA[CONH2] | <9 (n=2) | 3.36 | 0.25, 0.25, 0.25 |
| 28 | AC**F**FP**[HArg]**CPWVEGCA[CONH2] | <9 (n=2) | 1.12 | 0.125, 0.125, 0.25 |
| 29 | AC**N**FP**[HArg]**CPWVEGCA[CONH2] | <9 (n=2) | NT | 0.25, 1, 1 |
| 30 | AC**Q**FP**[HArg]**CPWVEGCA[CONH2] | <9 (n=2) | NT | 1, 1, 1 |
| 31 | AC**E**FP**[HArg]**CPWVEGCA[CONH2] | <9 (n=2) | NT | 0.25, 0.25, 0.5 |
| 32 | ACS**H**P**[HArg]**CPWVEGCA[CONH2] | 115 (70.1;161, n=3) | NT | 32, 64, 64 |
| 33 | ACSF**G[HArg]**CPWVEGCA[CONH2] | 53.5 (51.1;55.8, n=2) | NT | 4, 8, 8 |
| 34 | ACSF**Y[HArg]**CPWVEGCA[CONH2] | 27.1 (11.4;42.8, n=2) | NT | 4, 4, 4 |
| 35 | ACSF**S[HArg]**CPWVEGCA[CONH2] | 53.4 (32.7;74.1, n=2) | NT | 8, 16, 16 |
| 36 | ACSF**T[HArg]**CPWVEGCA[CONH2] | 75.0 (55.0;95.0, n=2) | NT | 8, 8, 16 |
| 37 | ACSF**H[HArg]**CPWVEGCA[CONH2] | 31.5 (13.3;49.7, n=2) | NT | 4, 4, 4 |
| 38 | ACSF**D[HArg]**CPWVEGCA[CONH2] | 167 (29.7;304, n=2) | NT | 32, 64, 64 |
| 39 | ACSFP**T**CPWVEGCA[CONH2] | <9, 21.6 (n=2) | NT | 0.5, 0.5, 1 |
| 40 | ACSFP**H**CPWVEGCA[CONH2] | <9, 19.4 (n=2) | NT | 0.5, 0.5 ,0.5 |
| 41 | ACSFP**[HArg]**C**G**WVEGCA[CONH2] | >5000 (n=2) | NT | NT |
| 42 | ACSFP**[HArg]**CP**Y**VEGCA[CONH2] | 209 (114;305, n=4) | NT | NT |
| 43 | ACSFP**[HArg]**CP**F**VEGCA[CONH2] | 252, >5000 (n=2) | NT | NT |
| 44 | ACSFP**[HArg]**CP**H**VEGCA[CONH2] | >5000 (n=2) | NT | NT |
| 45 | ACSFP**[HArg]**CPW**I**EGCA[CONH2] | <9 (n=2) | NT | 0.5,0.5,2 |
| 46 | ACSFP**[HArg]**CPW**T**EGCA[CONH2] | 15.5 (11.5;19.5, n=2) | NT | 4, 4, 4 |
| 47 | ACSFP**[HArg]**CPWV**P**GCA[CONH2] | 18.0 (11.7;24.3, n=2) | NT | 4, 8, 16 |

| Peptide no. | Sequence | *Ki* (nM) with BCY12820 as tracer, Geometric mean (Lower 95% CI of geo. Mean; Upper 95% CI of geo. Mean, n=number of values) | $K_d$ (nM) by SPR | MIC (µg/mL) *E. coli* GKCW102 (EcPore) |
|---|---|---|---|---|
| 48 | ACSFP**[HArg]**CPWV**L**GCA[CONH2] | 49.7 (29.2;70.2, n=2) | NT | 16, 16, 64 |
| 49 | ACSFP**[HArg]**CPWV**F**GCA[CONH2] | <9, 23.4 (n=2) | NT | 4, 8, 16 |
| 50 | ACSFP**[HArg]**CPWV**Q**GCA[CONH2] | <9, 32.7 (n=2) | NT | 2, 8, 8 |
| 51 | ACSFP**[HArg]**CPWVE**Y**CA[CONH2] | <9 (n=2) | 1.83 | 0.125, 0.25, 0.5 |
| 52 | ACSFP**[HArg]**CPWVE**N**CA[CONH2] | <9 (n=2) | NT | 1, 2, 4 |
| 53 | ACSFP**[HArg]**CPWVE**H**CA[CONH2] | <9 (n=2) | NT | 1, 1, 2 |
| 54 | ACS**[5FPhe]**PKCPWVEGCA[CONH2] | NT | 701 | 64, 64, >64 |
| 55 | ACS**[3FPhe]**PKCPWVEGCA[CONH2] | NT | 16.8 | 1, 1, 1 |
| 56 | ACS**[2FPhe]**PKCPWVEGCA[CONH2] | NT | 56.6 | 0.5, 2, 2 |
| 57 | ACS**[2MePhe]**PKCPWVEGCA[CONH2] | NT | 22.3 | 0.5, 1, 1 |
| 58 | ACS**[4MePhe]**PKCPWVEGCA[CONH2] | NT | 5.06 | 0.5, 1, 1 |
| 59 | ACS**[4Pal]**PKCPWVEGCA[CONH2] | NT | 121 | 1, 2, 4, |
| 60 | ACS**[2Pal]**PKCPWVEGCA[CONH2] | NT | 122 | 16, 32, 64 |
| 61 | ACS**[HPhe]**PKCPWVEGCA[CONH2] | NT | 232 | 16, 16, 64 |
| 62 | ACSFPKCP**[3QuinAla]**VEGCA[CONH2] | NT | 27 | 0.5, 0.5, 2 |
| 63 | ACSFPKCP**[5MeoTrp]**VEGCA[CONH2] | NT | 175 | 4, 4, 16 |
| 64 | ACSFPKCP**[4MeoTrp]**VEGCA[CONH2] | NT | NT | 64, >64, >64 |
| 65 | ACSFPKCP**[6ClTrp]**VEGCA[CONH2] | NT | 0.269 | 0.0625, 0.0625, 0.125 |
| 66 | ACSFPKCP**[6FTrp]**VEGCA[CONH2] | NT | 0.489 | 0.125, 0.125, 0.25 |
| 67 | ACSFPKCPW**[tBuGly]**EGCA[CONH2] | NT | 574 | 8, 16, >16 |
| 68 | ACSFPKCPW**[Cbg]**EGCA[CONH2] | NT | 187 | 4, 8, 8 |
| 69 | ACSFPKCPW**[dA]**EGCA[CONH2] | NT | NB[1] | >16, >16, >16 |

[1] No binding observed at 2500 nM

**Table S-7: MIC evaluation of peptides and conjugates against *E. coli*.** MICs were determined in caMHB for hyperporinated strains, and in MHB for *E. coli* ATCC 25922.

| Compound | MIC (µg/ml) | | | *Ki* (nM) by fluorescence polarisation with tracer **1**, Geometric mean (Lower 95% CI of geo. Mean; Upper 95% CI of geo. Mean, n=number of values) |
|---|---|---|---|---|
| | *E. coli* GKCW101 (WT) | *E. coli* GKCW102 (EcPore) | *E. coli* ATCC 25922 | |
| Peptide 2 | >128, >128, >128 | 0.5, 1, 2 | >16, >16, >16 | <9 (n=46) |
| Conjugate 2 | 4, 4, 8 | 0.5, 1, 1 | 1, 2, 2 | 27.7nM (12.5, 42.9) n=16 |
| Conjugate 3 | >64, >64, >64 | 16, 16, 16 | 64, 64, >64 | NT |

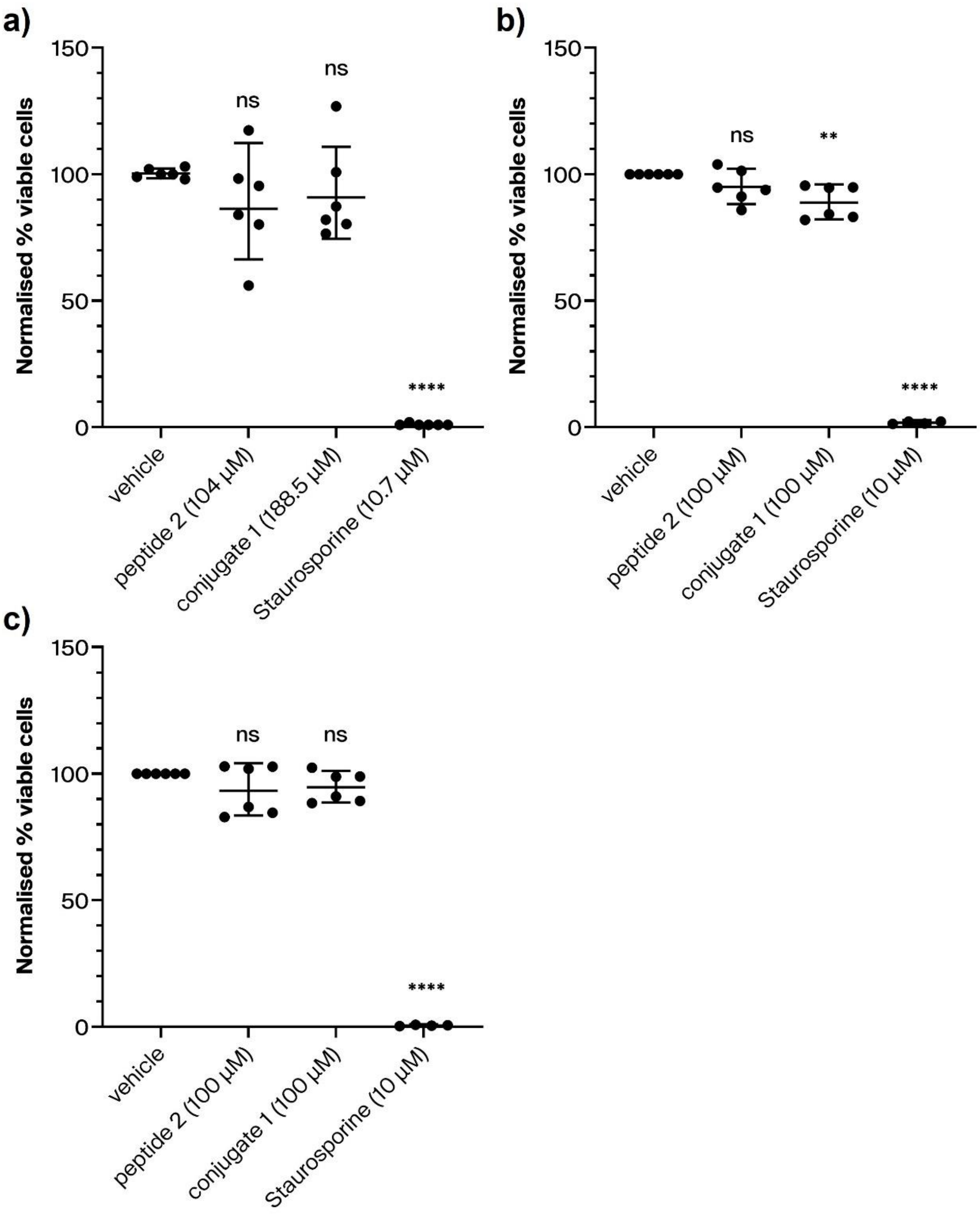

**Figure S-9: Cytotoxicity of peptide 2 and conjugate 1 against human cells.** Compounds were tested against a) HT1080 cells, b) A549 cells and c) HepG2 cells. Each point represents a technical replicate, derived from n≥2 biological replicates. Data were analyzed using an unpaired t-test (two-tailed) to compare compounds with the vehicle control. Whilst a significant difference was observed in cell viability of A549 cells treated with conjugate 1 compared to vehicle, the IC50 of this effect was >100µM, thereby indicating good specificity for bacterial killing compared to cytotoxicity. ns, no significant difference; **, *P* <0.01; ****, *P* < 0.0001.

**Table S-8: Plasma stability (mouse) of conjugates with DRAMP18563 variants**

| Conjugate | DRAMP variant | Plasma stability ($t_{1/2}$), h |
|---|---|---|
| 1 | L-amino acids | 0.3 |
| 2 | Retroinverso | 5.7 |
| 8 | D-amino acids | 5.3 |

71

**Table S-9: Blood stability (mouse) for SAR peptides.** Results are the average of two technical replicates. Data were generated with EDTA as the anticoagulant. MICs were generated in caMHB medium.

| Peptide | Sequence | Blood stability ($t_{1/2}$), h | MIC (µg/ml) *E. coli* GKCW102 |
|---|---|---|---|
| 2 | ACSFPKCPWVEGCA | NT | 0.5, 1, 2 |
| 25 | ACSFP[HArg]CPWVEGCA | 0.2 | 0.25, 0.5, 0.5 |
| 70 | ACSFP[HArg]CPWVEGC | 0.3 | 0.125, 0.125, 0.25 |
| 71 | CSFP[HArg]CPWVEGCA | 5.7 | 0.125, 0.25, 0.5 |
| 72 | CSFP[HArg]CPWVEGC | 3.0 | 0.5, 1, 1 |
| 73 | [-Ac]ACSFP[HArg]CPWVEGCA | 14.1 | 0.5, 1, 2 |
| 74 | [-Ac]CSFP[HArg]CPWVEGCA | 17.7 | 0.25, 0.5, 1 |
| 75 | [-Ac]ACSFP[HArg]CP[6ClTrp]VEGCA | 11.4 | 0.0625, 0.125, 0.125 |

72

**Table S-10: Blood stability (mouse) for SAR conjugates.** Results are the average of two technical replicates. Data were generated with heparin as the anticoagulant. MICs were generated in MHB medium.

| Conjugate | Sequence | Blood stability (t1/2), h | MIC (µg/ml) *E. coli* ATCC25922 |
|---|---|---|---|
| 2 | ACSFPKCPWVEGCA | 6.9 | 0.5, 1, 2 |
| 4 | ACSFP[HArg]CP[6ClTrp]VEGCA | 8.8 | 0.5, 2, 2 |
| 5 | [-Ac]ACSFP[HArg]CP[6ClTrp]VEGCA | 19.9 | 2, 4, 4 |
| 6 | [-Ac]CSFP[HArg]CP[6ClTrp]VEGCA | 4.4 | 0.5, 1, 2 |
| 7 | [-Ac]CSFP[HArg]CPWVEGCA | 2.8 | 1, 1, 4 |

73

74

#### Peptide sequences

| Tracers | |
|---|---|
| Tracer | Sequence |
| 1 | [-Fl]G[Sar5]ACSFPKCPWVEGCA[CONH2] |
| 2 | ACSFPKCPWVEGCA[Sar6][KFl][CONH2] |

| Conjugates | | |
|---|---|---|
| Conjugate | Peptide sequence | Vector sequence |
| 1 | ACSFPKCPWVEGCA[K(PYA)] | KSLRRVWRSWR[CAzal] |
| 2 | ACSFPKCPWVEGCA[K(PYA)] | [CAzal][dR][dW][dS][dR][dW][dV][dR][dR][dL][dS][dK][CONH2] |
| 3 | [dA][dC][dS][dF][dP][dK][dC][dP][dW][dV][dE]G[dC][dA][K(PYA)] | [CAzal][dR][dW][dS][dR][dW][dV][dR][dR][dL][dS][dK] |
| 4 | ACSFP[HArg]CP[6ClTrp]VEGCA[K(PYA)] | [CAzal][dR][dW][dS][dR][dW][dV][dR][dR][dL][dS][dK] |
| 5 | [-Ac]ACSFP[HArg]CP[6ClTrp]VEGCA[K(PYA)] | [CAzal][dR][dW][dS][dR][dW][dV][dR][dR][dL][dS][dK] |
| 6 | [-Ac]CSFP[HArg]CP[6ClTrp]VEGCA[K(PYA)] | [CAzal][dR][dW][dS][dR][dW][dV][dR][dR][dL][dS][dK] |
| 7 | [-Ac]CSFP[HArg]CPWVEGCA[K(PYA)] | [CAzal][dR][dW][dS][dR][dW][dV][dR][dR][dL][dS][dK] |
| 8 | ACSFPKCPWVEGCA[K(PYA)] | [dK][dS][dL][dR][dR][dV][dW][dR][dS][dW][dR][CAzal] |

| Peptides | |
|---|---|
| Peptide | Sequence |
| 1 | ACAADRLWCLAKNDWCA[CONH2] |
| 2 | ACSFPKCPWVEGCA[CONH2] |
| 3 | ACAKTPEWLCGFNAYCA[CONH2] |
| 4 | ACRGESSFCLMFPELCA[CONH2] |
| 5 | ACRTFGCWWEGCA[CONH2] |
| 6 | ACADTIYSLTCVPYPCA[CONH2] |
| 7 | ACATEFLYPLCWWDNCA[CONH2] |
| 8 | ACEVVYPLCYWSDCA[CONH2] |

| | |
|---|---|
| 9 | ACYKYPGLCLEFTNMCA[CONH2] |
| 10 | ACYPGLPPELCSPSFCA[CONH2] |
| 11 | ACPYAYPGLCLEFKSCA[CONH2] |
| 12 | ACPERLCALGLPTLRCA[CONH2] |
| 13 | ACLENCYYVPYYGYACA[CONH2] |
| 14 | ACIRERCWDLKENDWCA[CONH2] |
| 15 | ACARPVVLCYWPEDCA[CONH2] |
| 16 | ACAFPKCPWVEGCA[CONH2] |
| 17 | ACSAPKCPWVEGCA[CONH2] |
| 18 | ACSFAKCPWVEGCA[CONH2] |
| 19 | ACSFPACPWVEGCA[CONH2] |
| 20 | ACSFPKCAWVEGCA[CONH2] |
| 21 | ACSFPKCPAVEGCA[CONH2] |
| 22 | ACSFPKCPWAEGCA[CONH2] |
| 23 | ACSFPKCPWVAGCA[CONH2] |
| 24 | ACSFPKCPWVEACA[CONH2] |
| 25 | ACSFP[HArg]CPWVEGCA[CONH2] |
| 26 | ACGFP[HArg]CPWVEGCA[CONH2] |
| 27 | ACLFP[HArg]CPWVEGCA[CONH2] |
| 28 | ACFFP[HArg]CPWVEGCA[CONH2] |
| 29 | ACNFP[HArg]CPWVEGCA[CONH2] |
| 30 | ACQFP[HArg]CPWVEGCA[CONH2] |
| 31 | ACEFP[HArg]CPWVEGCA[CONH2] |
| 32 | ACSHP[HArg]CPWVEGCA[CONH2] |
| 33 | ACSFG[HArg]CPWVEGCA[CONH2] |
| 34 | ACSFY[HArg]CPWVEGCA[CONH2] |
| 35 | ACSFS[HArg]CPWVEGCA[CONH2] |
| 36 | ACSFT[HArg]CPWVEGCA[CONH2] |
| 37 | ACSFH[HArg]CPWVEGCA[CONH2] |
| 38 | ACSFD[HArg]CPWVEGCA[CONH2] |

| | |
|---|---|
| 39 | ACSFPTCPWVEGCA[CONH2] |
| 40 | ACSFPHCPWVEGCA[CONH2] |
| 41 | ACSFP[HArg]CGWVEGCA[CONH2] |
| 42 | ACSFP[HArg]CPYVEGCA[CONH2] |
| 43 | ACSFP[HArg]CPFVEGCA[CONH2] |
| 44 | ACSFP[HArg]CPHVEGCA[CONH2] |
| 45 | ACSFP[HArg]CPWIEGCA[CONH2] |
| 46 | ACSFP[HArg]CPWTEGCA[CONH2] |
| 47 | ACSFP[HArg]CPWVPGCA[CONH2] |
| 48 | ACSFP[HArg]CPWVLGCA[CONH2] |
| 49 | ACSFP[HArg]CPWVFGCA[CONH2] |
| 50 | ACSFP[HArg]CPWVQGCA[CONH2] |
| 51 | ACSFP[HArg]CPWVEYCA[CONH2] |
| 52 | ACSFP[HArg]CPWVENCA[CONH2] |
| 53 | ACSFP[HArg]CPWVEHCA[CONH2] |
| 54 | ACS[5FPhe]PKCPWVEGCA[CONH2] |
| 55 | ACS[3FPhe]PKCPWVEGCA[CONH2] |
| 56 | ACS[2FPhe]PKCPWVEGCA[CONH2] |
| 57 | ACS[2MePhe]PKCPWVEGCA[CONH2] |
| 58 | ACS[4MePhe]PKCPWVEGCA[CONH2] |
| 59 | ACS[4Pal]PKCPWVEGCA[CONH2] |
| 60 | ACS[2Pal]PKCPWVEGCA[CONH2] |
| 61 | ACS[HPhe]PKCPWVEGCA[CONH2] |
| 62 | ACSFPKCP[3QuinAla]VEGCA[CONH2] |
| 63 | ACSFPKCP[5MeoTrp]VEGCA[CONH2] |
| 64 | ACSFPKCP[4MeoTrp]VEGCA[CONH2] |
| 65 | ACSFPKCP[6ClTrp]VEGCA[CONH2] |
| 66 | ACSFPKCP[6FTrp]VEGCA[CONH2] |
| 67 | ACSFPKCPW[tBuGly]EGCA[CONH2] |
| 68 | ACSFPKCPW[Cbg]EGCA[CONH2] |

| | |
|---|---|
| 69 | ACSFPKCPW[dA]EGCA[CONH2] |
| 70 | ACSFP[HArg]CPWVEGC[CONH2] |
| 71 | CSFP[HArg]CPWVEGCA[CONH2] |
| 72 | CSFP[HArg]CPWVEGC[CONH2] |
| 73 | [-Ac]ACSFP[HArg]CPWVEGCA[CONH2] |
| 74 | [-Ac]CSFP[HArg]CPWVEGCA[CONH2] |
| 75 | [-Ac]ACSFP[HArg]CP[6ClTrp]VEGCA[CONH2] |
| 76 | [dA][dC][dS][dF][dP][dK][dC][dP][dW][dV][dE]G[dC][dA][CONH2] |

### Supplementary methods

#### Bocillin FL staining of bacterial membranes

Adaption of the method of Kocaoglu and Carlson.[6] E. coli strain DC2 cells were grown overnight at 37 °C then diluted 1 in 100 and allowed to grow at 37 °C to an OD600 of 0.5 in Luria broth. The cells were then centrifuged (3,220 x g for 10 min at room temperature) and washed once in PBS (pH 7.4) then pelleted and resuspended in ice-cold PBS. Cells were sonicated (Fisherbrand™ Model 505 Sonic Dismembrator) on ice for five 10 s intervals with 15 s cooling time in between rounds. The membranes were collected by centrifugation (21,000 x g for 15 min at 4ºC) and resuspended the pellet in 100 µL PBS using a blunt needle. The resuspended membranes were incubated with the indicated compound for 15 minutes at room temperature then 10 µg/ml BOCILLIN-Fl (Invitrogen) was added and reacted with the resuspended membranes for 10 minutes. The membranes were washed once as above in cold PBS and resuspended the pellet in 100 µL PBS using a blunt needle. The total protein content of each condition was normalised to 2.5 mg/ml by measuring the absorbance at A280 and adjusting where necessary with the addition of PBS. Samples were heated at 95 °C for 5 minutes in SDS-PAGE loading buffer and 10 µL was loaded onto a 10% Bis-Tris gel (NuPAGE Novex, Invitrogen). The gel was run for 45 min at 180 V, 400 mA, then imaged with an iBright FL1500 (excitation 455-485 nm, emission filter 508-557 nm, software version 1.8.0). Following fluorescence imaging, the gel was stained with InstantBlue Ultrafast protein stain (Abcam), washed twice in water and imaged in brightfield with an iBright FL1500.

#### Fluorescence Polarisation

Competition fluorescence polarisation assays were run by observing the fluorescence polarisation of a fluorescent "tracer" Bicycle molecule which competes against an unlabelled peptide for binding to the *Ec*PBP3 target, as previously described.[7] 10 μL of protein (9.4 nM) was mixed with either 5 μL of peptide or 5 μL of buffer and then the assay was initiated by the addition of 10 μL of tracer (1 nM) (final volume 25 μL), in a black 384 low bind, low volume plate (Corning). The buffer was 10 mM HEPES (Sigma), 2% (v/v) glycerol (Sigma), 300 mM NaCl (Sigma), adjusted to pH 8. The plate was incubated for 1 hour at 25 °C and then read in a PHERAstar FS (BMG) using with an "FP 485 520 520" optic module (excitation 485 nm, emission 520 nm). The gain was determined immediately prior to the read on a tracer-only well.

The $K_d$ of the tracer molecule was found using the above method with the exclusion of the competing peptide and titration of the protein concentration (0.8 μM top concentration) against a fixed concentration of the tracer. The $EC_{80}$ of the interaction was found using a 4-point logistic model:

$$fluorescence\ polarisation = unbound + \frac{(max - unbound) \times [PBP3]^{slope}}{[PBP3]^{slope} + EC80^{slope}}$$

Where *unbound* is the fluorescence polarisation value corresponding to the unbound tracer; *max* is the fluorescence polarisation value corresponding to maximally bound tracer and *slope* is the hill slope used.

In a competition fluorescence polarisation assay typically 12 concentrations of unlabelled peptide in a 2-fold titration from 5 μM to 2 nM were used to generate a dose-response curve against a fixed concentration of tracer and PBP3. The

Cheng-Prusoff equation was then used to find the $K_i$ using a value for the tracer $K_d$ of 2 nM:

$$K_i = \frac{IC_{50}}{1 + ([tracer]/K_d}$$

Where $IC_{50}$ is the inflection point of the dose-response curve.

Dotmatics (Dotmatics) workflows performing the above calculations were used to process the data. Values quoted are the geometric means of at least 2 repeats.

**Haemolysis**

Haemolysis assays were performed using human blood from healthy volunteers, with $K_2$-EDTA as the anticoagulant. Blood was preincubated at 37 °C for 15 mins with orbital shaking at 60rpm. In a 96-deep well plate, 4 µL compound at 100x final concentration was prepared, to which 200µL whole blood and either 200 µL phosphate-buffered saline (PBS) (test and negative control wells) or 200µL 2% (w/v) saponin in PBS (positive control wells) was added. Plates were incubated at 37°C for 10mins with orbital shaking at 60rpm, and then centrifuged at 500 xg for 5 mins at room temperature. 50 µL of supernatant was diluted 1:1 with PBS in a 96 well flat-bottom plate, and absorbance at A540nm was read with a SpectraMax plate reader. Absorbance values were normalised to the mean value for the 1% (w/v) saponin wells, to give a percentage haemolysis readout. Data are representative of 3 technical replicates.

**Cytotoxicity**

Cytotoxicity against HT1080, A549 and HepG2 cells was screened using the CellTiter-Glo luminescent cell viability assay kit (Promega). Cells were seeded at 10000 cells per well into 96-well plates (Corning #734-1660), with outer wells containing media only, and incubated for 24 hours at 37 °C with 5% $CO_2$. Test compounds were diluted to final concentration in cell culture media plus 0.5% (v/v) DMSO. All media was removed from the plate containing cells, and replaced with 100 µL of media containing test compounds or controls. Staurosporine (Sigma Aldrich) was used as a positive control, and DMSO (Sigma Aldrich) as a negative control. Plates were sealed with a breathable plate seal and incubated as before for 72 hours. Following incubation, 100 µL of CellTiter-Glo reagent (prepared according to manufacturer's instructions) was added to all wells. Plates were sealed with optically clear seals and incubated at room temperature for 10 minutes prior to luminescence reading using a LUM plus module in a Pherastar plate reader (BMG Labtech).

### Peptide QC data

#### Peptide 1

**Data file:** D:\Chemstation\1\Data\Peptides\4800-4899_BCY00012129_01 4840S1Cy1P1F2 Z (1) 2019-04-19 15-04-59.D

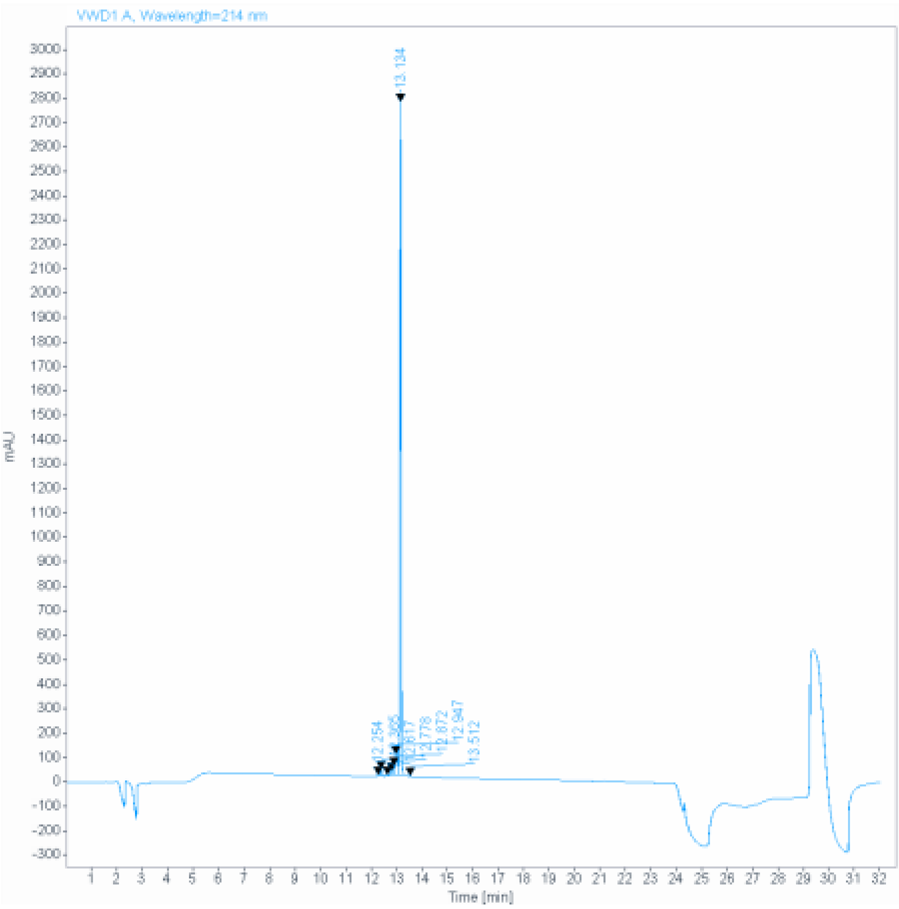

**Signal:** VWD1 A, Wavelength=214 nm

| RT [min] | Type | Width [min] | Area | Height | Area% | Name |
|---|---|---|---|---|---|---|
| 12.254 | BV | 0.0419 | 16.1391 | 5.7258 | 0.1683 | |
| 12.365 | VB | 0.0458 | 85.3088 | 28.6163 | 0.8894 | |
| 12.617 | BB | 0.0478 | 9.9602 | 2.9970 | 0.1038 | |
| 12.778 | BV E | 0.0537 | 66.6822 | 18.2018 | 0.6952 | |
| 12.872 | VV E | 0.0723 | 191.8802 | 39.4078 | 2.0005 | |
| 12.947 | VV E | 0.0816 | 541.4348 | 87.0055 | 5.6450 | |
| 13.134 | VB R | 0.0492 | 8669.7451 | 2757.6914 | 90.3903 | |
| 13.512 | BB | 0.0430 | 10.3085 | 3.8196 | 0.1075 | |
| | | Sum | 9591.4589 | | | |

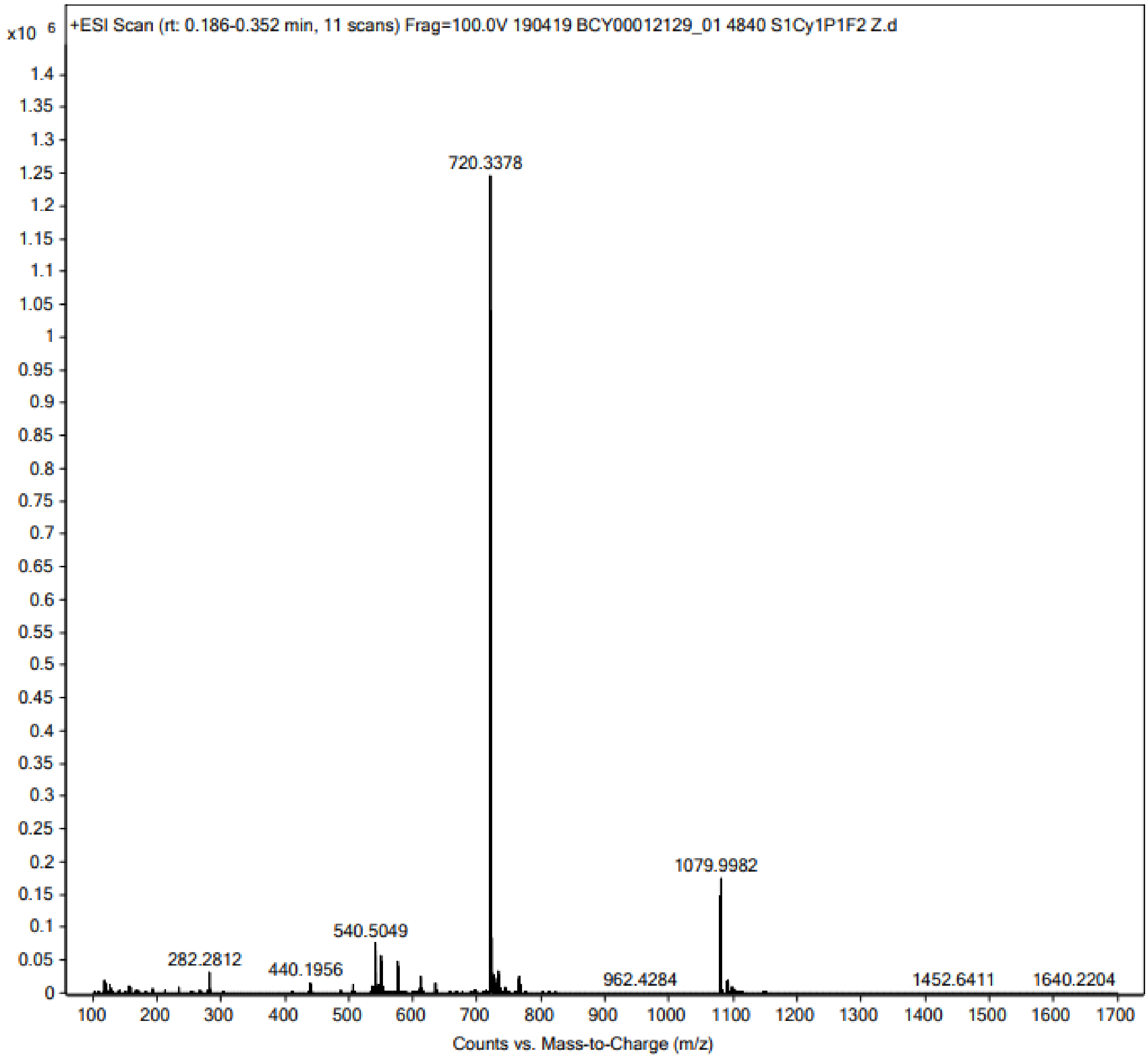

+ESI Scan (rt: 0.186-0.352 min, 11 scans) Frag=100.0V 190419 BCY00012129_01 4840 S1Cy1P1F2 Z.d
x10 6
720.3378
1079.9982
540.5049
282.2812
440.1956
962.4284
1452.6411
1640.2204
Counts vs. Mass-to-Charge (m/z)

#### Peptide 2

**Data file:** D:\Chemstation\1\Data\Peptides_facon\7000-7999
_BCY00012130_05_7557_A_zHPLC 16-07-49.D

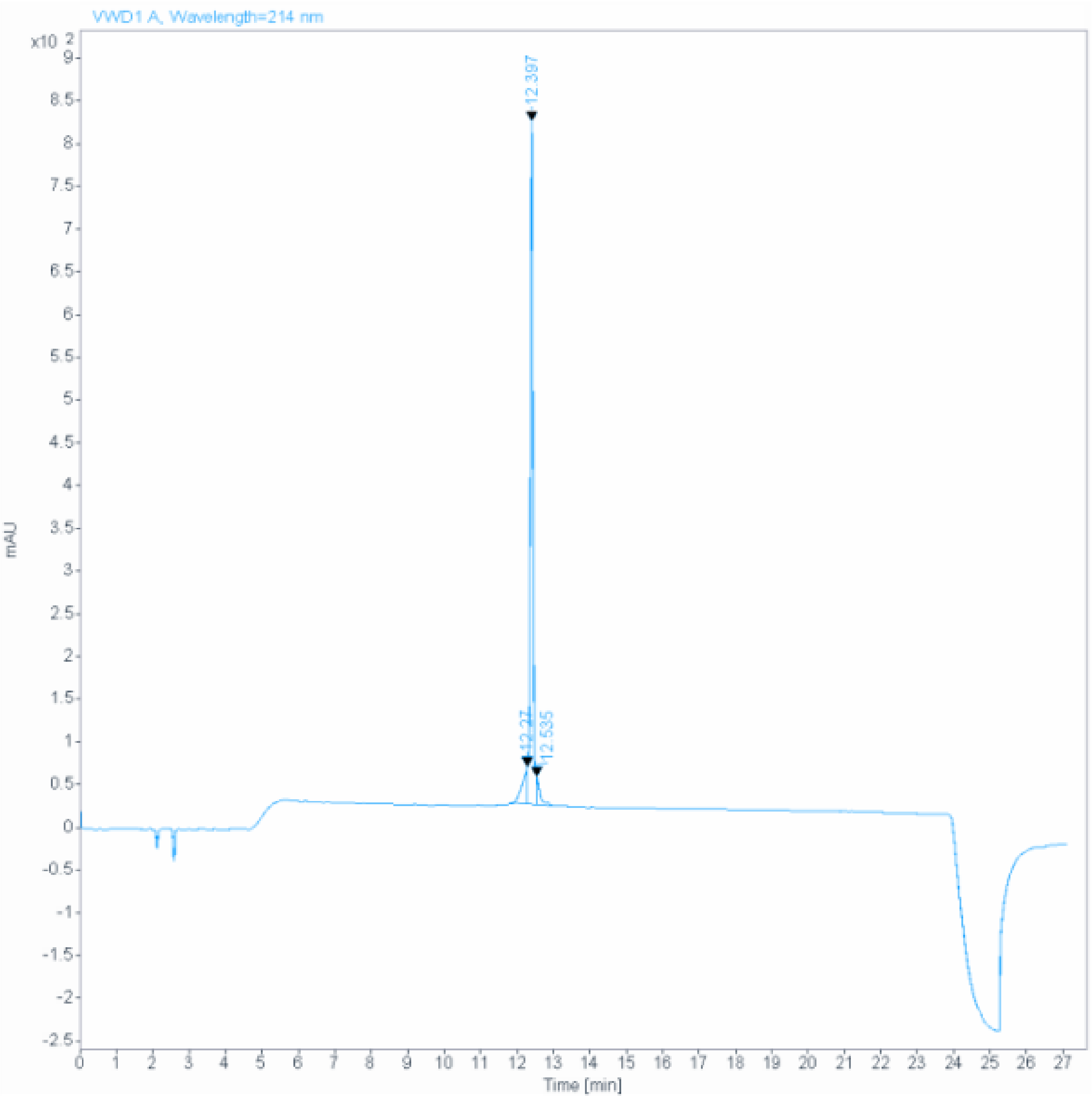

**Signal:** VWD1 A, Wavelength=214 nm

| RT [min] | Type | Width [min] | Area | Height | Area% | Name |
|---|---|---|---|---|---|---|
| 12.270 | MF | 0.1590 | 424.4482 | 44.4815 | 8.3924 | |
| 12.397 | MF | 0.0919 | 4412.6670 | 799.9210 | 87.2493 | |
| 12.535 | FM | 0.1100 | 220.4254 | 33.3902 | 4.3584 | |
| | | Sum | 5057.5407 | | | |

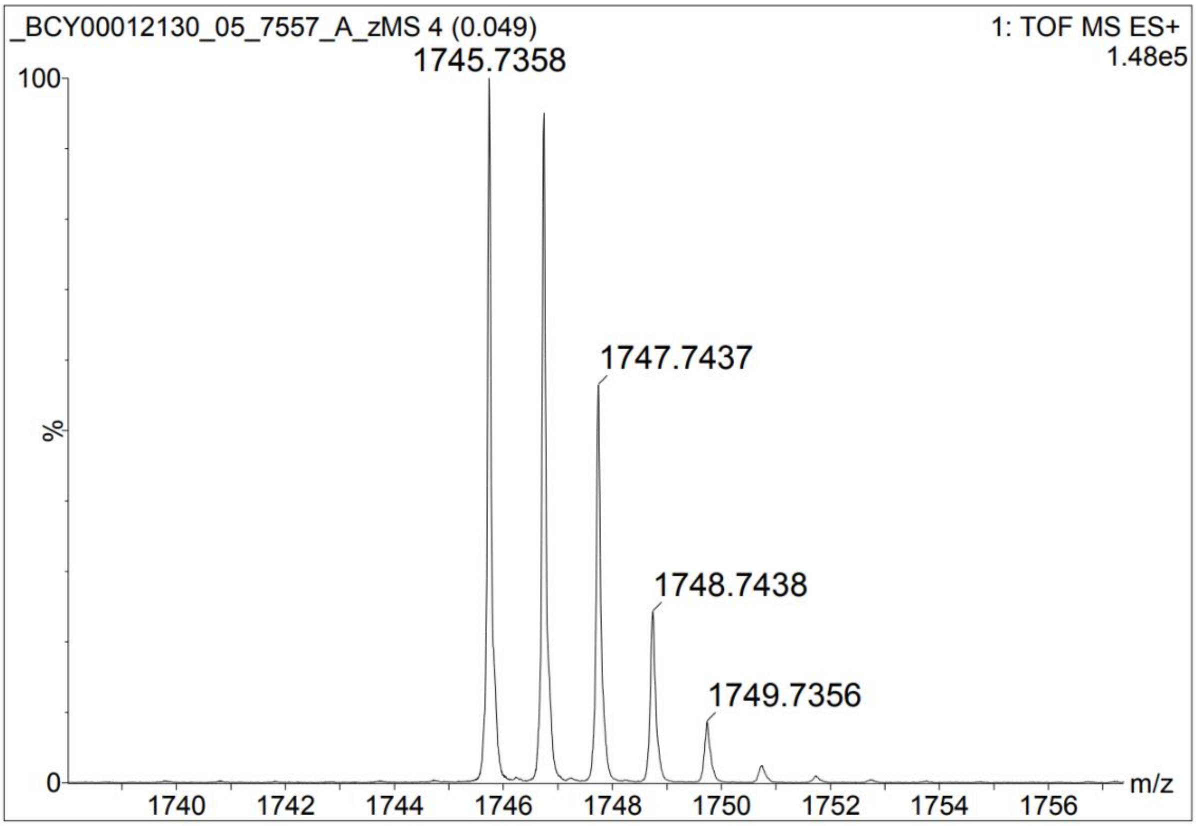
_BCY00012130_05_7557_A_zMS 4 (0.049)
1: TOF MS ES+
1.48e5
1745.7358
1747.7437
1748.7438
1749.7356
100
%
0
1740
1742
1744
1746
1748
1750
1752
1754
1756
m/z

#### Peptide 3

**Data file:** C:\Chem32\1\Data\Peptides\4800-4899_BCY00012131_01 4867 S1Cy1P1F1Z 2019-05-20 18-18-37.D

**Sample name:** _BCY00012131_01 4867 S1Cy1P1F1Z

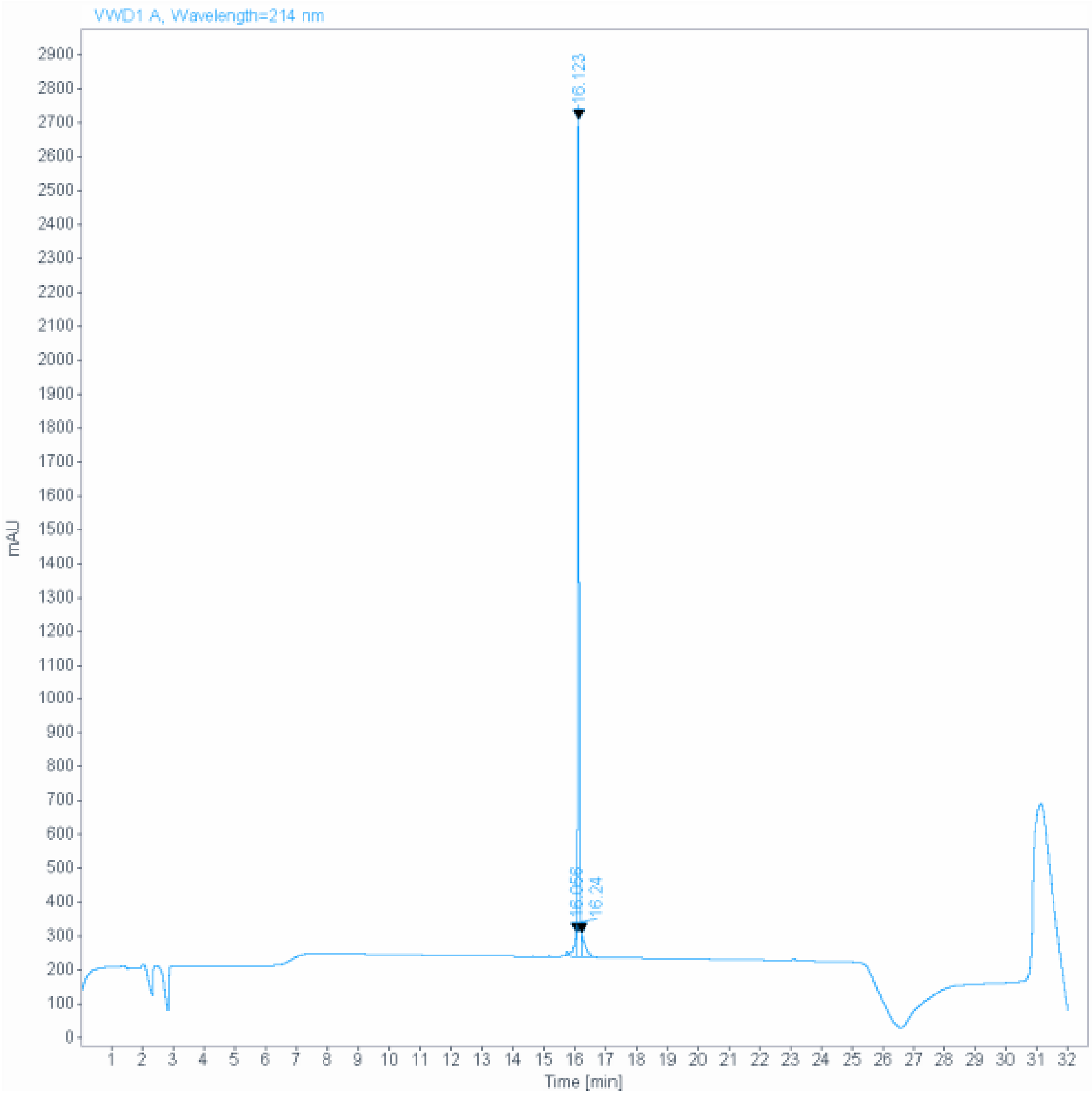

**Signal:** VWD1 A, Wavelength=214 nm

| RT [min] | Type | Width [min] | Area | Height | Area% | Name |
|---|---|---|---|---|---|---|
| 16.056 | MF | 0.1077 | 447.4658 | 69.2520 | 4.6339 | |
| 16.123 | MF | 0.0587 | 8704.0625 | 2472.5249 | 90.1375 | |
| 16.240 | FM | 0.1221 | 504.8945 | 68.8973 | 5.2286 | |
| | | Sum | 9656.4228 | | | |

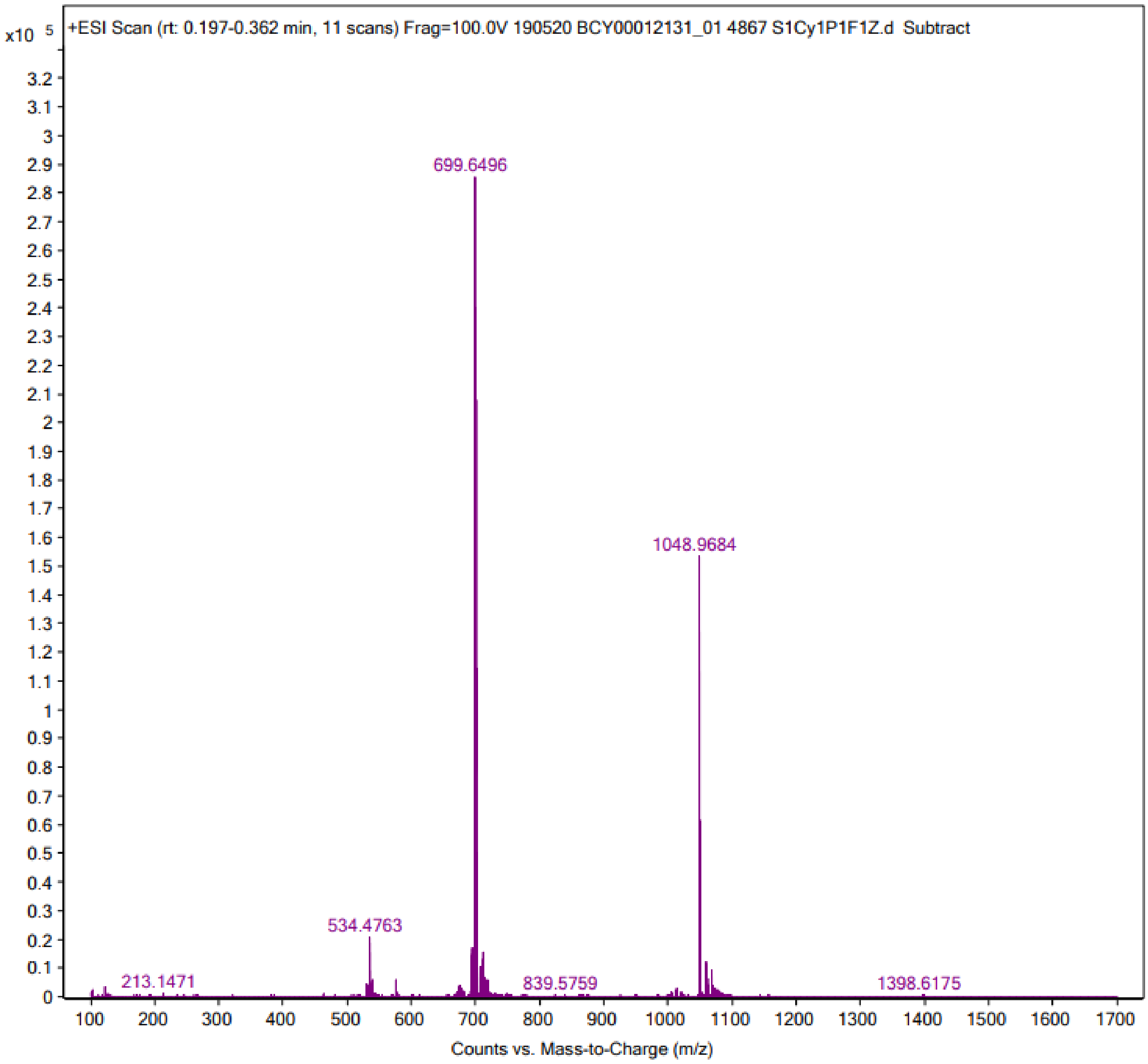
x10 5
+ESI Scan (rt: 0.197-0.362 min, 11 scans) Frag=100.0V 190520 BCY00012131_01 4867 S1Cy1P1F1Z.d Subtract
699.6496
1048.9684
534.4763
213.1471
839.5759
1398.6175
Counts vs. Mass-to-Charge (m/z)

205

#### Peptide 4

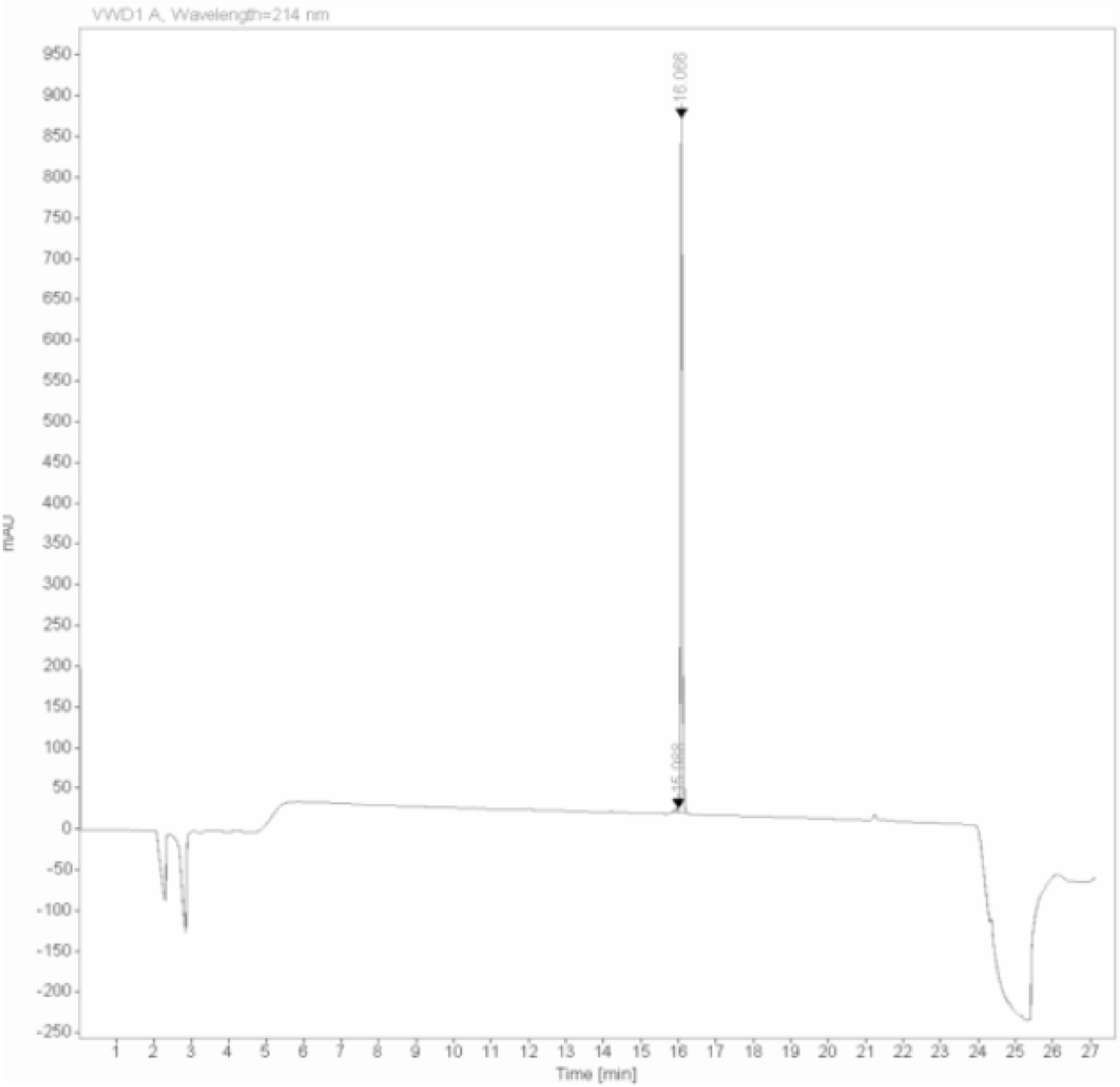

Signal: VWD1 A, Wavelength=214 nm

| RT [min] | Type | Width [min] | Area | Height | Area% | Name |
|---|---|---|---|---|---|---|
| 15.988 | MF | 0.0900 | 29.1654 | 5.4000 | 0.9969 | |
| 16.066 | FM | 0.0566 | 2896.3469 | 853.4639 | 99.0031 | |
| | | Sum | 2925.5123 | | | |

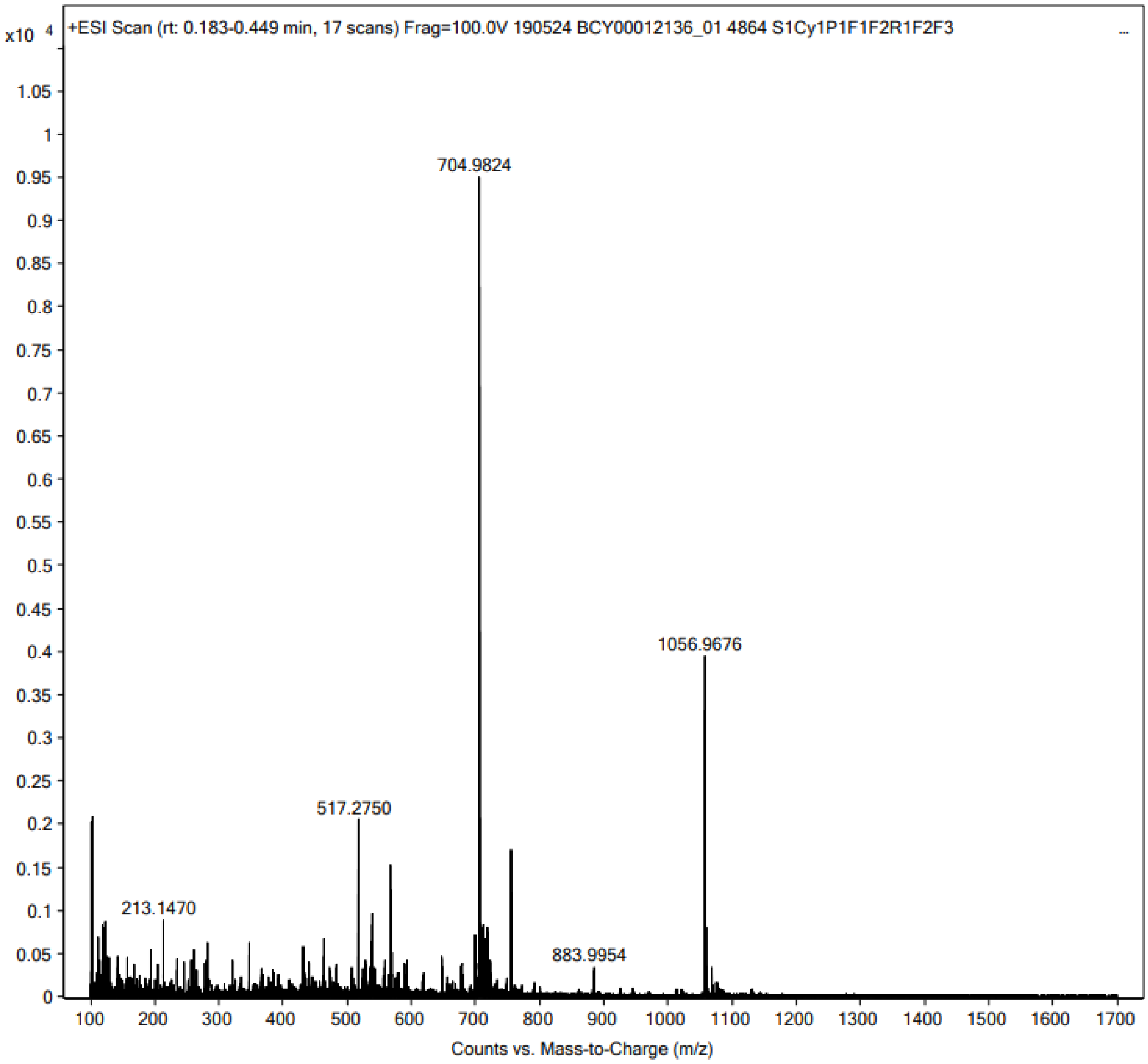
x10 4
+ESI Scan (rt: 0.183-0.449 min, 17 scans) Frag=100.0V 190524 BCY00012136_01 4864 S1Cy1P1F1F2R1F2F3
704.9824
1056.9676
517.2750
213.1470
883.9954
Counts vs. Mass-to-Charge (m/z)

208

#### Peptide 5

**Data file:** C:\Chem32\1\Data\Peptides\4800-4899_BCY00012132_01 4868 S1Cy1P1F1Z 2019-05-20 18-18-50.D

**Sample name:** _BCY00012132_01 4868 S1Cy1P1F1Z

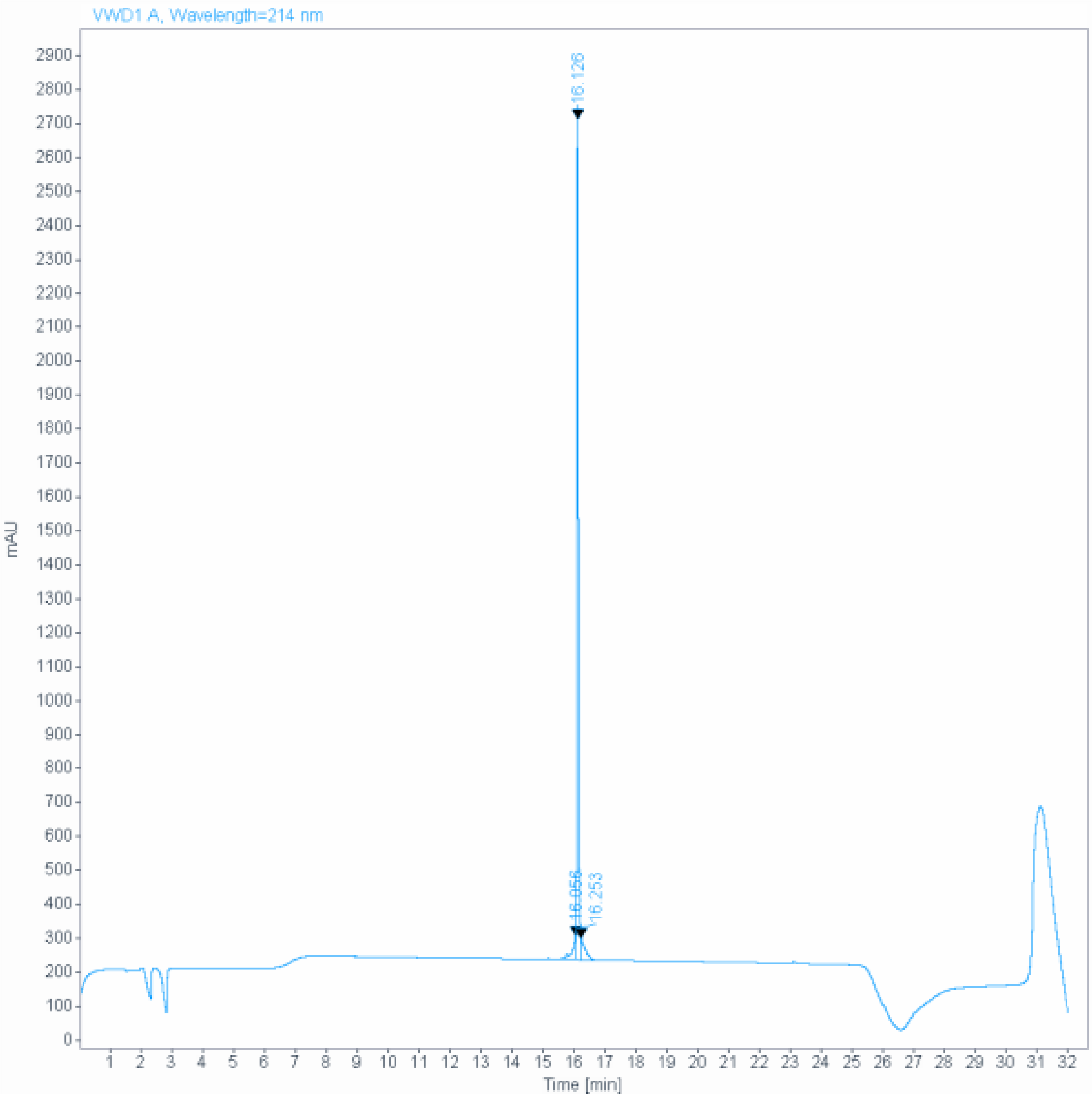

**Signal:** VWD1 A, Wavelength=214 nm

| RT [min] | Type | Width [min] | Area | Height | Area% | Name |
|---|---|---|---|---|---|---|
| 16.056 | MF | 0.1088 | 448.8487 | 68.7475 | 4.6234 | |
| 16.126 | MF | 0.0591 | 8772.6650 | 2475.7566 | 90.3627 | |
| 16.253 | FM | 0.1280 | 486.7630 | 63.3937 | 5.0139 | |
| | | Sum | 9708.2768 | | | |

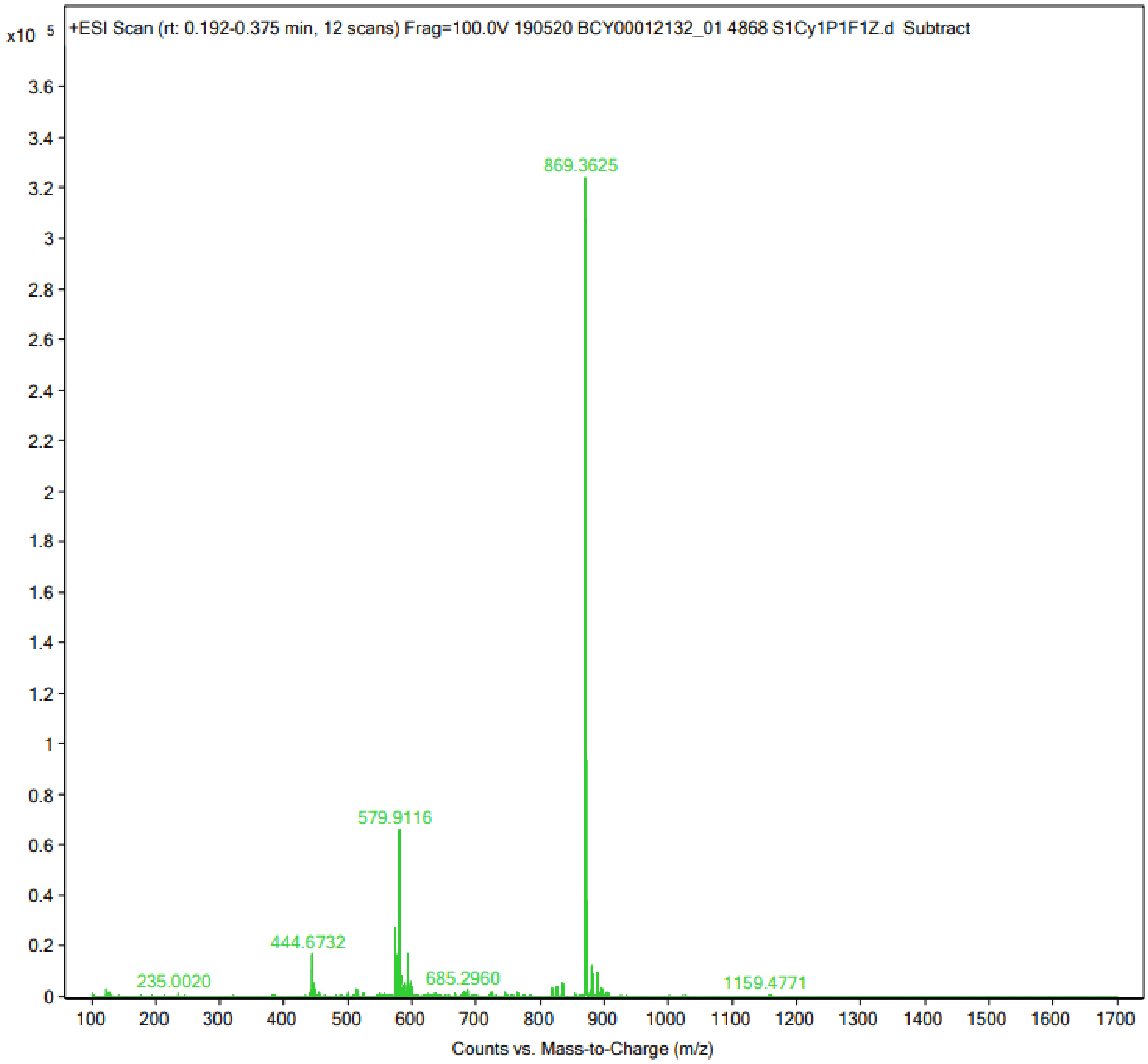

x10 5
+ESI Scan (rt: 0.192-0.375 min, 12 scans) Frag=100.0V 190520 BCY00012132_01 4868 S1Cy1P1F1Z.d Subtract
869.3625
579.9116
444.6732
235.0020
685.2960
1159.4771
Counts vs. Mass-to-Charge (m/z)

211

#### Peptide 6

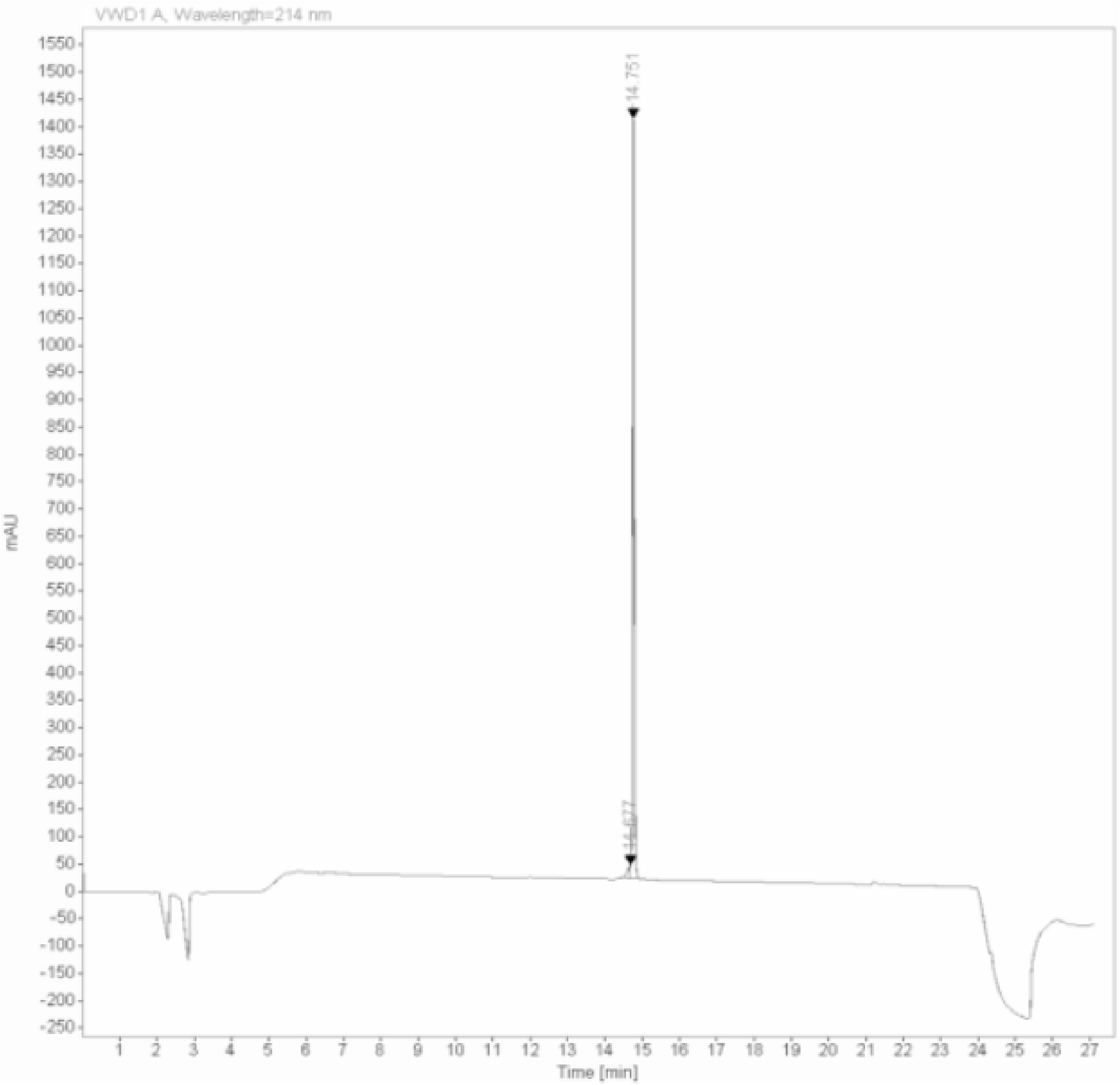

Signal: VWD1 A, Wavelength=214 nm

| RT [min] | Type | Width [min] | Area | Height | Area% | Name |
|---|---|---|---|---|---|---|
| 14.677 | MF | 0.0886 | 130.1320 | 24.4681 | 2.6617 | |
| 14.751 | FM | 0.0569 | 4758.9902 | 1392.9545 | 97.3383 | |
| | | Sum | 4889.1222 | | | |

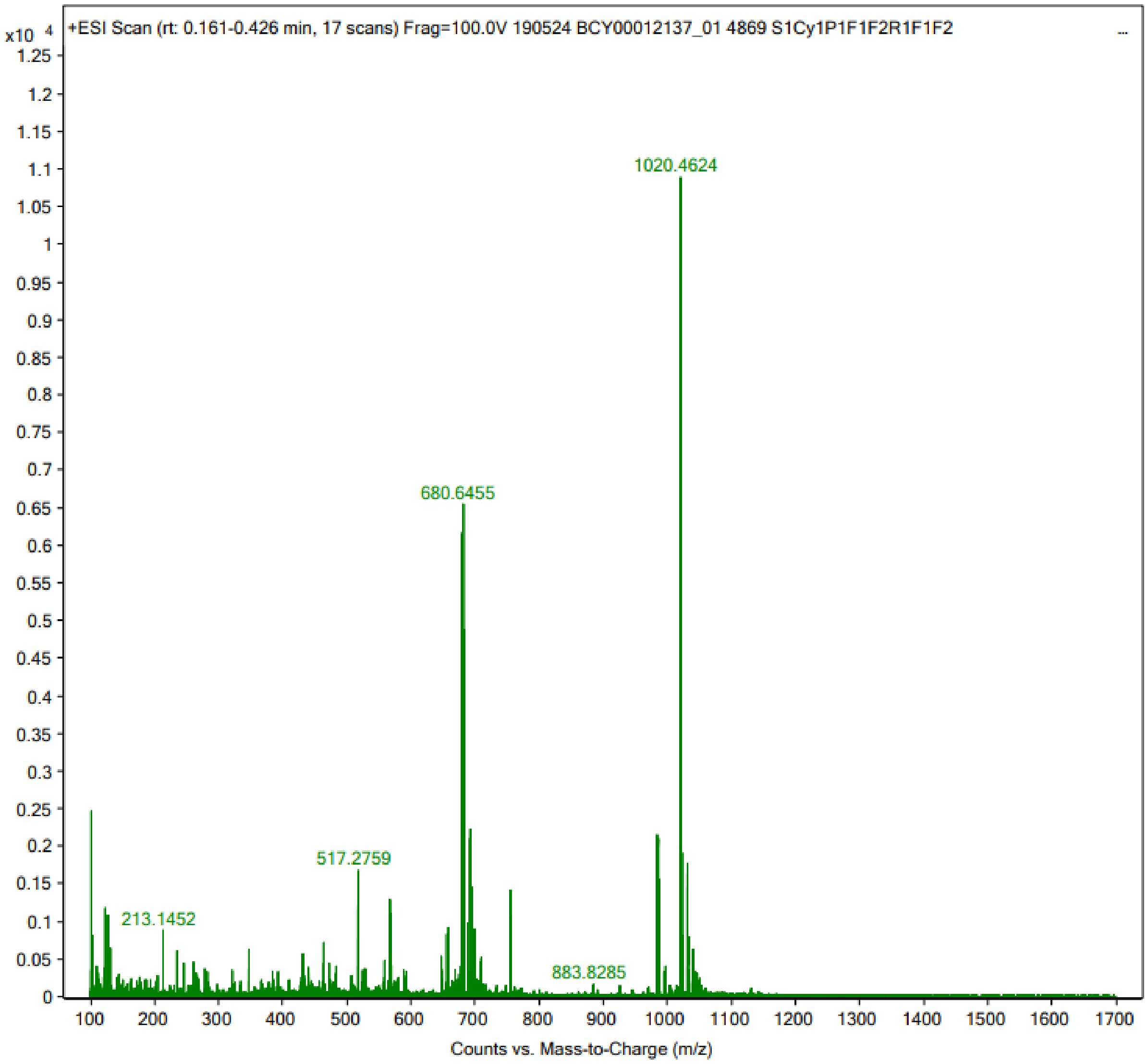
x10 4
+ESI Scan (rt: 0.161-0.426 min, 17 scans) Frag=100.0V 190524 BCY00012137_01 4869 S1Cy1P1F1F2R1F1F2
1020.4624
680.6455
517.2759
213.1452
883.8285
Counts vs. Mass-to-Charge (m/z)

214

**Peptide 7**

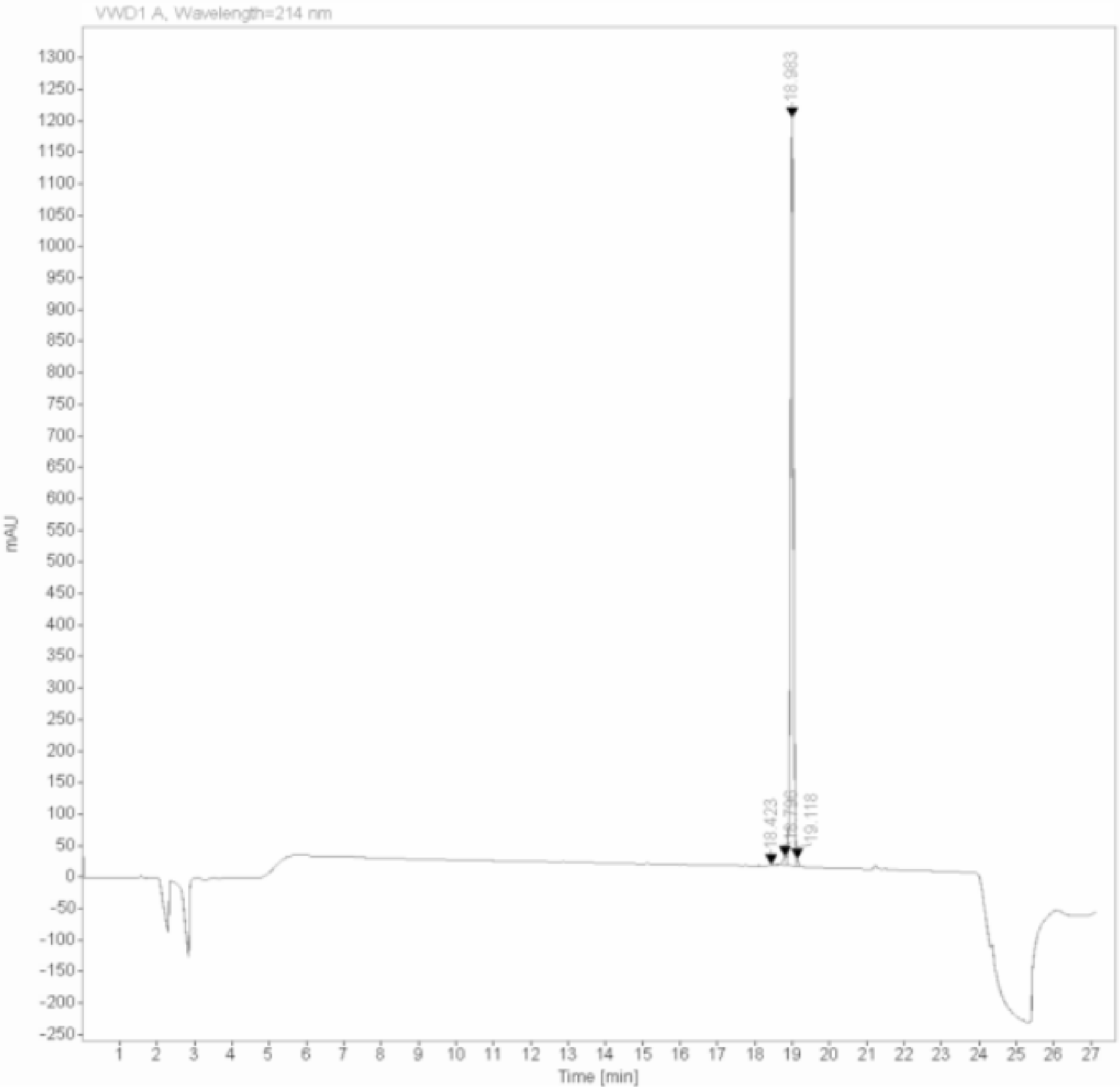

| Signal: | VWD1 A, Wavelength=214 nm | | | | | |
|---|---|---|---|---|---|---|
| RT [min] | Type | Width [min] | Area | Height | Area% | Name |
| 18.423 | MM | 0.0547 | 8.2297 | 2.5094 | 0.1208 | |
| 18.796 | MF | 0.1131 | 105.5613 | 15.5575 | 1.5490 | |
| 18.983 | MF | 0.0934 | 6663.8960 | 1188.6801 | 97.7828 | |
| 19.118 | FM | 0.0474 | 37.3108 | 13.1298 | 0.5475 | |
| | | Sum | 6814.9978 | | | |

217

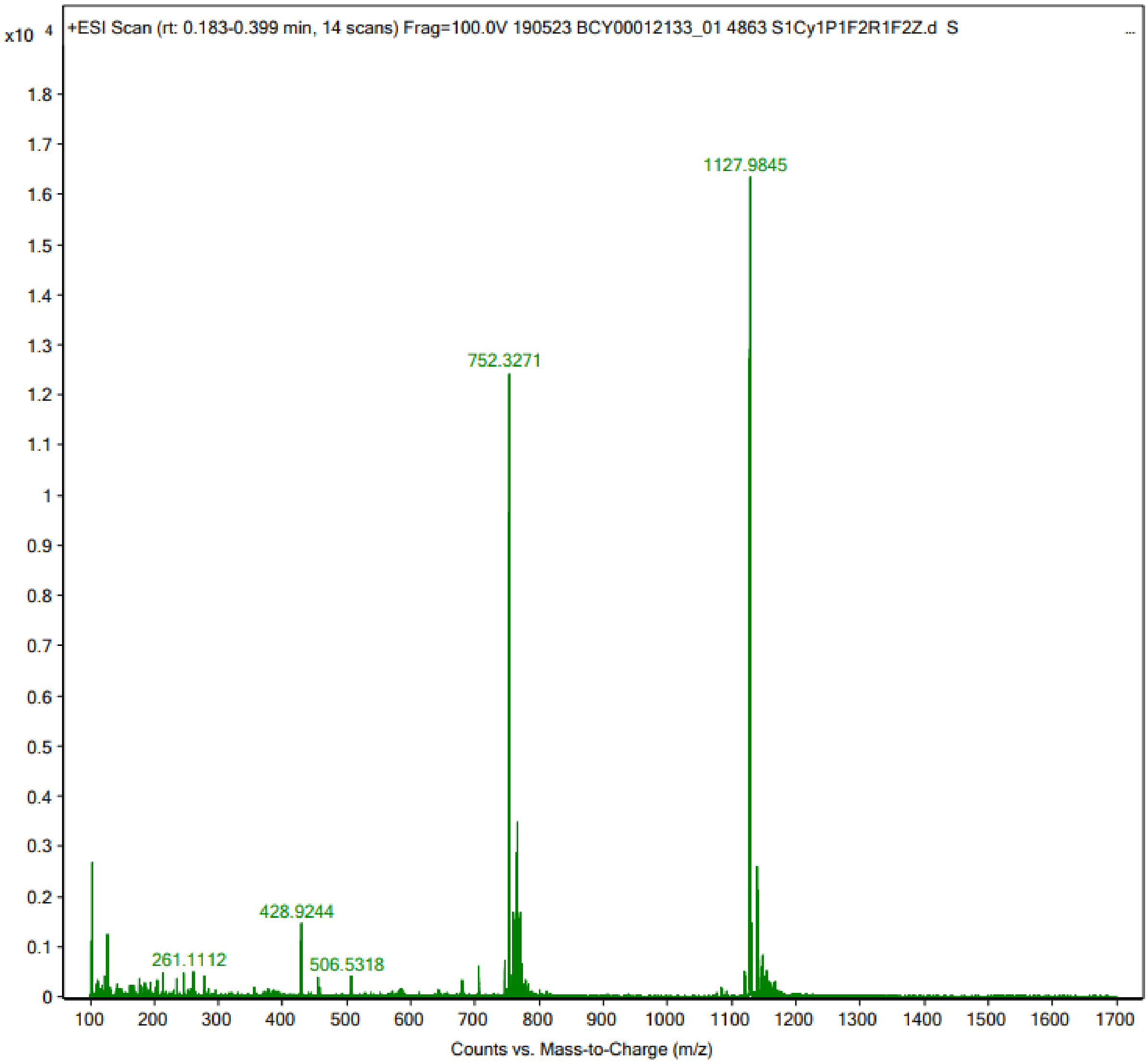

x10 4
+ESI Scan (rt: 0.183-0.399 min, 14 scans) Frag=100.0V 190523 BCY00012133_01 4863 S1Cy1P1F2R1F2Z.d S
1127.9845
752.3271
428.9244
261.1112
506.5318
1.8
1.7
1.6
1.5
1.4
1.3
1.2
1.1
1
0.9
0.8
0.7
0.6
0.5
0.4
0.3
0.2
0.1
0
100 200 300 400 500 600 700 800 900 1000 1100 1200 1300 1400 1500 1600 1700
Counts vs. Mass-to-Charge (m/z)

#### Peptide 8

**Data file:** Y:\Cunegonde\Chem32\1\Data\Peptides\4800-4899_BCY-00012125_01 4843 S1Cy1P1F1Z 2019-05-03 18-08-28.D

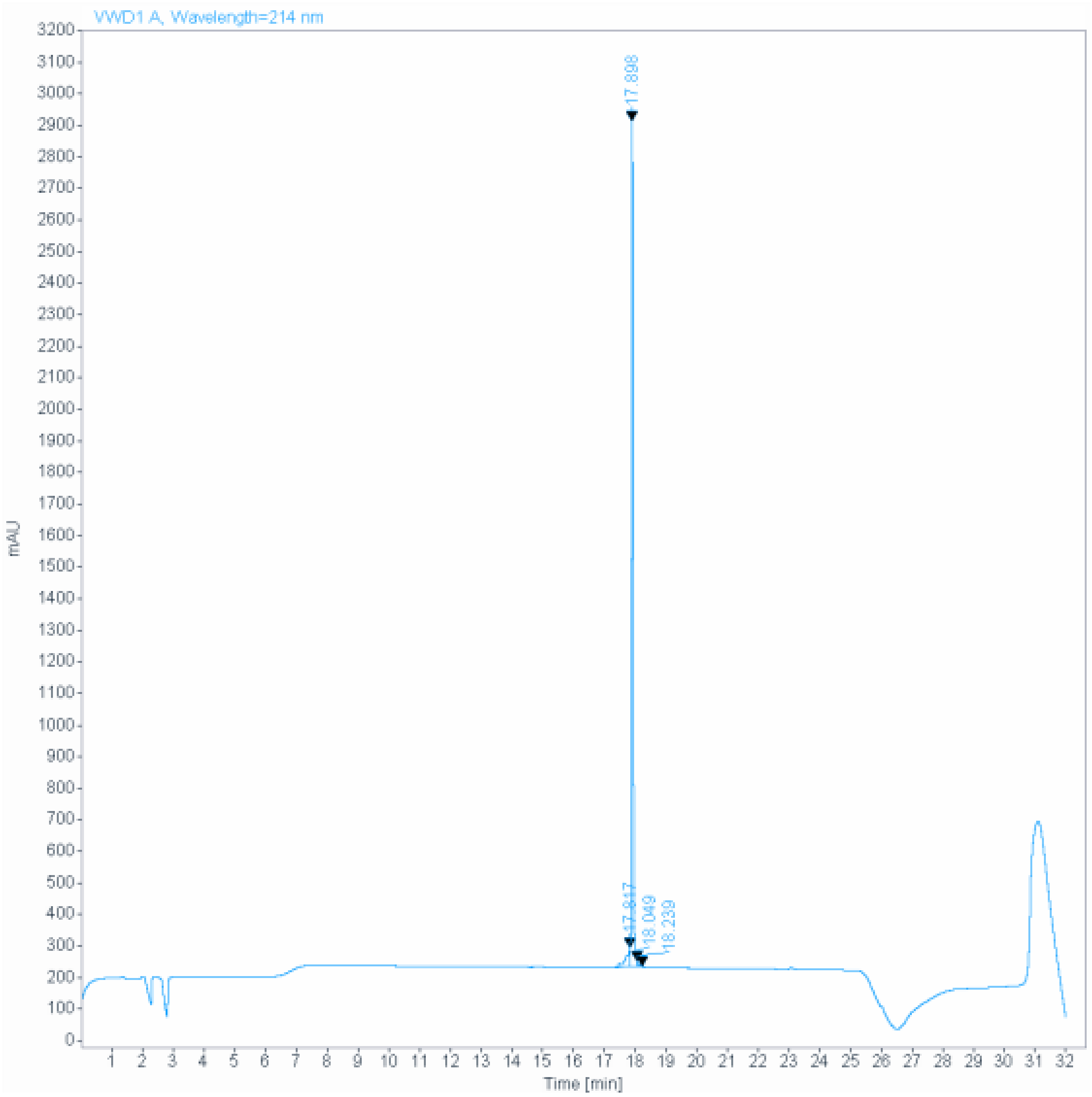

**Signal:** VWD1 A, Wavelength=214 nm

| RT [min] | Type | Width [min] | Area | Height | Area% | Name |
|---|---|---|---|---|---|---|
| 17.817 | MF | 0.1405 | 490.2820 | 58.1721 | 4.9663 | |
| 17.898 | MF | 0.0579 | 9314.9258 | 2680.0886 | 94.3553 | |
| 18.049 | FM | 0.0592 | 62.0055 | 17.4606 | 0.6281 | |
| 18.239 | MM | 0.0497 | 4.9650 | 1.6666 | 0.0503 | |
| | | Sum | 9872.1783 | | | |

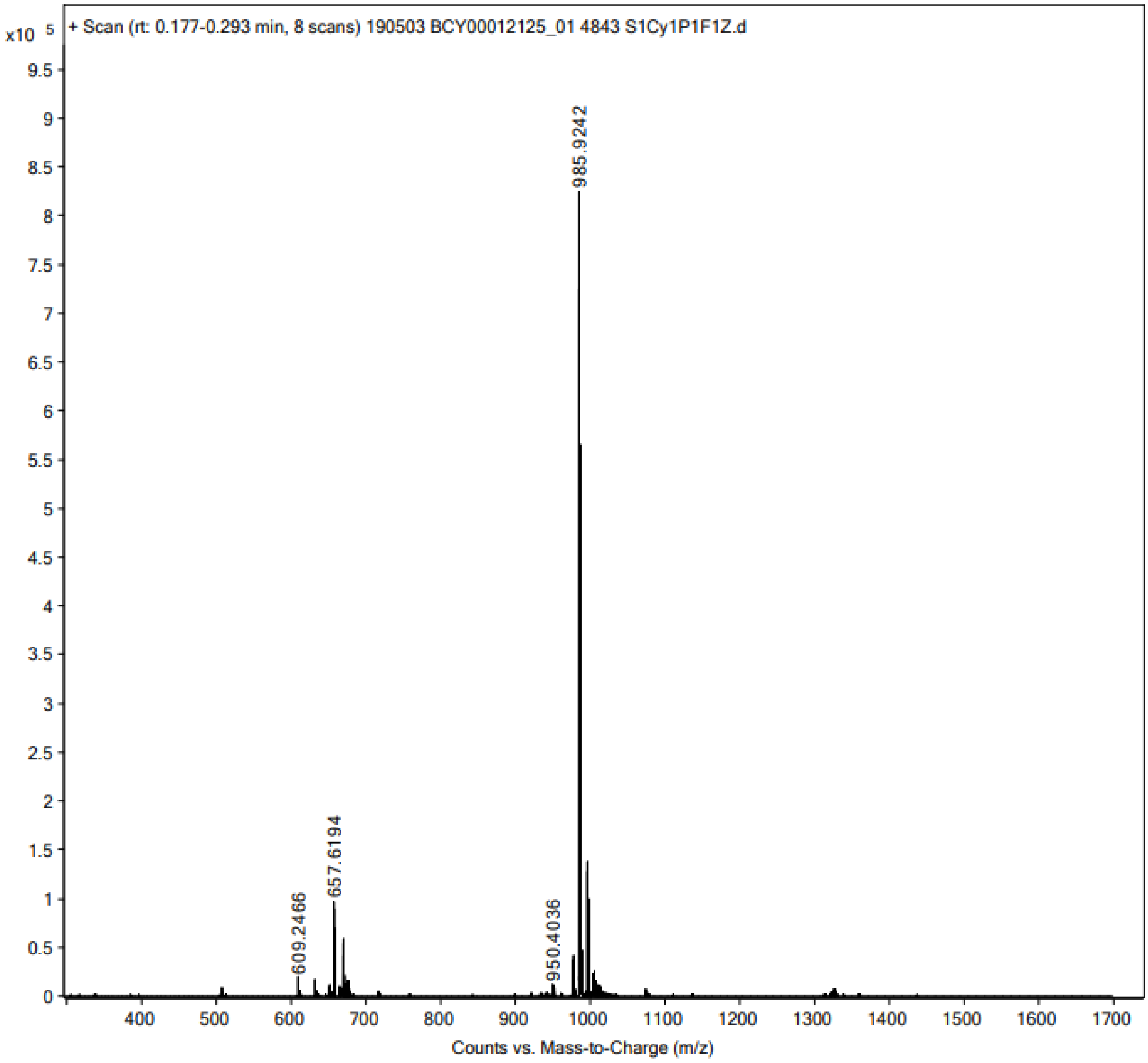
x10 5
+ Scan (rt: 0.177-0.293 min, 8 scans) 190503 BCY00012125_01 4843 S1Cy1P1F1Z.d
985.9242
657.6194
609.2466
950.4036
Counts vs. Mass-to-Charge (m/z)

220

221 **Peptide 9**

**Data file:** Y:\Cunegonde\Chem32\1\Data\Peptides\4800-4899_BCY-00012124_01 4842 S1Cy1P1F1Z 2019-05-03 18-07-57.D

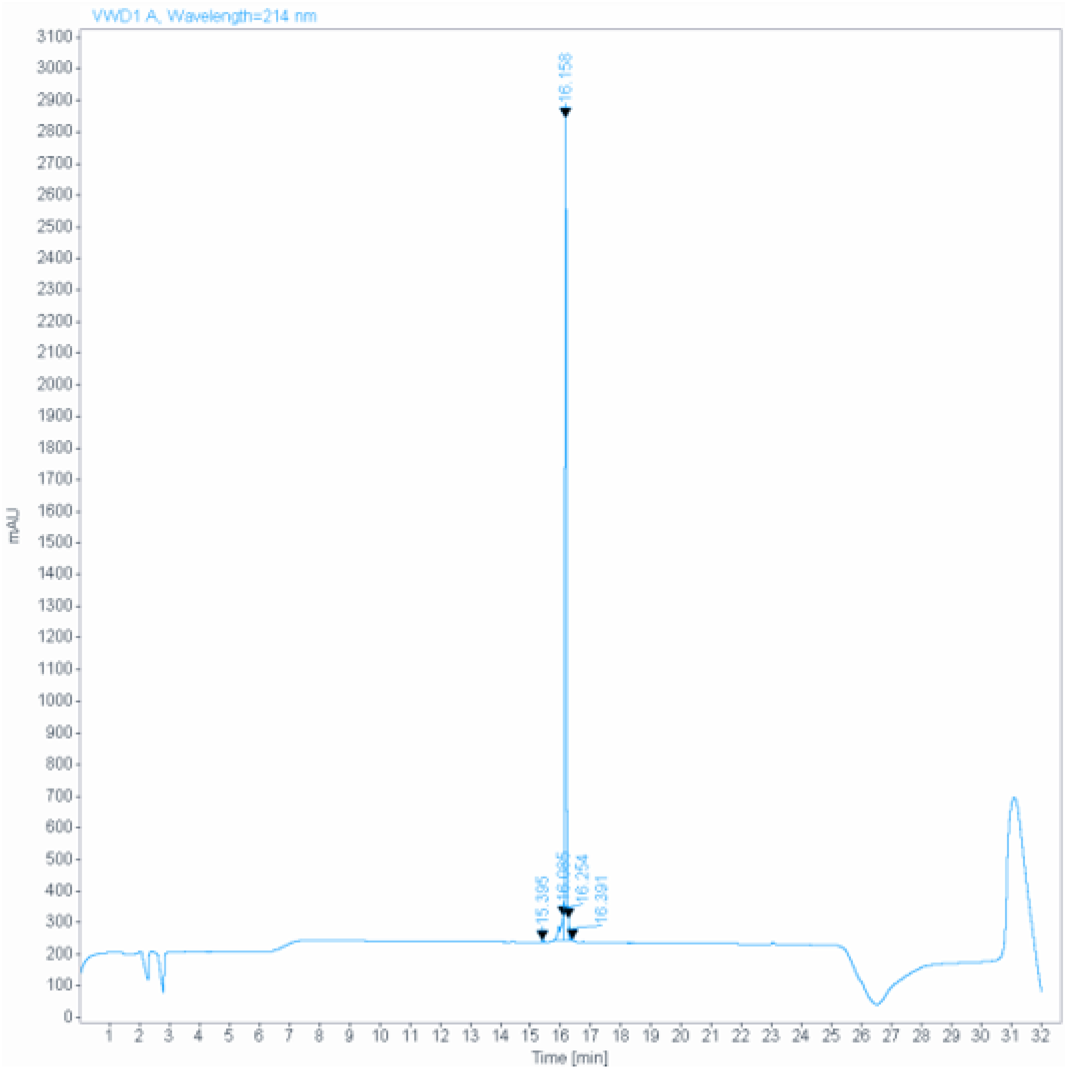

**Signal:** VWD1 A, Wavelength=214 nm

| RT [min] | Type | Width [min] | Area | Height | Area% | Name |
|---|---|---|---|---|---|---|
| 15.395 | VV R | 0.0632 | 15.6337 | 3.7458 | 0.1671 | |
| 16.085 | MF | 0.1321 | 591.7968 | 74.6883 | 6.3262 | |
| 16.158 | MF | 0.0553 | 8624.3271 | 2601.4370 | 92.1927 | |
| 16.254 | FM | 0.0269 | 109.8465 | 68.1846 | 1.1742 | |
| 16.391 | MM | 0.0430 | 13.0673 | 5.0638 | 0.1397 | |
| | | Sum | 9354.6715 | | | |

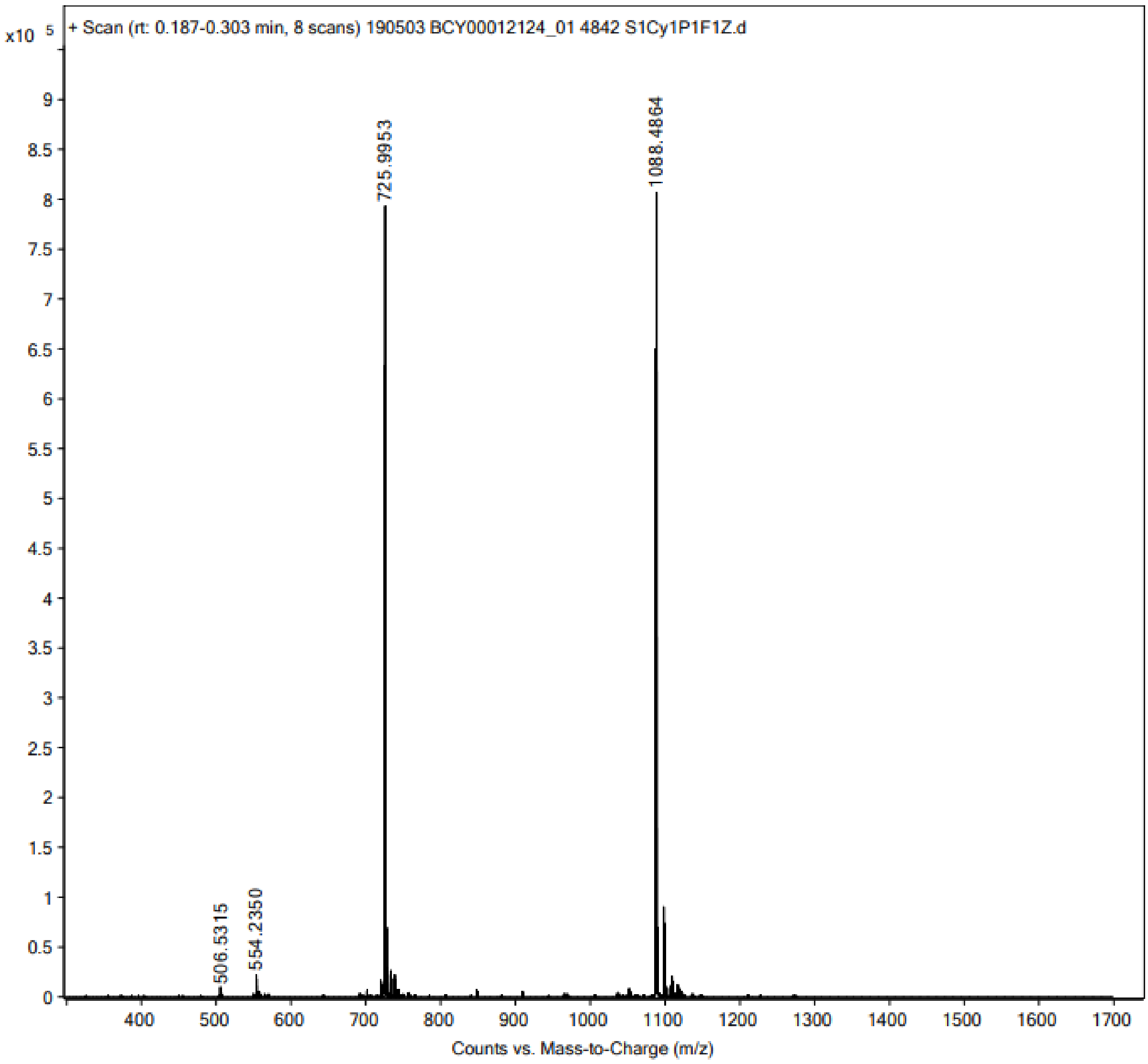
x10 5
+ Scan (rt: 0.187-0.303 min, 8 scans) 190503 BCY00012124_01 4842 S1Cy1P1F1Z.d
725.9953
1088.4864
506.5315
554.2350
Counts vs. Mass-to-Charge (m/z)

222

#### Peptide 10

**Data file:** C:\Chem32\1\Data\Peptides\4800-4899_BCY00012135_01 4865 S1Cy1P1F1F2Z 2019-05-20 18-18-18.D

**Sample name:** _BCY00012135_01 4865 S1Cy1P1F1F2Z

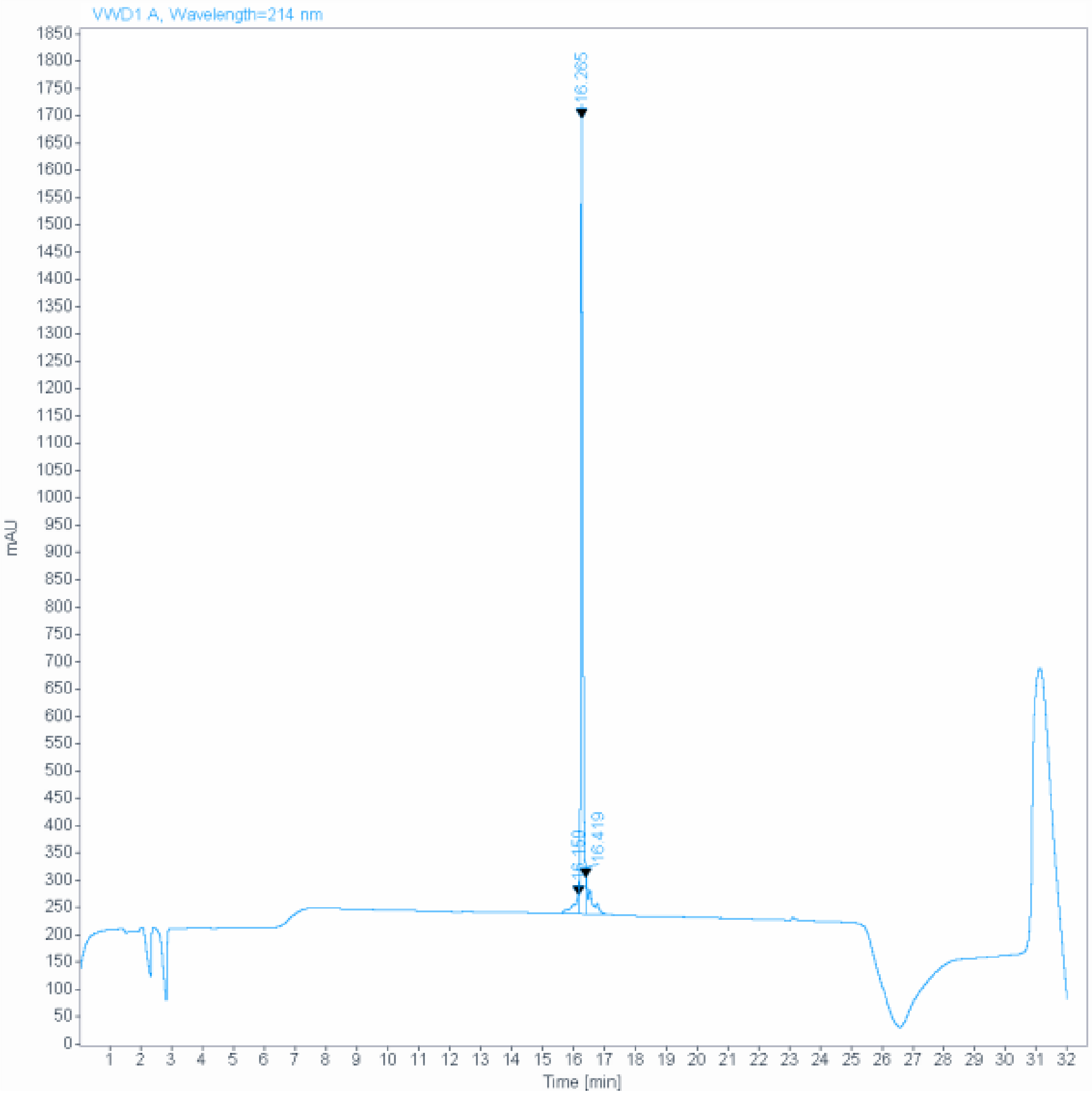

**Signal:** VWD1 A, Wavelength=214 nm

| RT [min] | Type | Width [min] | Area | Height | Area% | Name |
|---|---|---|---|---|---|---|
| 16.159 | MF | 0.1862 | 373.6817 | 33.4513 | 4.7803 | |
| 16.265 | MF | 0.0773 | 6764.0708 | 1458.1731 | 86.5283 | |
| 16.419 | FM | 0.1727 | 679.4210 | 65.5497 | 8.6914 | |
| | | Sum | 7817.1735 | | | |

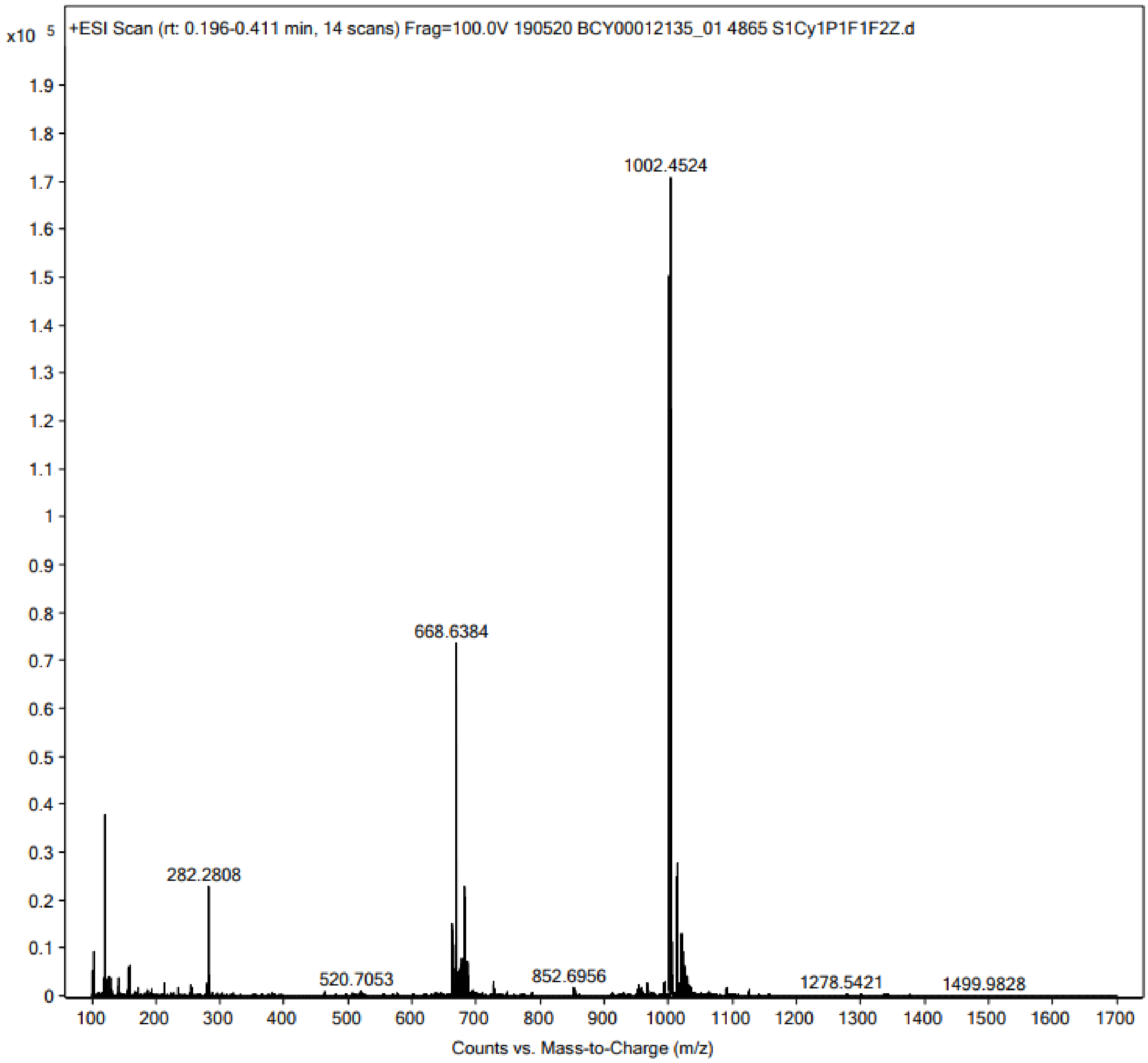
x10 5
+ESI Scan (rt: 0.196-0.411 min, 14 scans) Frag=100.0V 190520 BCY00012135_01 4865 S1Cy1P1F1F2Z.d
1002.4524
668.6384
282.2808
520.7053
852.6956
1278.5421
1499.9828
Counts vs. Mass-to-Charge (m/z)

#### Peptide 11

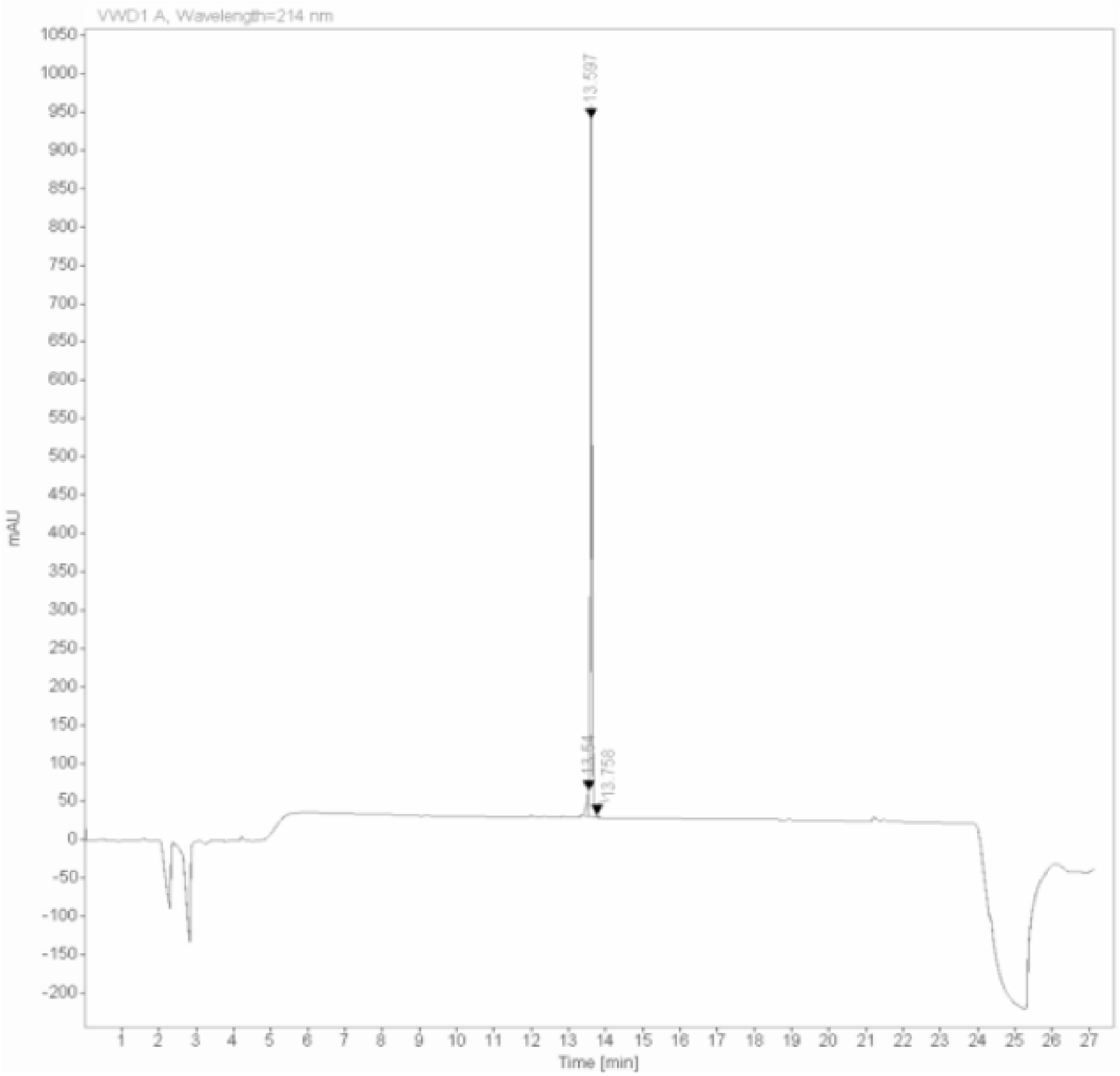

Signal: VWD1 A, Wavelength=214 nm

| RT [min] | Type | Width [min] | Area | Height | Area% | Name |
|---|---|---|---|---|---|---|
| 13.540 | MF | 0.0685 | 143.5043 | 34.9315 | 4.8870 | |
| 13.597 | FM | 0.0507 | 2773.6589 | 912.5562 | 94.4570 | |
| 13.758 | FM | 0.0673 | 19.2617 | 4.7713 | 0.6560 | |
| | | Sum | 2936.4249 | | | |

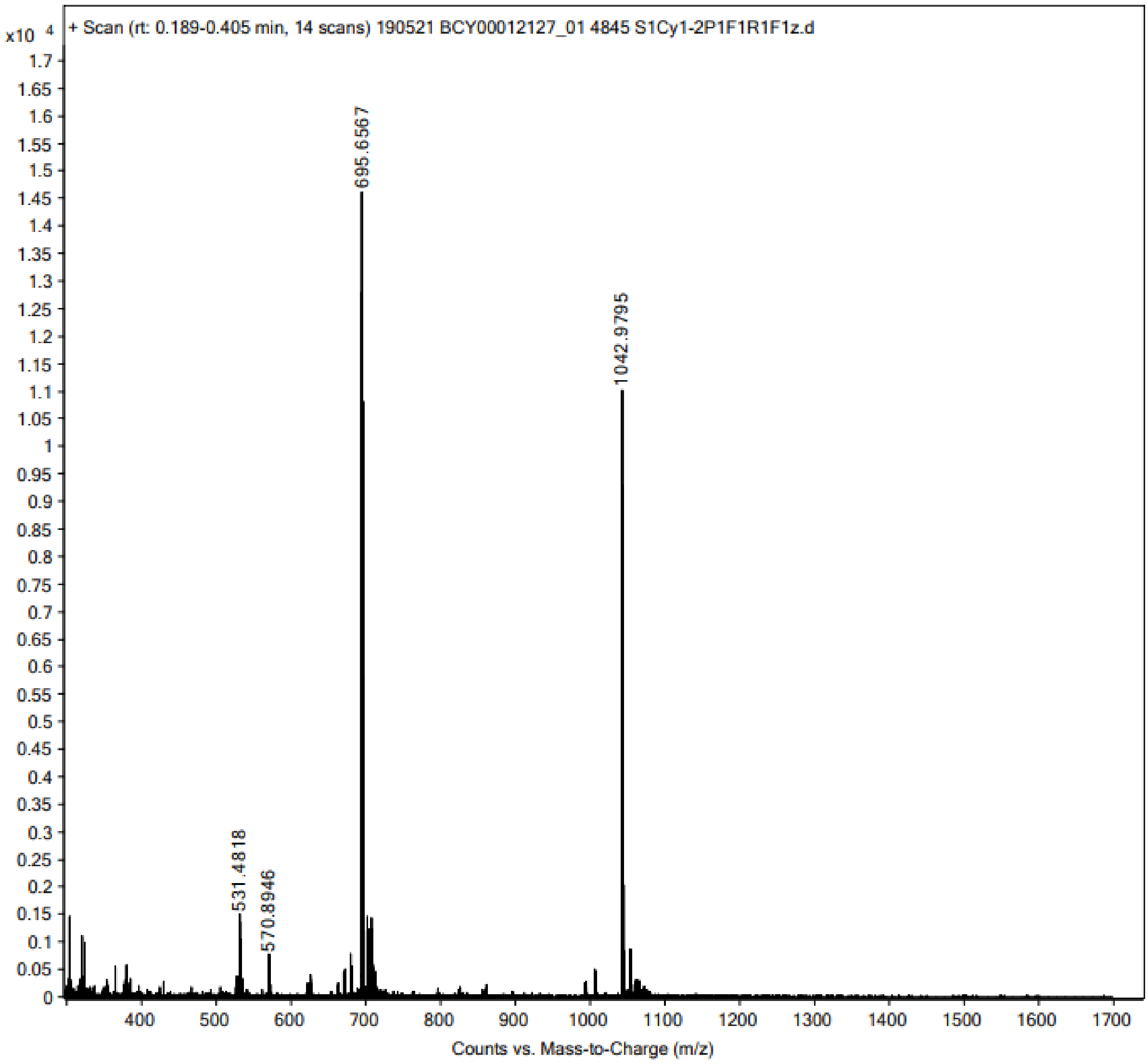

x10 4
+ Scan (rt: 0.189-0.405 min, 14 scans) 190521 BCY00012127_01 4845 S1Cy1-2P1F1R1F1z.d
695.6567
1042.9795
531.4818
570.8946
Counts vs. Mass-to-Charge (m/z)

229

#### Peptide 12

**Data file:** C:\Chem32\1\Data\Peptides\4800-4899_BCY00012134_01 4861 S1Cy1P1F1Z 2019-05-20 18-17-43.D

**Sample name:** _BCY00012134_01 4861 S1Cy1P1F1Z

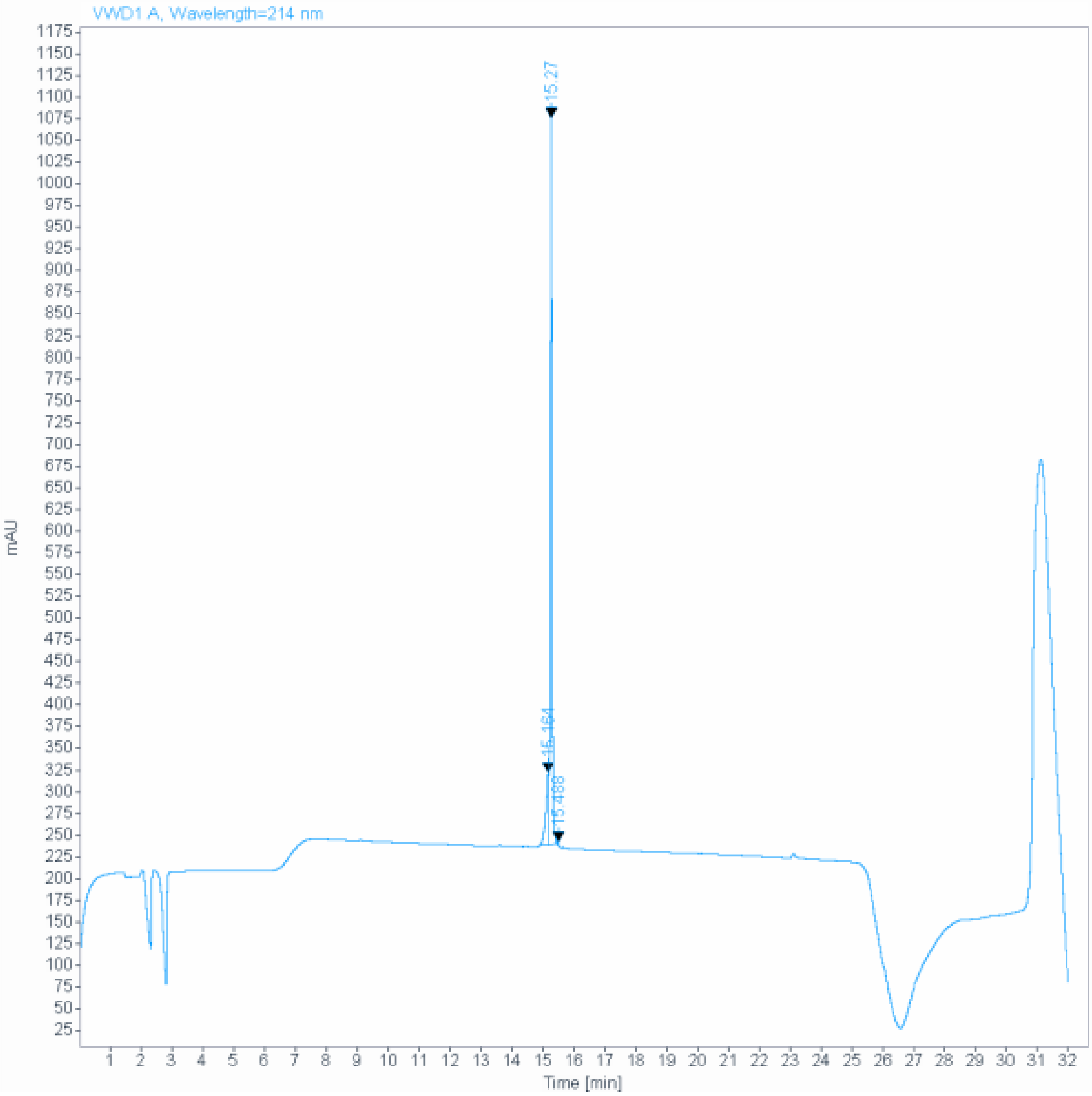

**Signal:** VWD1 A, Wavelength=214 nm

| RT [min] | Type | Width [min] | Area | Height | Area% | Name |
|---|---|---|---|---|---|---|
| 15.164 | MF | 0.1023 | 511.0786 | 83.2671 | 14.0753 | |
| 15.270 | FM | 0.0618 | 3108.8955 | 838.0609 | 85.6204 | |
| 15.488 | MM | 0.0458 | 11.0467 | 4.0182 | 0.3042 | |
| | | Sum | 3631.0207 | | | |

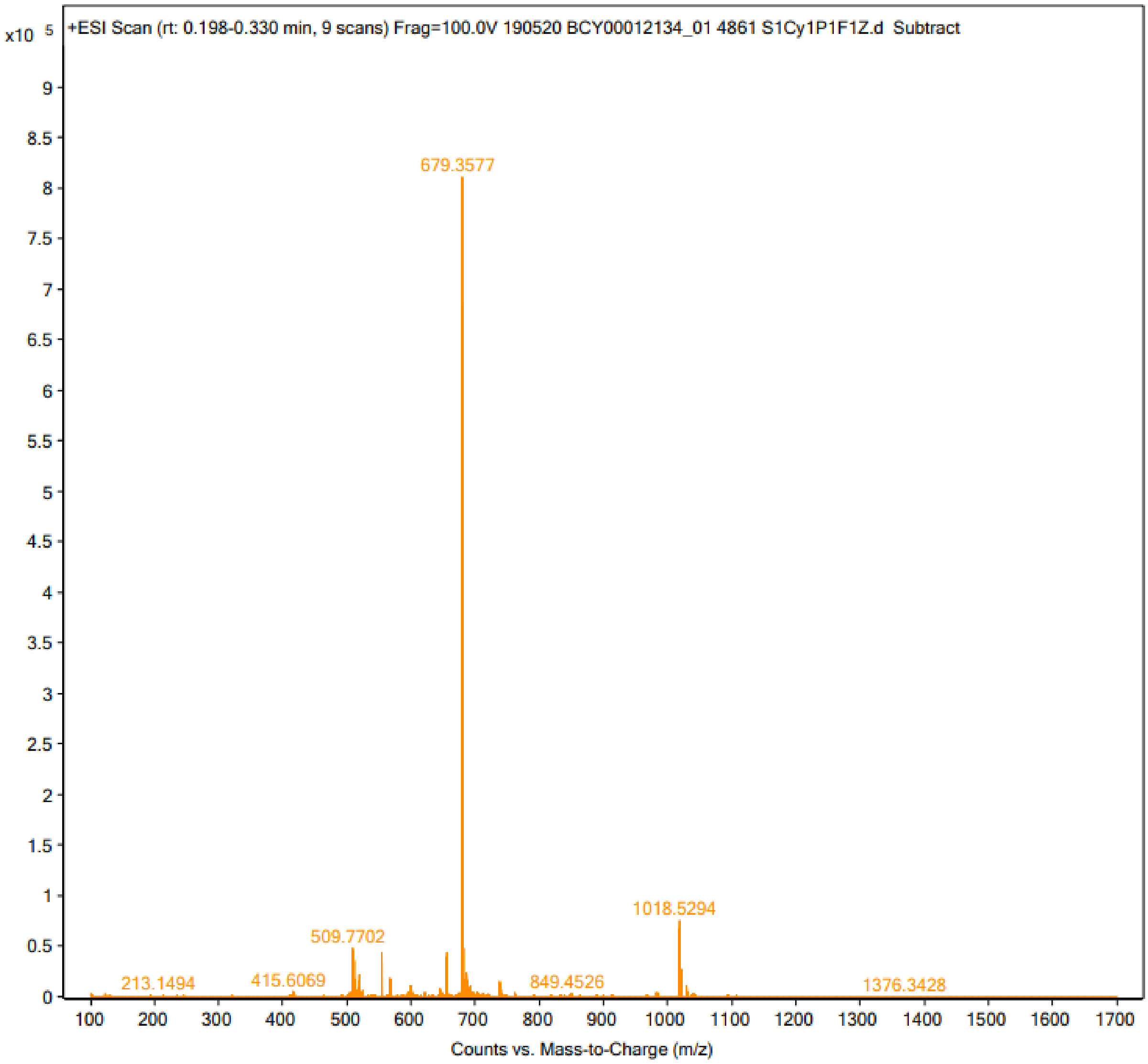
x10 5
+ESI Scan (rt: 0.198-0.330 min, 9 scans) Frag=100.0V 190520 BCY00012134_01 4861 S1Cy1P1F1Z.d Subtract
679.3577
1018.5294
509.7702
213.1494
415.6069
849.4526
1376.3428
0
0.5
1
1.5
2
2.5
3
3.5
4
4.5
5
5.5
6
6.5
7
7.5
8
8.5
9
100
200
300
400
500
600
700
800
900
1000
1100
1200
1300
1400
1500
1600
1700
Counts vs. Mass-to-Charge (m/z)

232

#### Peptide 13

**Data file:** Y:\Cunegonde\Chem32\1\Data\Peptides\4800-4899_BCY00012123_01 4841 S1Cy1P1F1z 2019-05-02 17-25-45.D

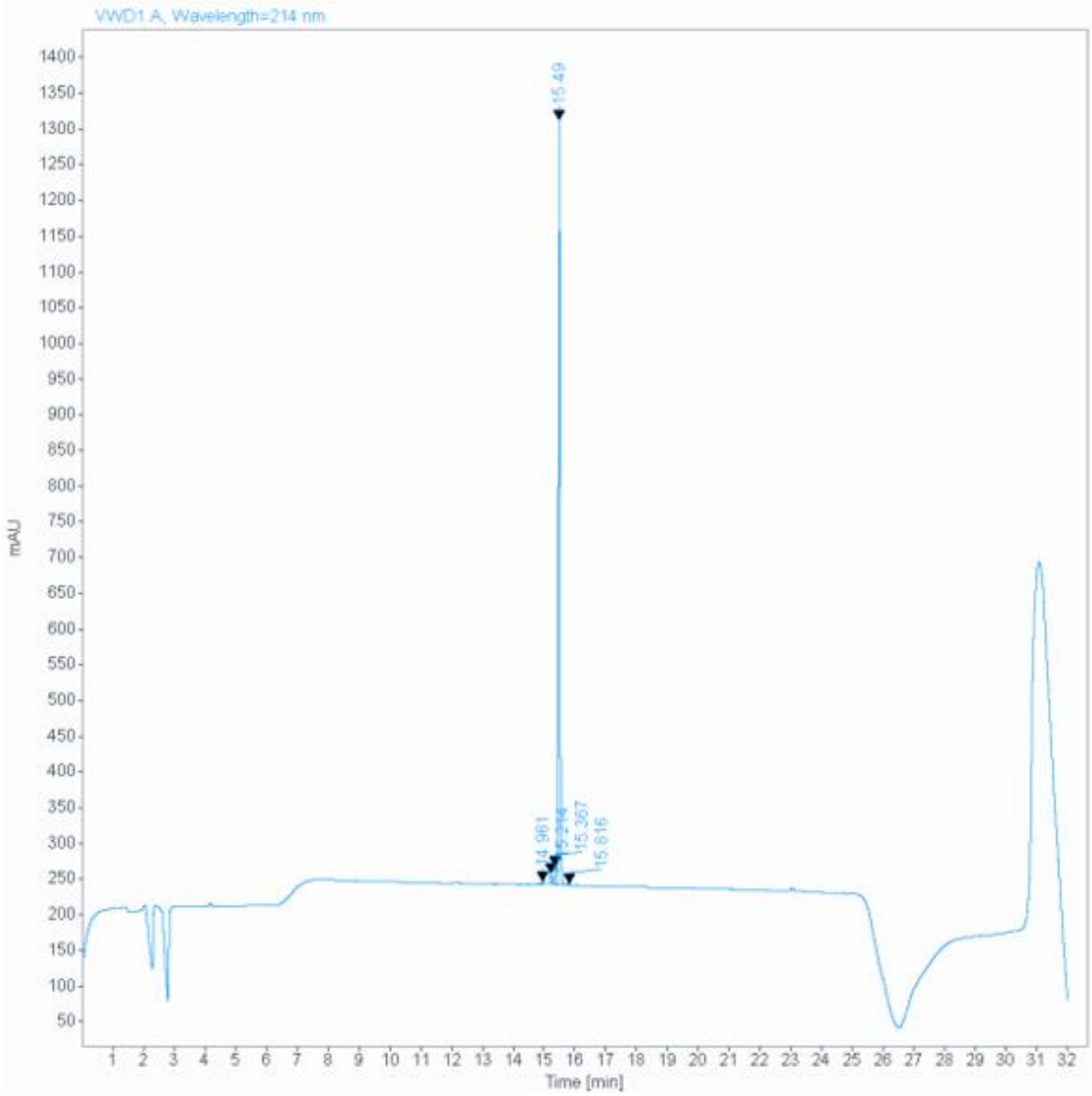

**Signal:** VWD1 A, Wavelength=214 nm

| RT [min] | Type | Width [min] | Area | Height | Area% | Name |
|---|---|---|---|---|---|---|
| 14.961 | VB | 0.0630 | 13.6901 | 3.1022 | 0.3796 | |
| 15.214 | BV E | 0.0557 | 55.0858 | 15.0598 | 1.5273 | |
| 15.367 | VV E | 0.0784 | 126.9883 | 25.1406 | 3.5208 | |
| 15.490 | VB R | 0.0486 | 3402.8162 | 1070.3527 | 94.3445 | |
| 15.816 | VB | 0.0464 | 8.2186 | 2.7497 | 0.2279 | |
| | | Sum | 3606.7990 | | | |

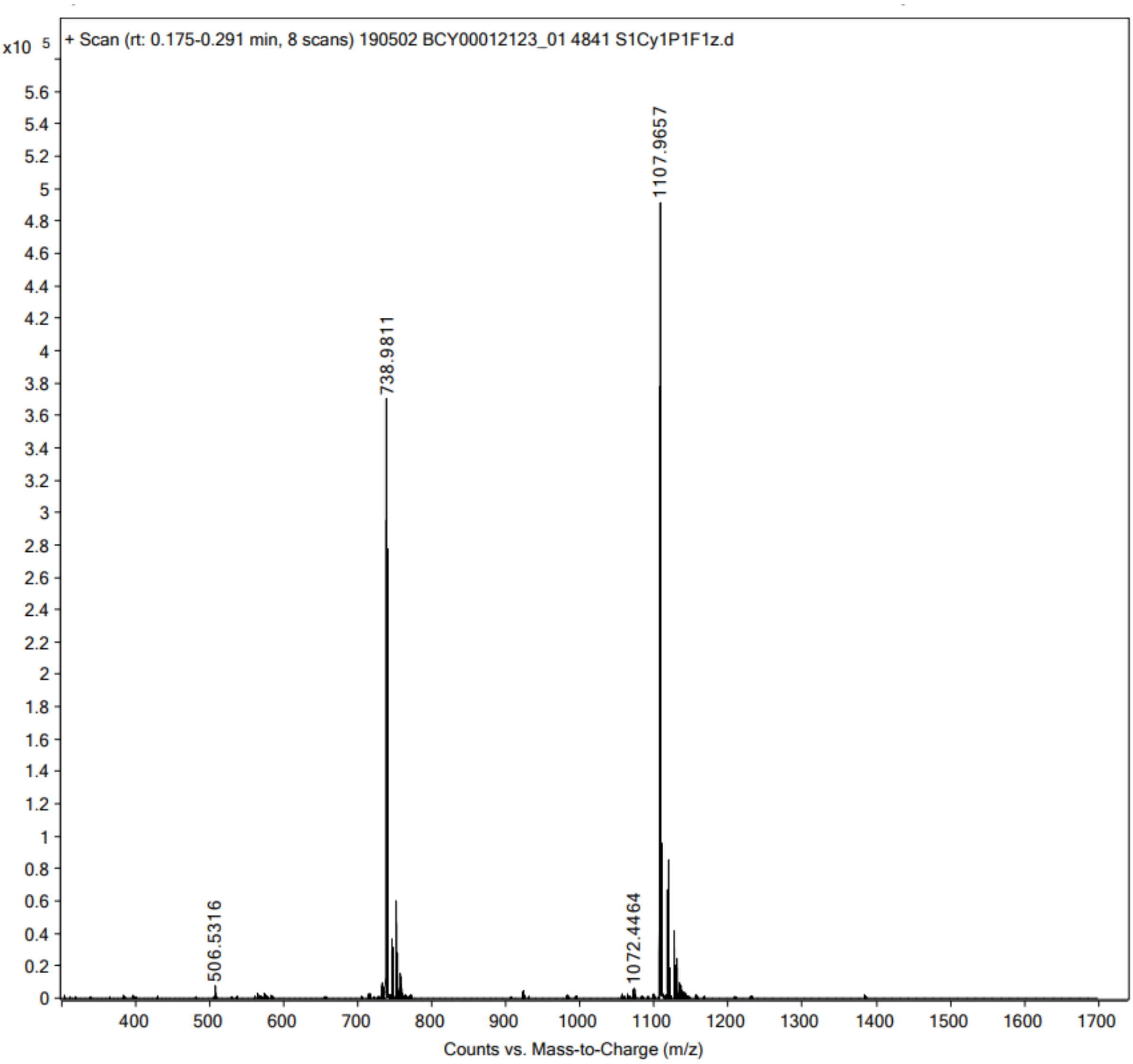

x10 5
+ Scan (rt: 0.175-0.291 min, 8 scans) 190502 BCY00012123_01 4841 S1Cy1P1F1z.d
1107.9657
738.9811
506.5316
1072.4464
Counts vs. Mass-to-Charge (m/z)

235

#### Peptide 14

**Data file:** D:\Chemstation\1\Data\Peptides\4800-4899_BCY00012126_01 4844 S1Cy1P1F1z 2019-05-16 15-45-06.D

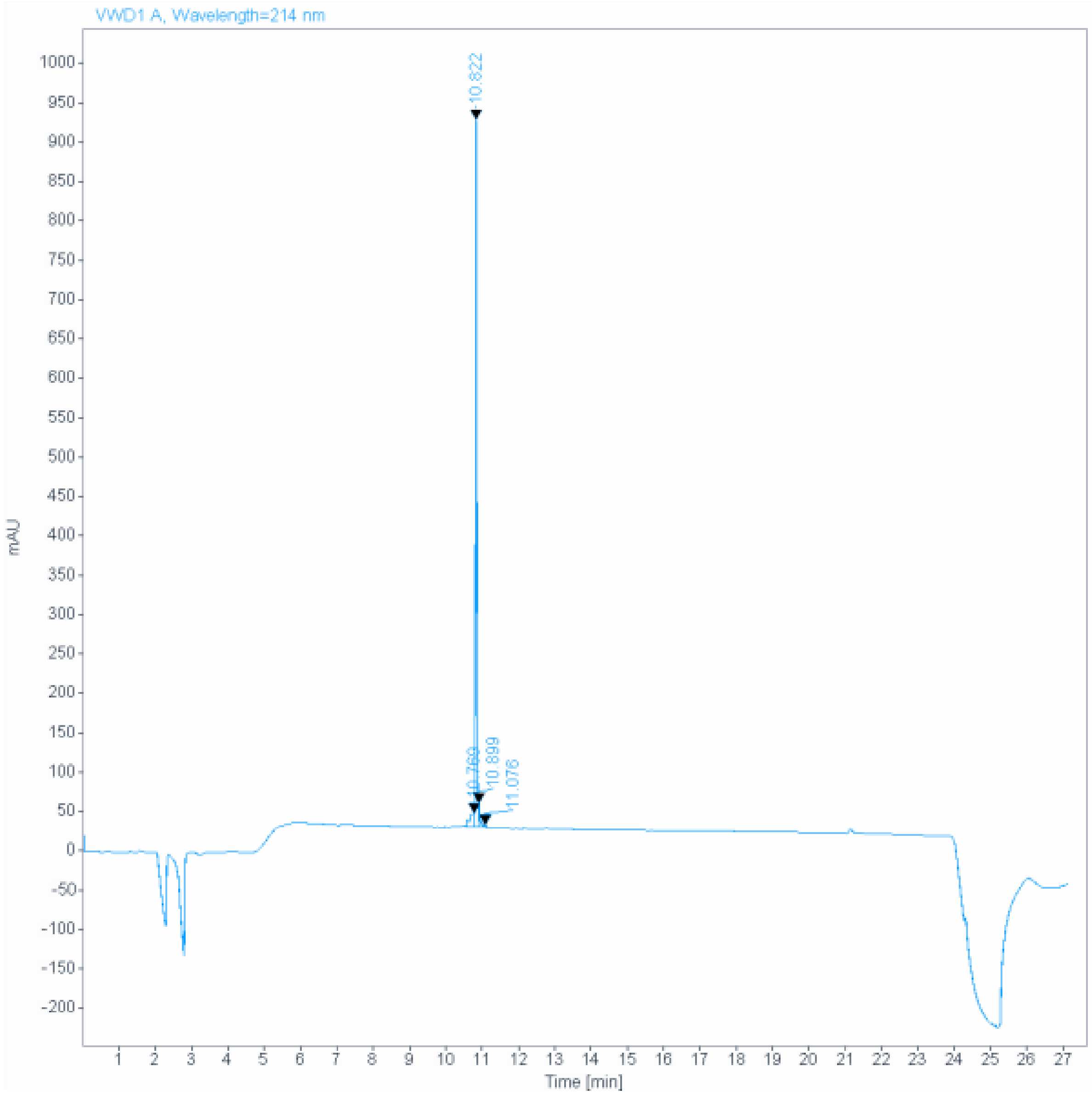

**Signal:** VWD1 A, Wavelength=214 nm

| RT [min] | Type | Width [min] | Area | Height | Area% | Name |
|---|---|---|---|---|---|---|
| 10.769 | MF | 0.1517 | 158.2035 | 17.3859 | 5.6179 | |
| 10.822 | FM | 0.0477 | 2572.5610 | 898.4066 | 91.3524 | |
| 10.899 | FM | 0.0397 | 71.8466 | 30.1981 | 2.5513 | |
| 11.076 | MM | 0.0654 | 13.4723 | 3.4346 | 0.4784 | |
| | | Sum | 2816.0835 | | | |

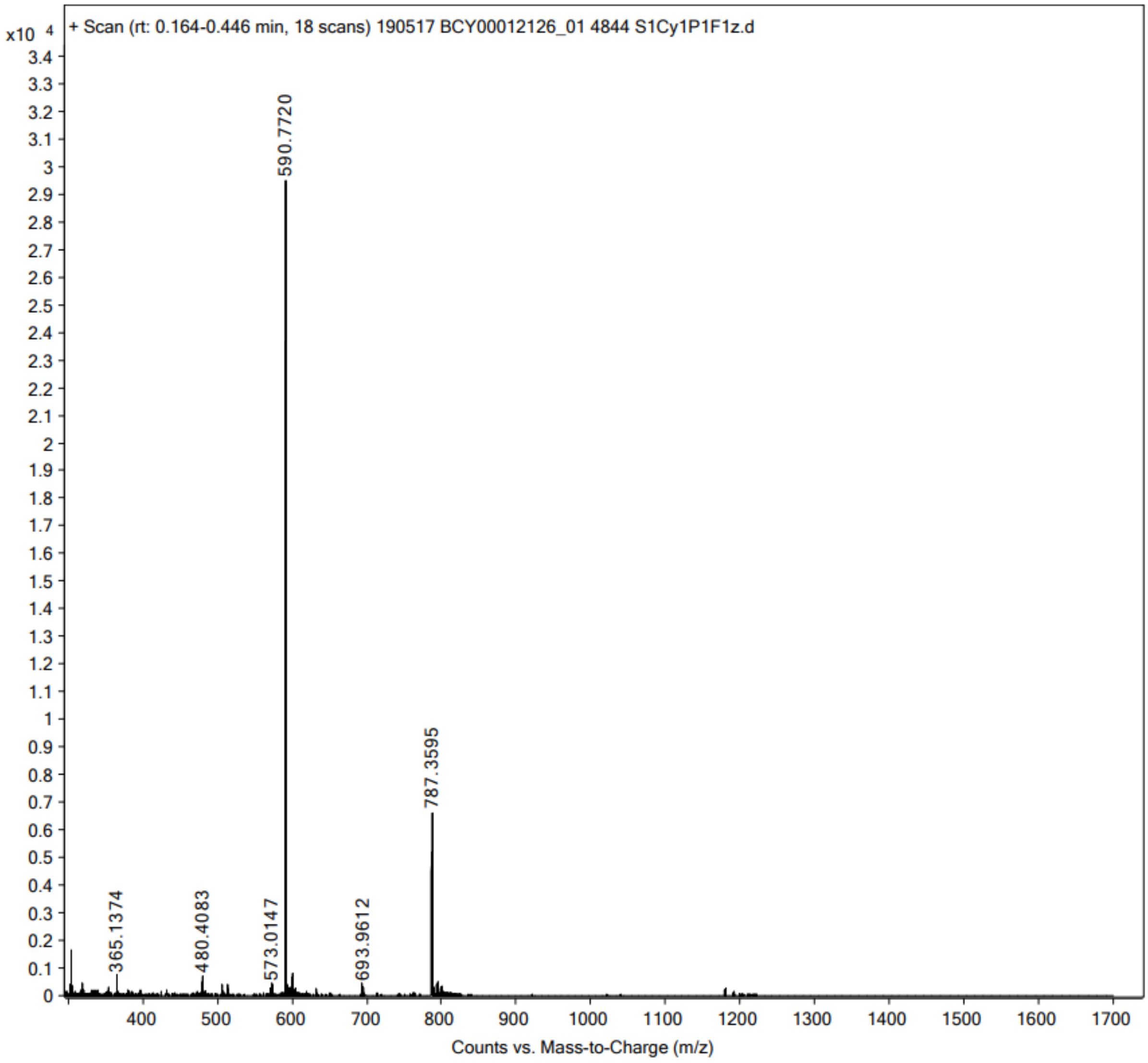

x10 4
+ Scan (rt: 0.164-0.446 min, 18 scans) 190517 BCY00012126_01 4844 S1Cy1P1F1z.d
590.7720
787.3595
365.1374
480.4083
573.0147
693.9612
Counts vs. Mass-to-Charge (m/z)

238

239 **Peptide 15**

**Data file:** C:\Chem32\1\Data\Peptides\4800-4899_BCY00012128_01 4862 S1Cy1P1F1F2Z 2019-05-20 18-18-03.D

**Sample name:** _BCY00012128_01 4862 S1Cy1P1F1F2Z

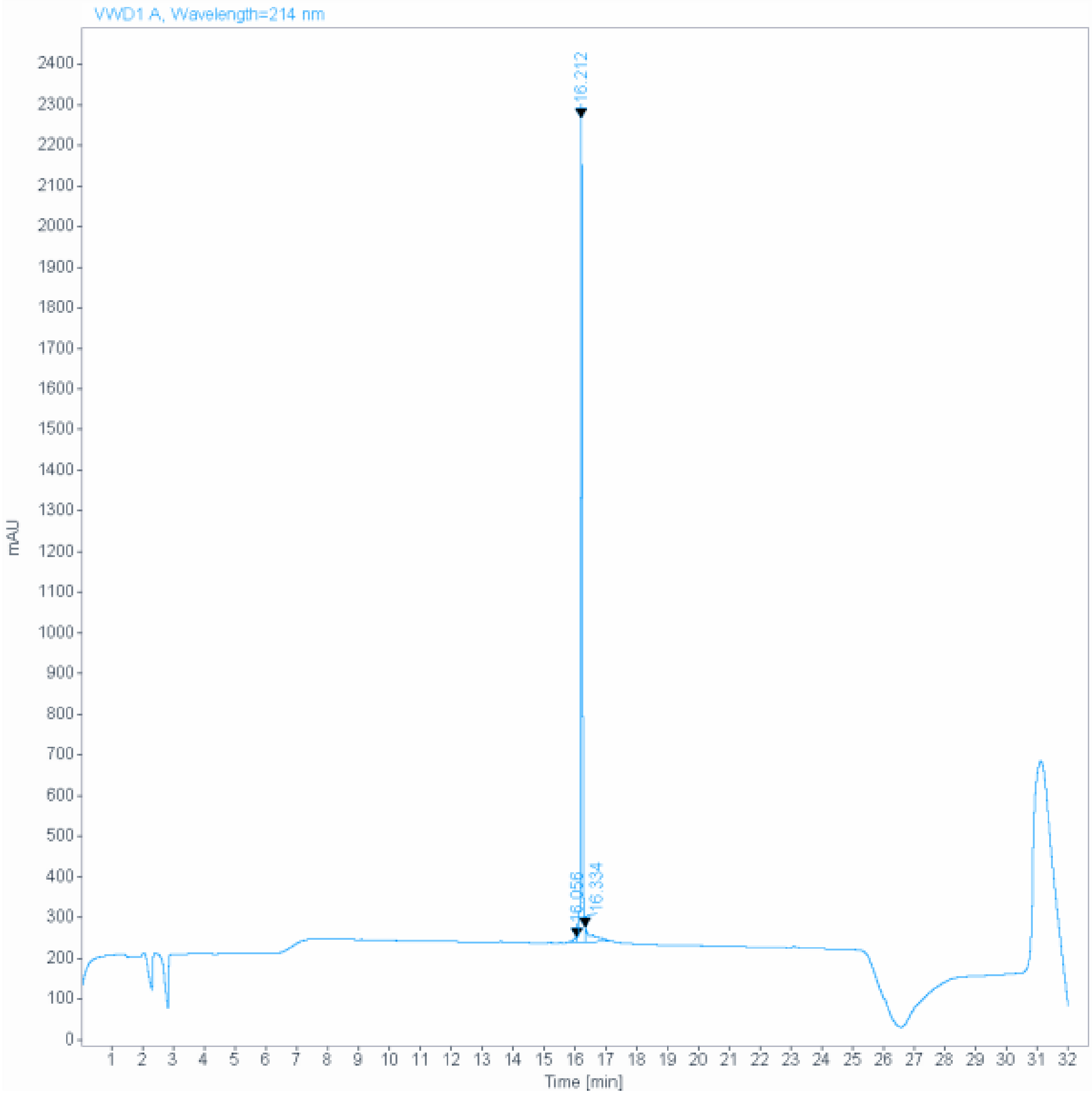

**Signal:** VWD1 A, Wavelength=214 nm

| RT [min] | Type | Width [min] | Area | Height | Area% | Name |
|---|---|---|---|---|---|---|
| 16.056 | MF | 0.1189 | 76.1543 | 10.6751 | 0.9295 | |
| 16.212 | MF | 0.0626 | 7619.6328 | 2027.8024 | 92.9980 | |
| 16.334 | FM | 0.2442 | 497.5469 | 33.9556 | 6.0726 | |
| | | Sum | 8193.3340 | | | |

240

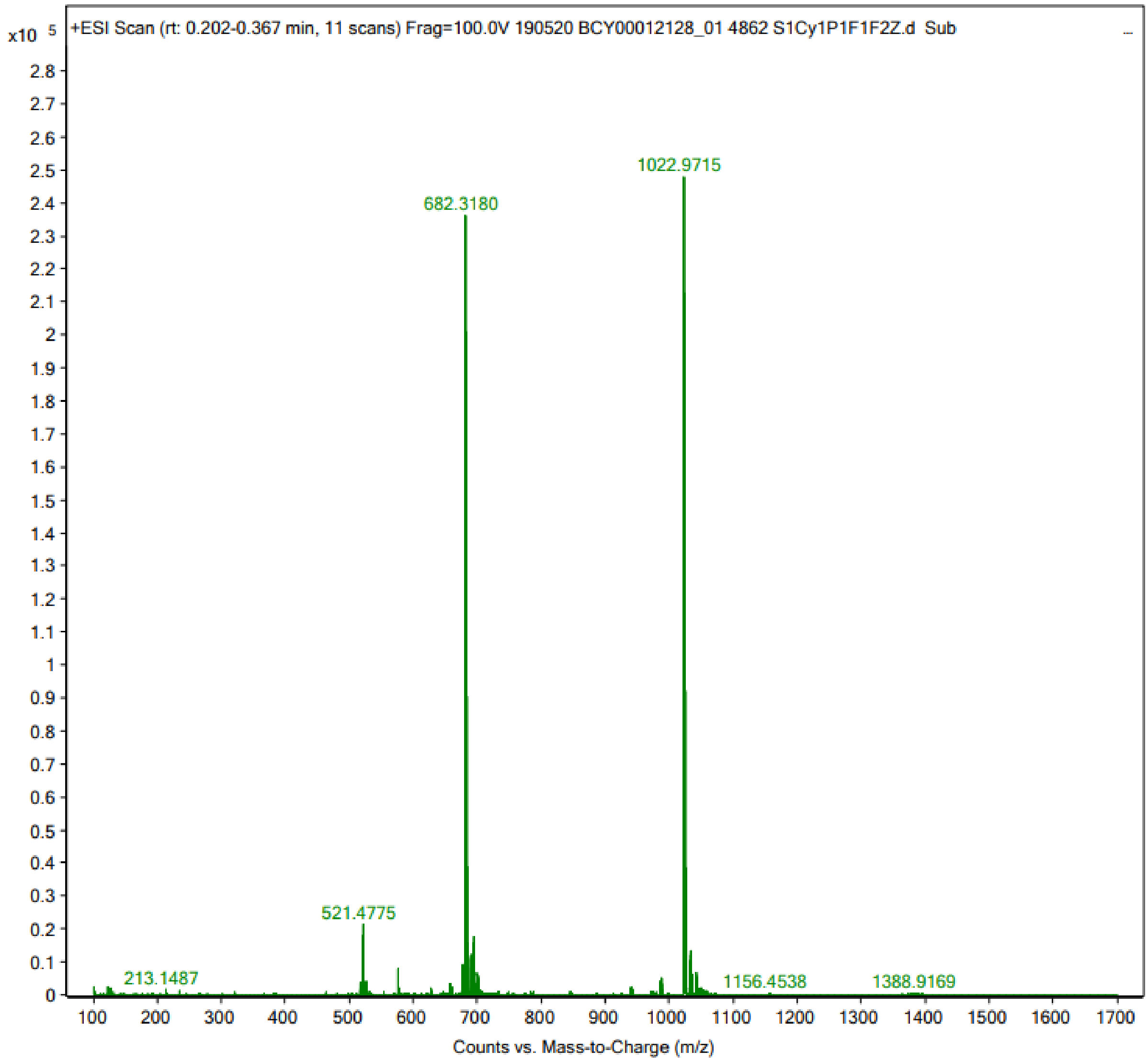
x10 5
+ESI Scan (rt: 0.202-0.367 min, 11 scans) Frag=100.0V 190520 BCY00012128_01 4862 S1Cy1P1F1F2Z.d Sub
1022.9715
682.3180
521.4775
213.1487
1156.4538
1388.9169
Counts vs. Mass-to-Charge (m/z)

### Peptide 16

**Sample name:** (101-08-00) Ala1 (TATA) QC
**Instrument:** 1260_1

**Injection date:** 11/4/2019 2:52:52 PM
**Acq. method:** 0595B_AB_Poroshell1
20_15.5min.M

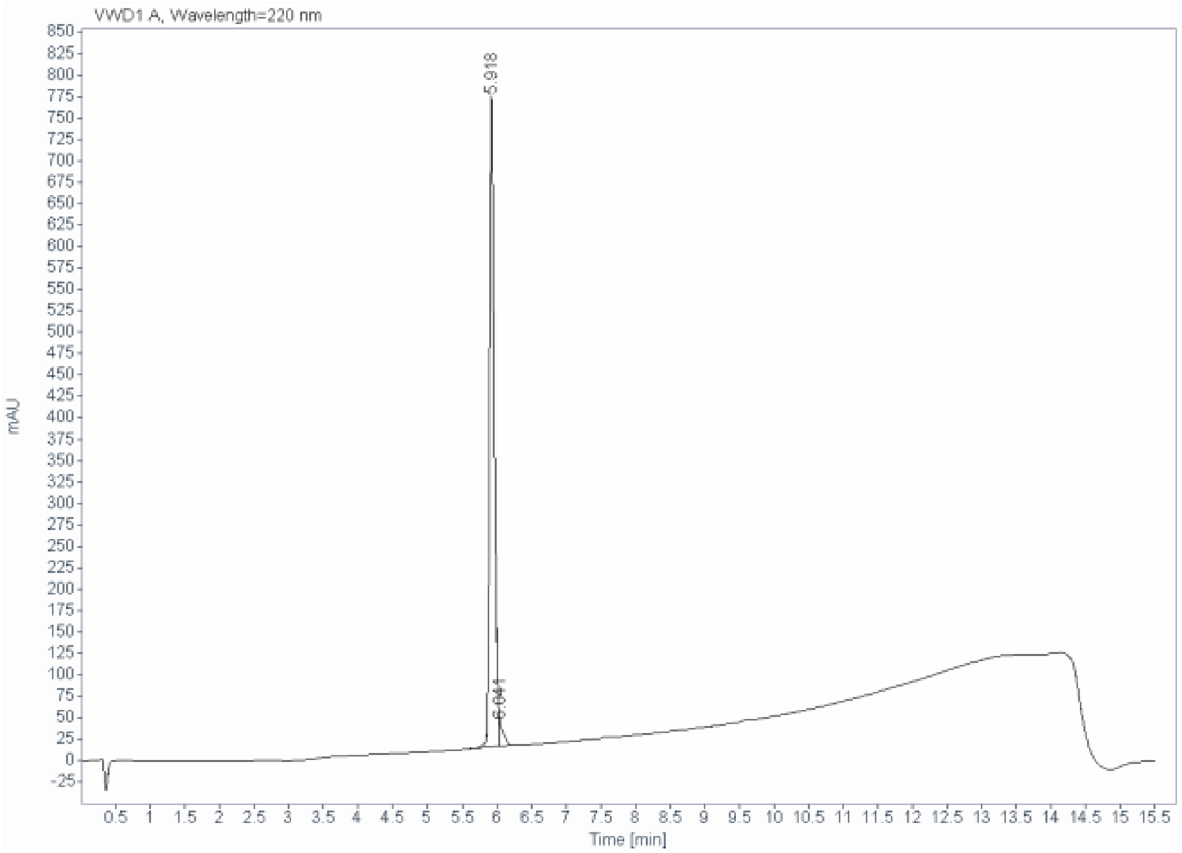

**Signal:** VWD1 A, Wavelength=220 nm

| RT [min] | Type | Width [min] | Area | Height | Area% |
|---|---|---|---|---|---|
| 5.918 | BF | 0.0724 | 3666.1338 | 757.6926 | 97.2293 |
| 6.041 | VB | 0.0617 | 104.4730 | 28.2120 | 2.7707 |
| | | Sum | 3770.6067 | | |

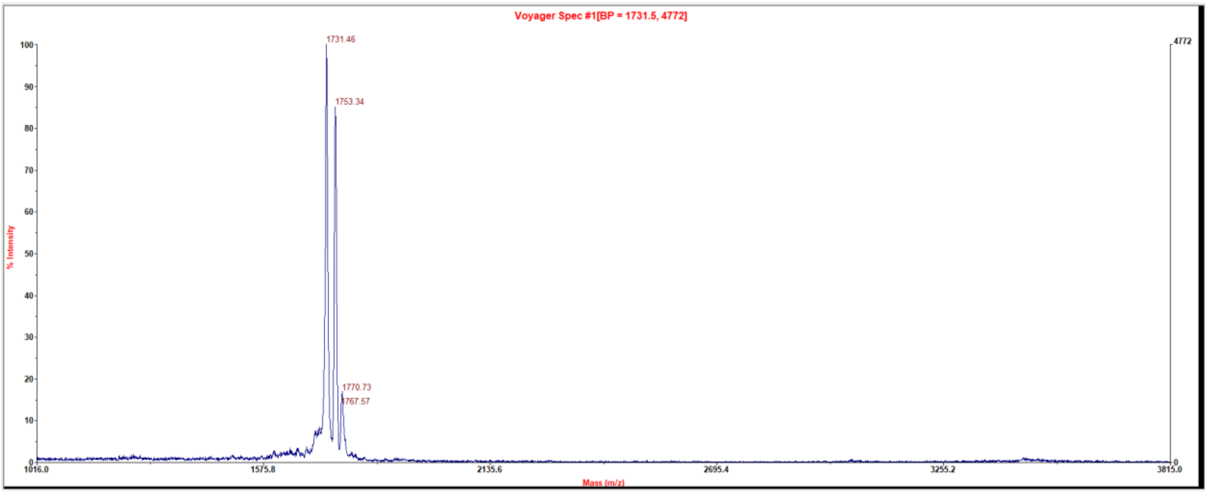

### Peptide 17

| | |
|---|---|
| **Sample name:** | (101-08-00) Ala2 (TATA) QC |
| **Instrument:** | 1260_1 |
| **Injection date:** | 11/4/2019 3:09:26 PM |
| **Acq. method:** | 0595B_AB_Poroshell120_15.5min.M |

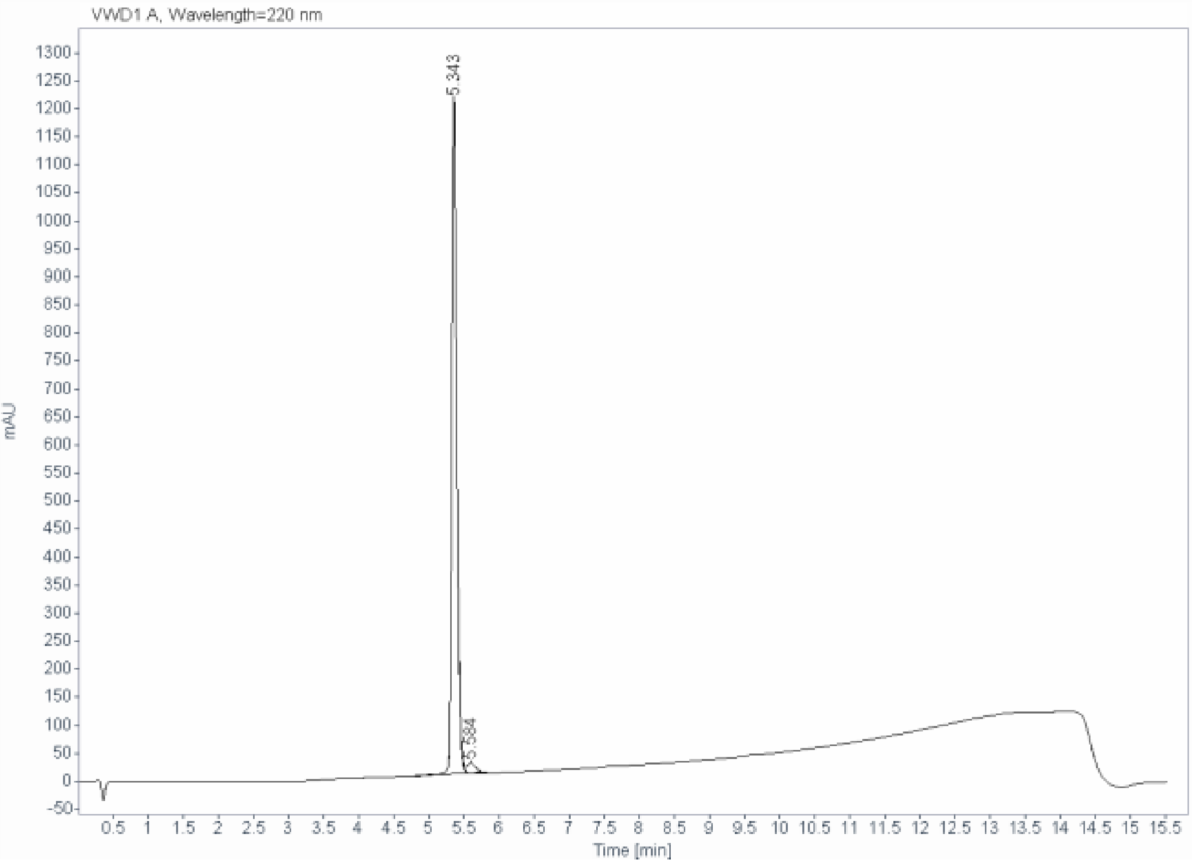

**Signal:** VWD1 A, Wavelength=220 nm

| RT [min] | Type | Width [min] | Area | Height | Area% |
|---|---|---|---|---|---|
| 5.343 | BV R | 0.0744 | 5935.8984 | 1206.6854 | 97.1012 |
| 5.584 | VB E | 0.1295 | 177.2058 | 18.7091 | 2.8988 |
| | | Sum | 6113.1042 | | |

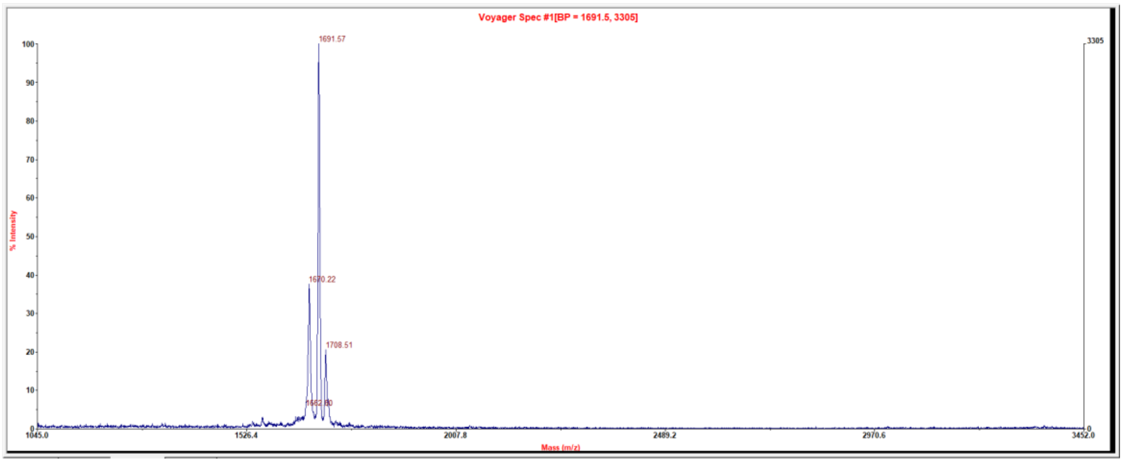

#### Peptide 18

| | |
|---|---|
| **Sample name:** | (101-08-00) Ala3 (TATA) QC |
| **Instrument:** | 1260_1 |
| **Injection date:** | 11/4/2019 3:25:45 PM |
| **Acq. method:** | 0595B_AB_Poroshell120_15.5min.M |

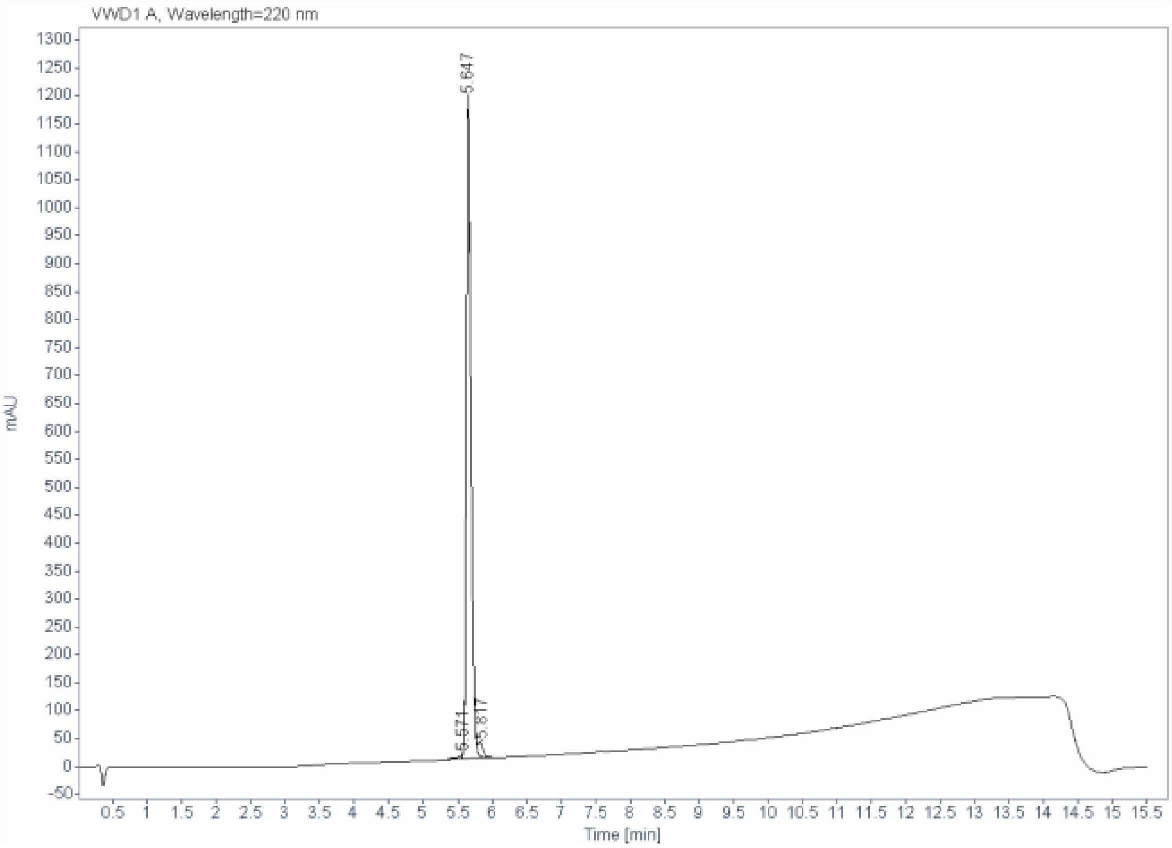

**Signal:** VWD1 A, Wavelength=220 nm

| RT [min] | Type | Width [min] | Area | Height | Area% |
|---|---|---|---|---|---|
| 5.571 | BV | 0.1024 | 52.6781 | 8.5738 | 0.9037 |
| 5.647 | VV R | 0.0716 | 5646.2817 | 1185.5745 | 96.8662 |
| 5.817 | VB E | 0.0786 | 129.9880 | 24.6271 | 2.2300 |
| | | Sum | 5828.9479 | | |

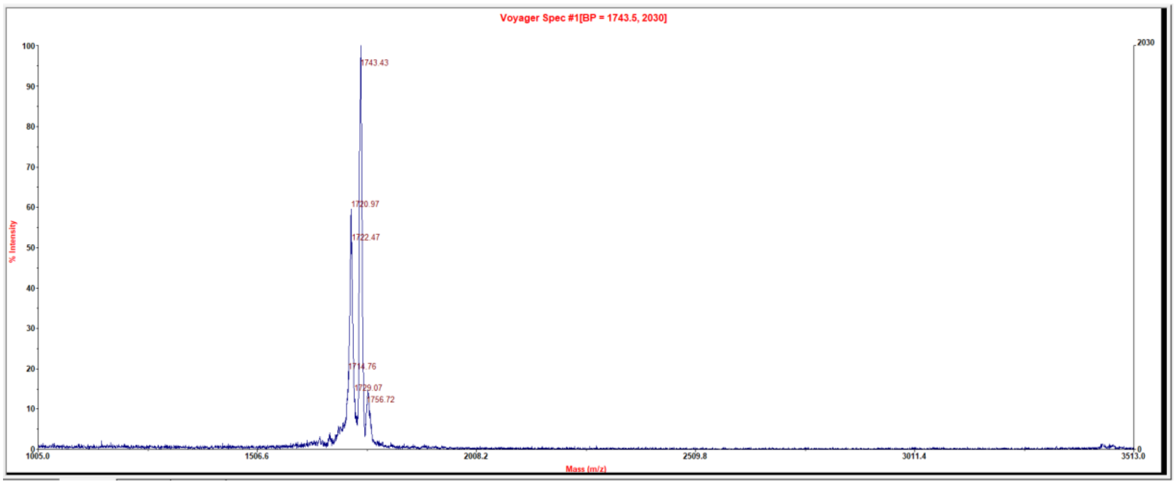

### Peptide 19

| | |
|---|---|
| **Sample name:** | (101-08-00) Ala4 (TATA) QC |
| **Instrument:** | 1260_1 |
| **Injection date:** | 11/4/2019 3:42:20 PM |
| **Acq. method:** | 0595B_AB_Poroshell120_15.5min.M |

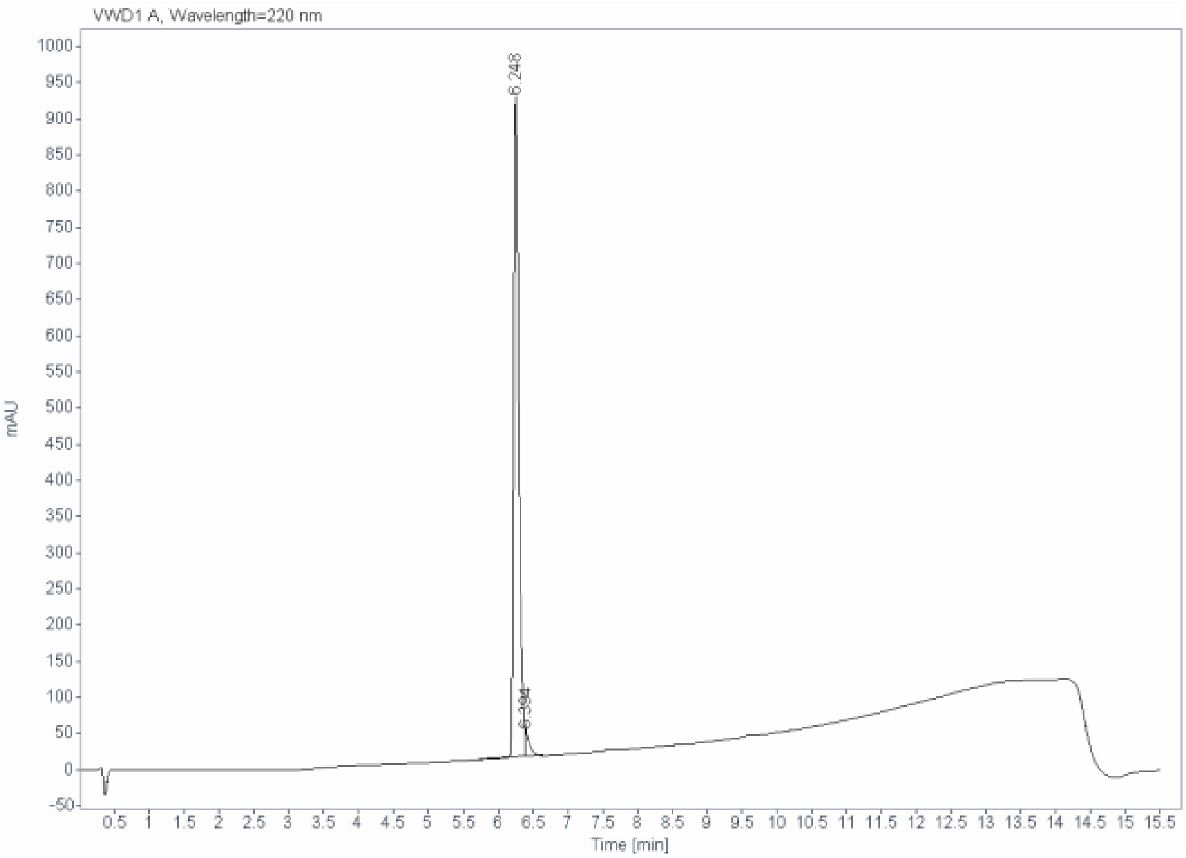

**Signal:** VWD1 A, Wavelength=220 nm

| RT [min] | Type | Width [min] | Area | Height | Area% |
|---|---|---|---|---|---|
| 6.248 | BF | 0.0717 | 4504.1519 | 910.2987 | 97.1590 |
| 6.394 | VB | 0.0654 | 131.7032 | 33.5718 | 2.8410 |
| | | Sum | 4635.8550 | | |

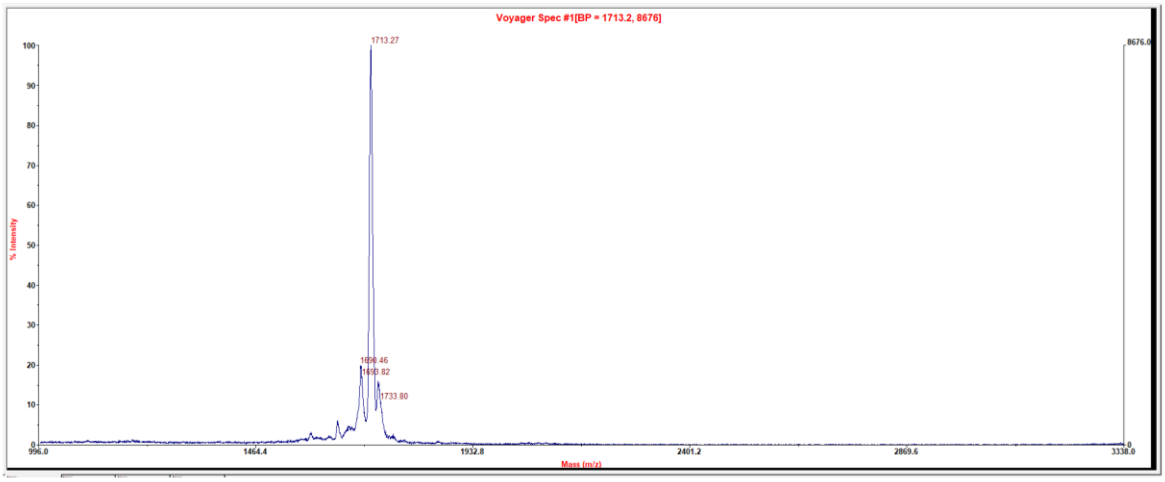

### Peptide 20

| | |
|---|---|
| **Sample name:** | (101-08-00) Ala5 (TATA) QC2 |
| **Instrument:** | 1260_1 |
| **Injection date:** | 11/4/2019 4:14:59 PM |
| **Acq. method:** | 0595B_AB_Poroshell120_15.5min.M |

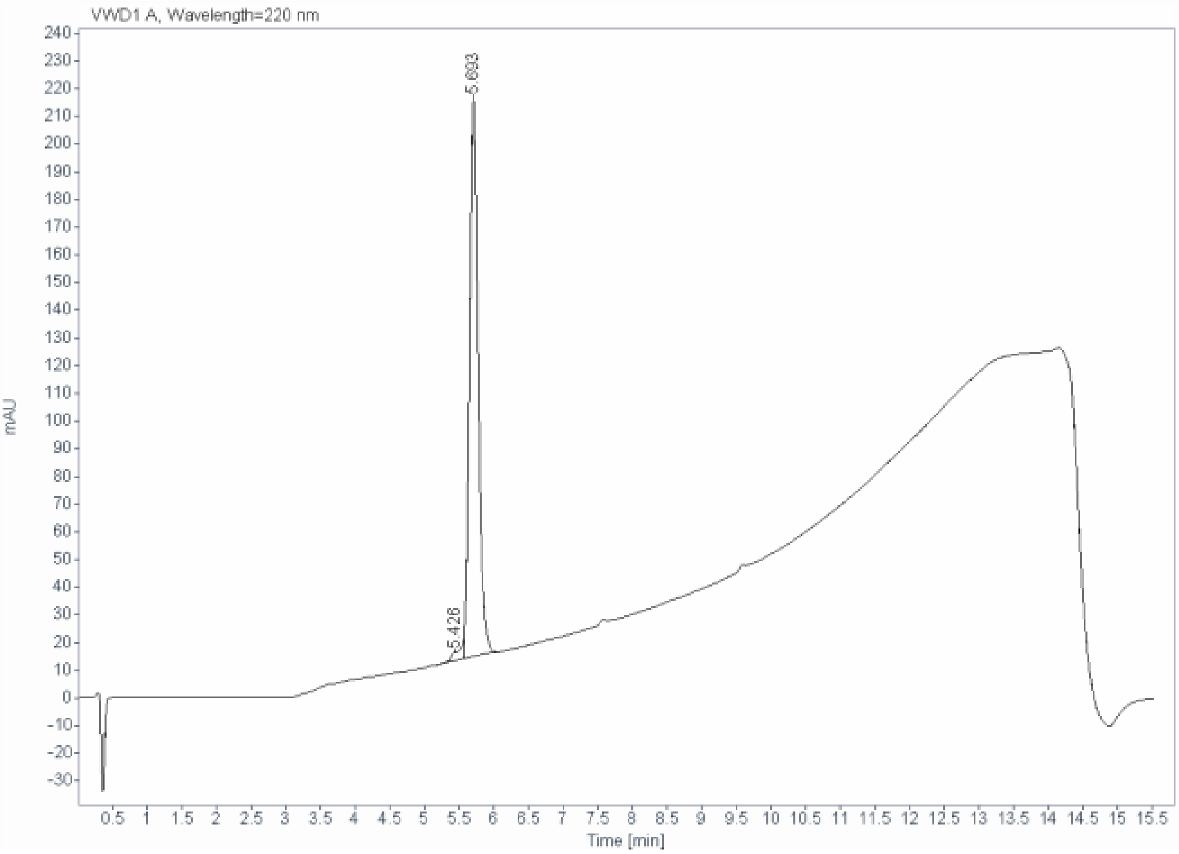

**Signal:** VWD1 A, Wavelength=220 nm

| RT [min] | Type | Width [min] | Area | Height | Area% |
|---|---|---|---|---|---|
| 5.426 | BV | 0.1952 | 50.1846 | 3.2858 | 2.7684 |
| 5.693 | VB | 0.1358 | 1762.5721 | 201.9507 | 97.2316 |
| | | Sum | 1812.7568 | | |

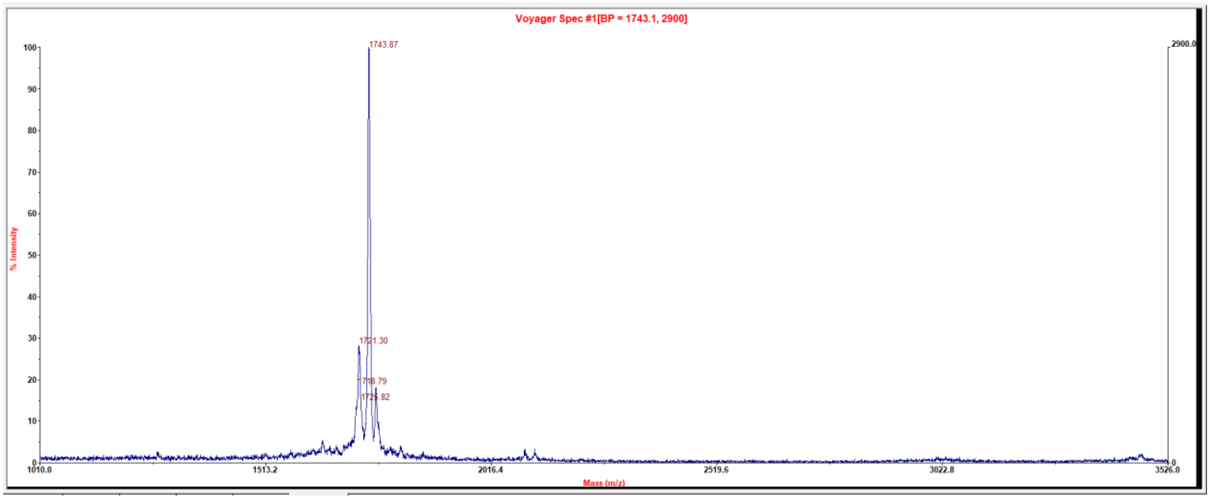

#### Peptide 21

| | |
|---|---|
| Sample name: | (101-08-00) Ala6 (TATA) QC2 |
| Instrument: | 1260_1 |
| Injection date: | 11/4/2019 4:55:17 PM |
| Acq. method: | 0595B_AB_Poroshell120_15.5min.M |

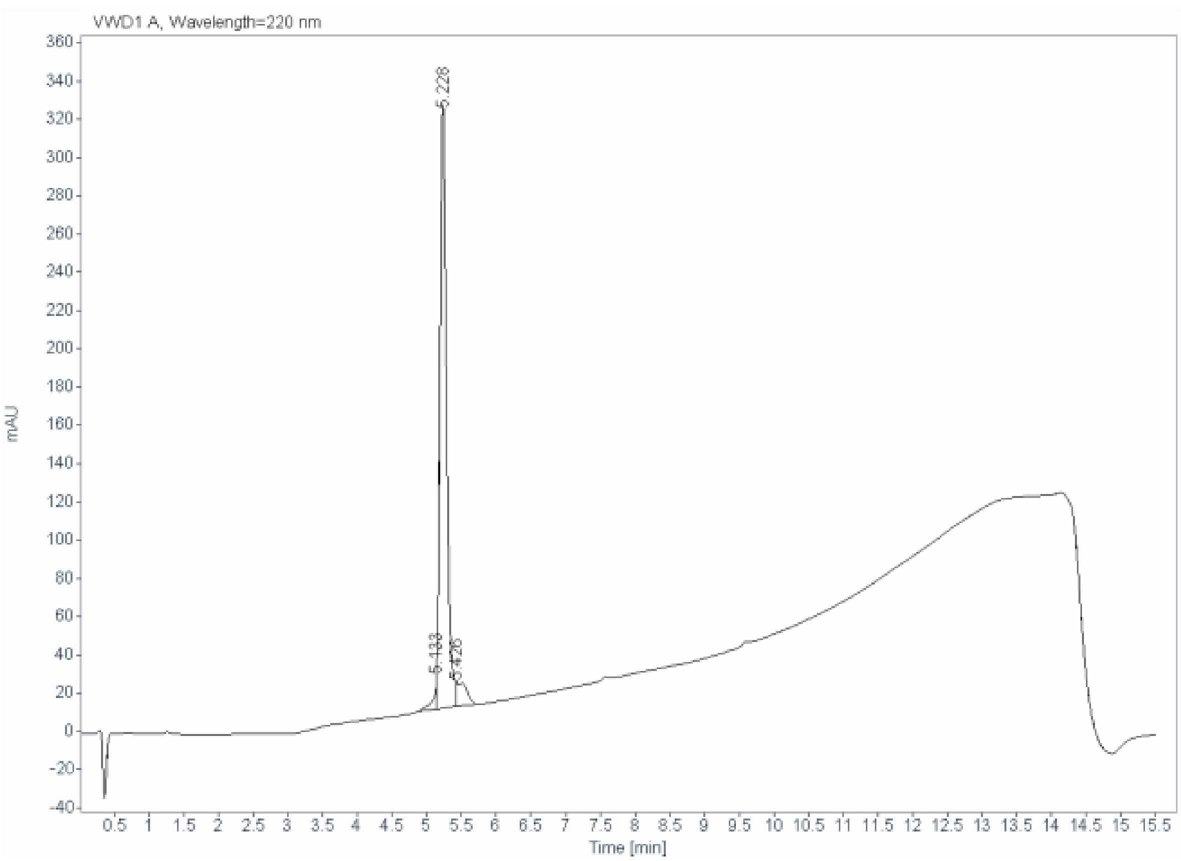

Signal: VWD1 A, Wavelength=220 nm

| RT [min] | Type | Width [min] | Area | Height | Area% |
|---|---|---|---|---|---|
| 5.133 | BV | 0.0480 | 52.2967 | 18.1773 | 2.3051 |
| 5.228 | VF | 0.1159 | 2094.3215 | 291.1203 | 92.3112 |
| 5.426 | VB | 0.1706 | 122.1438 | 11.9302 | 5.3837 |
| | | Sum | 2268.7620 | | |

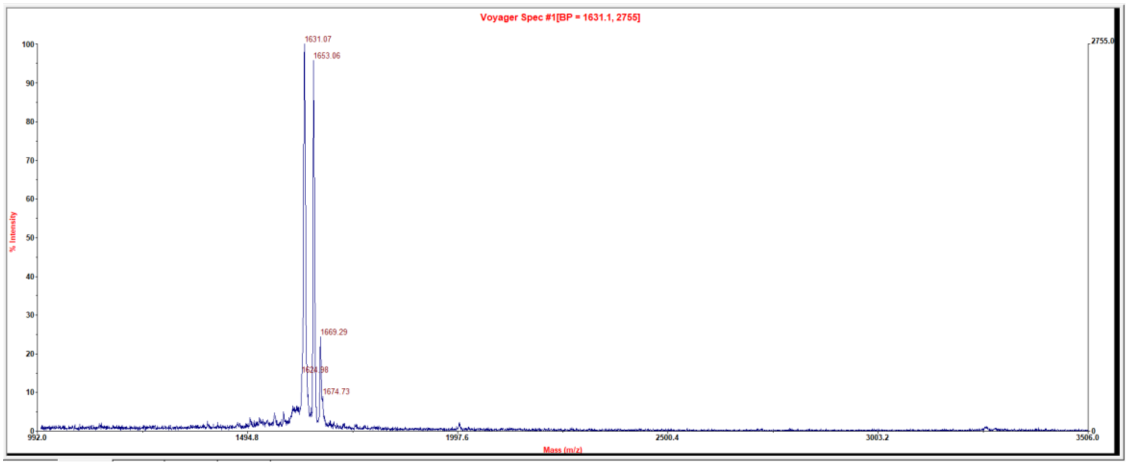

#### Peptide 22

| | |
|---|---|
| Sample name: | (101-08-00) Ala7 (TATA) QC2 |
| Instrument: | 1260_1 |
| Injection date: | 11/4/2019 5:11:36 PM |
| Acq. method: | 0595B_AB_Poroshell1 20_15.5min.M |

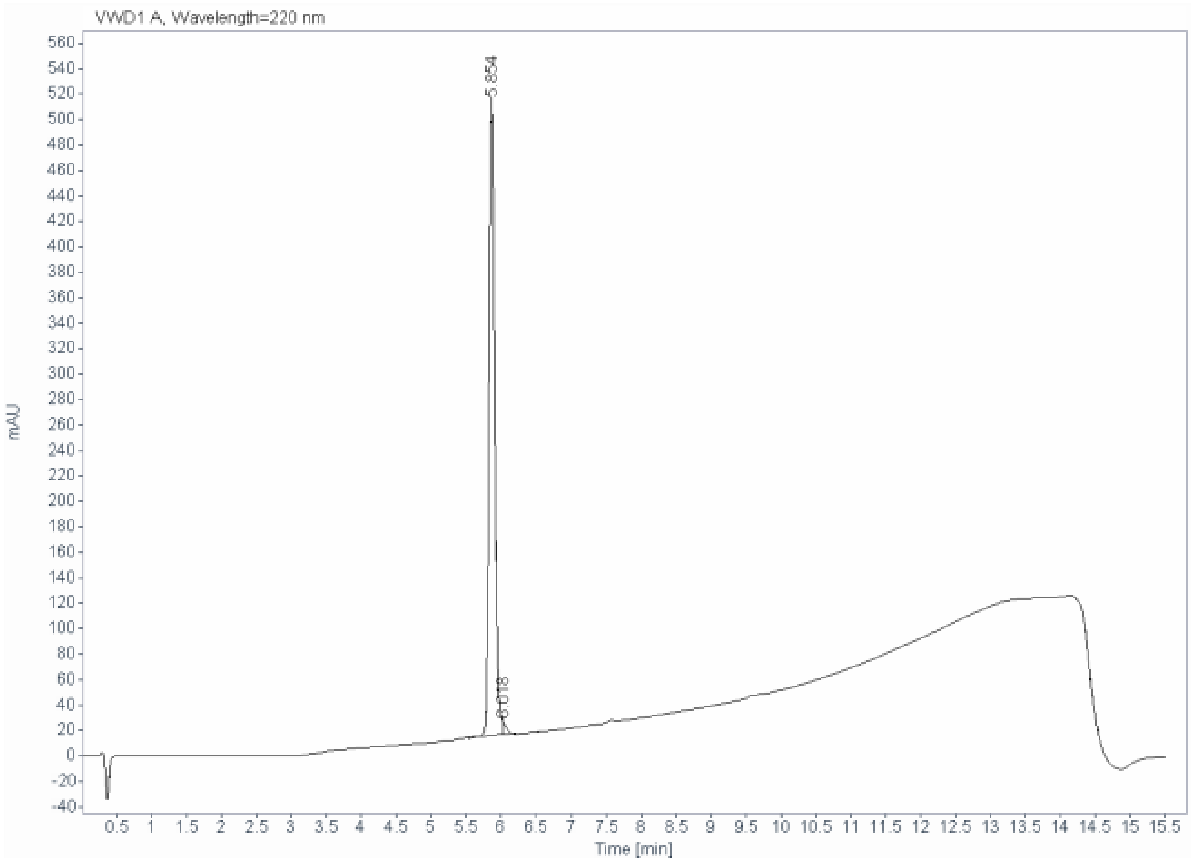

Signal: VWD1 A, Wavelength=220 nm

| RT [min] | Type | Width [min] | Area | Height | Area% |
|---|---|---|---|---|---|
| 5.854 | BF | 0.0871 | 2826.5938 | 498.8858 | 98.7241 |
| 6.018 | VB | 0.0584 | 36.5300 | 10.4305 | 1.2759 |
| | | Sum | 2863.1238 | | |

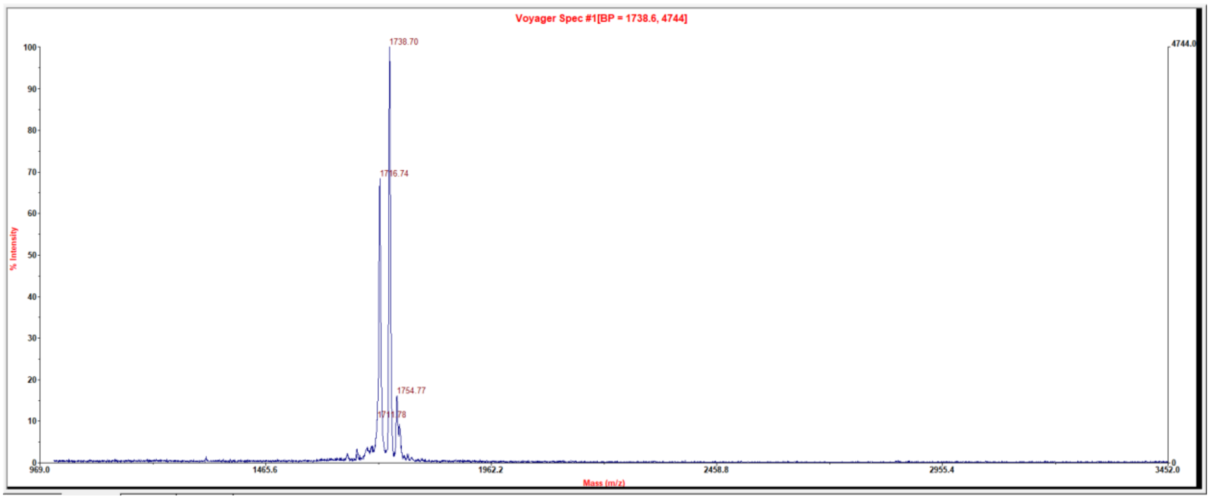

#### Peptide 23

| | |
|---|---|
| Sample name: | (101-08-00) Ala8 (TATA) QC |
| Instrument: | 1260_2 |
| Injection date: | 11/4/2019 5:15:56 PM |
| Acq. method: | 0595B_AB_Poroshell1 20_15.5min.M |

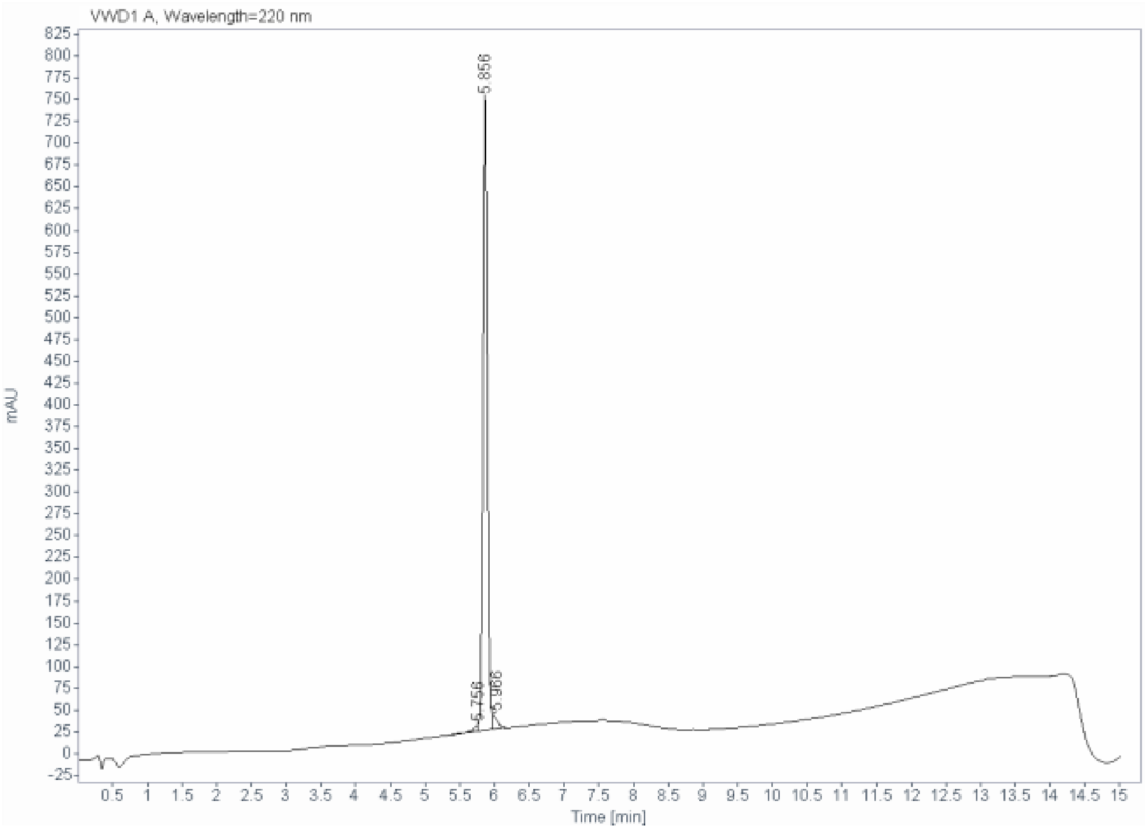

Signal: VWD1 A, Wavelength=220 nm

| RT [min] | Type | Width [min] | Area | Height | Area% |
|---|---|---|---|---|---|
| 5.756 | BV | 0.0858 | 39.7773 | 7.7236 | 1.2113 |
| 5.856 | VF | 0.0678 | 3165.6328 | 727.2114 | 96.3973 |
| 5.966 | VB | 0.0731 | 78.5320 | 17.9154 | 2.3914 |
| | | Sum | 3283.9422 | | |

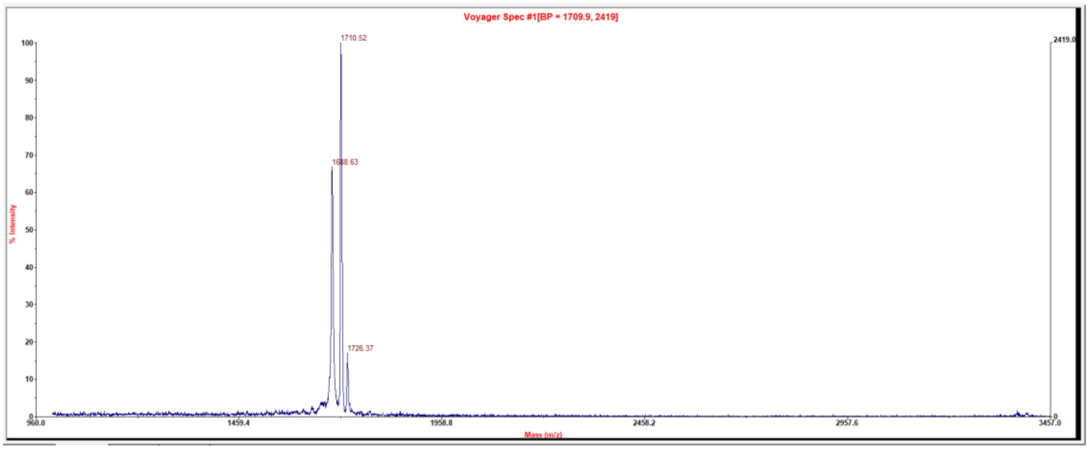

### Peptide 24

| | |
|---|---|
| Sample name: | (101-08-00) Ala9 (TATA) QC |
| Instrument: | 1260_1 |
| Injection date: | 11/4/2019 5:29:28 PM |
| Acq. method: | 0595B_AB_Poroshell120_15.5min.M |

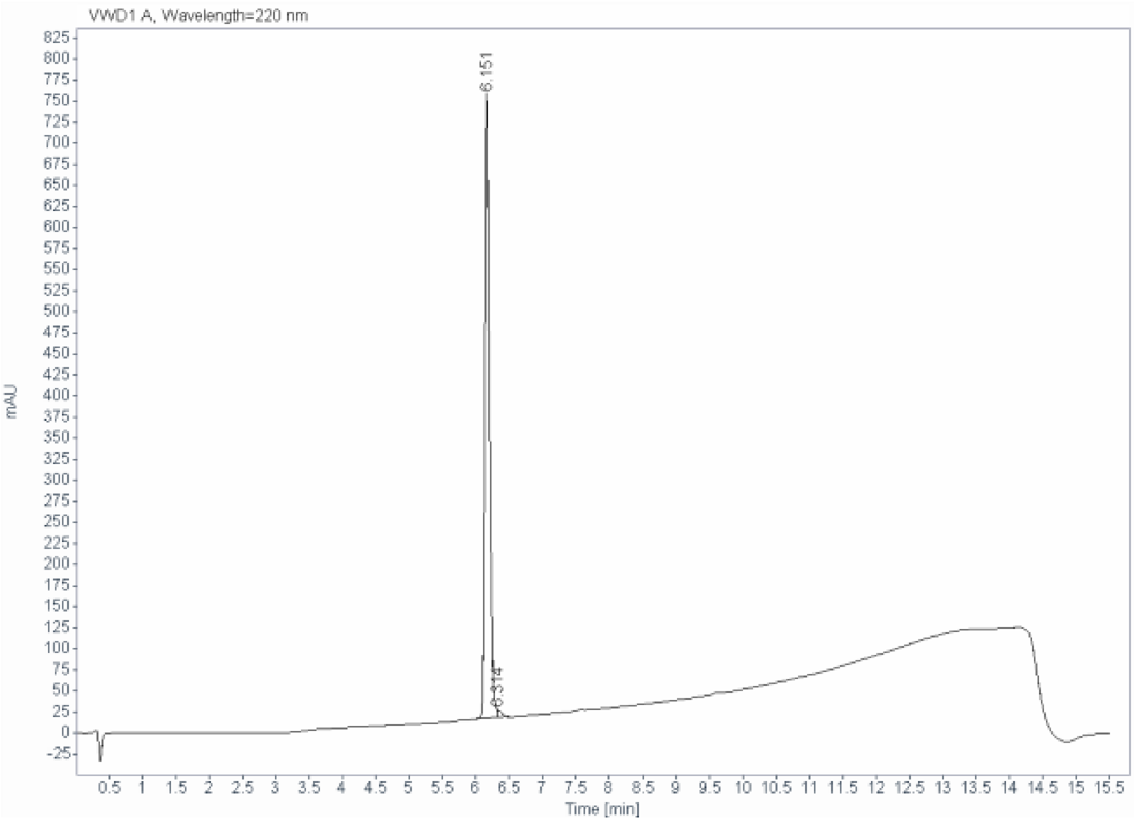

Signal: VWD1 A, Wavelength=220 nm

| RT [min] | Type | Width [min] | Area | Height | Area% |
|---|---|---|---|---|---|
| 6.151 | VF R | 0.0730 | 3612.3682 | 740.0170 | 98.9778 |
| 6.314 | VB | 0.0715 | 37.3083 | 8.6952 | 1.0222 |
| | | Sum | 3649.6765 | | |

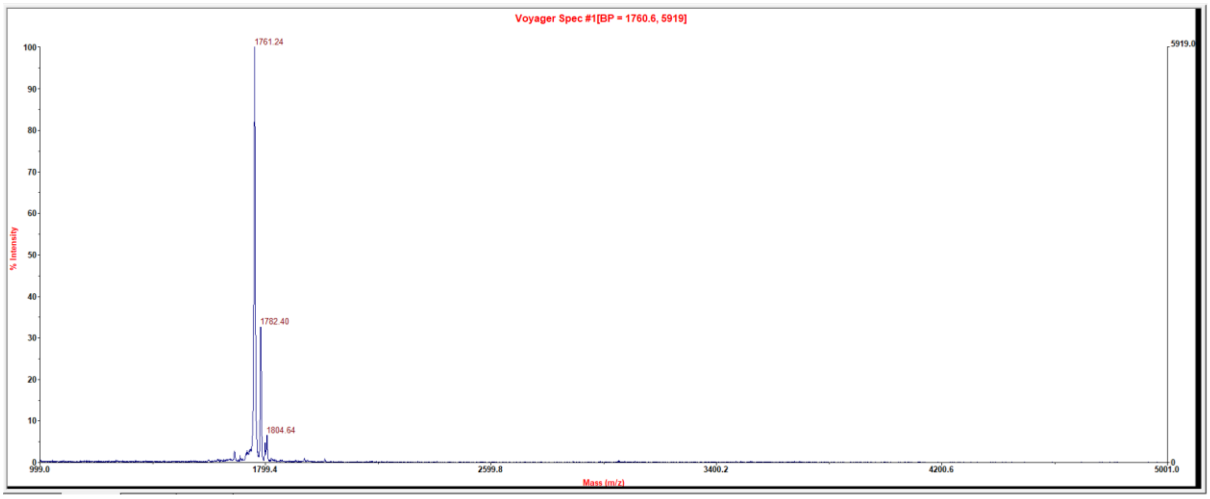

#### Peptide 25

| | |
|---|---|
| **Sample name:** | (101-08-00)HArg4 (TATA) QC |
| **Instrument:** | 1260_1 |
| **Injection date:** | 10/10/2019 12:34:59 PM |
| **Acq. method:** | 0595B_AB_Poroshell120_15.5min.M |

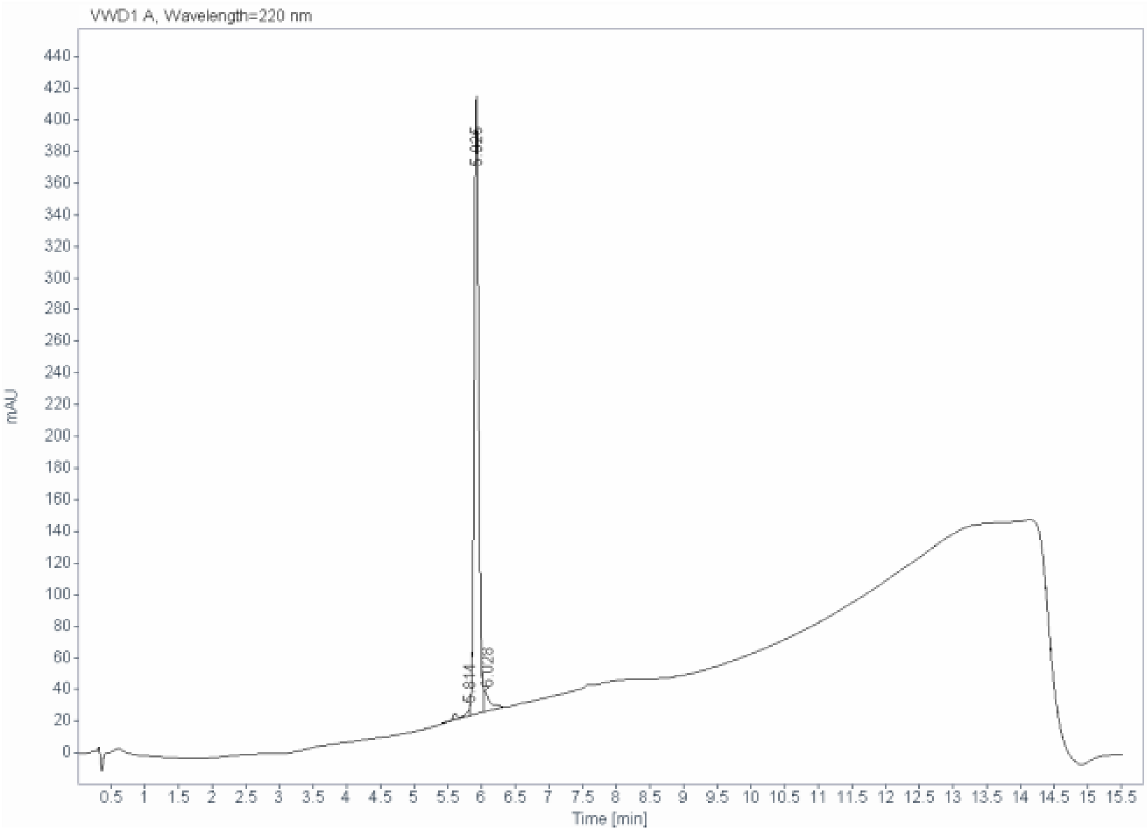

**Signal:** VWD1 A, Wavelength=220 nm

| RT [min] | Type | Width [min] | Area | Height | Area% |
|---|---|---|---|---|---|
| 5.814 | BV | 0.0328 | 44.0170 | 22.3527 | 2.2270 |
| 5.925 | VV | 0.1205 | 1848.8356 | 255.6557 | 93.5394 |
| 6.028 | VB | 0.0746 | 83.6783 | 18.6860 | 4.2336 |
| | | Sum | 1976.5309 | | |

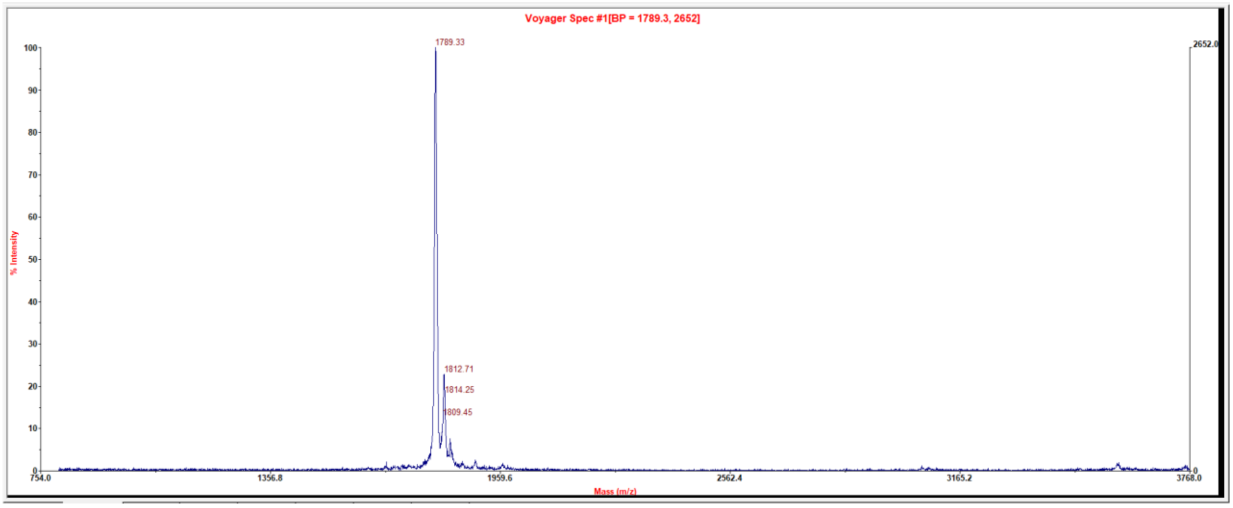

#### Peptide 26

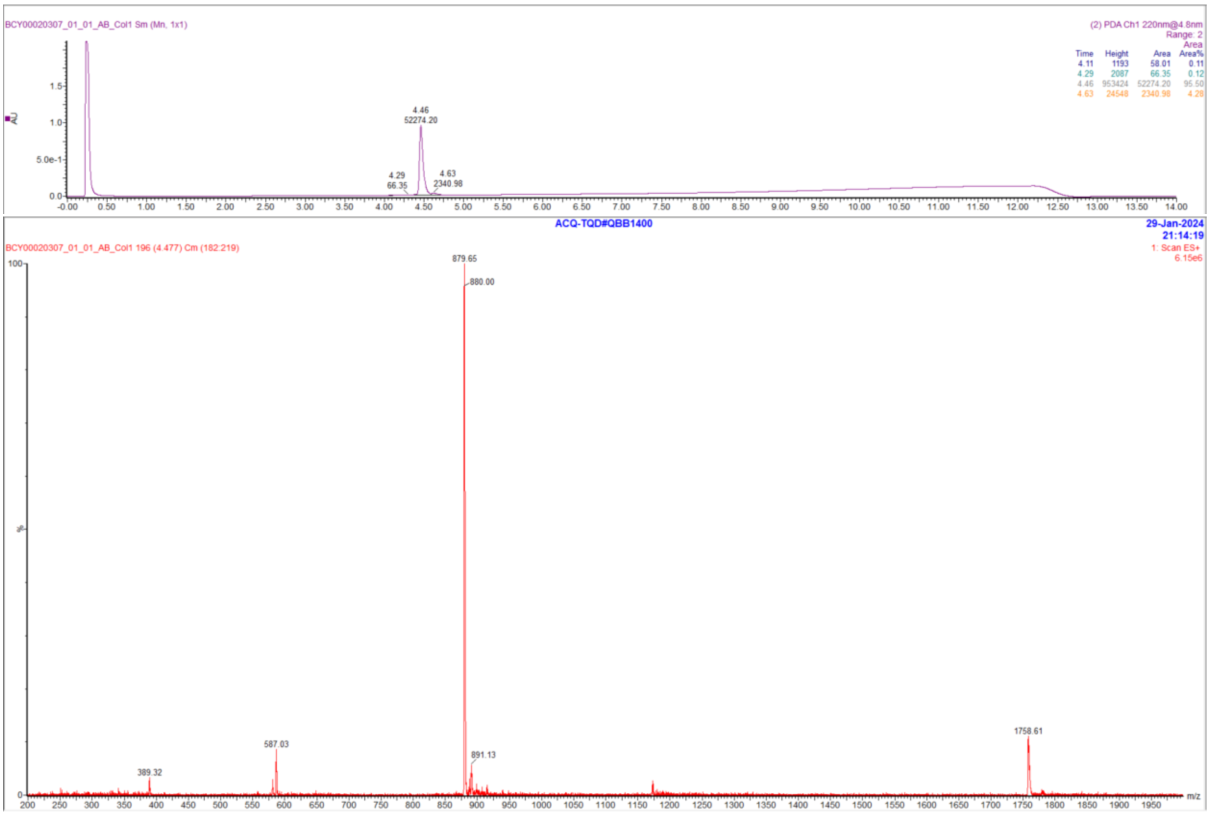

#### Peptide 27

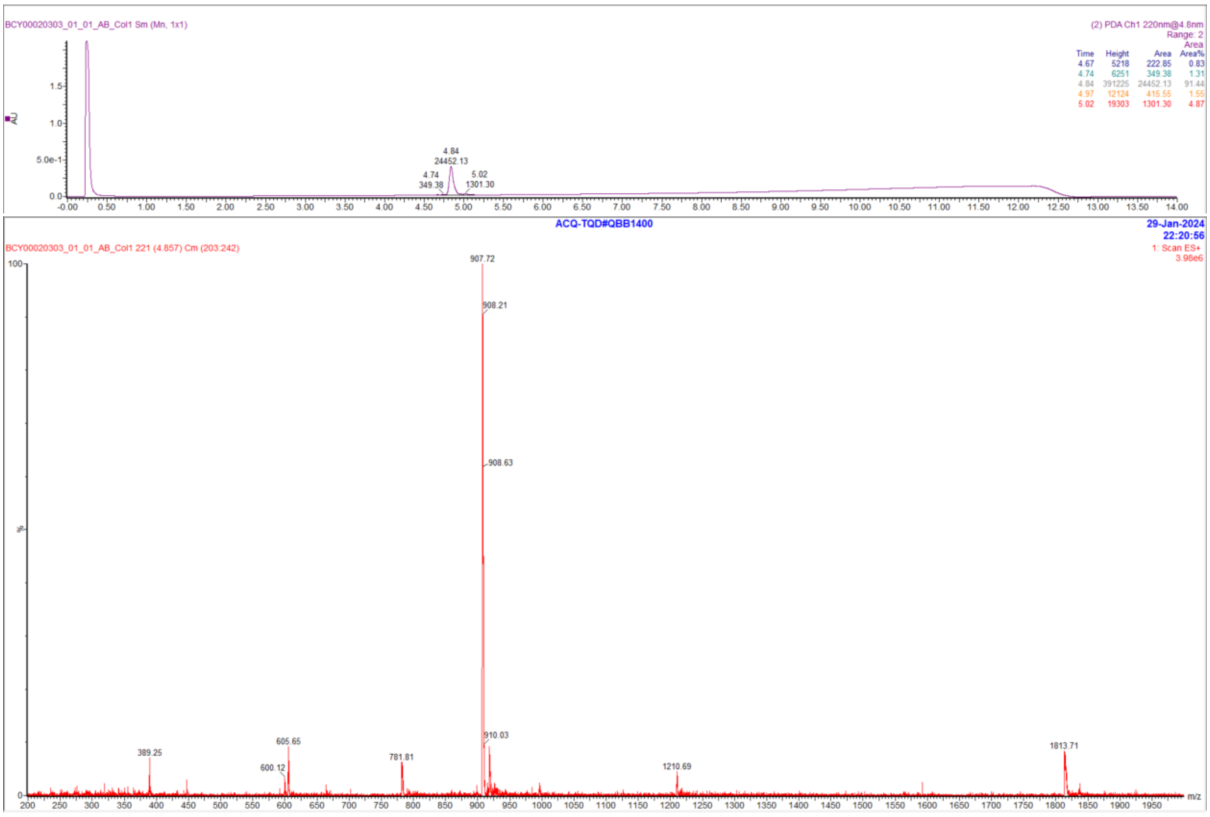

#### Peptide 28

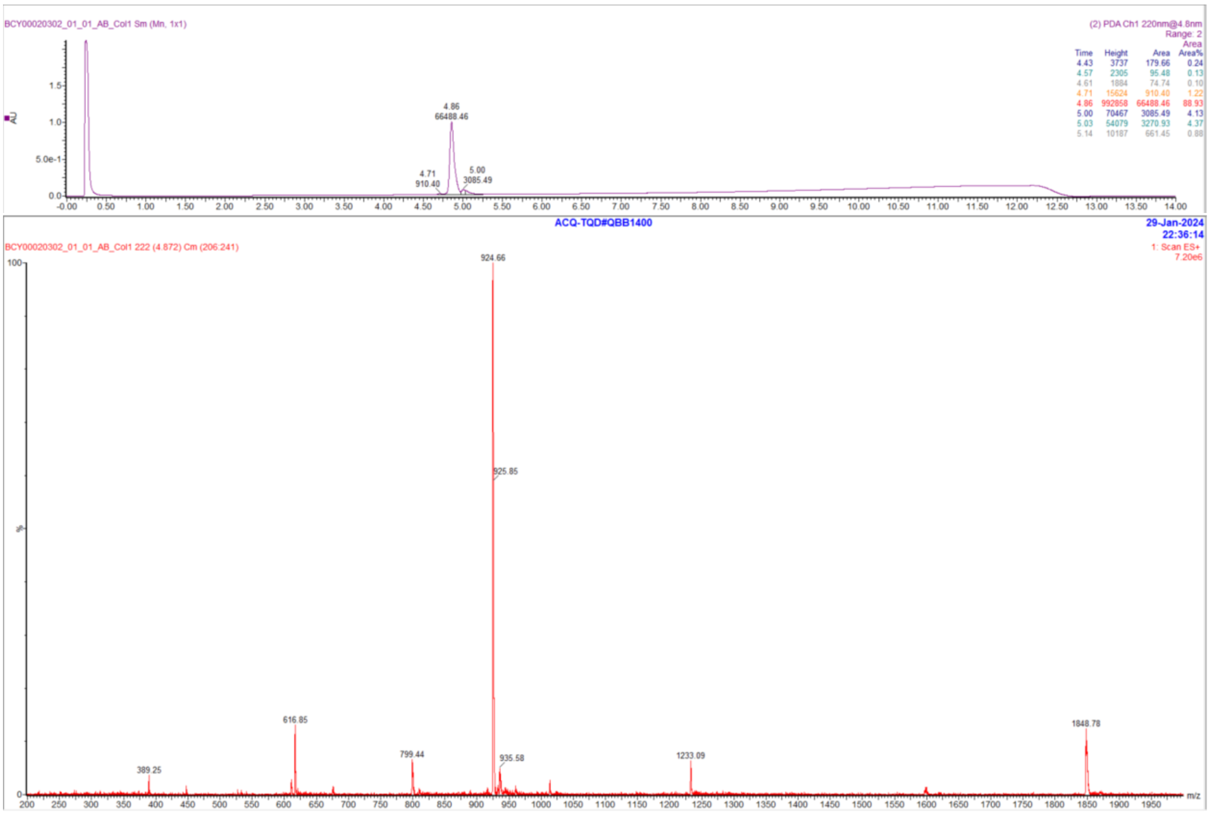

#### Peptide 29

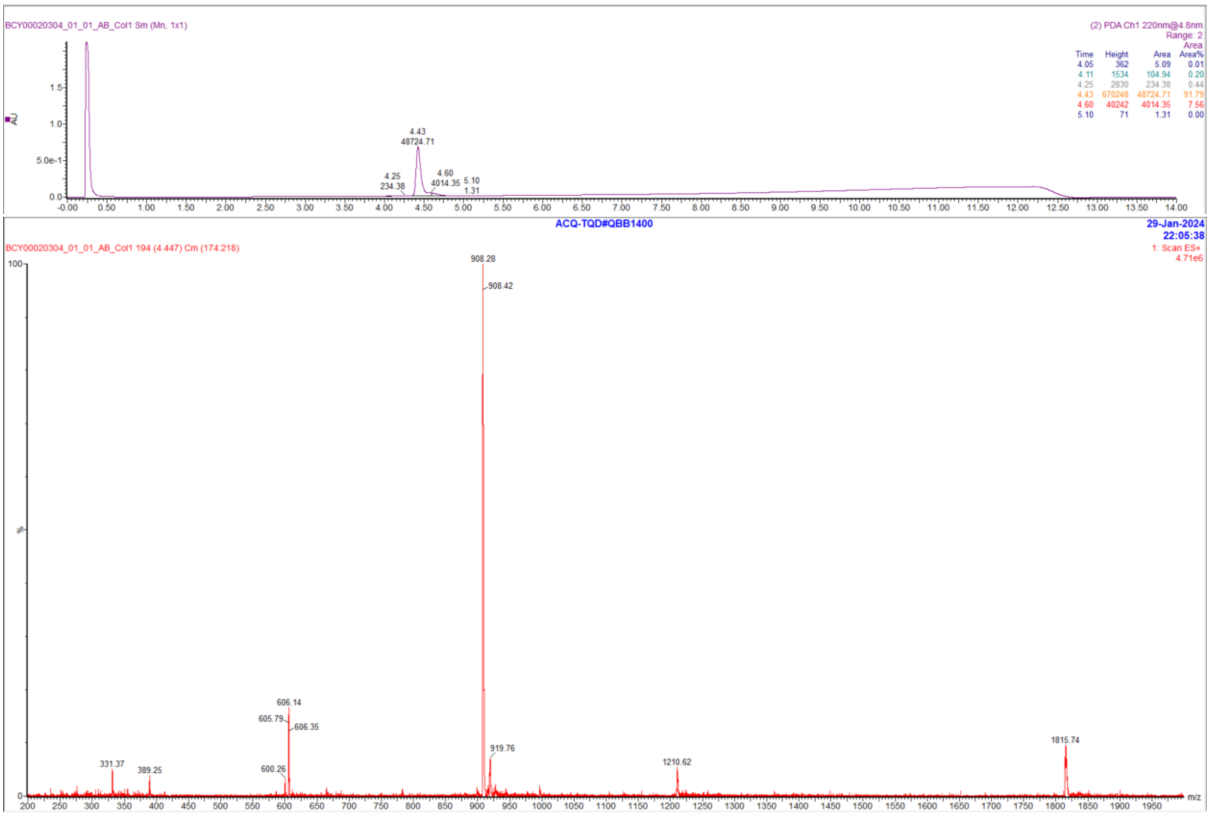

### Peptide 30

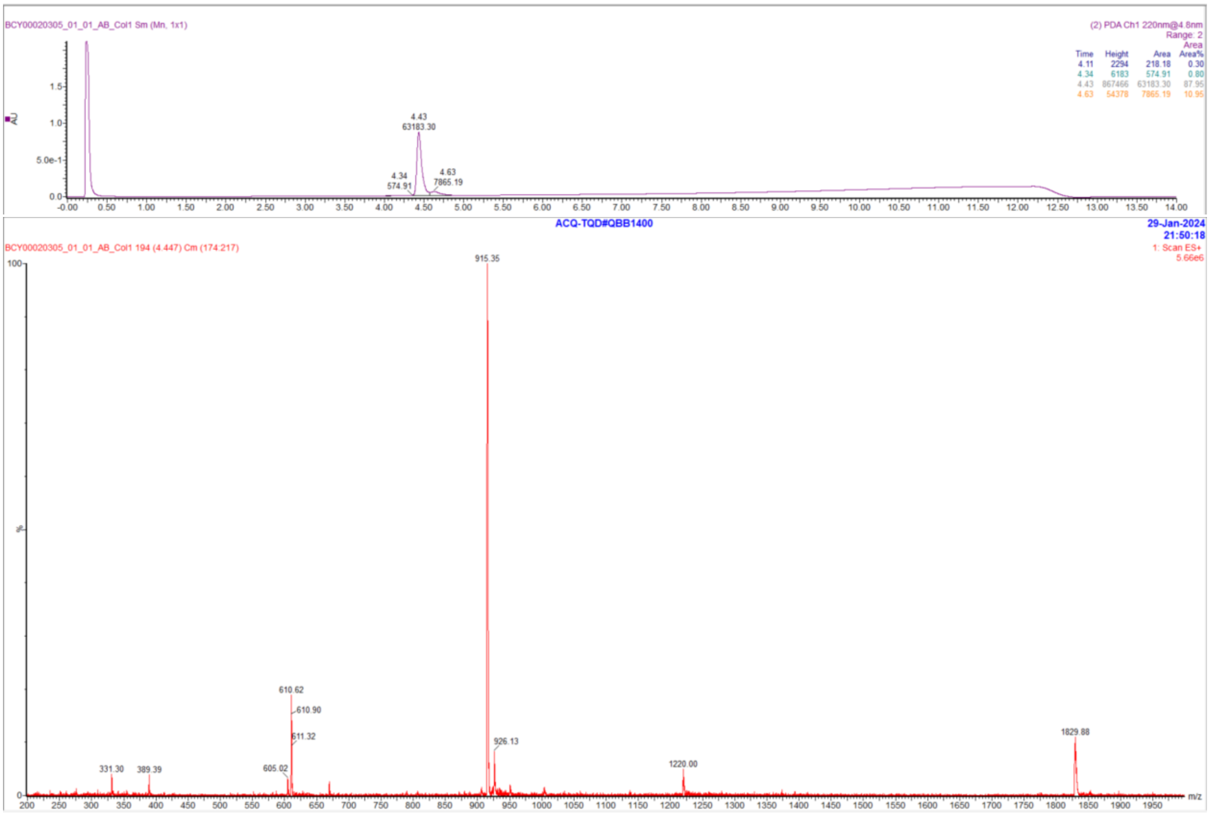

### Peptide 31

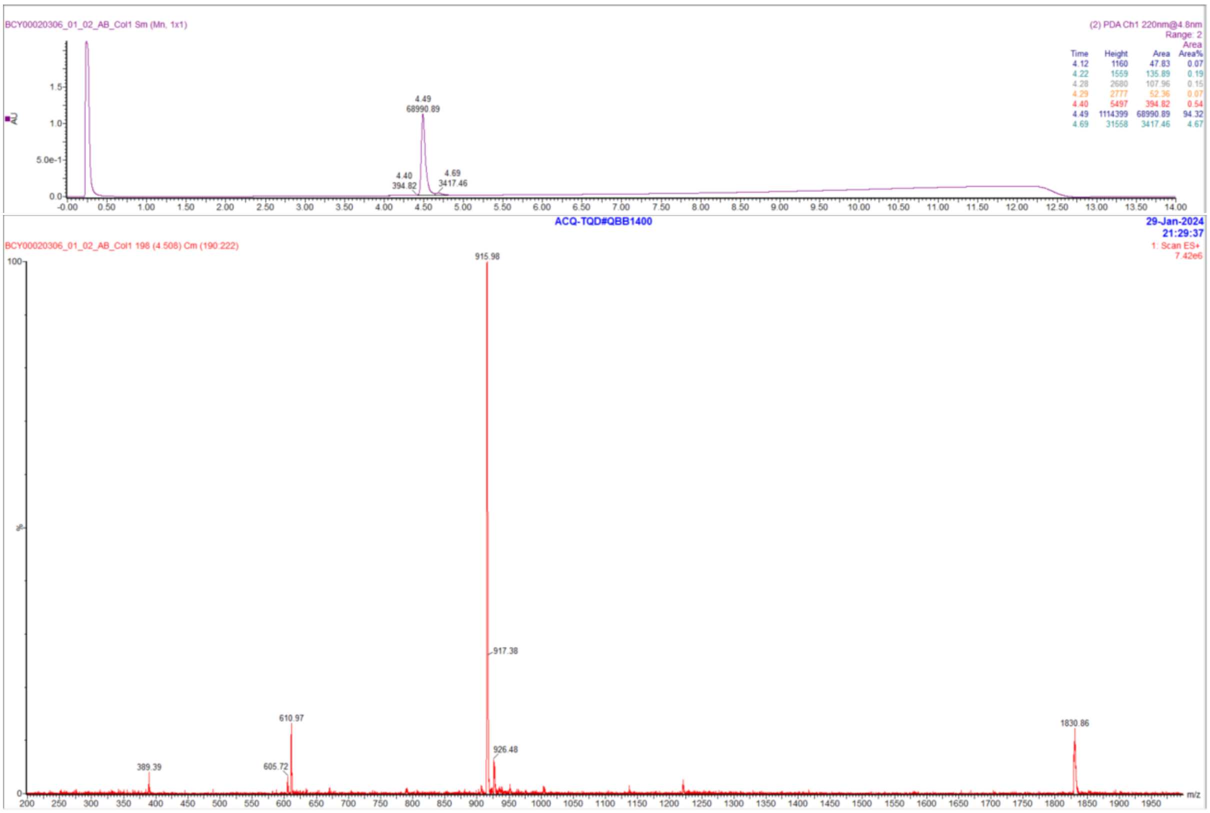

### Peptide 32

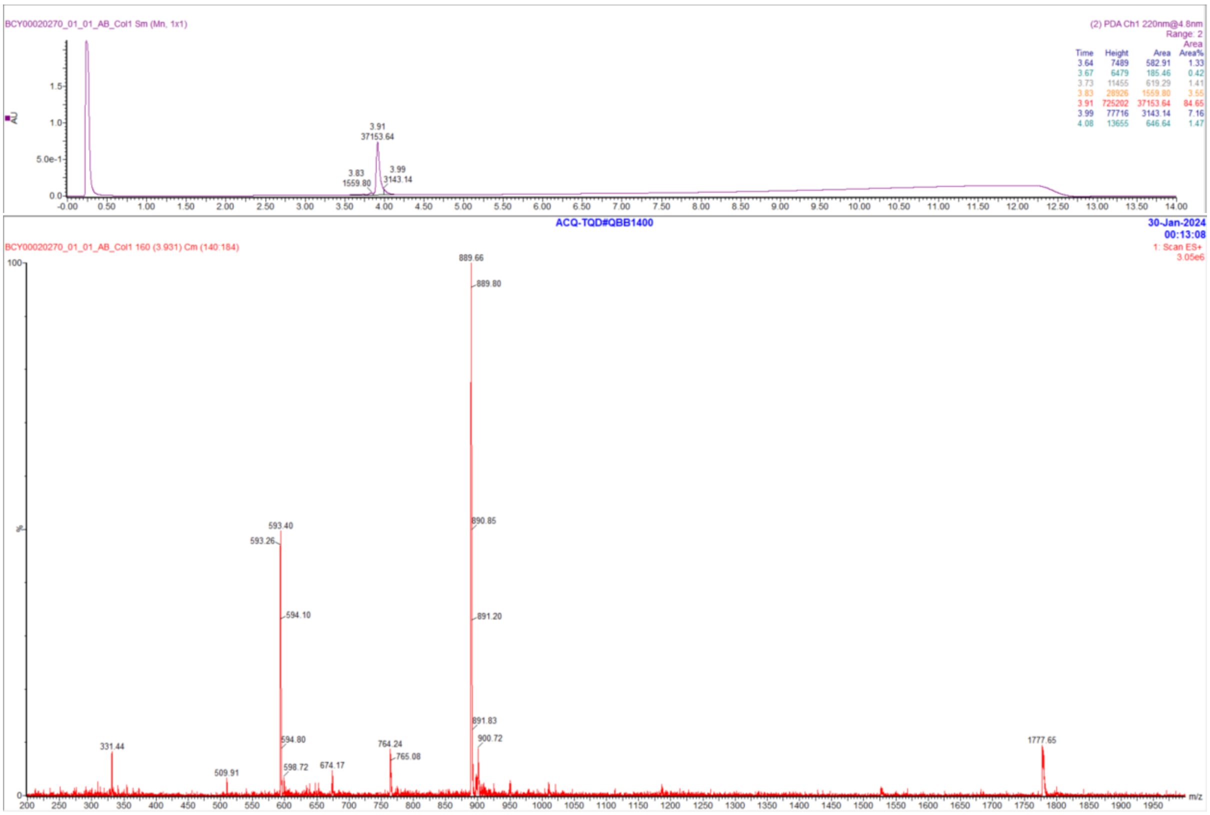

### Peptide 33

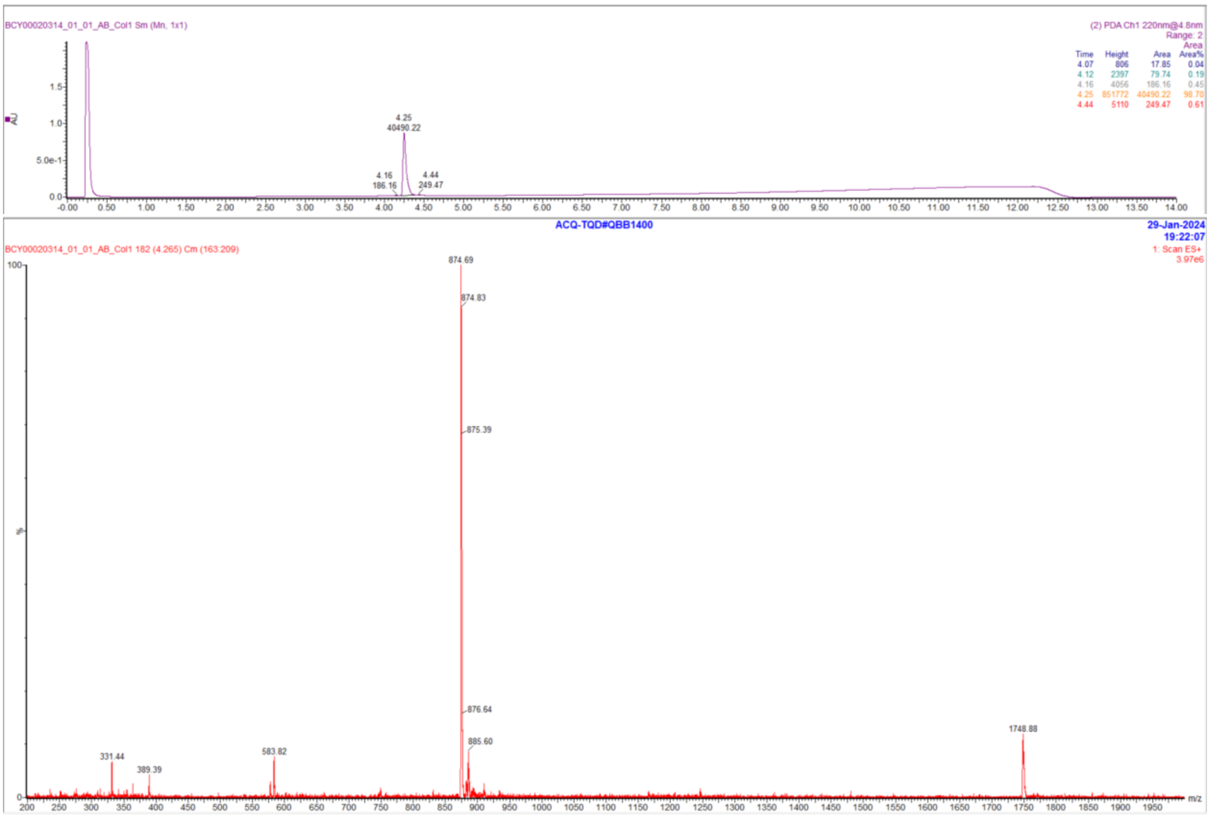

#### Peptide 34

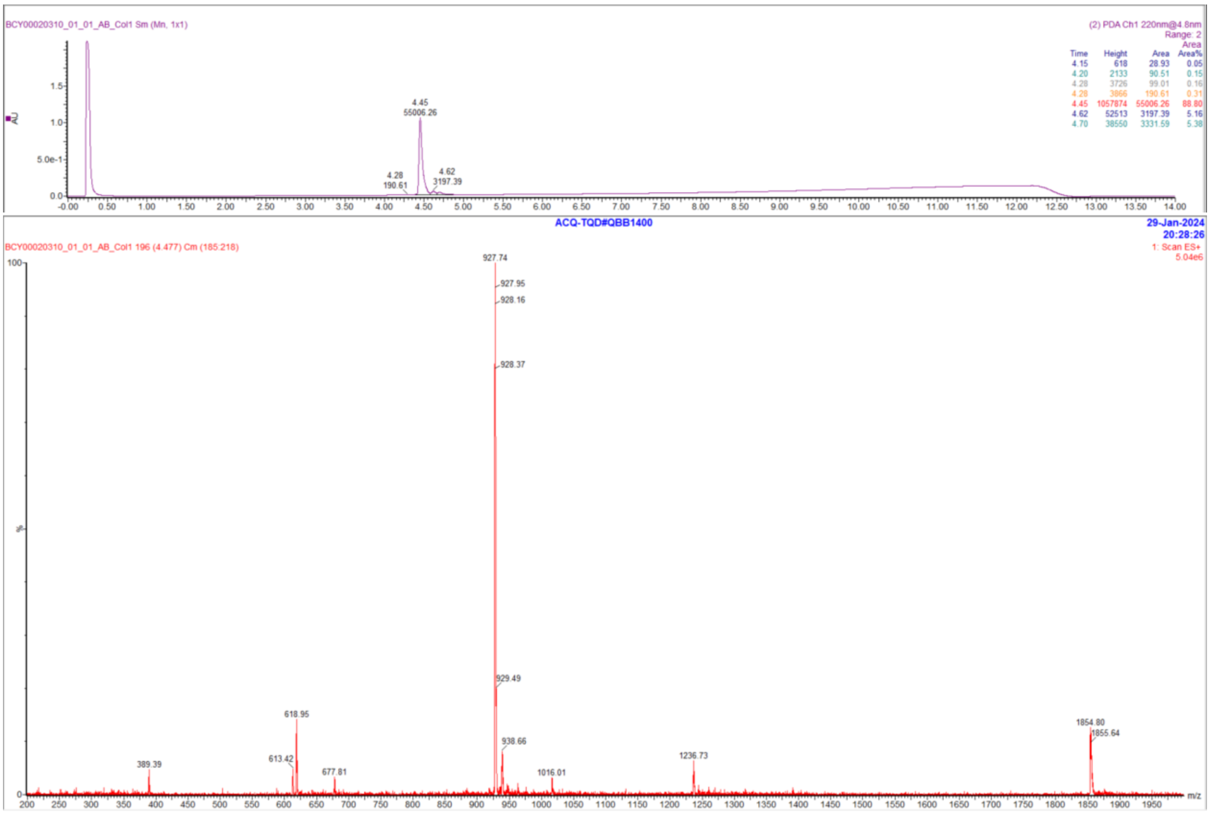
BCY00020310_01_01_AB_Col1 Sm (Mn, 1x1)
(2) PDA Ch1 220nm@4.8nm
Range: 2
Area
Time Height Area Area%
4.15 618 28.93 0.05
4.20 2133 90.51 0.15
4.28 3726 99.01 0.16
4.28 3866 190.61 0.31
4.45 1057874 55006.26 88.80
4.62 52513 3197.39 5.16
4.70 38550 3331.59 5.38
AU
4.45
55006.26
4.28
190.61
4.62
3197.39
ACQ-TQD#QBB1400
29-Jan-2024
20:28:26
BCY00020310_01_01_AB_Col1 196 (4.477) Cm (185:218)
1: Scan ES+
5.04e6
927.74
927.95
928.16
928.37
929.49
938.66
618.95
613.42
389.39
677.81
1016.01
1236.73
1854.80
1855.64
%
m/z

#### Peptide 35

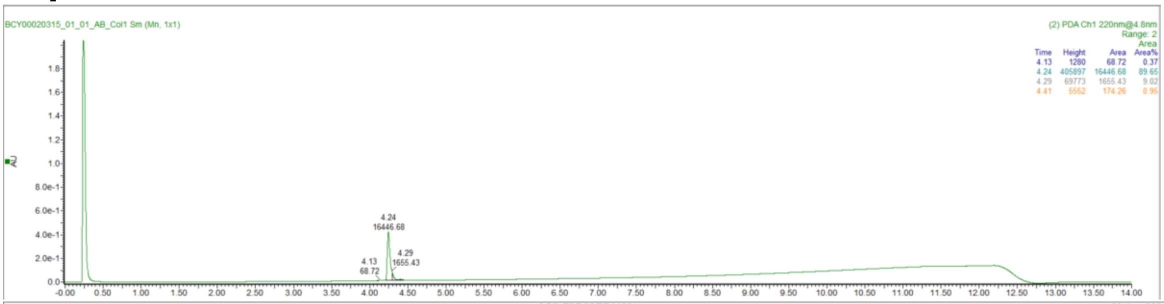

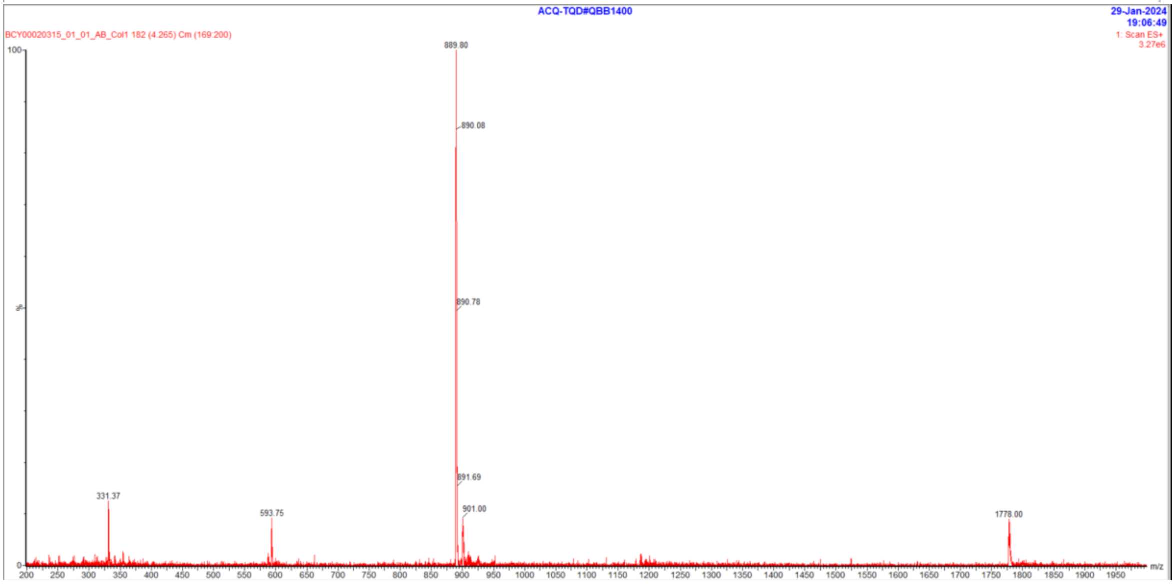

#### Peptide 36

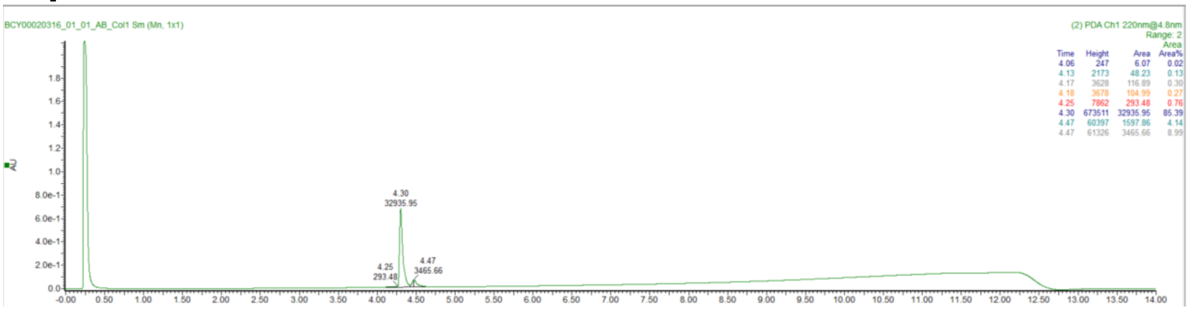

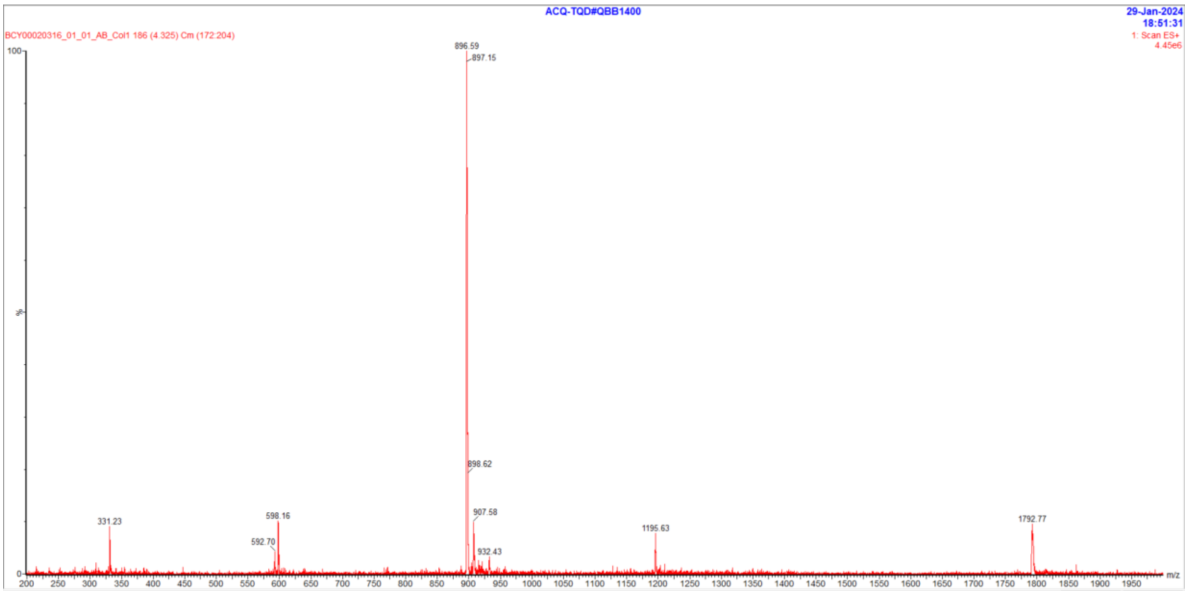
ACQ-TQD#QBB1400
29-Jan-2024
18:51:31
BCY00020316_01_01_AB_Col1 186 (4.325) Cm (172:204)
1: Scan ES+
4.45e6
896.59
897.15
898.62
907.58
932.43
331.23
598.16
592.70
1195.63
1792.77
m/z

### Peptide 37

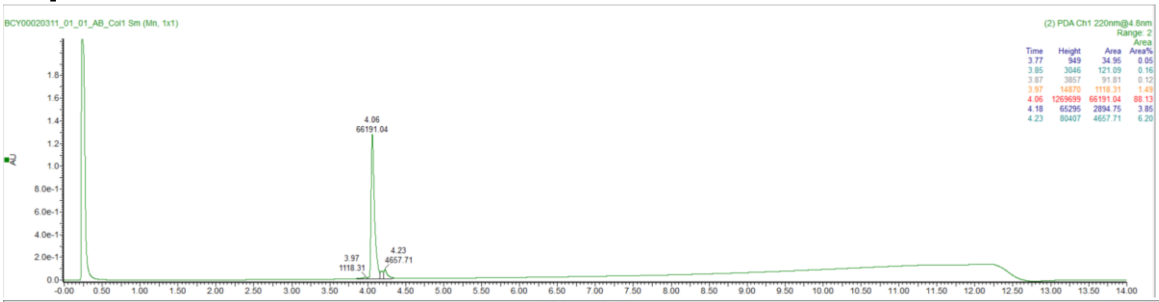
BCY00020311_01_01_AB_Col1 Sm (Mn, 1x1)
(2) PDA Ch1 220nm@4.8nm
4.06
66191.04
3.97
1118.31
4.23
4657.71

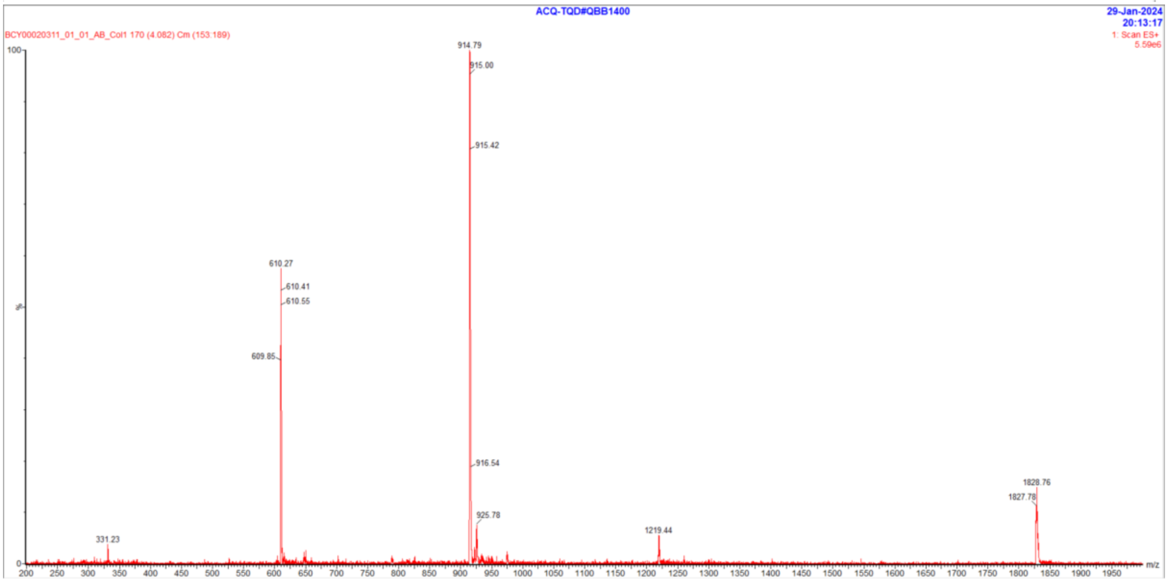
ACQ-TQD#QBB1400
29-Jan-2024
20:13:17
BCY00020311_01_01_AB_Col1 170 (4.082) Cm (153:189)
1: Scan ES+
5.59e6
914.79
915.00
915.42
916.54
925.78
610.27
610.41
610.55
609.85
331.23
1219.44
1828.76
1827.78
m/z

### Peptide 38

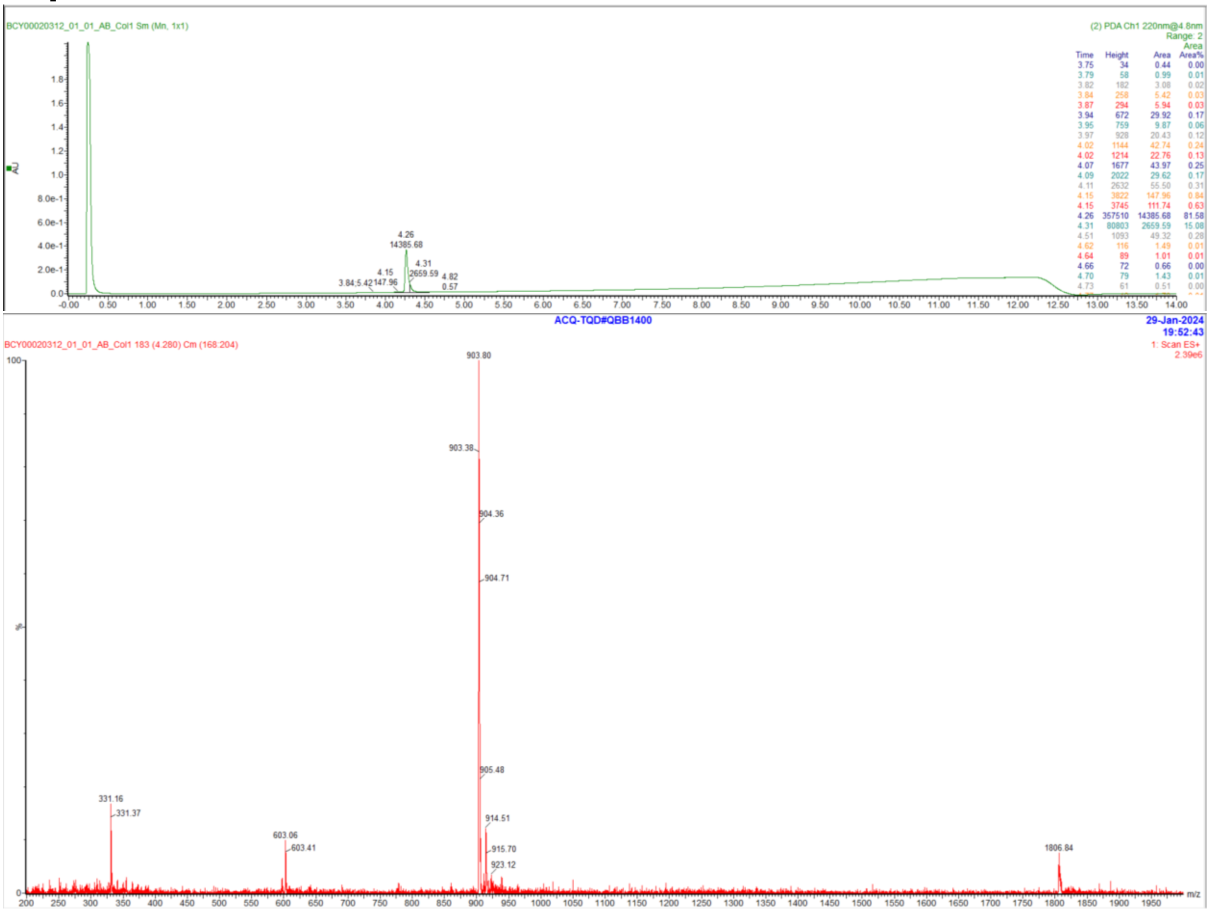
BCY00020312_01_01_AB_Col1 Sm (Mn, 1x1)
(2) PDA Ch1 220nm@4.8nm
Range: 2
Time Height Area Area%
3.75 34 0.44 0.00
3.79 58 0.99 0.01
3.82 182 3.08 0.02
3.84 258 5.42 0.03
3.87 294 5.94 0.03
3.94 672 29.92 0.17
3.95 759 9.87 0.06
3.97 928 20.43 0.12
4.02 1144 42.74 0.24
4.02 1214 22.76 0.13
4.07 1677 43.97 0.25
4.09 2022 29.62 0.17
4.11 2632 55.50 0.31
4.15 3822 147.96 0.84
4.15 3745 111.74 0.63
4.26 357510 14385.68 81.58
4.31 80803 2659.59 15.08
4.51 1093 49.32 0.28
4.62 116 1.49 0.01
4.64 89 1.01 0.01
4.66 72 0.66 0.00
4.70 79 1.43 0.01
4.73 61 0.51 0.00
AU
ACQ-TQD#QBB1400
29-Jan-2024
19:52:43
BCY00020312_01_01_AB_Col1 183 (4.280) Cm (168:204)
1: Scan ES+
2.39e6
903.80
903.38
904.36
904.71
905.48
914.51
915.70
923.12
331.16
331.37
603.06
603.41
1806.84
m/z

#### Peptide 39

| | |
|---|---|
| Sample name: | 101-08-26 (TATA) QC |
| Instrument: | 1260_2 |
| Injection date: | 12/11/2019 2:17:40 PM |
| Acq. method: | 0595B_AB_Poroshell120_15.5min.M |

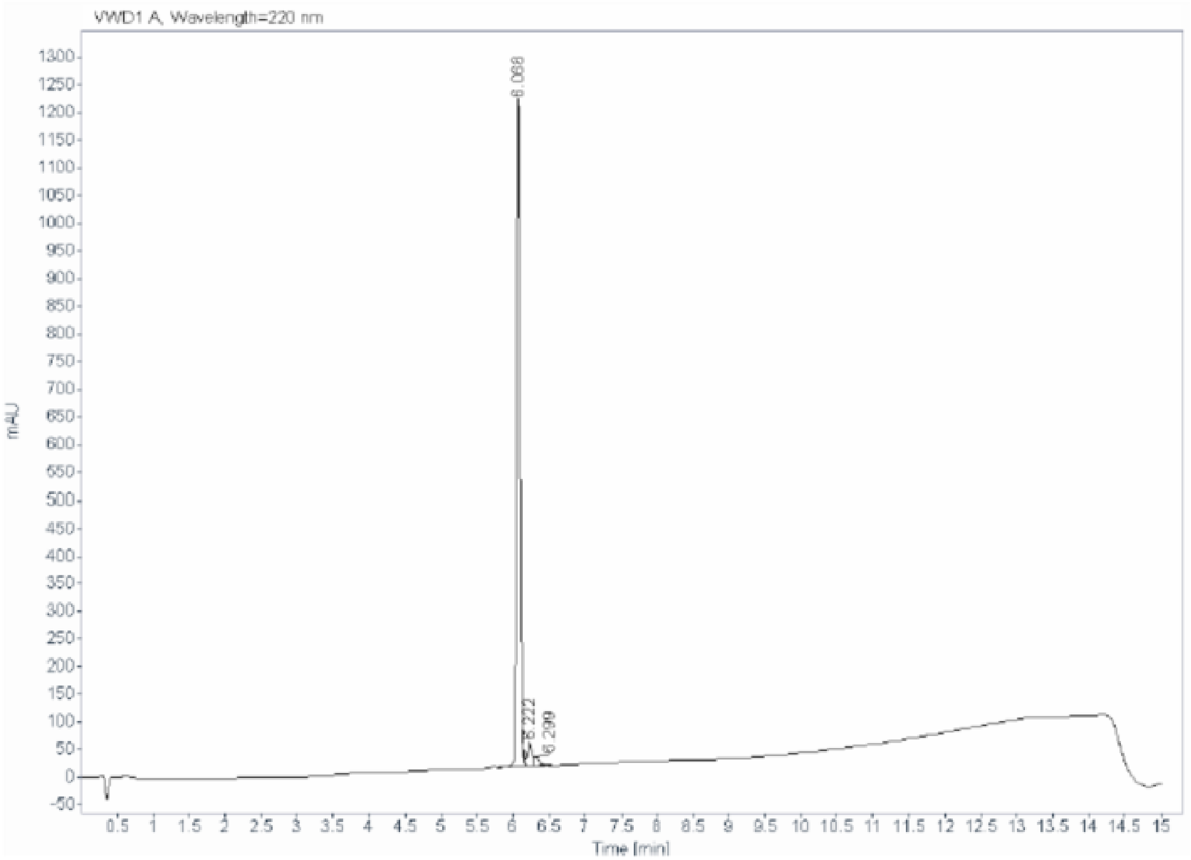

Signal: VWD1 A, Wavelength=220 nm

| RT [min] | Type | Width [min] | Area | Height | Area% |
|---|---|---|---|---|---|
| 6.068 | BV R | 0.0520 | 4026.8979 | 1203.2932 | 93.4396 |
| 6.222 | VV E | 0.0667 | 190.5921 | 41.4232 | 4.4225 |
| 6.299 | VB E | 0.0834 | 92.1372 | 16.9520 | 2.1379 |
| | | Sum | 4309.6273 | | |

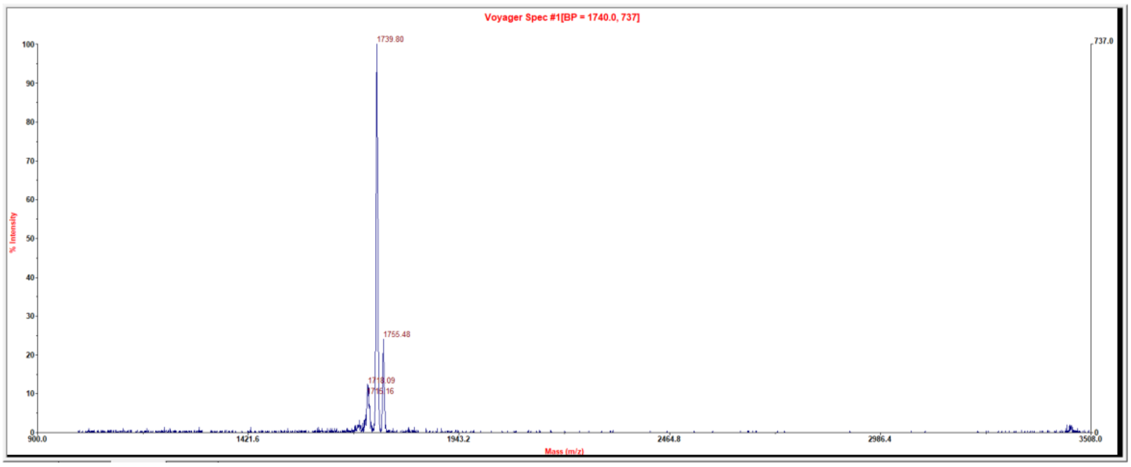

### Peptide 40

**Sample name:** 101-08-29 (TATA) QC
**Instrument:** 1260_2
**Injection date:** 12/11/2019 3:21:52 PM
**Acq. method:** 0595B_AB_Poroshell120_15.5min.M

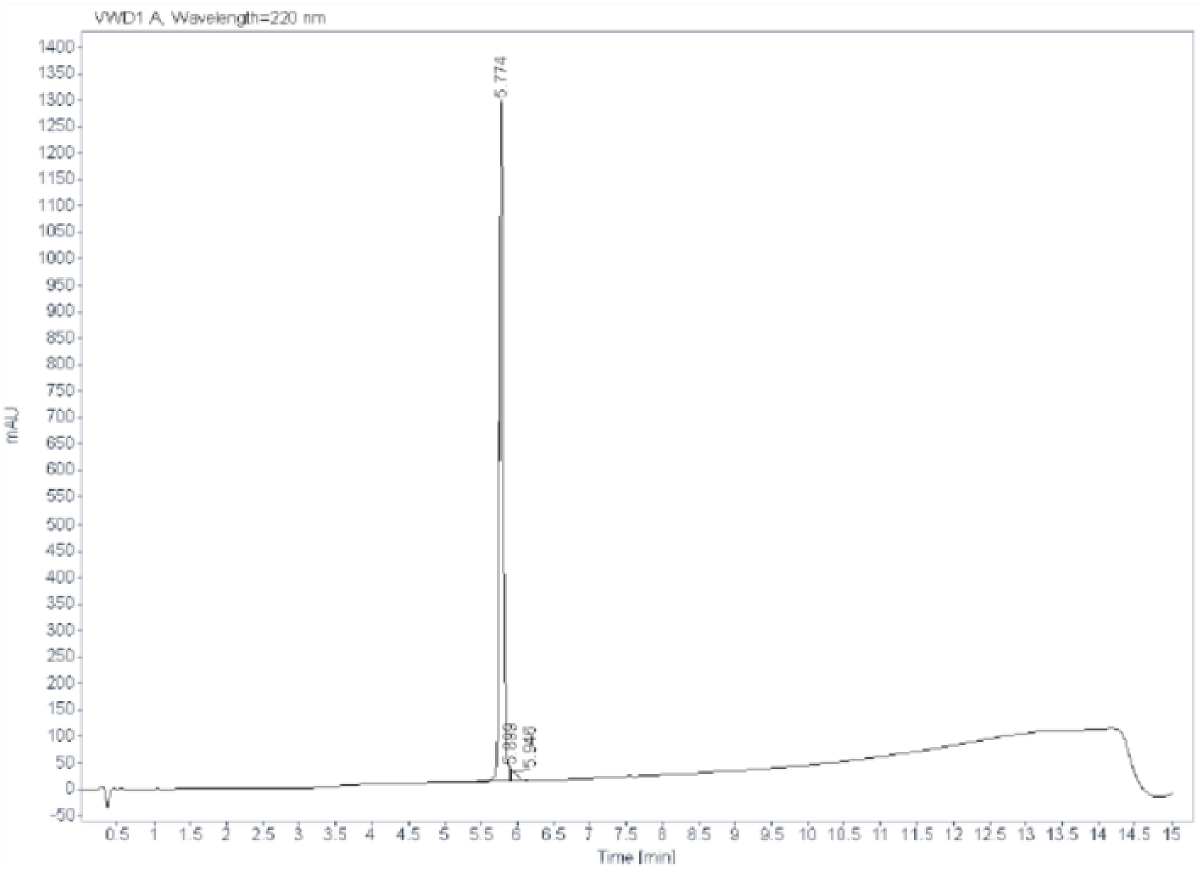

**Signal:** VWD1 A, Wavelength=220 nm

| RT [min] | Type | Width [min] | Area | Height | Area% |
|---|---|---|---|---|---|
| 5.774 | VF R | 0.0617 | 5143.9707 | 1284.1719 | 97.8242 |
| 5.899 | VV | 0.0157 | 22.5725 | 24.0308 | 0.4293 |
| 5.946 | VV B | 0.0747 | 91.8406 | 17.0967 | 1.7466 |
| | | Sum | 5258.3838 | | |

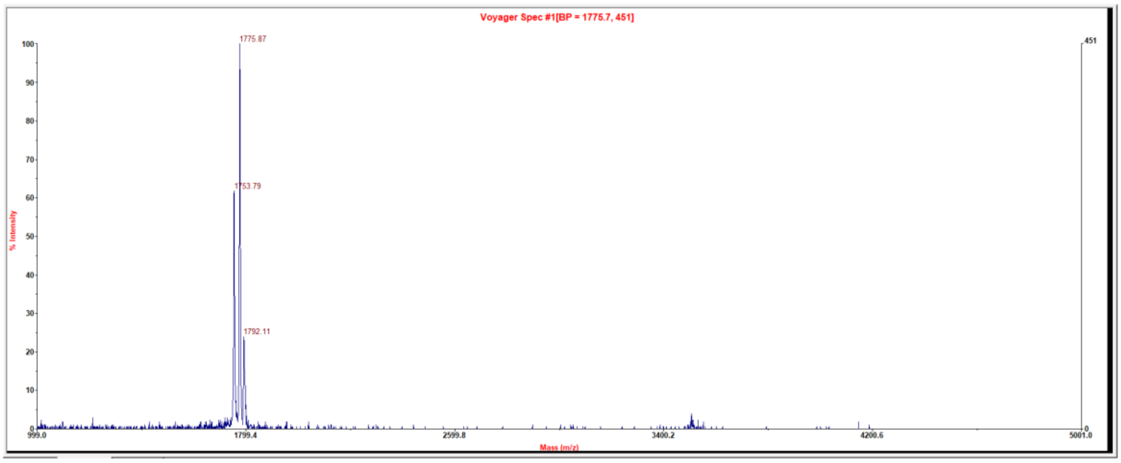

### Peptide 41

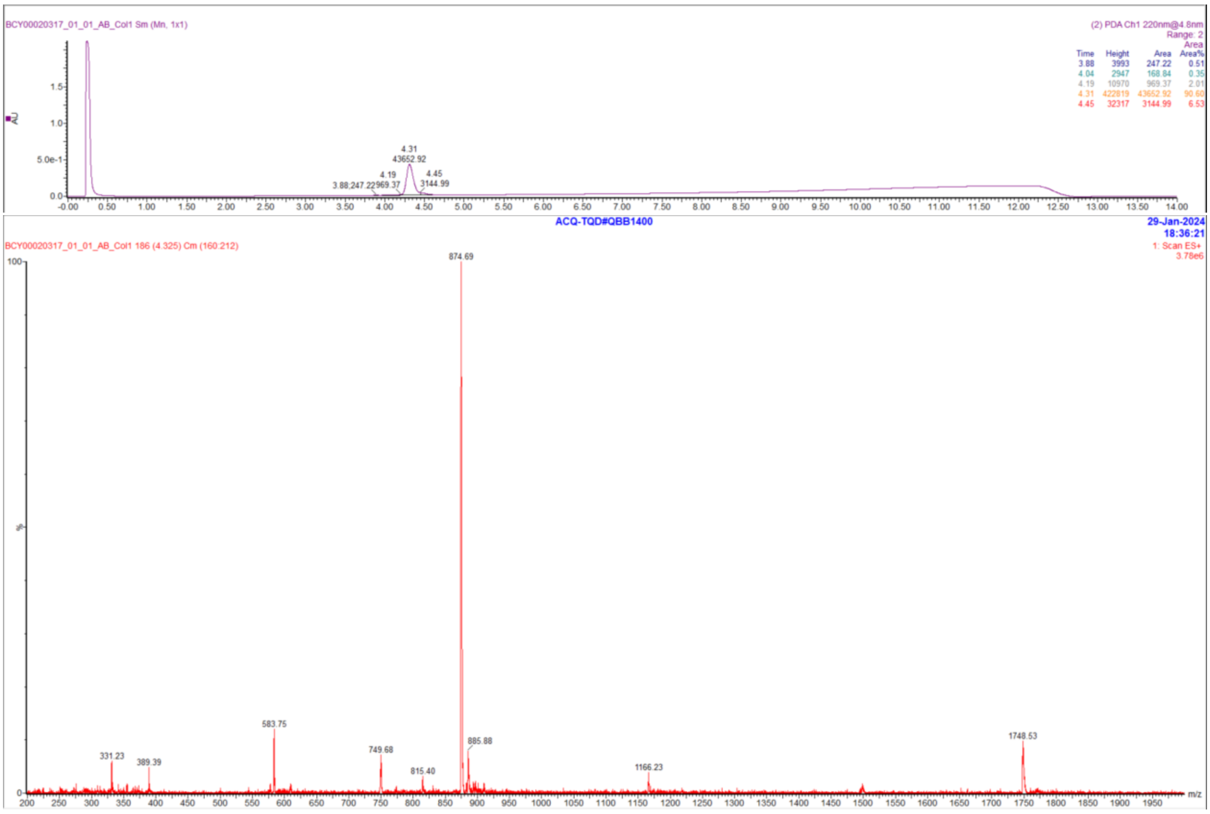

### Peptide 42

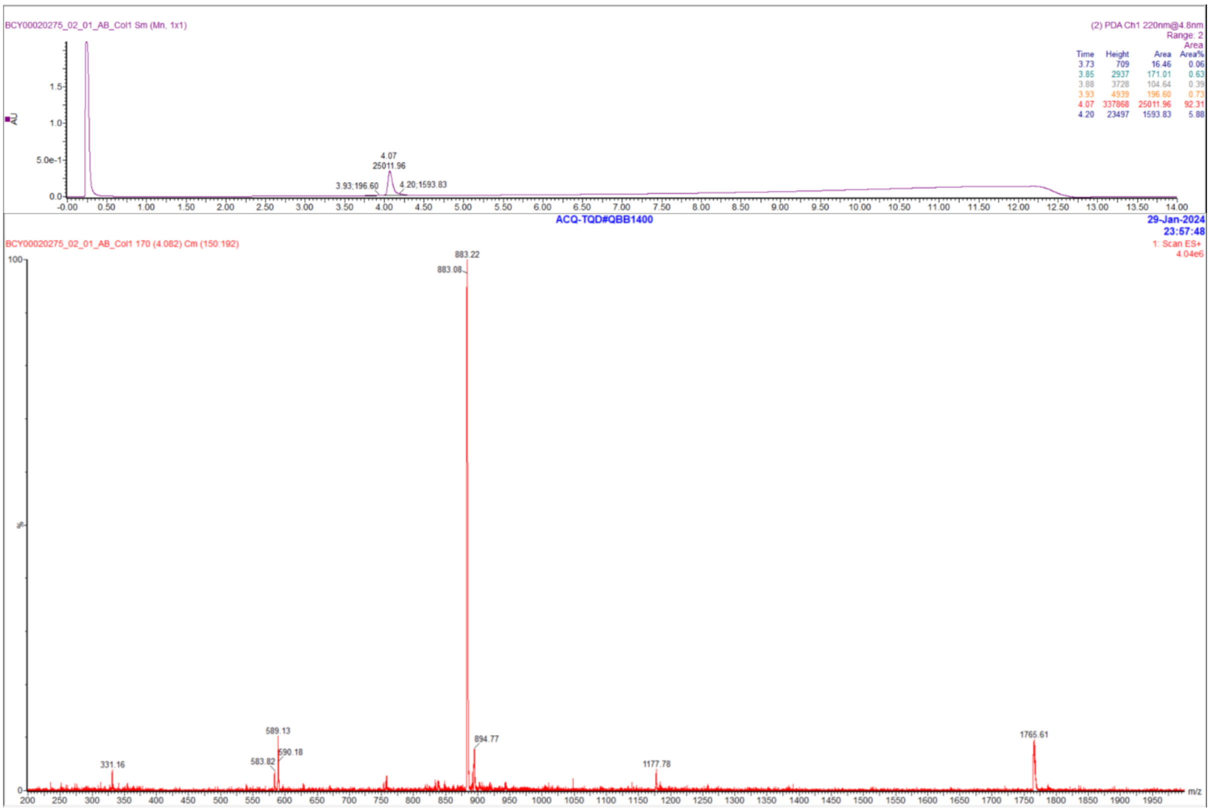

#### Peptide 43

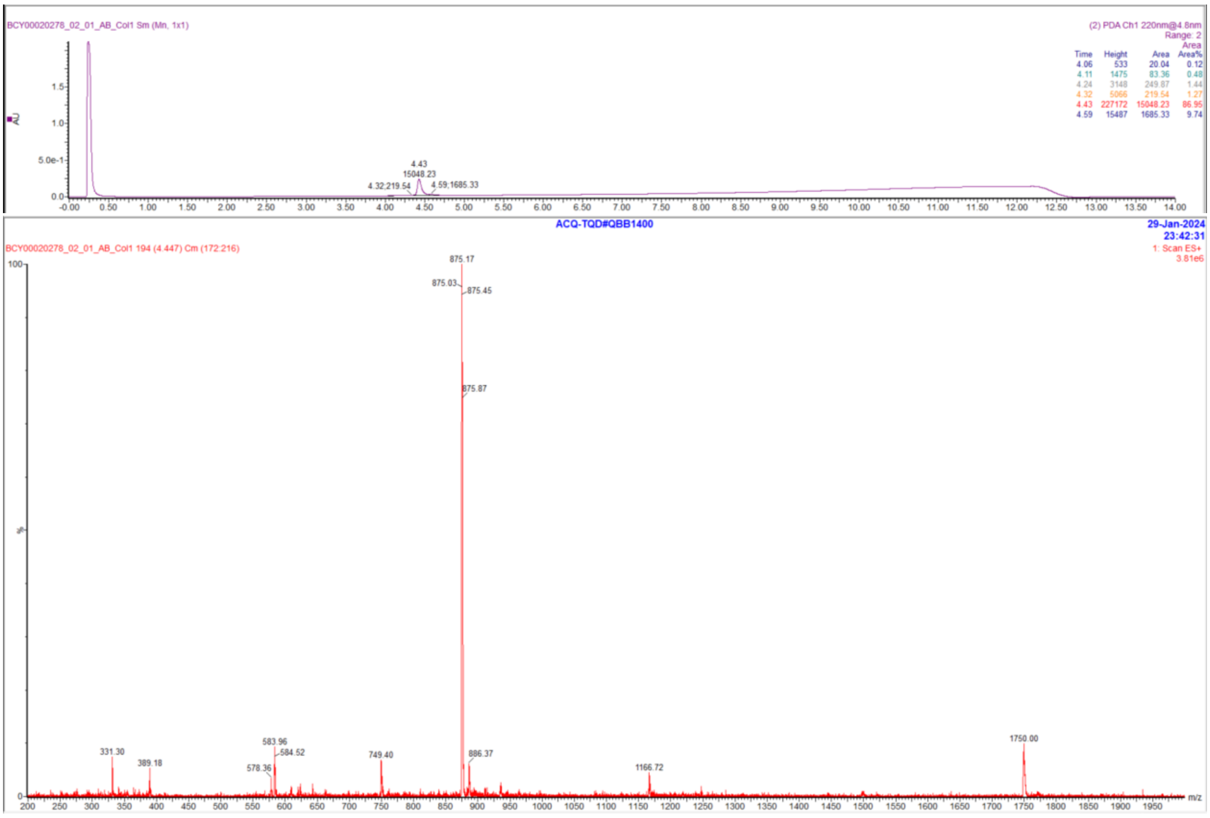

#### Peptide 44

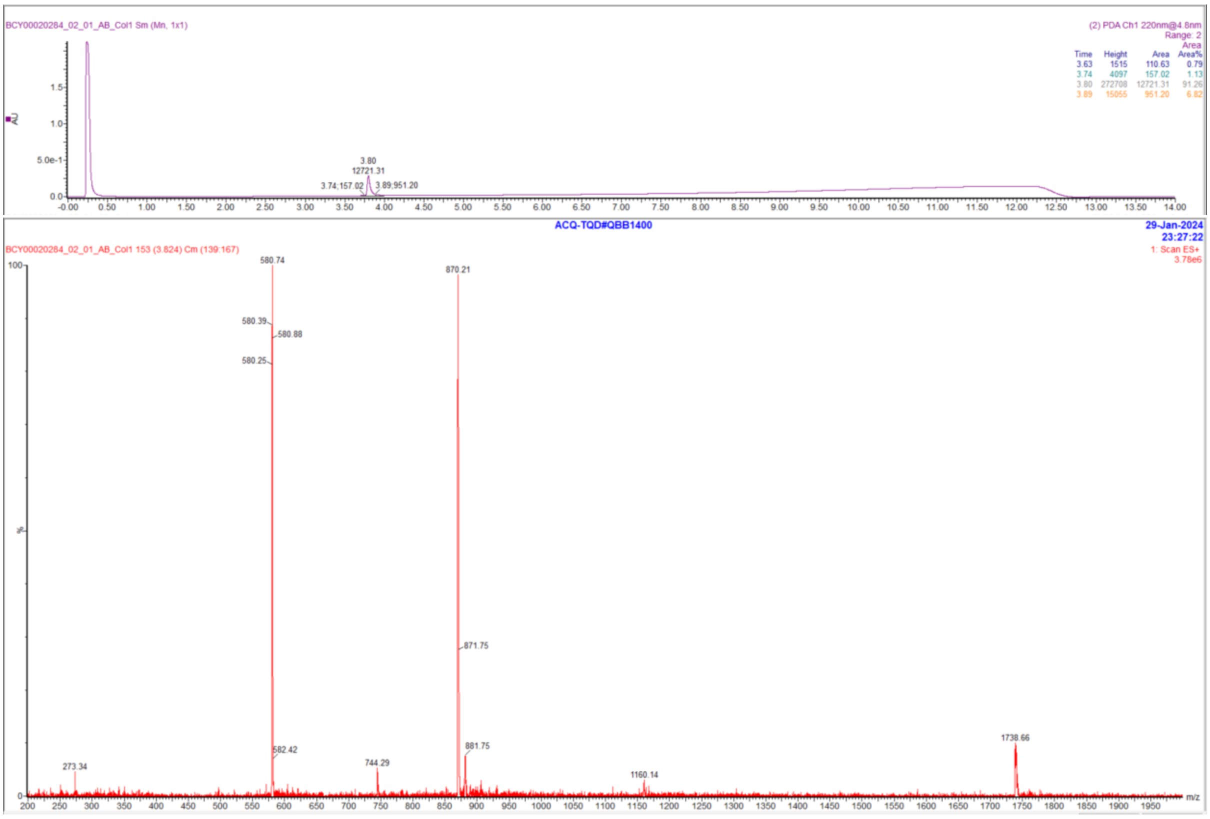

#### Peptide 45

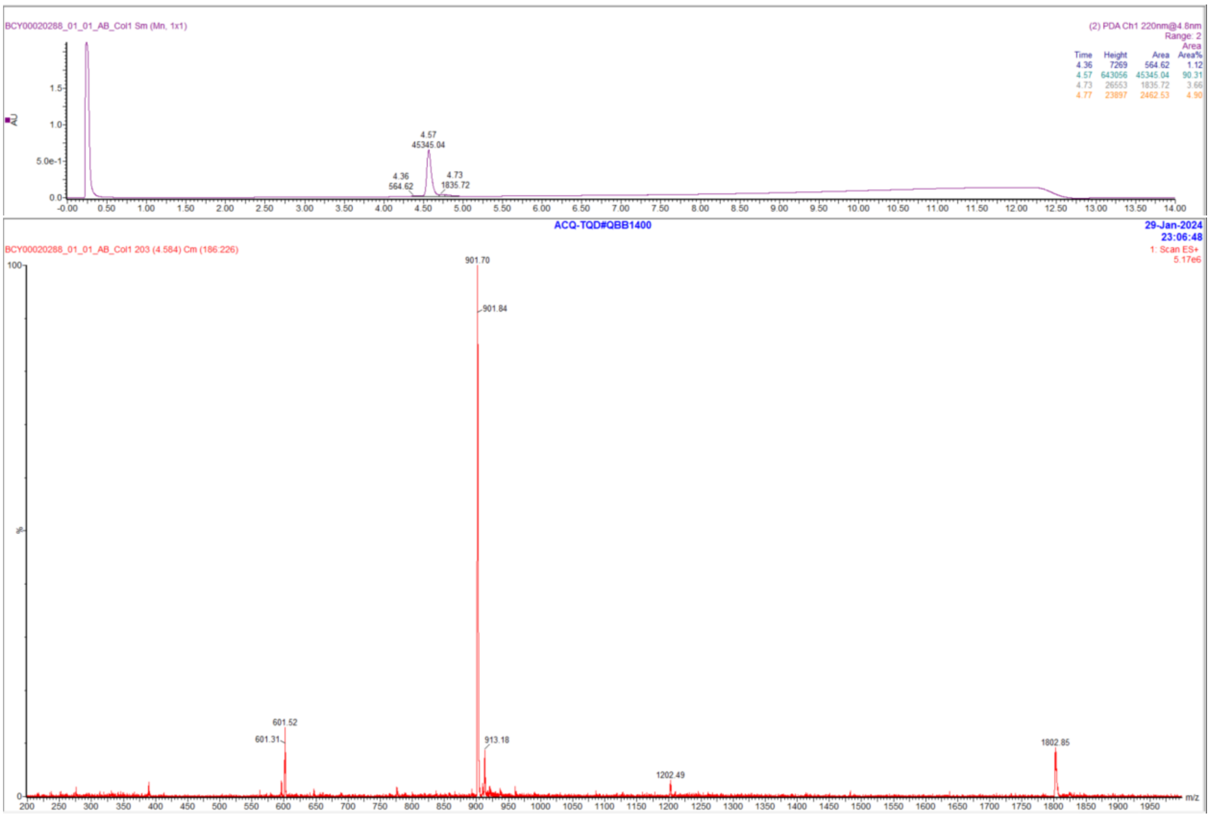

#### Peptide 46

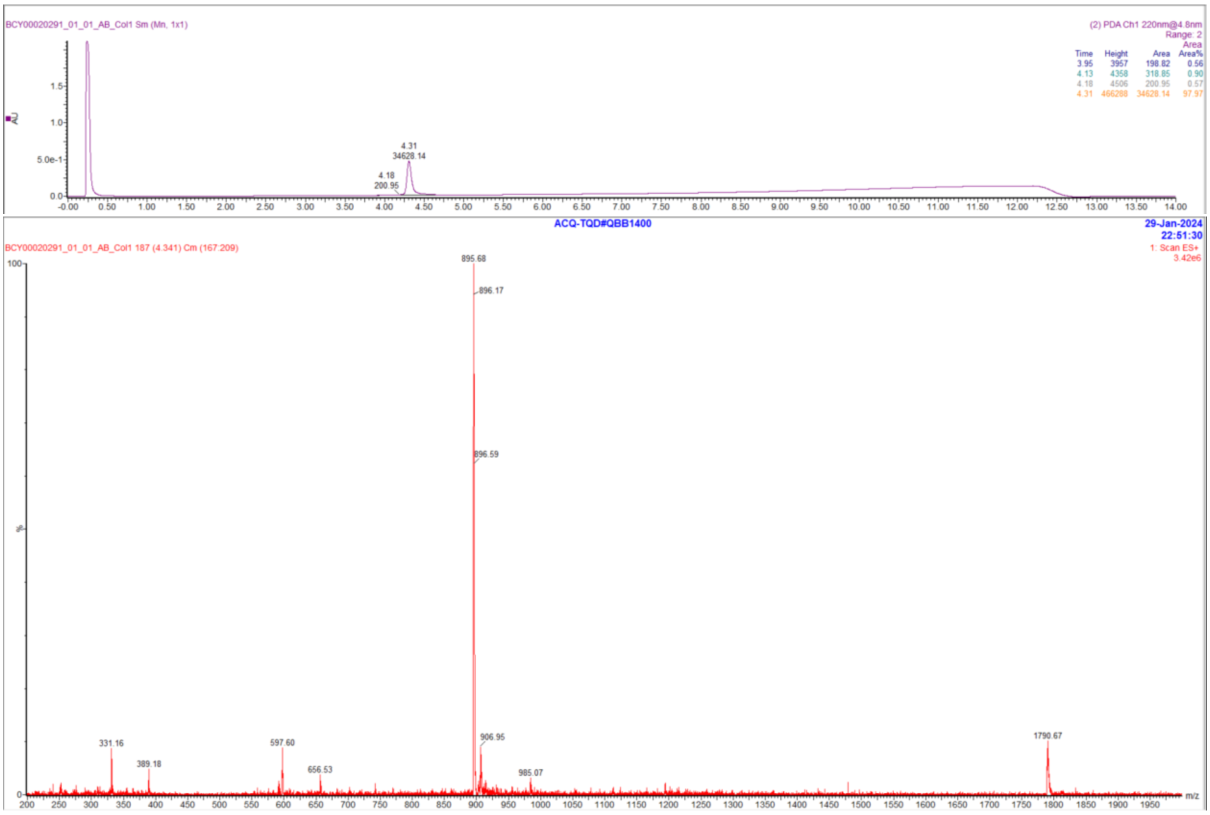

#### Peptide 47

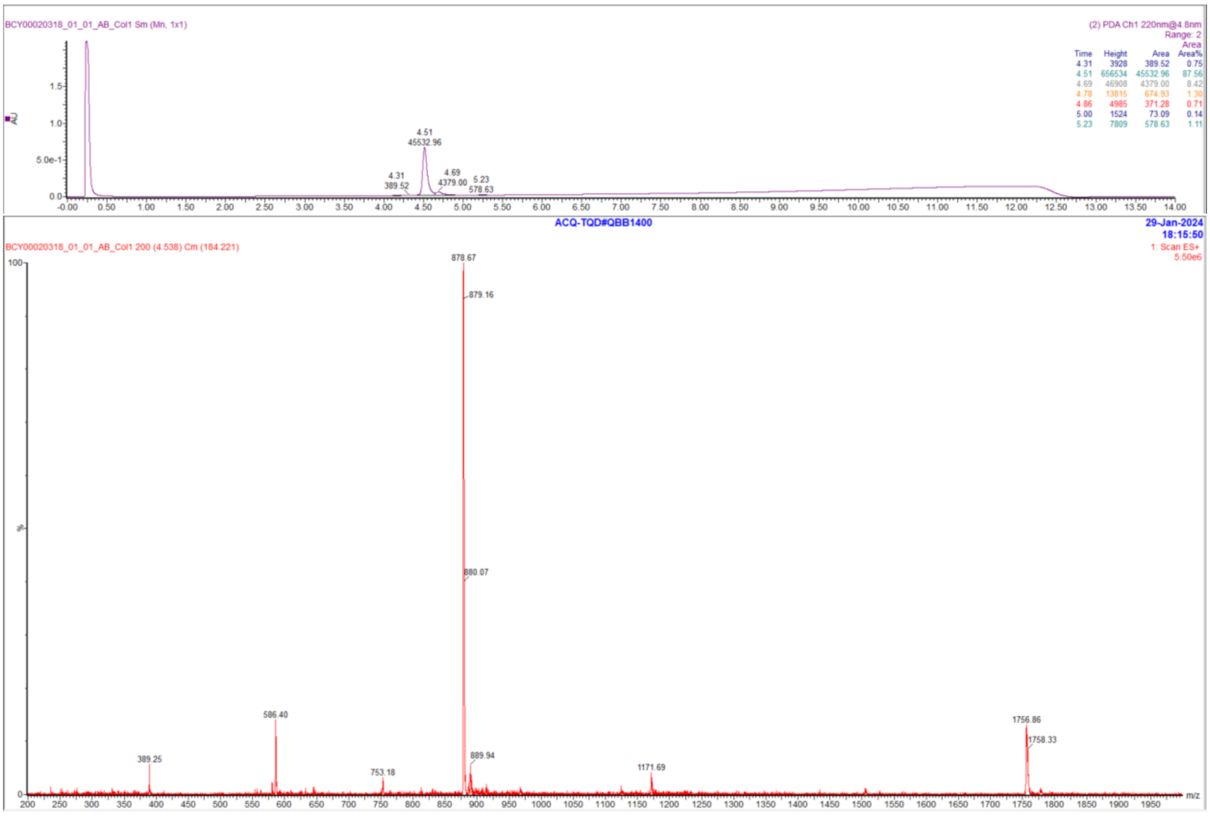

#### Peptide 48

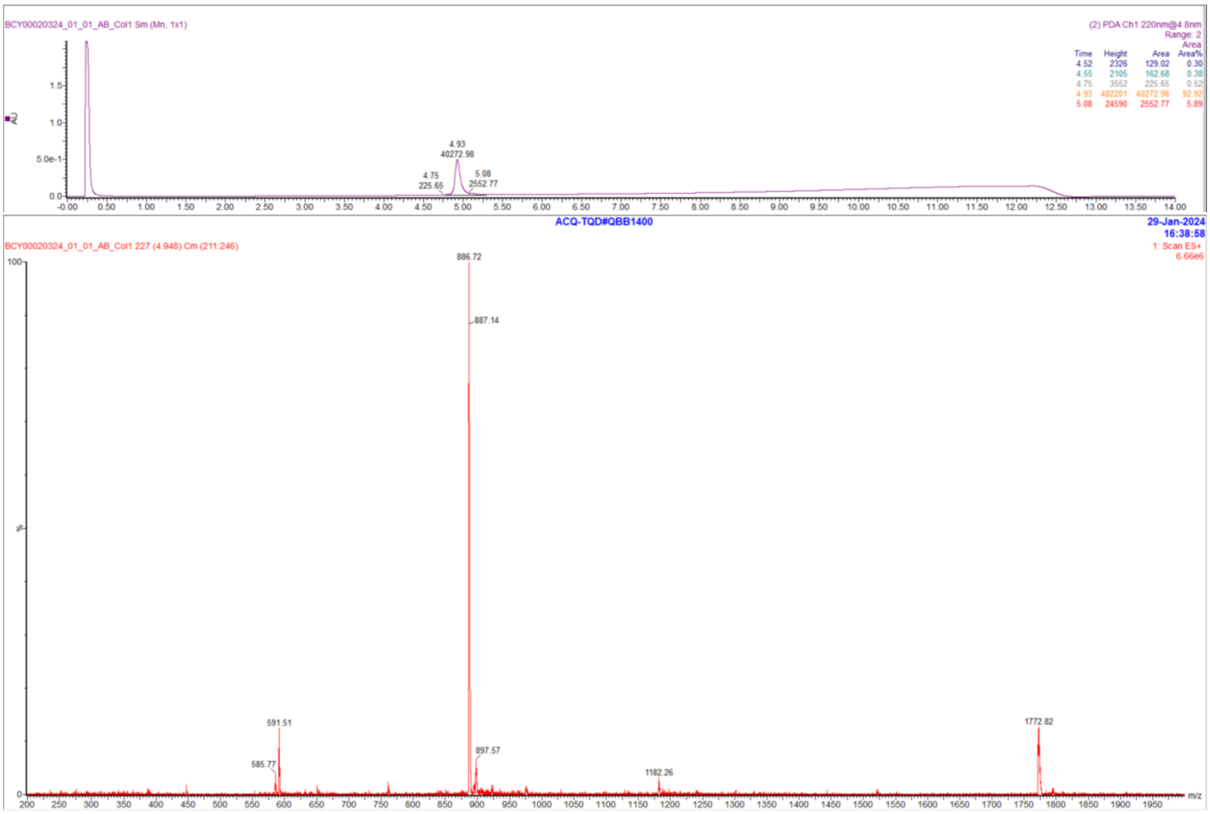

### Peptide 49

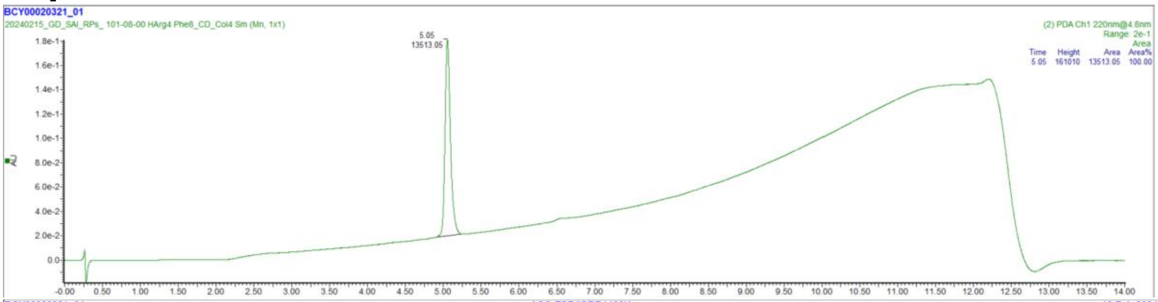

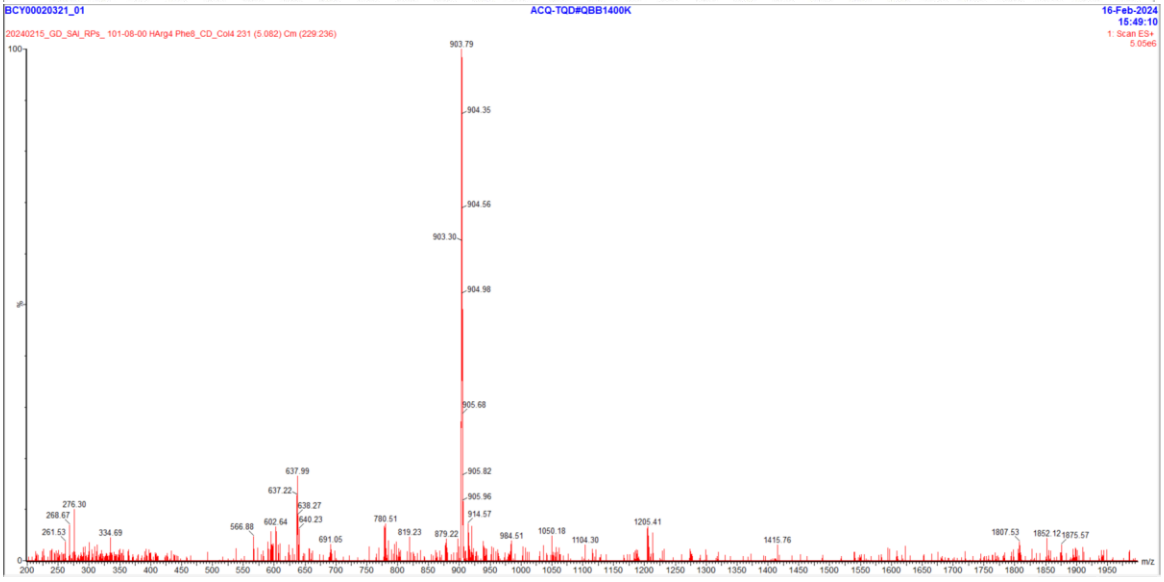

### Peptide 50

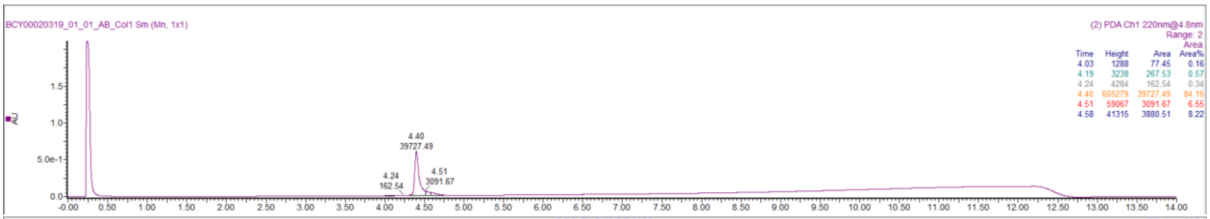

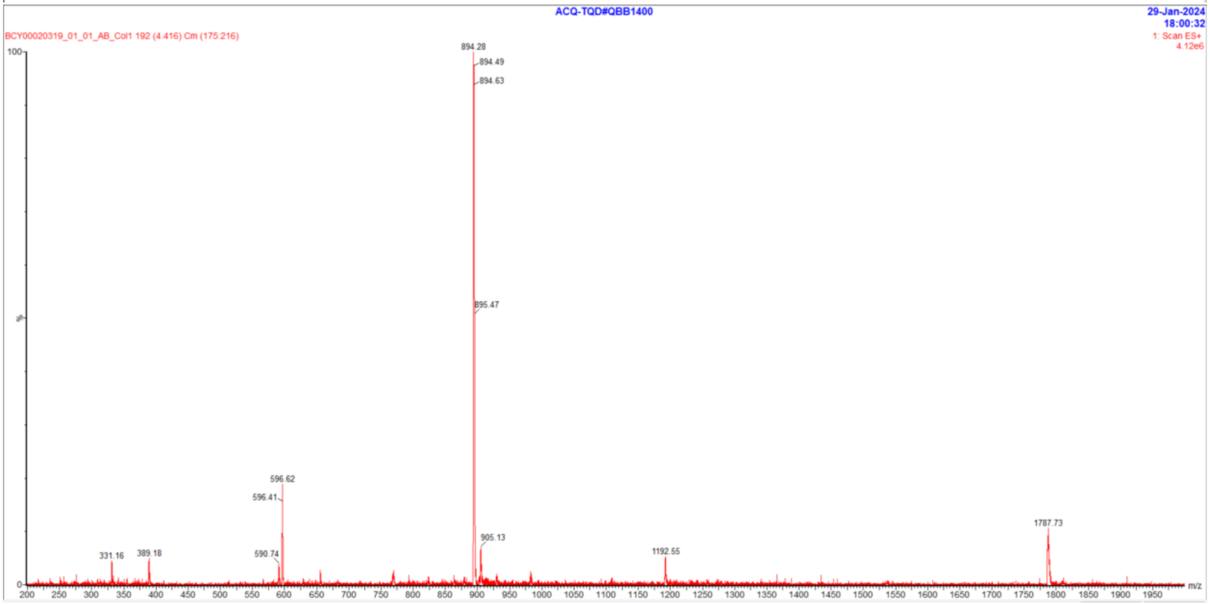

### Peptide 51

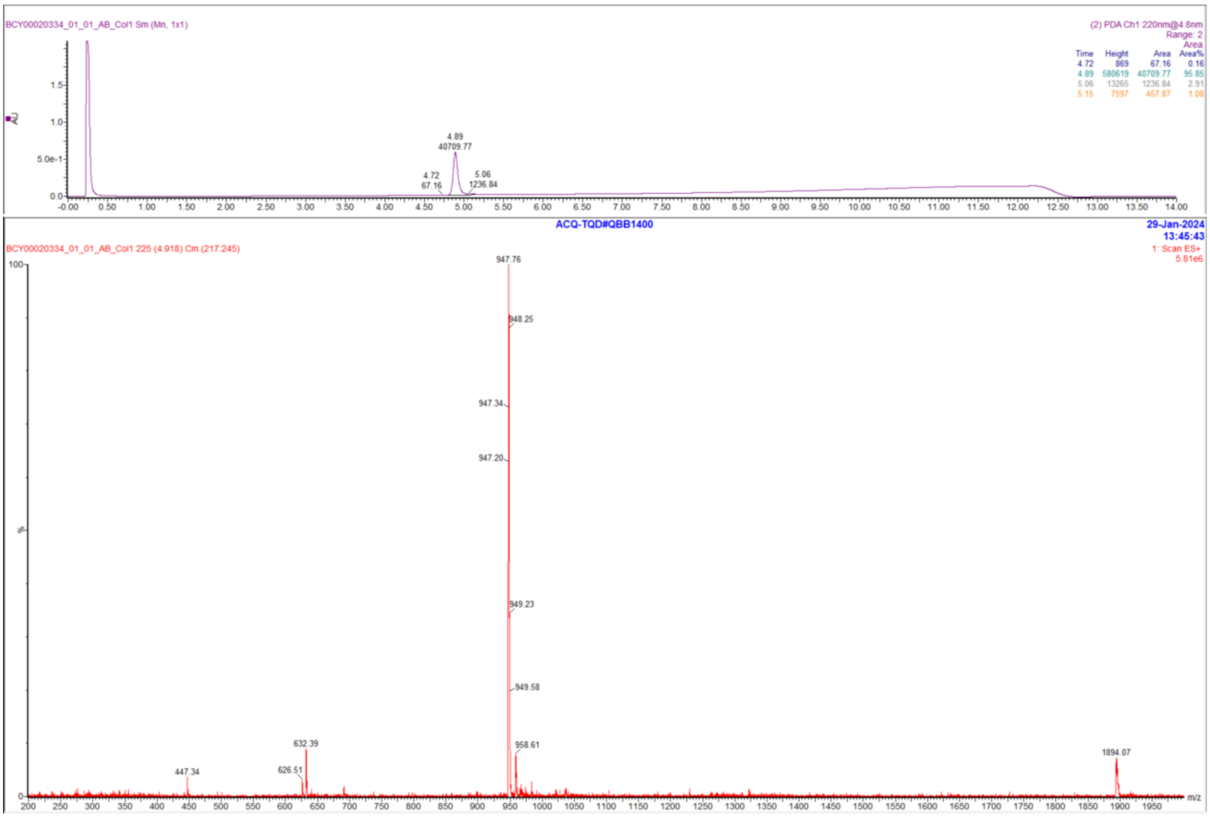

### Peptide 52

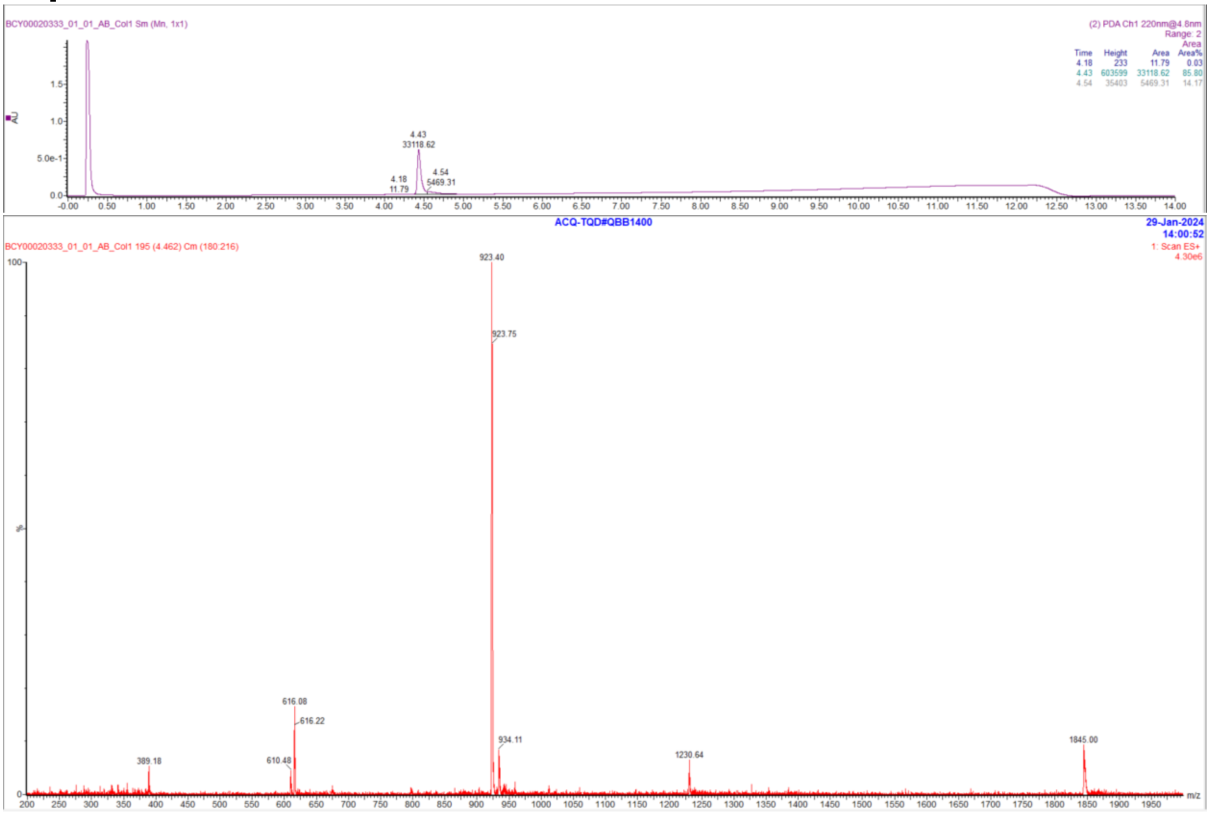

### Peptide 53

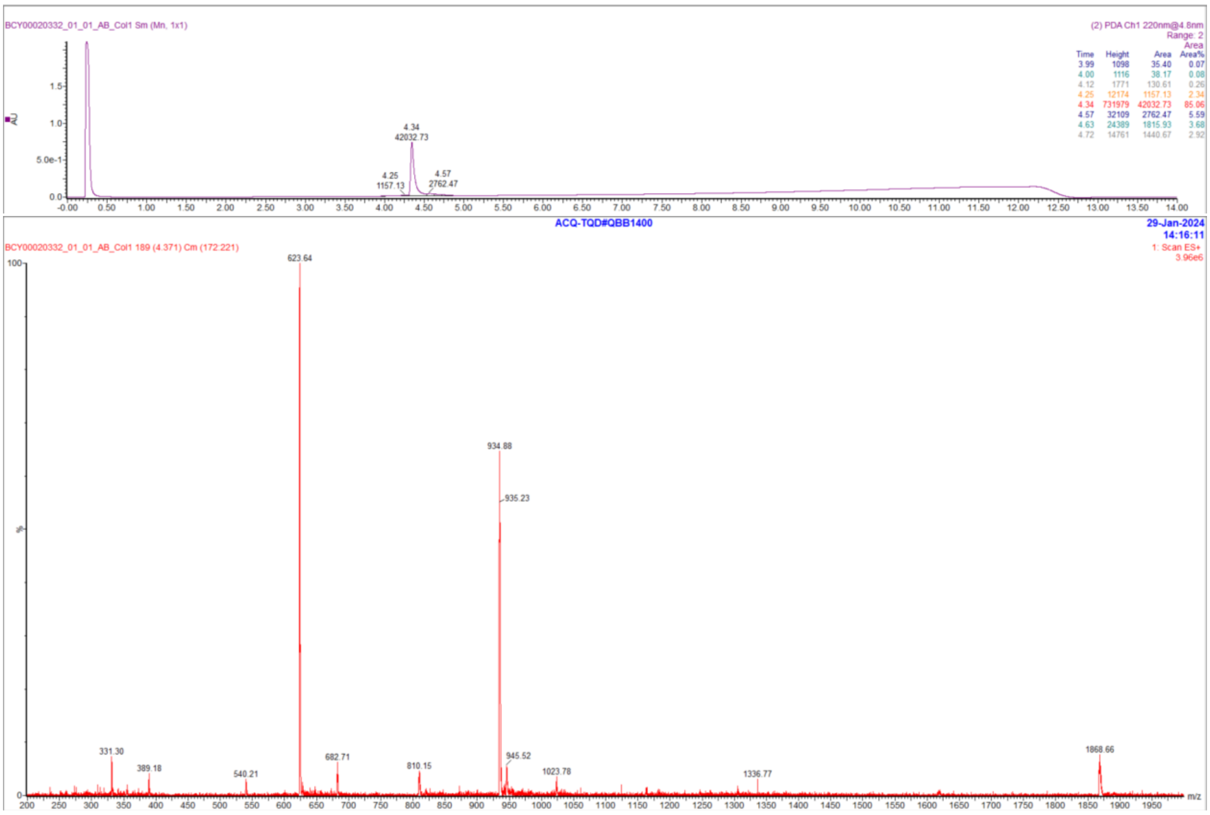

BCY00020332_01_01_AB_Col1 Sm (Mn, 1x1)
(2) PDA Ch1 220nm@4.8nm
Range: 2
Area
Time Height Area Area%
3.99 1098 35.40 0.07
4.00 1116 38.17 0.08
4.12 1771 130.61 0.26
4.25 12174 1157.13 2.34
4.34 731979 42032.73 85.06
4.57 32109 2762.47 5.59
4.63 24389 1815.93 3.68
4.72 14761 1440.67 2.92
4.34
42032.73
4.25
1157.13
4.57
2762.47
ACQ-TQD#QBB1400
29-Jan-2024
14:16:11
BCY00020332_01_01_AB_Col1 189 (4.371) Cm (172:221)
1: Scan ES+
3.96e6
623.64
934.88
935.23
331.30
389.18
540.21
682.71
810.15
945.52
1023.78
1336.77
1868.66
m/z

#### Peptide 54

**Data file:** D:\Chemstation\1\Data\Peptides_facon\7000-7999_BCY00016387_01_7564_A_zHPLC 13-03-54.D

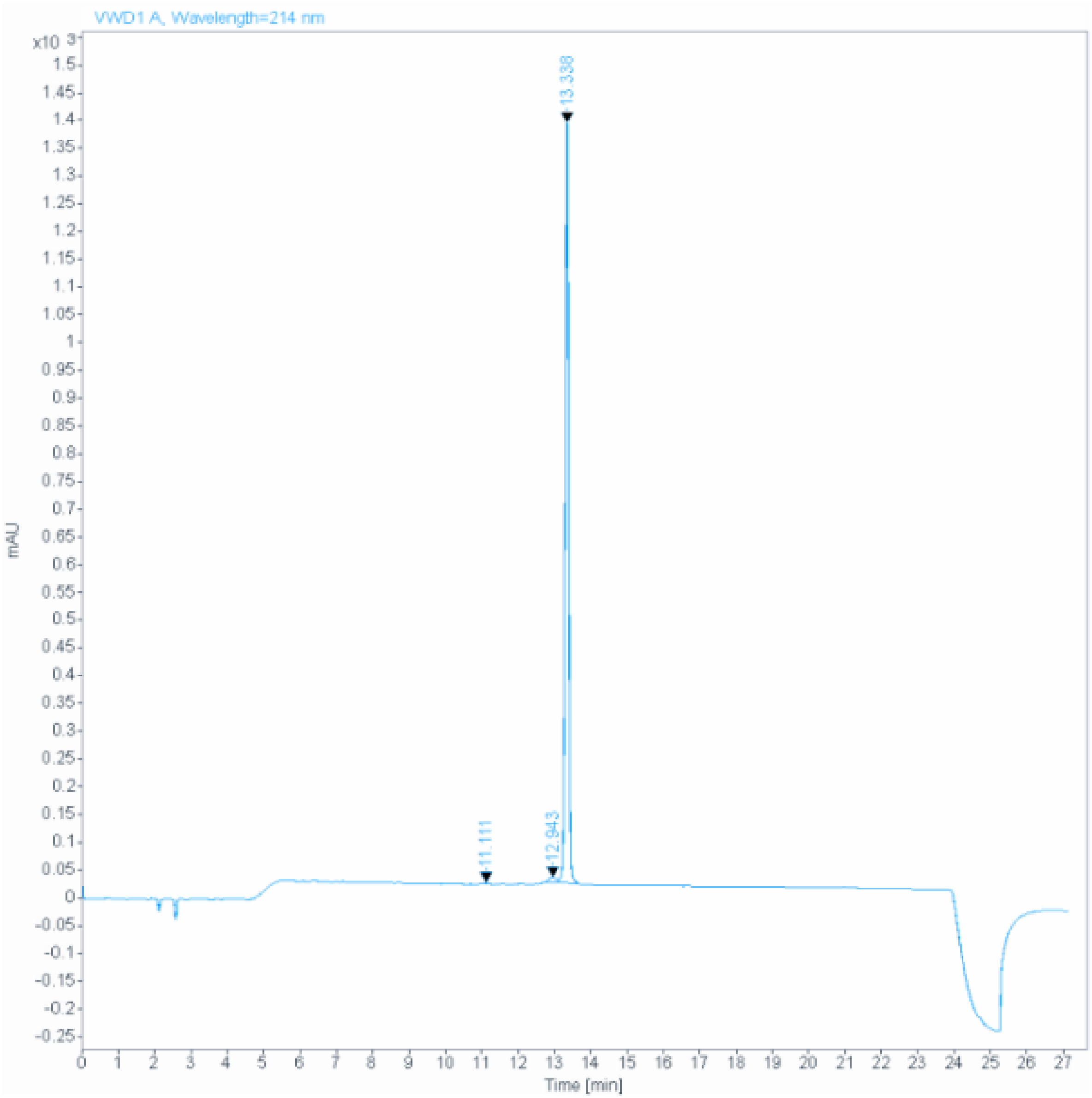

**Signal:** VWD1 A, Wavelength=214 nm

| RT [min] | Type | Width [min] | Area | Height | Area% | Name |
|---|---|---|---|---|---|---|
| 11.111 | BB | 0.0569 | 12.7126 | 3.2343 | 0.1433 | |
| 12.943 | MF | 0.2437 | 166.6561 | 11.3989 | 1.8791 | |
| 13.338 | FM | 0.1056 | 8689.7803 | 1371.4633 | 97.9776 | |
| | | Sum | 8869.1490 | | | |

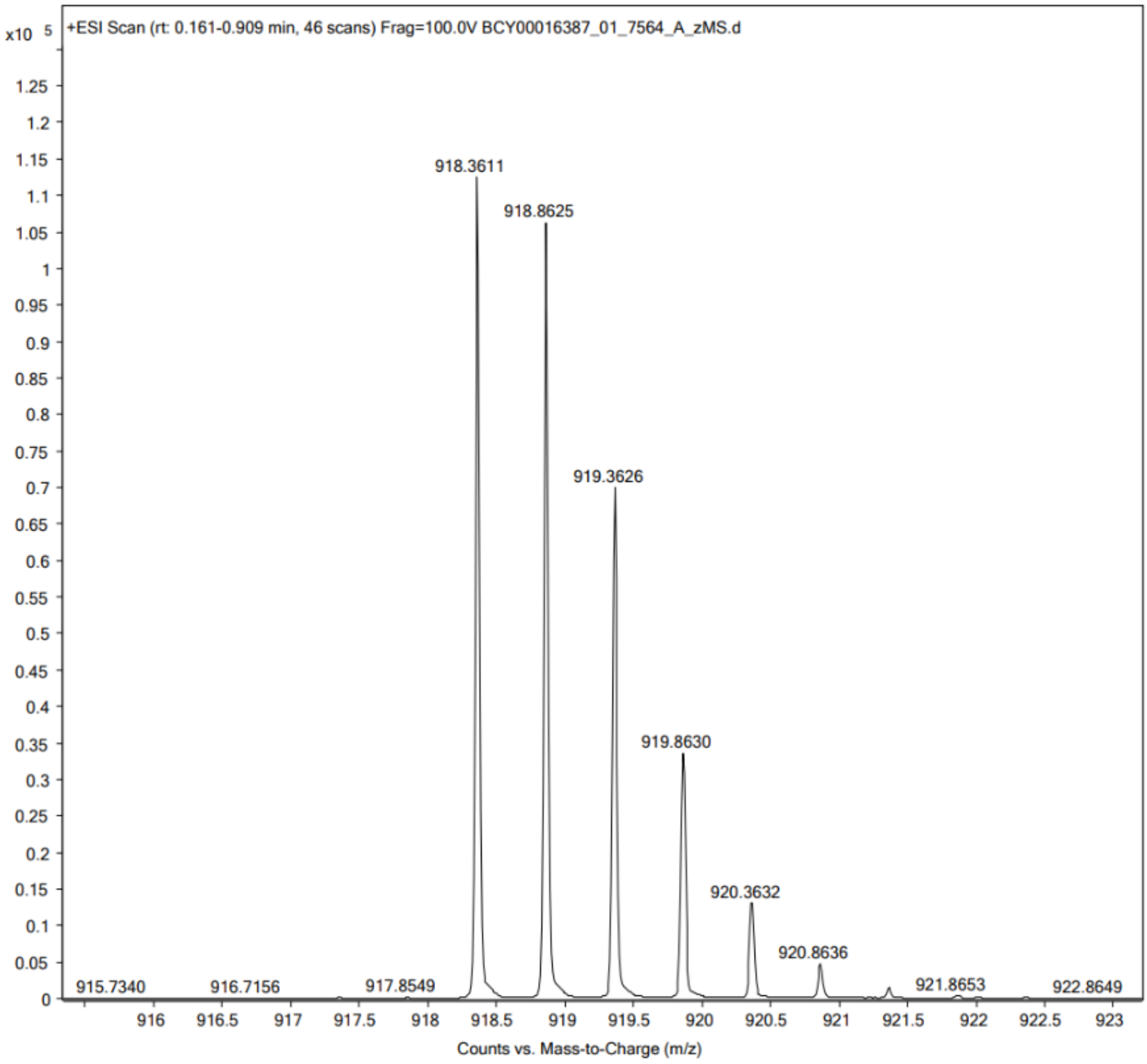

+ESI Scan (rt: 0.161-0.909 min, 46 scans) Frag=100.0V BCY00016387_01_7564_A_zMS.d
x10 5
918.3611
918.8625
919.3626
919.8630
920.3632
920.8636
915.7340
916.7156
917.8549
921.8653
922.8649
Counts vs. Mass-to-Charge (m/z)

369

#### Peptide 55

**Data file:** D:\Chemstation\1\Data\Peptides_facon\7000-7999_BCY00016385_01_7562_A_zHPLC 11-55-44.D

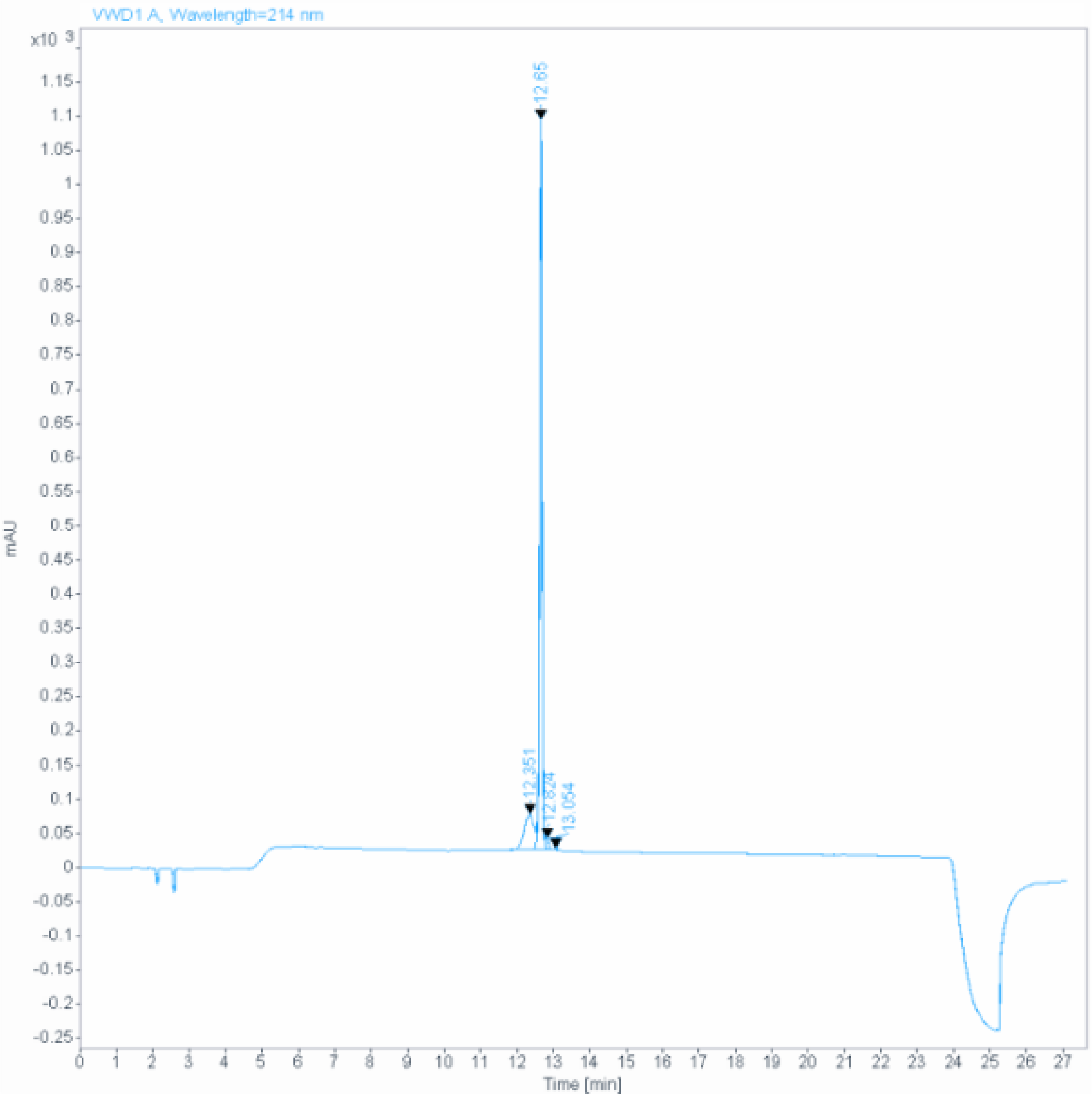

**Signal:** VWD1 A, Wavelength=214 nm

| RT [min] | Type | Width [min] | Area | Height | Area% | Name |
|---|---|---|---|---|---|---|
| 12.351 | BV E | 0.2492 | 1018.3990 | 51.9577 | 16.6186 | |
| 12.650 | VV R | 0.0716 | 5004.1411 | 1068.6205 | 81.6593 | |
| 12.824 | VB E | 0.0832 | 96.2061 | 16.3371 | 1.5699 | |
| 13.054 | MM | 0.0538 | 9.3291 | 2.8900 | 0.1522 | |
| | | Sum | 6128.0753 | | | |

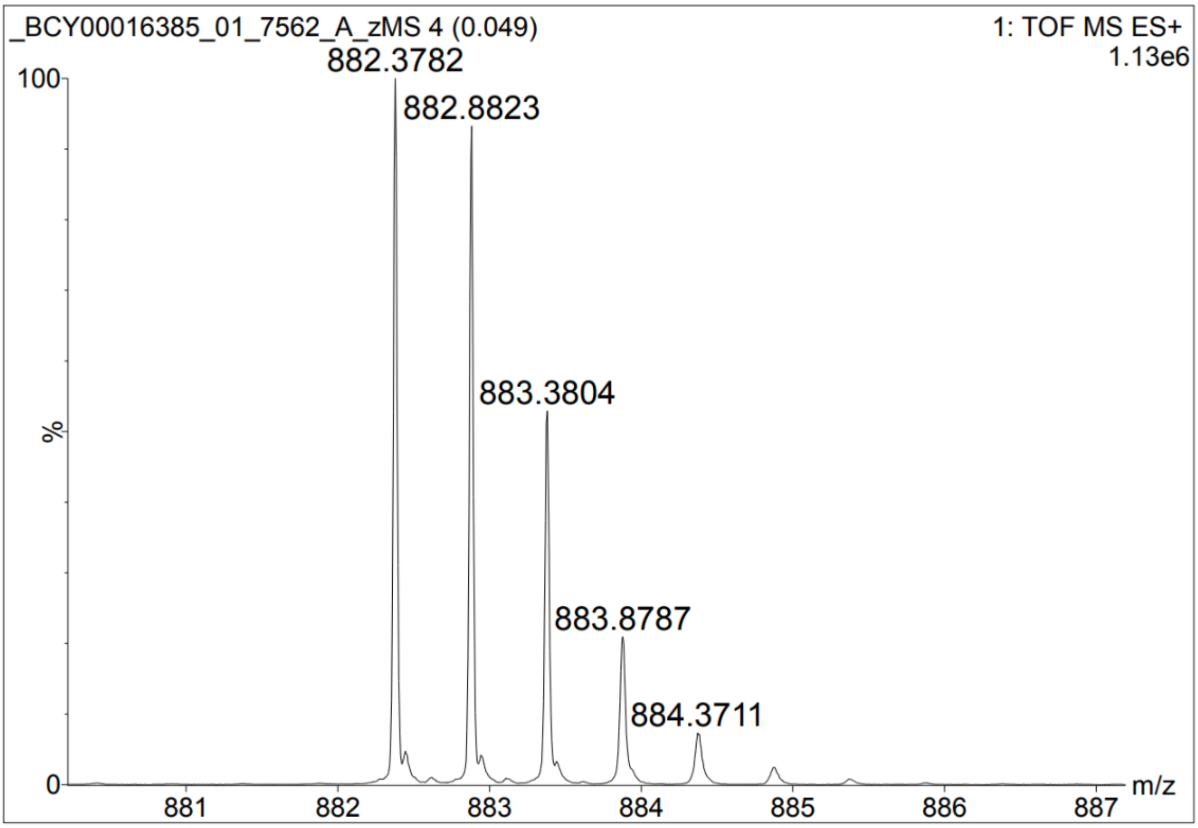

_BCY00016385_01_7562_A_zMS 4 (0.049)
1: TOF MS ES+
1.13e6
882.3782
882.8823
883.3804
883.8787
884.3711
100
%
0
881
882
883
884
885
886
887
m/z

#### Peptide 56

**Data file:** D:\Chemstation\1\Data\Peptides_facon\7000-7999_BCY00016384_01_7561_A_zHPLC 12-38-51.D

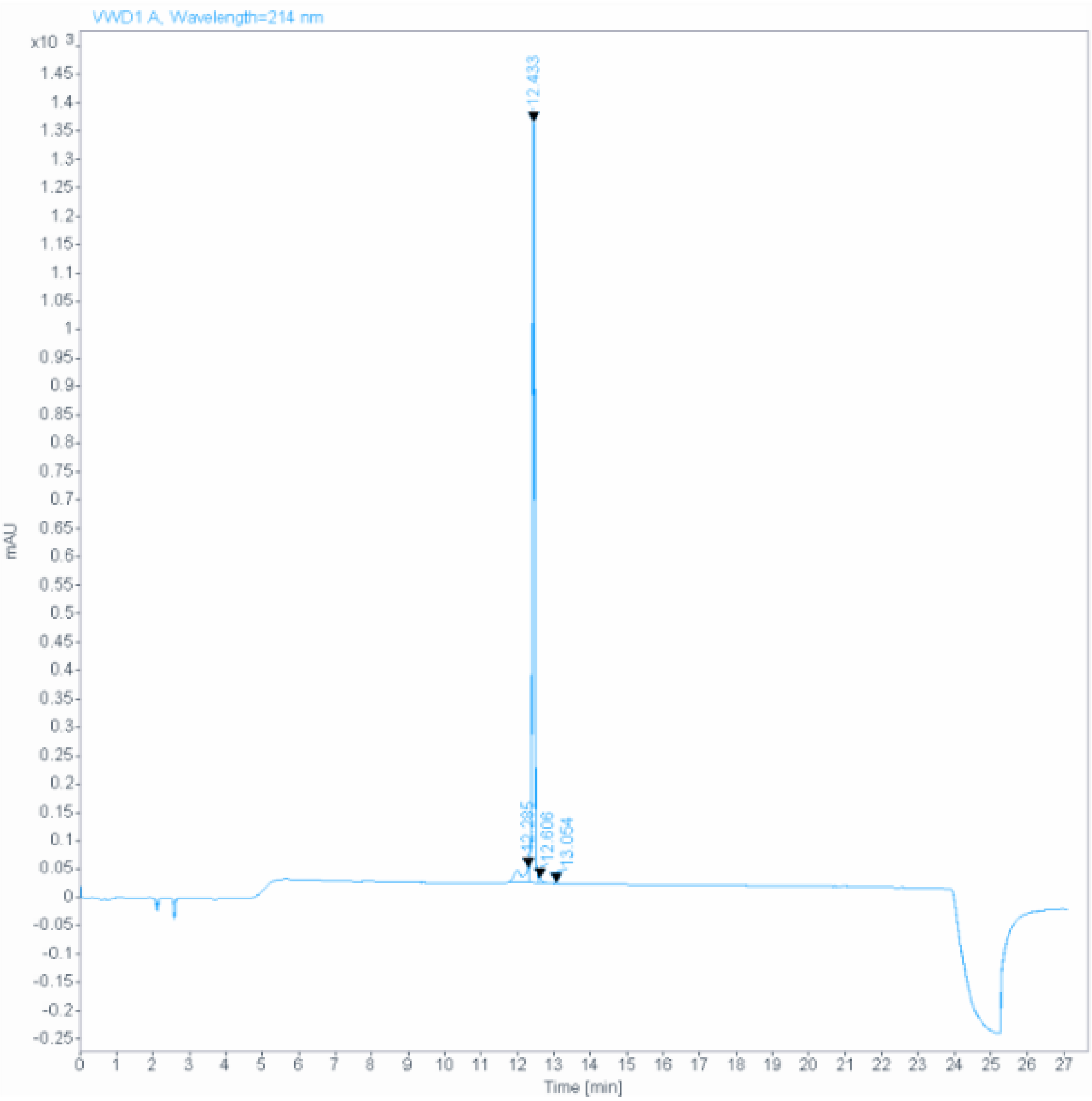

**Signal:** VWD1 A, Wavelength=214 nm

| RT [min] | Type | Width [min] | Area | Height | Area% | Name |
|---|---|---|---|---|---|---|
| 12.285 | MF | 0.3117 | 496.0348 | 26.5213 | 7.5037 | |
| 12.433 | MF | 0.0753 | 6061.4360 | 1341.2587 | 91.6936 | |
| 12.606 | FM | 0.0872 | 47.8876 | 9.1495 | 0.7244 | |
| 13.054 | MM | 0.0490 | 5.1769 | 1.7622 | 0.0783 | |
| | | Sum | 6610.5353 | | | |

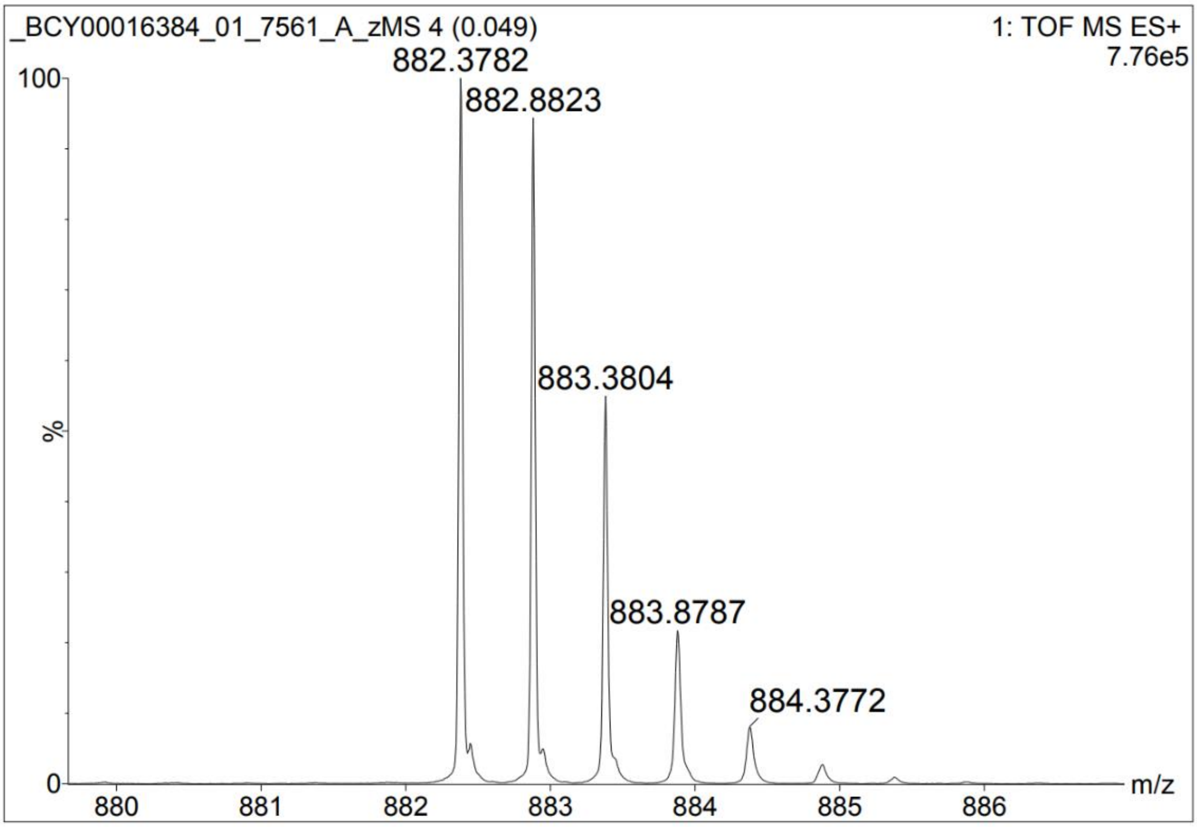
_BCY00016384_01_7561_A_zMS 4 (0.049)
1: TOF MS ES+
7.76e5
882.3782
882.8823
883.3804
883.8787
884.3772
100
%
0
880
881
882
883
884
885
886
m/z

#### Peptide 57

**Data file:** D:\Chemstation\1\Data\Peptides_facon\7000-7999 _BCY00016381_01_7558_A_zHPLC 11-35-52.D

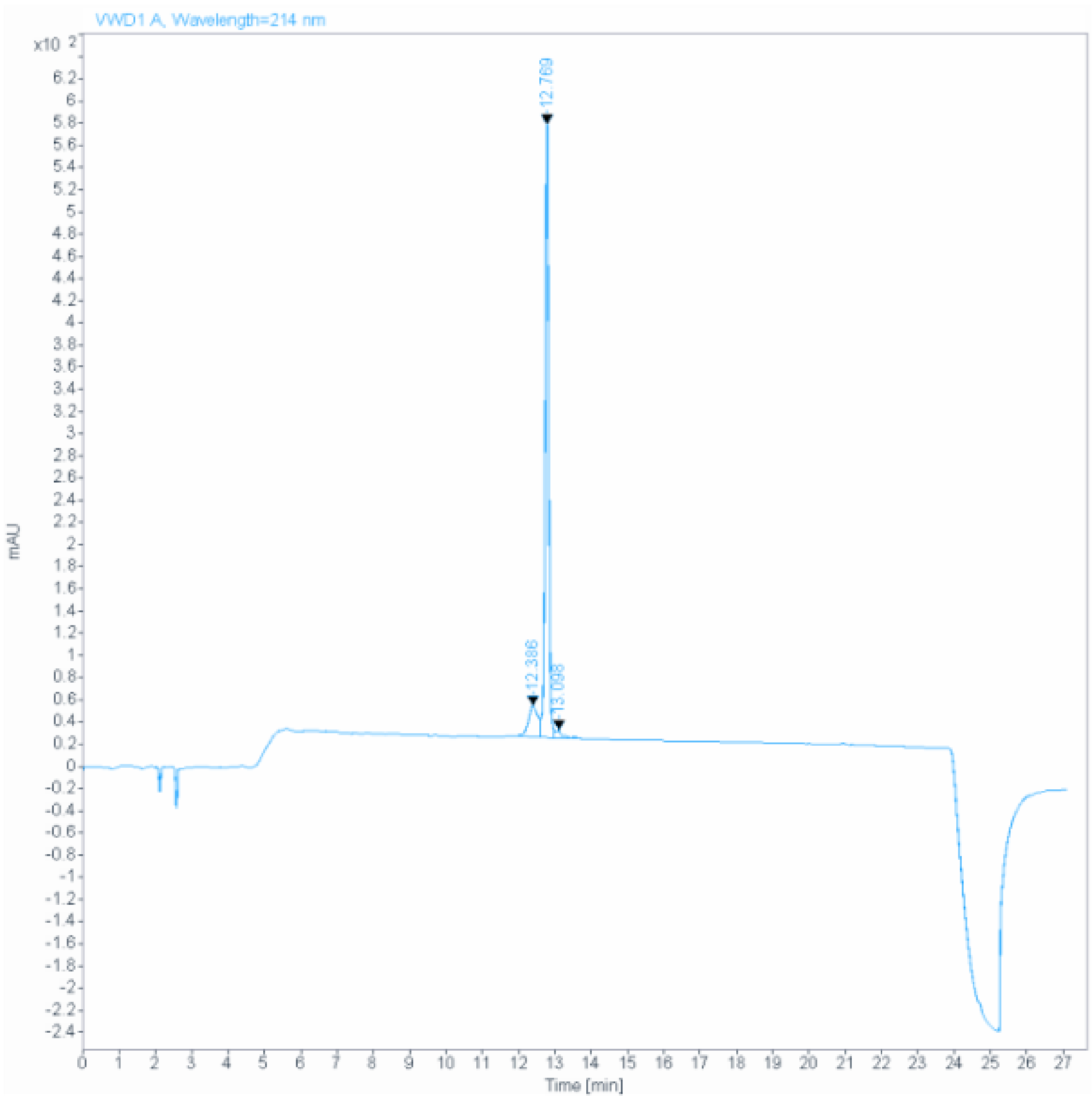

**Signal:** VWD1 A, Wavelength=214 nm

| RT [min] | Type | Width [min] | Area | Height | Area% | Name |
|---|---|---|---|---|---|---|
| 12.386 | MF | 0.3190 | 547.3634 | 28.5988 | 12.5099 | |
| 12.769 | FM | 0.1119 | 3717.5630 | 553.5273 | 84.9645 | |
| 13.098 | FM | 0.2575 | 110.5052 | 7.1529 | 2.5256 | |
| | | Sum | 4375.4316 | | | |

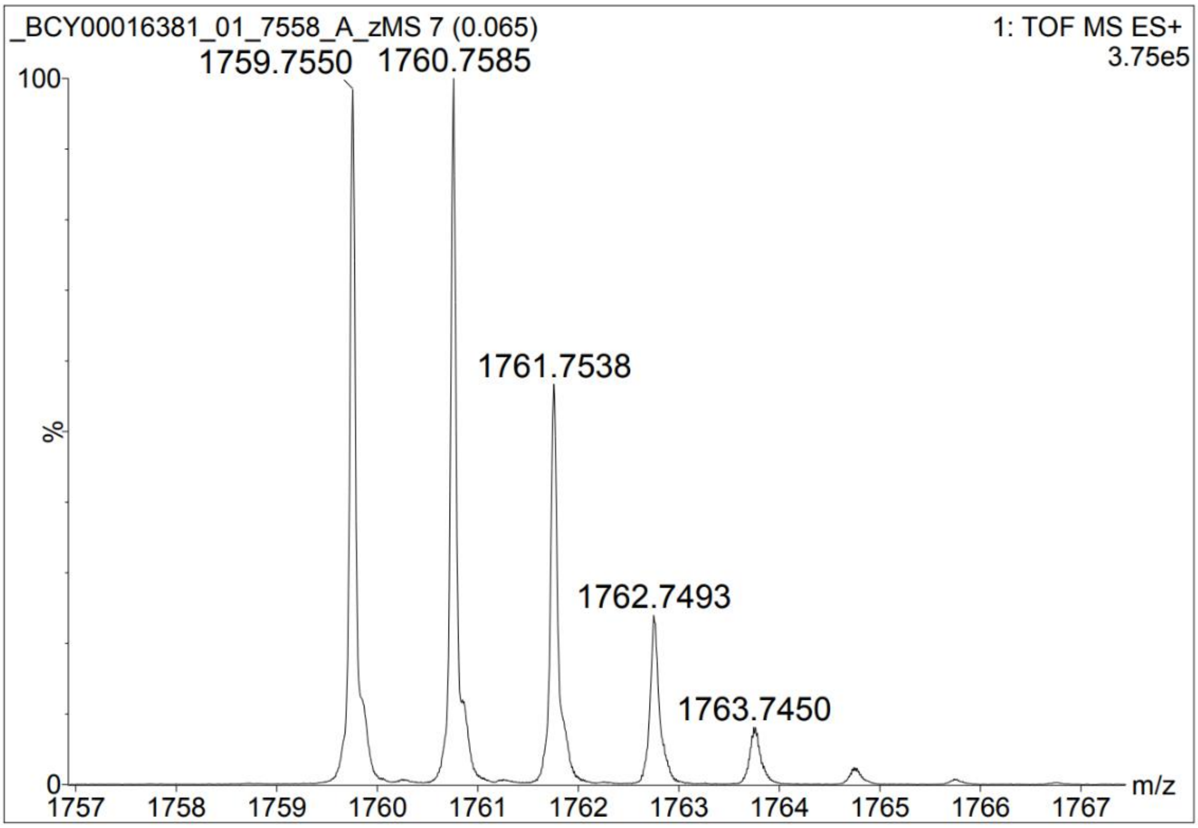
_BCY00016381_01_7558_A_zMS 7 (0.065)
1: TOF MS ES+
3.75e5
1759.7550
1760.7585
1761.7538
1762.7493
1763.7450
100
%
0
m/z

#### Peptide 58

**Data file:** D:\Chemstation\1\Data\Peptides_facon\7000-7999_BCY00016383_01_7560_A_zHPLC 12-14-55.D

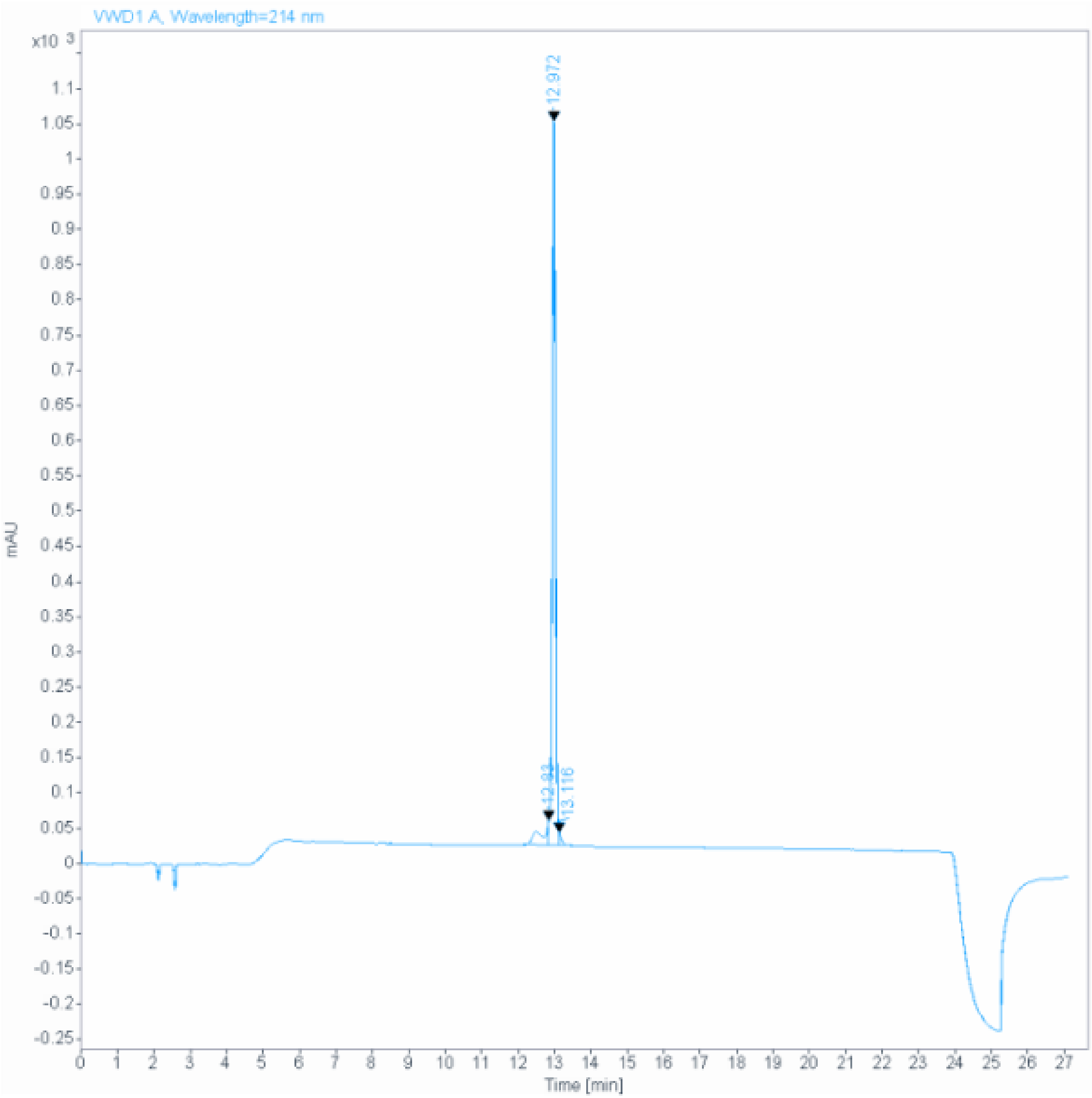

**Signal:** VWD1 A, Wavelength=214 nm

| RT [min] | Type | Width [min] | Area | Height | Area% | Name |
|---|---|---|---|---|---|---|
| 12.830 | MF | 0.2309 | 484.2571 | 34.9524 | 7.3136 | |
| 12.972 | MF | 0.0982 | 6056.5078 | 1028.3544 | 91.4700 | |
| 13.116 | FM | 0.0731 | 80.5382 | 18.3675 | 1.2163 | |
| | | Sum | 6621.3031 | | | |

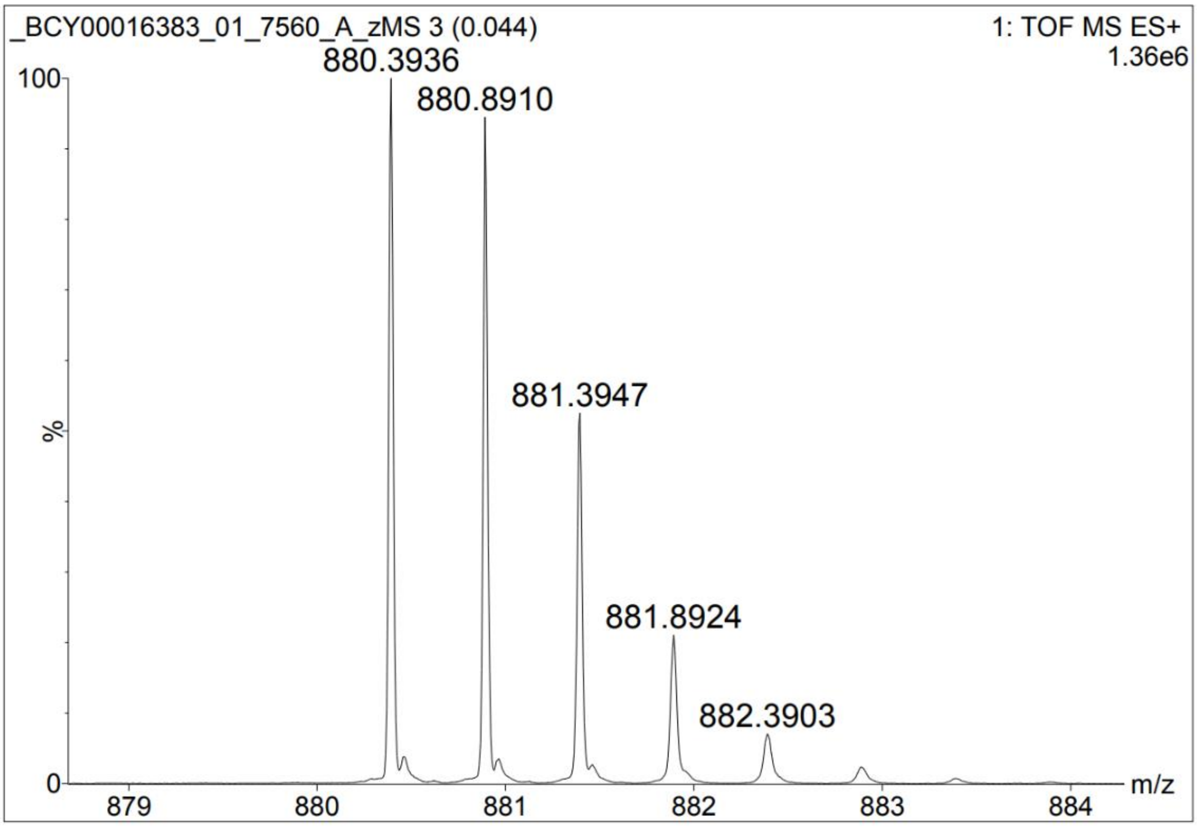
_BCY00016383_01_7560_A_zMS 3 (0.044)
1: TOF MS ES+
1.36e6
880.3936
880.8910
881.3947
881.8924
882.3903
100
%
0
879
880
881
882
883
884
m/z

#### Peptide 59

**Data file:** D:\Chemstation\1\Data\Peptides_facon\7000-7999_BCY00016392_01_7569_A_zHPLC 12-32-22.D

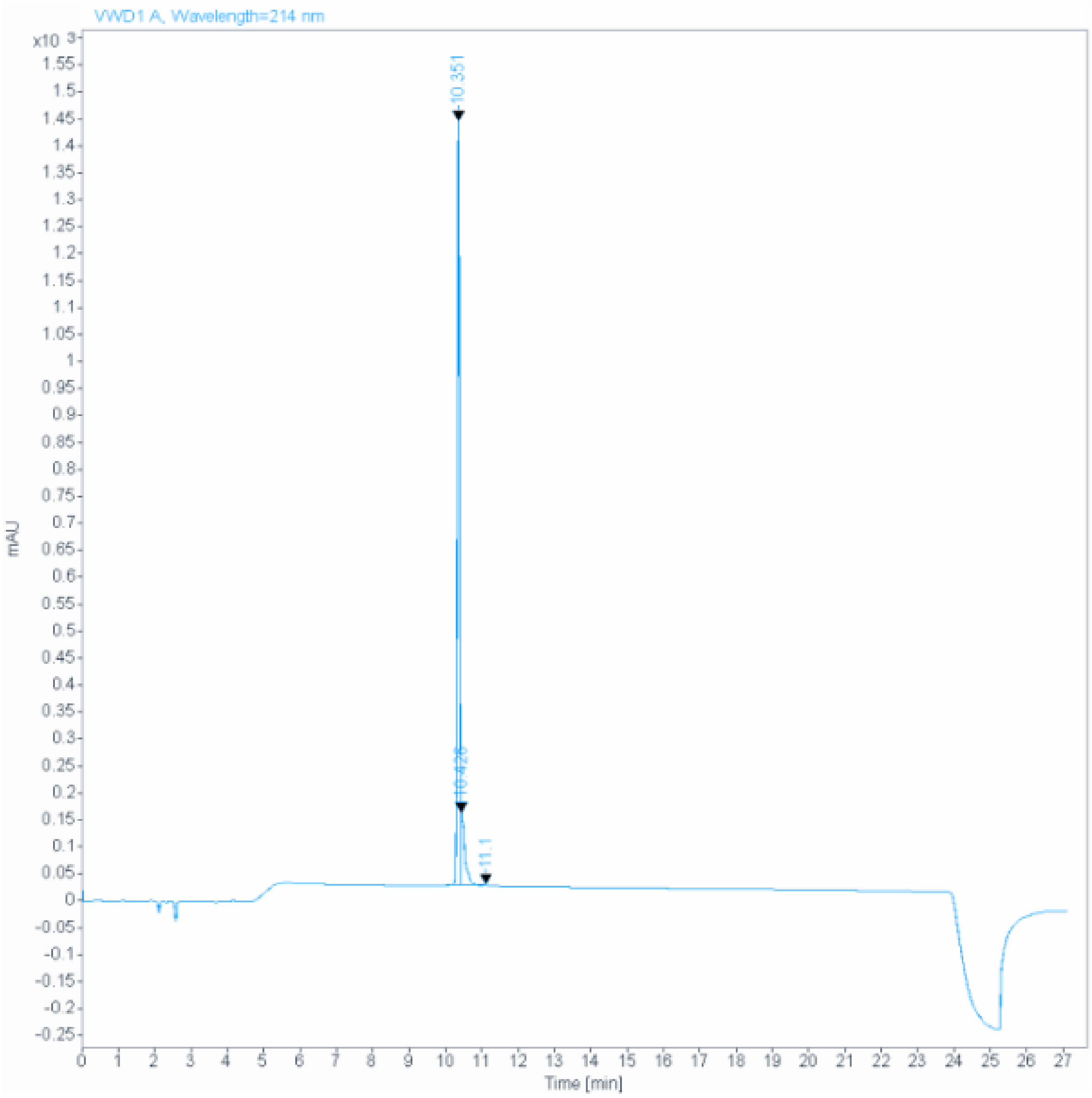

**Signal:** VWD1 A, Wavelength=214 nm

| RT [min] | Type | Width [min] | Area | Height | Area% | Name |
|---|---|---|---|---|---|---|
| 10.351 | MF | 0.0660 | 5622.1333 | 1419.0532 | 85.9266 | |
| 10.426 | FM | 0.1119 | 906.7505 | 135.0147 | 13.8584 | |
| 11.100 | BB | 0.0593 | 14.0652 | 3.5067 | 0.2150 | |
| | | Sum | 6542.9490 | | | |

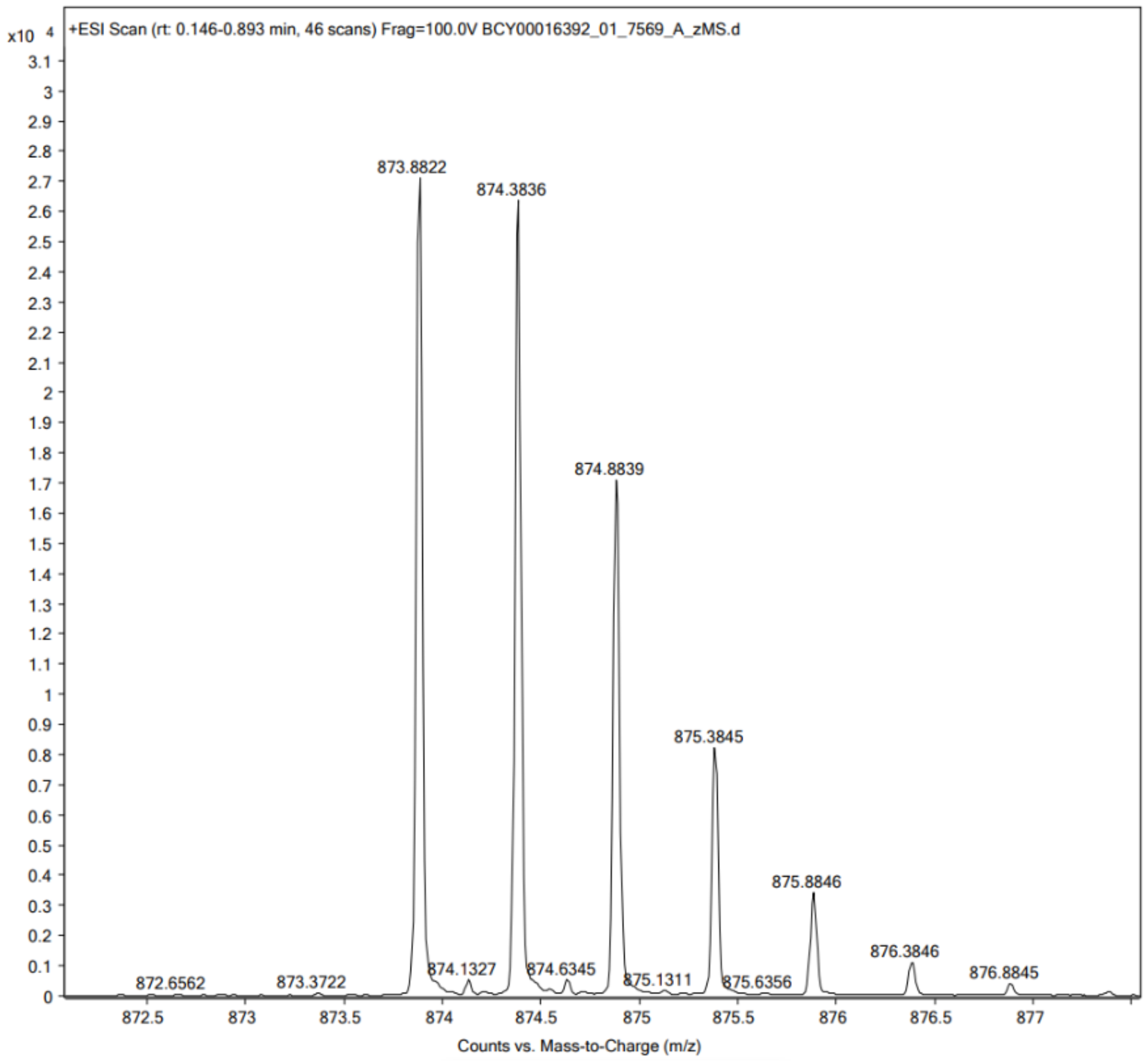

x10 4
+ESI Scan (rt: 0.146-0.893 min, 46 scans) Frag=100.0V BCY00016392_01_7569_A_zMS.d
873.8822
874.3836
874.8839
875.3845
875.8846
876.3846
876.8845
872.6562
873.3722
874.1327
874.6345
875.1311
875.6356
Counts vs. Mass-to-Charge (m/z)

#### Peptide 60

**Data file:** D:\Chemstation\1\Data\Peptides_facon\7000-7999_BCY00016390_01_7567_A_zHPLC 17-49-38.D

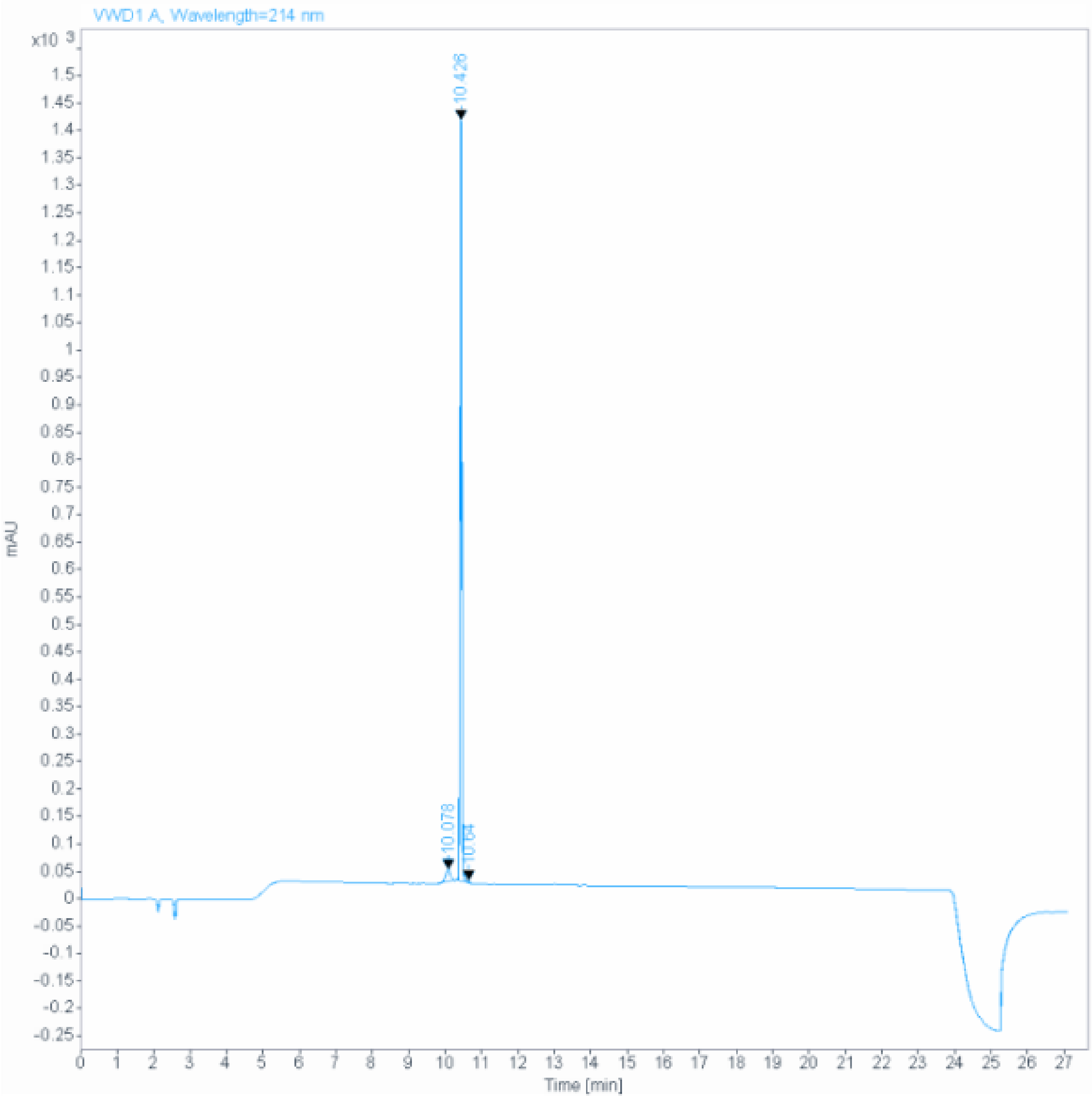

**Signal:** VWD1 A, Wavelength=214 nm

| RT [min] | Type | Width [min] | Area | Height | Area% | Name |
|---|---|---|---|---|---|---|
| 10.078 | BB | 0.1423 | 228.6930 | 23.2463 | 4.3871 | |
| 10.426 | BV R | 0.0558 | 4966.4780 | 1386.9453 | 95.2733 | |
| 10.640 | VB E | 0.0492 | 17.7032 | 5.5542 | 0.3396 | |
| | | Sum | 5212.8743 | | | |

388 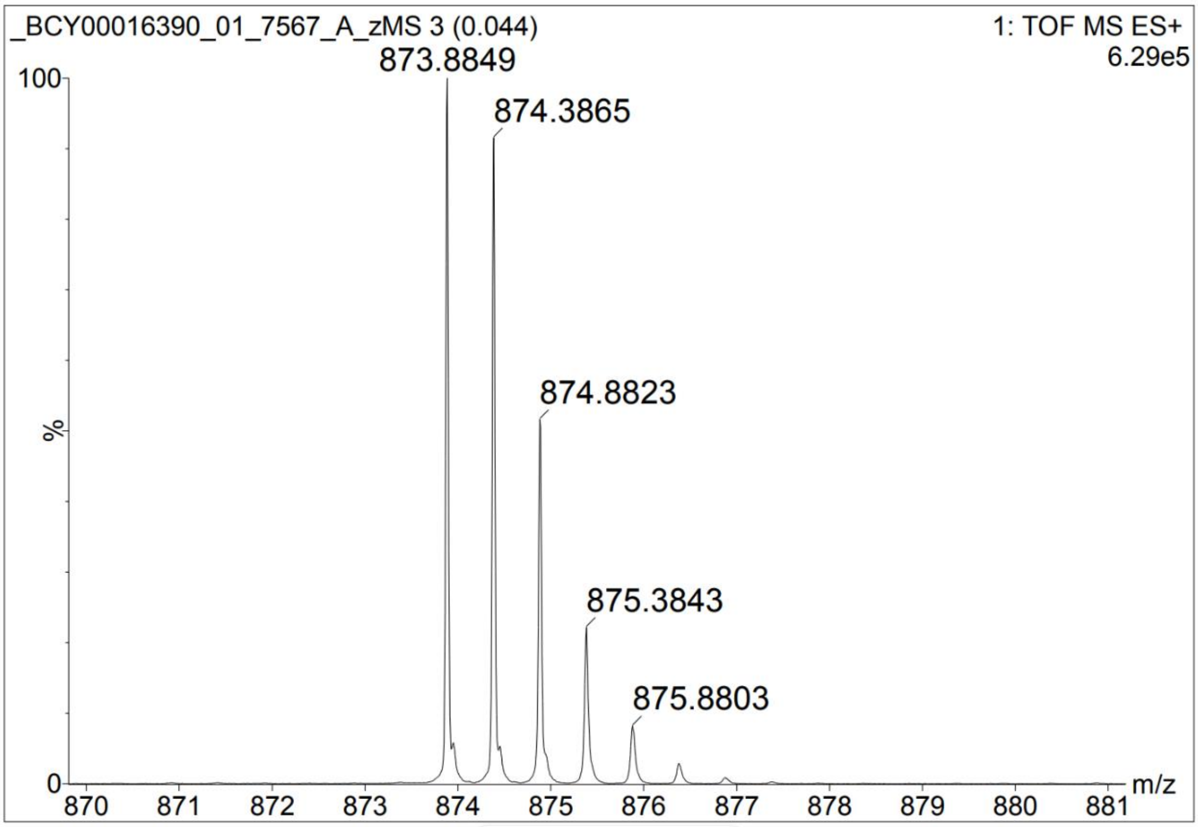

_BCY00016390_01_7567_A_zMS 3 (0.044)
1: TOF MS ES+
6.29e5
873.8849
874.3865
874.8823
875.3843
875.8803
100
%
0
m/z

389 **Peptide 61**

**Data file:** D:\Chemstation\1\Data\Peptides_facon\7000-7999_BCY00016420_01_7572_A_zHPLC 15-26-48.D

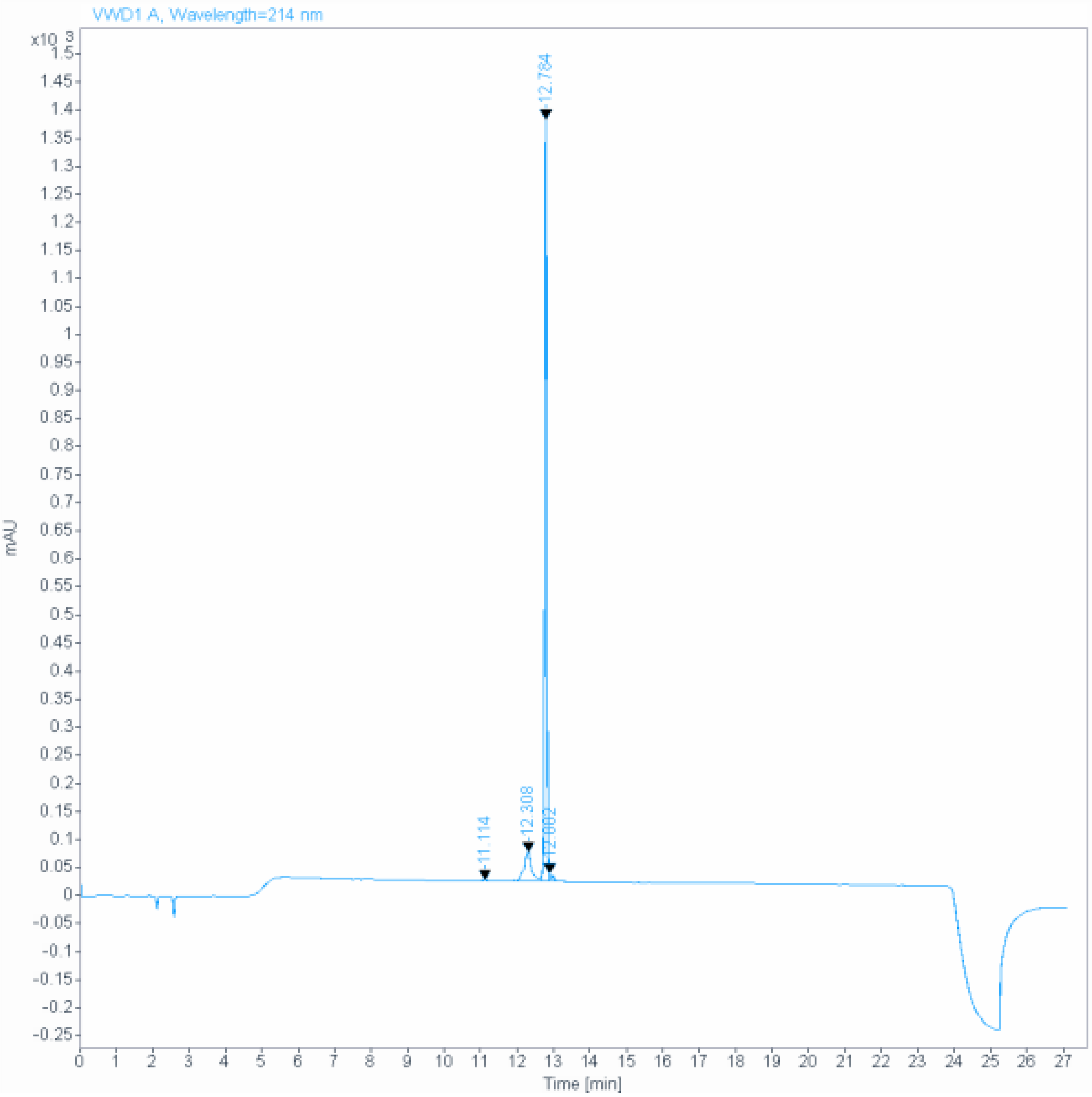

**Signal:** VWD1 A, Wavelength=214 nm

| RT [min] | Type | Width [min] | Area | Height | Area% | Name |
|---|---|---|---|---|---|---|
| 11.114 | MM | 0.0529 | 4.7925 | 1.5107 | 0.0821 | |
| 12.308 | MF | 0.2497 | 769.0922 | 51.3441 | 13.1703 | |
| 12.784 | MF | 0.0612 | 4988.9927 | 1358.3730 | 85.4340 | |
| 12.882 | FM | 0.0772 | 76.7119 | 14.2774 | 1.3137 | |
| | | Sum | 5839.5893 | | | |

390

391

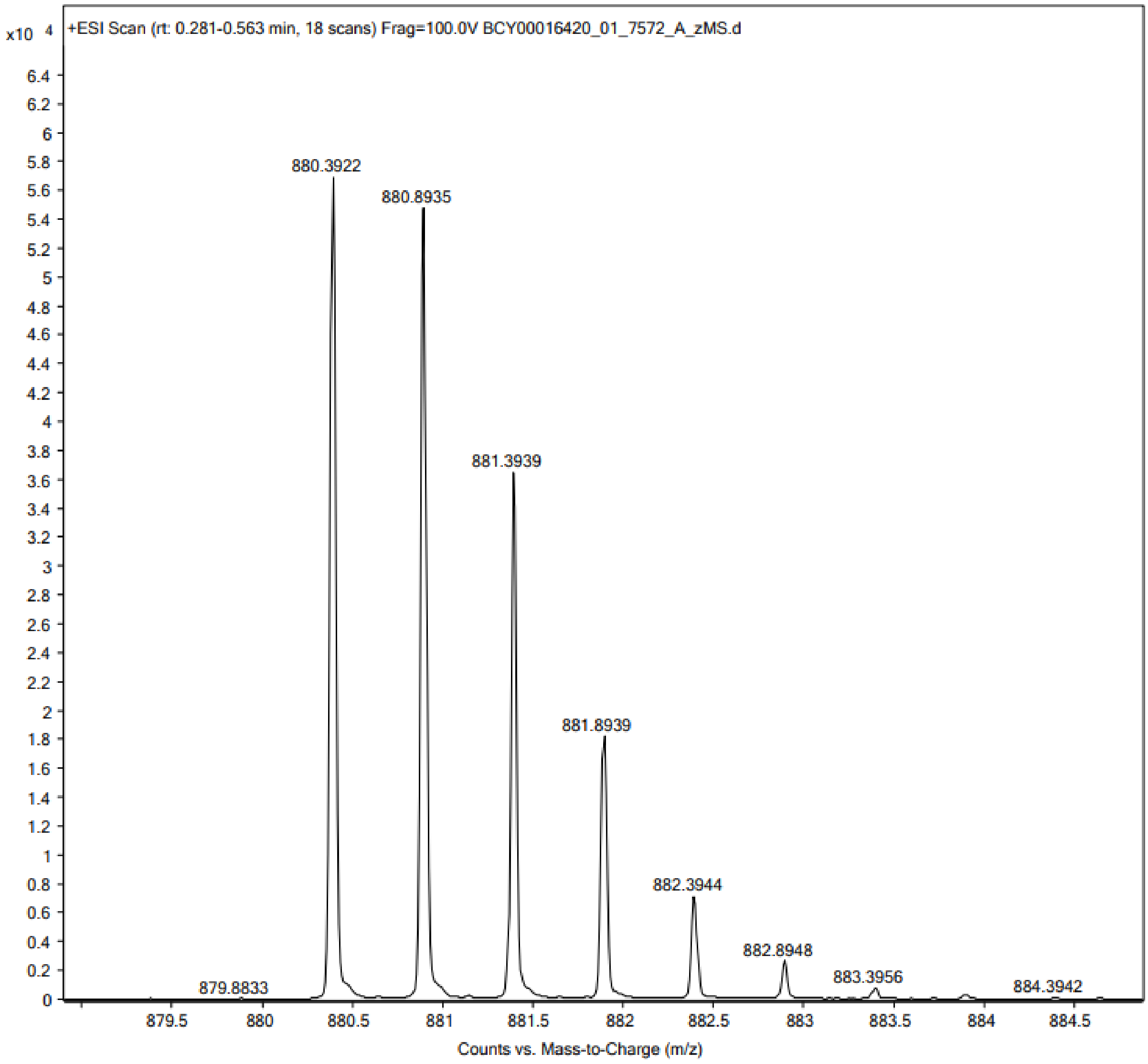

x10 4
+ESI Scan (rt: 0.281-0.563 min, 18 scans) Frag=100.0V BCY00016420_01_7572_A_zMS.d
880.3922
880.8935
881.3939
881.8939
882.3944
882.8948
883.3956
884.3942
879.8833
6.4
6.2
6
5.8
5.6
5.4
5.2
5
4.8
4.6
4.4
4.2
4
3.8
3.6
3.4
3.2
3
2.8
2.6
2.4
2.2
2
1.8
1.6
1.4
1.2
1
0.8
0.6
0.4
0.2
0
879.5
880
880.5
881
881.5
882
882.5
883
883.5
884
884.5
Counts vs. Mass-to-Charge (m/z)

392 **Peptide 62**

**Data file:** D:\Chemstation\1\Data\Peptides_facon\7000-7999_BCY00016414_01_7599_A_zHPLC 17-10-12.D

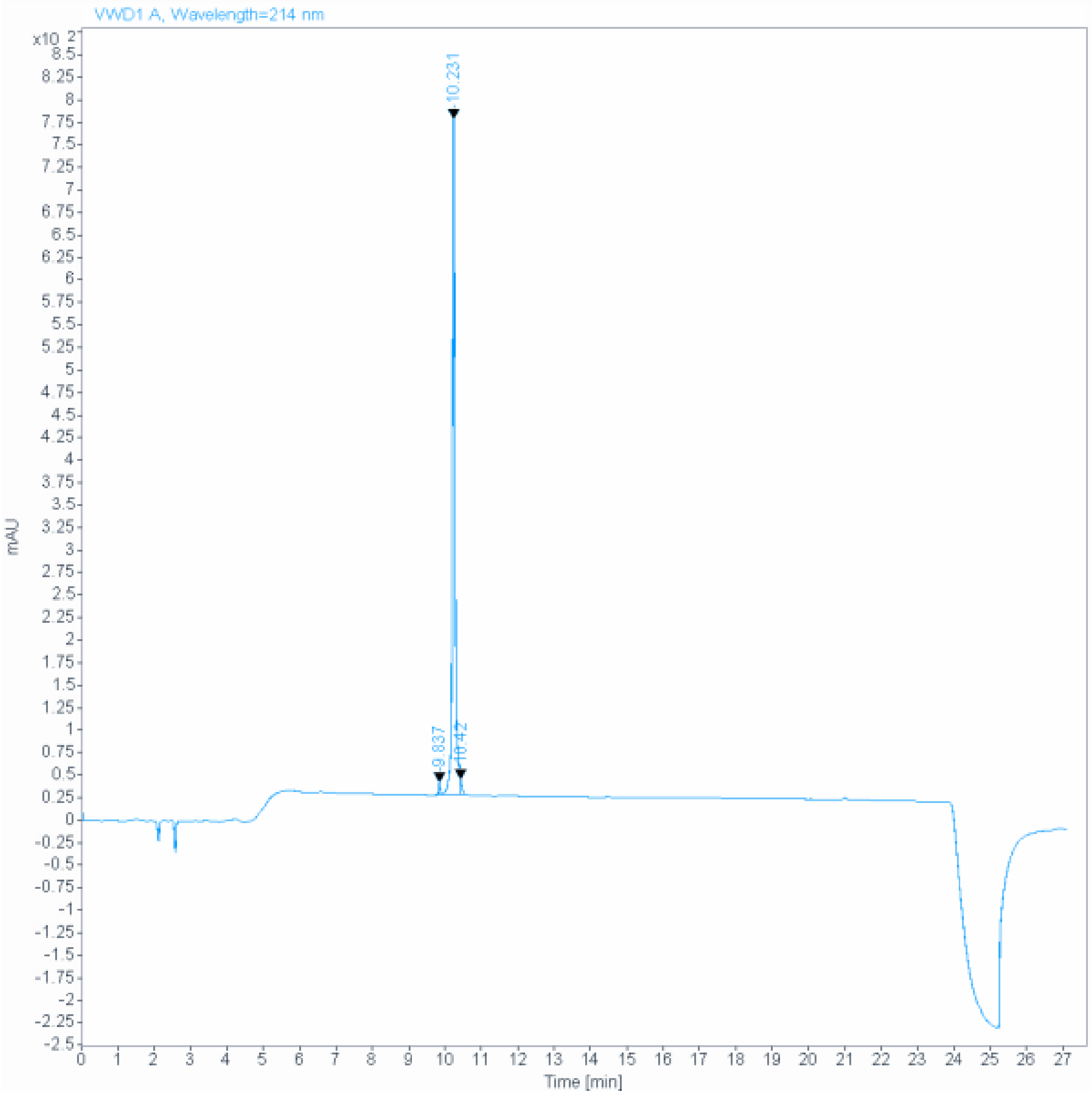

**Signal:** VWD1 A, Wavelength=214 nm

| RT [min] | Type | Width [min] | Area | Height | Area% | Name |
|---|---|---|---|---|---|---|
| 9.837 | MF | 0.0500 | 43.0186 | 14.3529 | 1.0656 | |
| 10.231 | MF | 0.0873 | 3936.6382 | 751.3096 | 97.5109 | |
| 10.420 | FM | 0.0543 | 57.4695 | 17.6266 | 1.4235 | |
| | | Sum | 4037.1262 | | | |

393

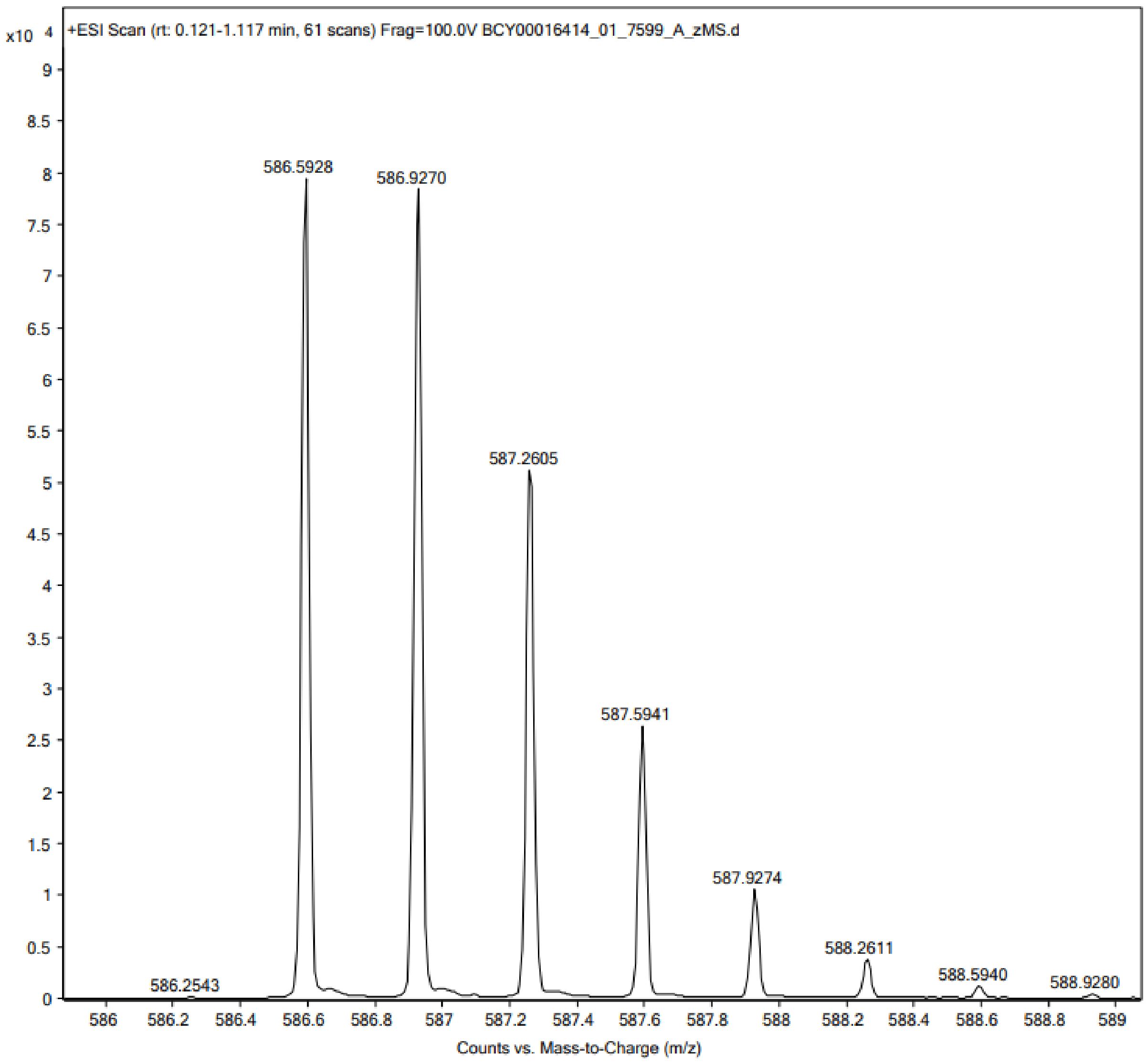
x10 4
+ESI Scan (rt: 0.121-1.117 min, 61 scans) Frag=100.0V BCY00016414_01_7599_A_zMS.d
586.5928
586.9270
587.2605
587.5941
587.9274
588.2611
588.5940
588.9280
586.2543
Counts vs. Mass-to-Charge (m/z)

394

#### Peptide 63

**Data file:** D:\Chemstation\1\Data\Peptides_facon\7000-7999 _BCY00016424_01_7590_A_zHPLC 12-15-21.D

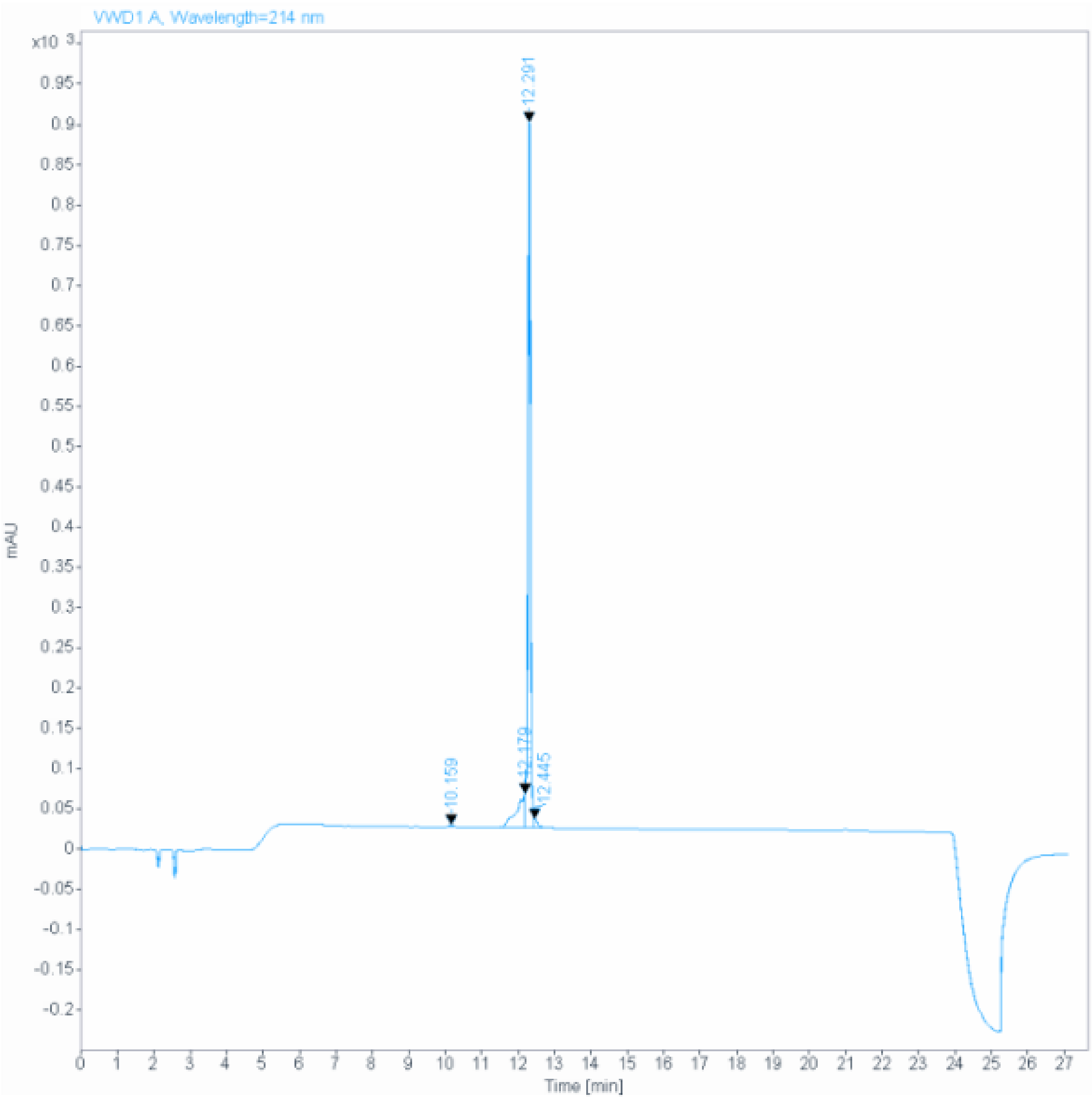

**Signal:** VWD1 A, Wavelength=214 nm

| RT [min] | Type | Width [min] | Area | Height | Area% | Name |
|---|---|---|---|---|---|---|
| 10.159 | MM | 0.1100 | 18.7838 | 2.8456 | 0.3526 | |
| 12.179 | MF | 0.2596 | 666.1787 | 42.7704 | 12.5038 | |
| 12.291 | MF | 0.0866 | 4556.5186 | 876.4865 | 85.5230 | |
| 12.445 | FM | 0.1265 | 86.3453 | 11.3775 | 1.6206 | |
| | | Sum | 5327.8263 | | | |

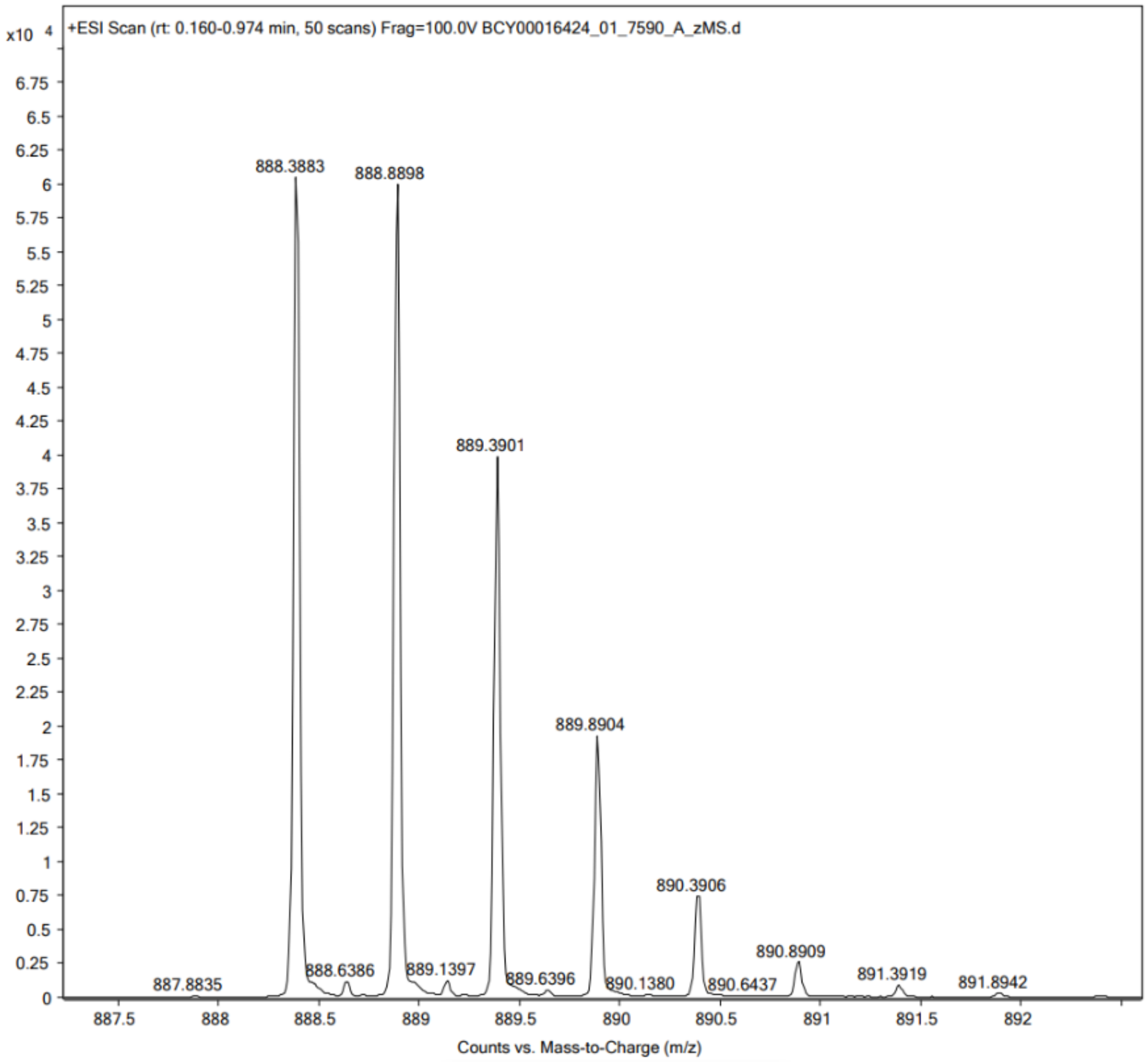
x10 4
+ESI Scan (rt: 0.160-0.974 min, 50 scans) Frag=100.0V BCY00016424_01_7590_A_zMS.d
888.3883
888.8898
889.3901
889.8904
890.3906
890.8909
891.3919
891.8942
887.8835
888.6386
889.1397
889.6396
890.1380
890.6437
Counts vs. Mass-to-Charge (m/z)

#### Peptide 64

**Data file:** D:\Chemstation\1\Data\Peptides_facon\7000-7999_BCY00016425_01_7591_A_zHPLC 12-51-13.D

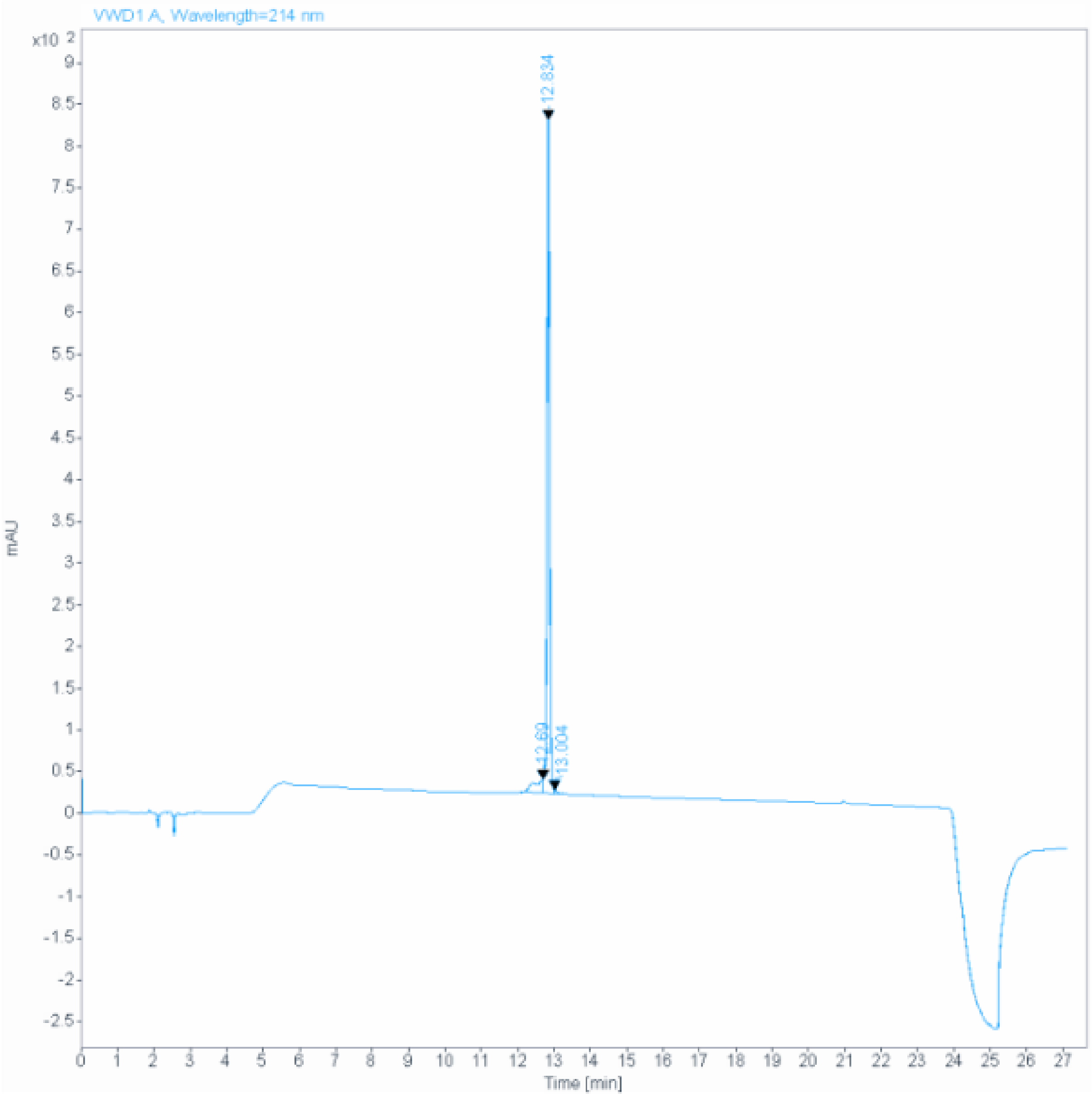

**Signal:** VWD1 A, Wavelength=214 nm

| RT [min] | Type | Width [min] | Area | Height | Area% | Name |
|---|---|---|---|---|---|---|
| 12.690 | MF | 0.3195 | 313.4438 | 16.3531 | 7.0477 | |
| 12.834 | MF | 0.0848 | 4111.5620 | 807.7789 | 92.4476 | |
| 13.004 | FM | 0.0878 | 22.4476 | 4.2594 | 0.5047 | |
| | | Sum | 4447.4534 | | | |

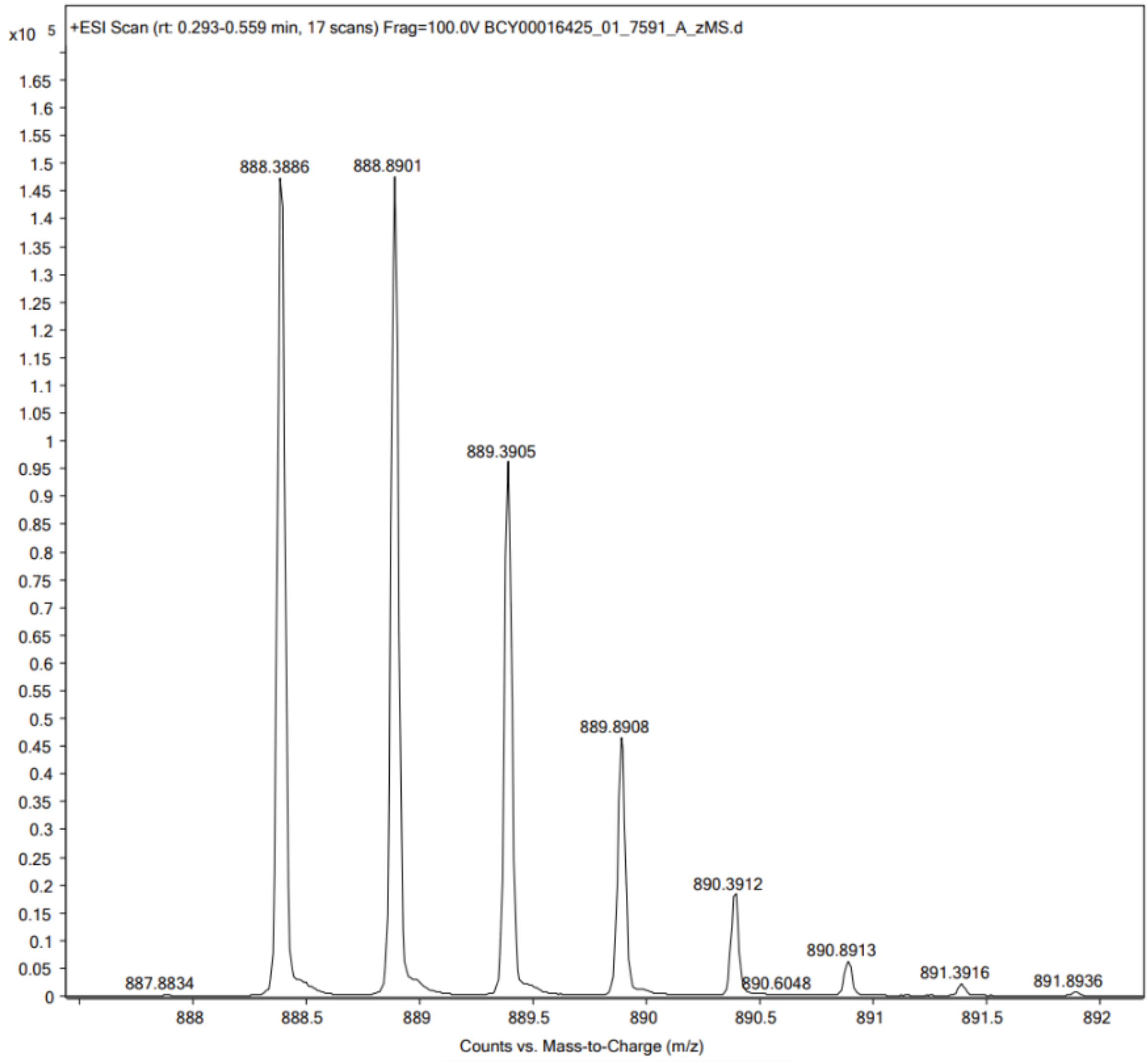
x10 5
+ESI Scan (rt: 0.293-0.559 min, 17 scans) Frag=100.0V BCY00016425_01_7591_A_zMS.d
888.3886
888.8901
889.3905
889.8908
890.3912
890.8913
891.3916
891.8936
887.8834
890.6048
Counts vs. Mass-to-Charge (m/z)

400

#### Peptide 65

**Data file:** D:\Chemstation\1\Data\Peptides_facon\7000-7999_BCY00016423_01_7585_A_zHPLC 15-52-38.D

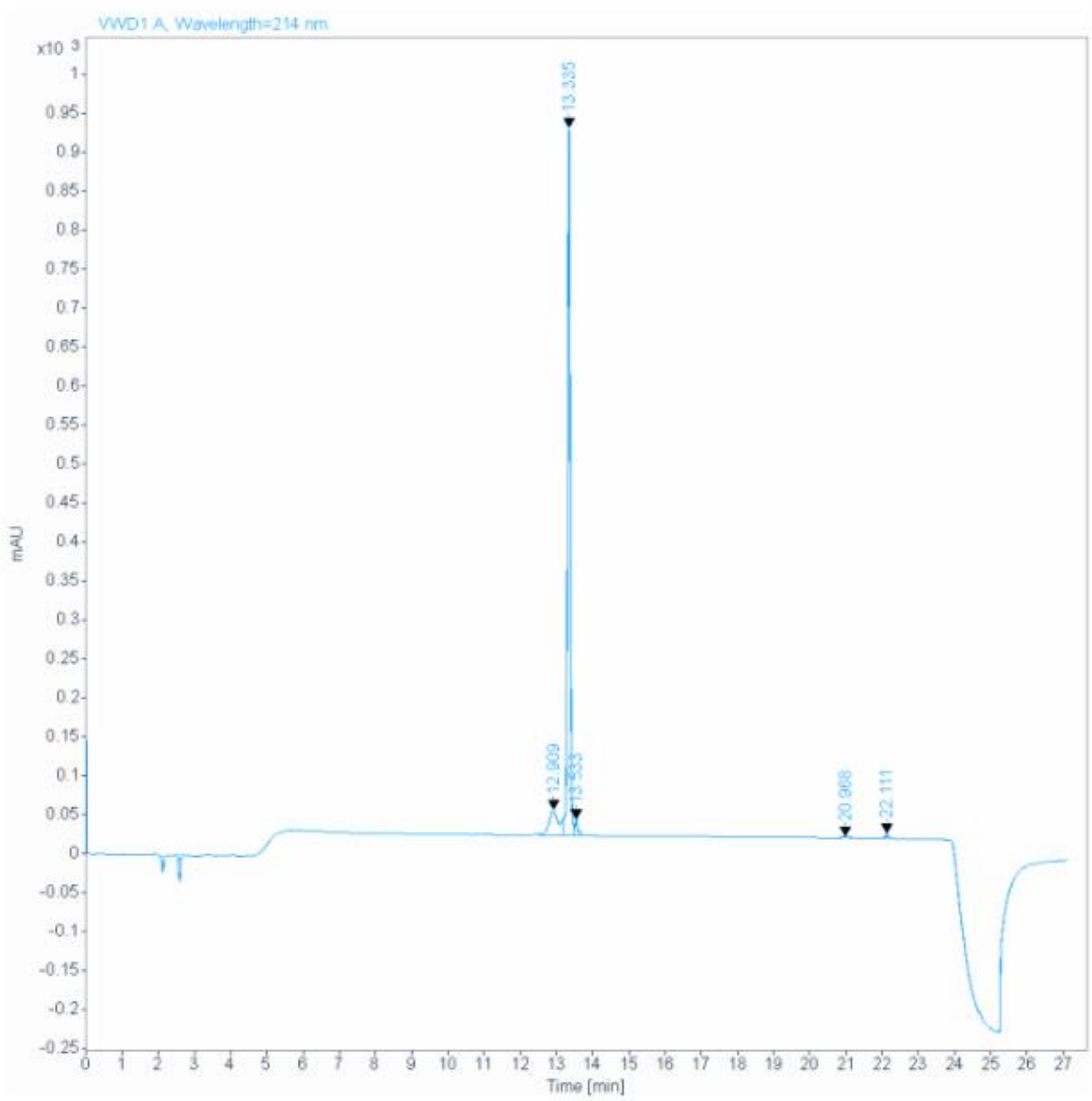

**Signal:** VWD1 A, Wavelength=214 nm

| RT [min] | Type | Width [min] | Area | Height | Area% | Name |
|---|---|---|---|---|---|---|
| 12.909 | MF | 0.3389 | 679.0752 | 33.3977 | 12.3935 | |
| 13.335 | MF | 0.0849 | 4632.4487 | 908.9108 | 84.5449 | |
| 13.533 | FM | 0.0988 | 127.4004 | 21.4998 | 2.3251 | |
| 20.968 | MM | 0.1232 | 17.8466 | 2.4138 | 0.3257 | |
| 22.111 | MM | 0.0615 | 22.5069 | 6.0992 | 0.4108 | |
| | | Sum | 5479.2778 | | | |

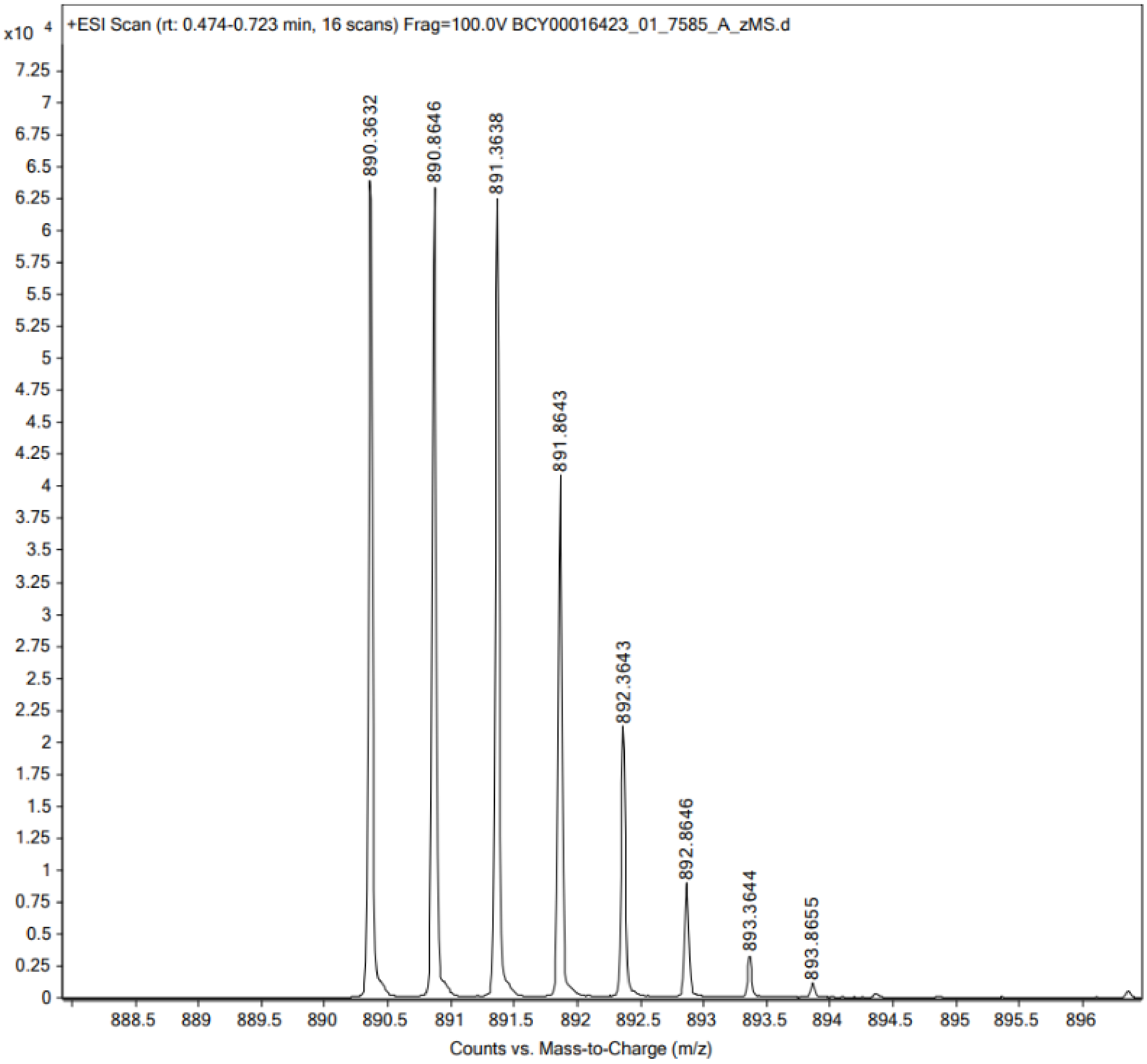
+ESI Scan (rt: 0.474-0.723 min, 16 scans) Frag=100.0V BCY00016423_01_7585_A_zMS.d
x10 4
890.3632
890.8646
891.3638
891.8643
892.3643
892.8646
893.3644
893.8655
Counts vs. Mass-to-Charge (m/z)

403

404 **Peptide 66**

**Data file:** D:\Chemstation\1\Data\Peptides_facon\7000-7999_BCY00016422_01_7584_A_zHPLC 15-26-42.D

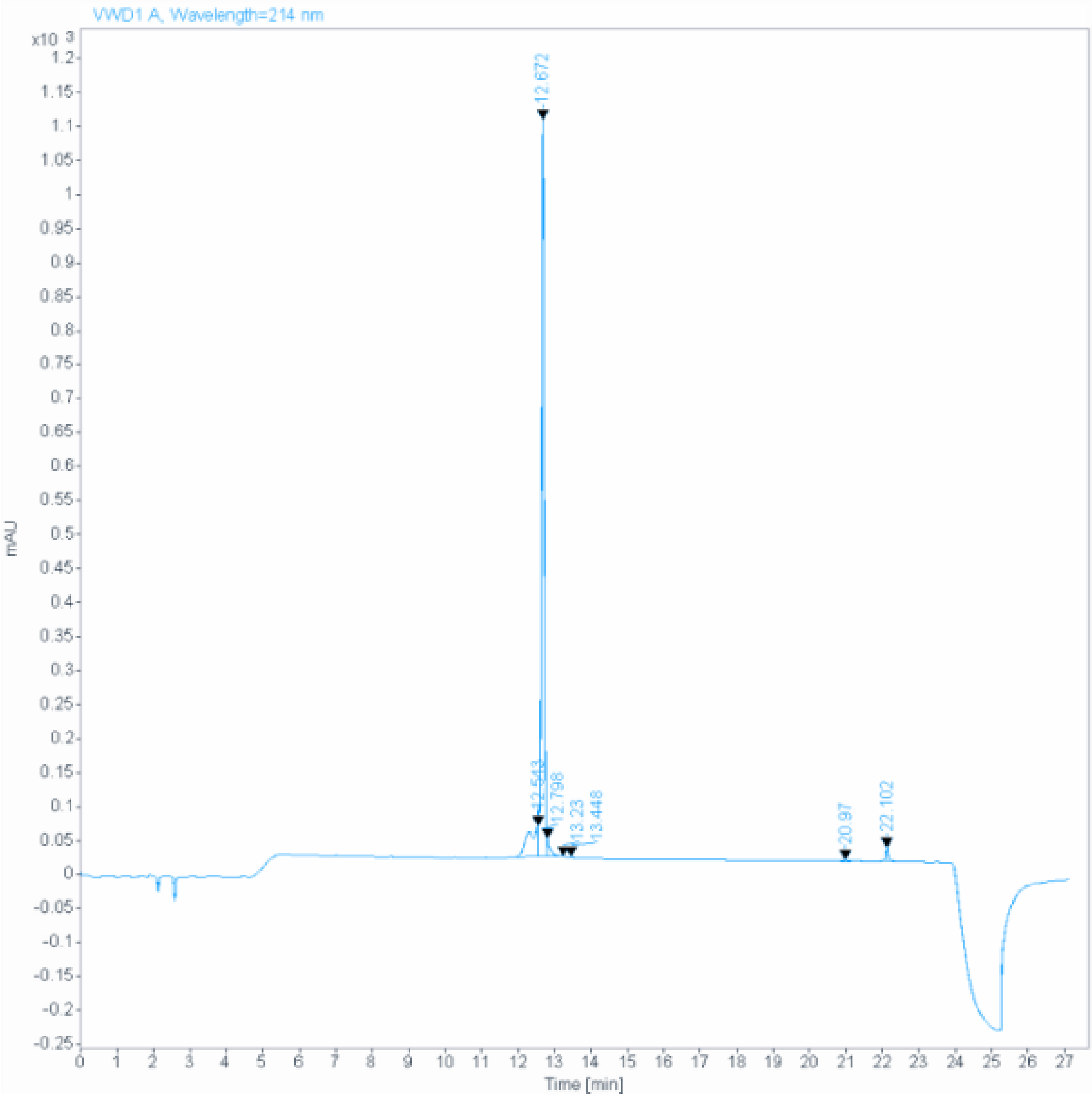

**Signal:** VWD1 A, Wavelength=214 nm

| RT [min] | Type | Width [min] | Area | Height | Area% | Name |
|---|---|---|---|---|---|---|
| 12.543 | MF | 0.3063 | 858.9743 | 46.7413 | 12.1752 | |
| 12.672 | FM | 0.0909 | 5917.3608 | 1084.9646 | 83.8735 | |
| 12.798 | FM | 0.0853 | 140.6630 | 27.4843 | 1.9938 | |
| 13.230 | MM | 0.1219 | 9.4092 | 1.2863 | 0.1334 | |
| 13.448 | MM | 0.0623 | 6.7239 | 1.8000 | 0.0953 | |
| 20.970 | MM | 0.2149 | 38.4804 | 2.9842 | 0.5454 | |
| 22.102 | MM | 0.0690 | 83.4887 | 20.1703 | 1.1834 | |
| | | Sum | 7055.1002 | | | |

405

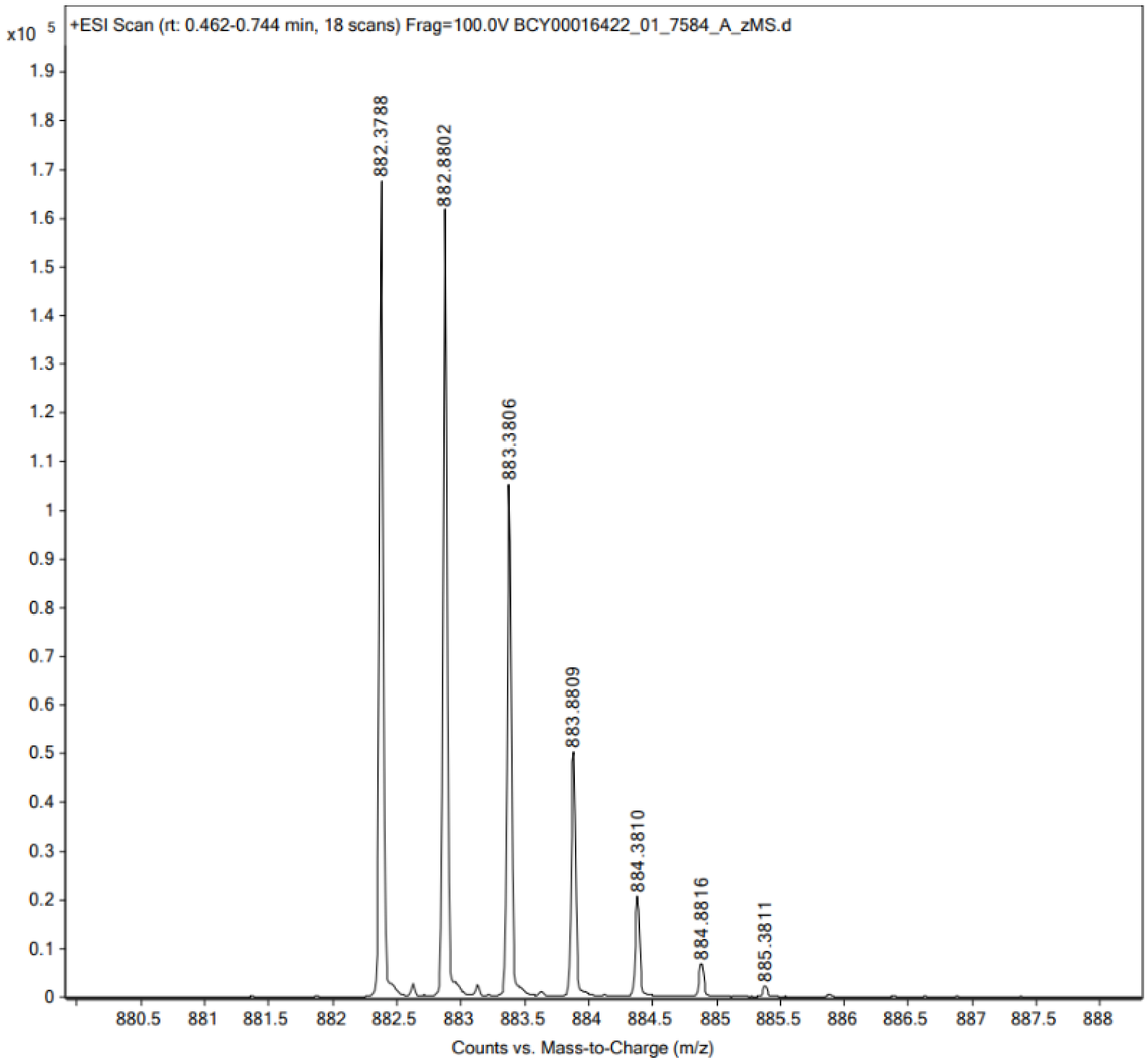

+ESI Scan (rt: 0.462-0.744 min, 18 scans) Frag=100.0V BCY00016422_01_7584_A_zMS.d
x10 5
882.3788
882.8802
883.3806
883.8809
884.3810
884.8816
885.3811
Counts vs. Mass-to-Charge (m/z)

406

407 **Peptide 67**

**Data file:** D:\Chemstation\1\Data\Peptides_facon\7000-7999_BCY00016415_01_7600_A_zHPLC 15-12-56.D

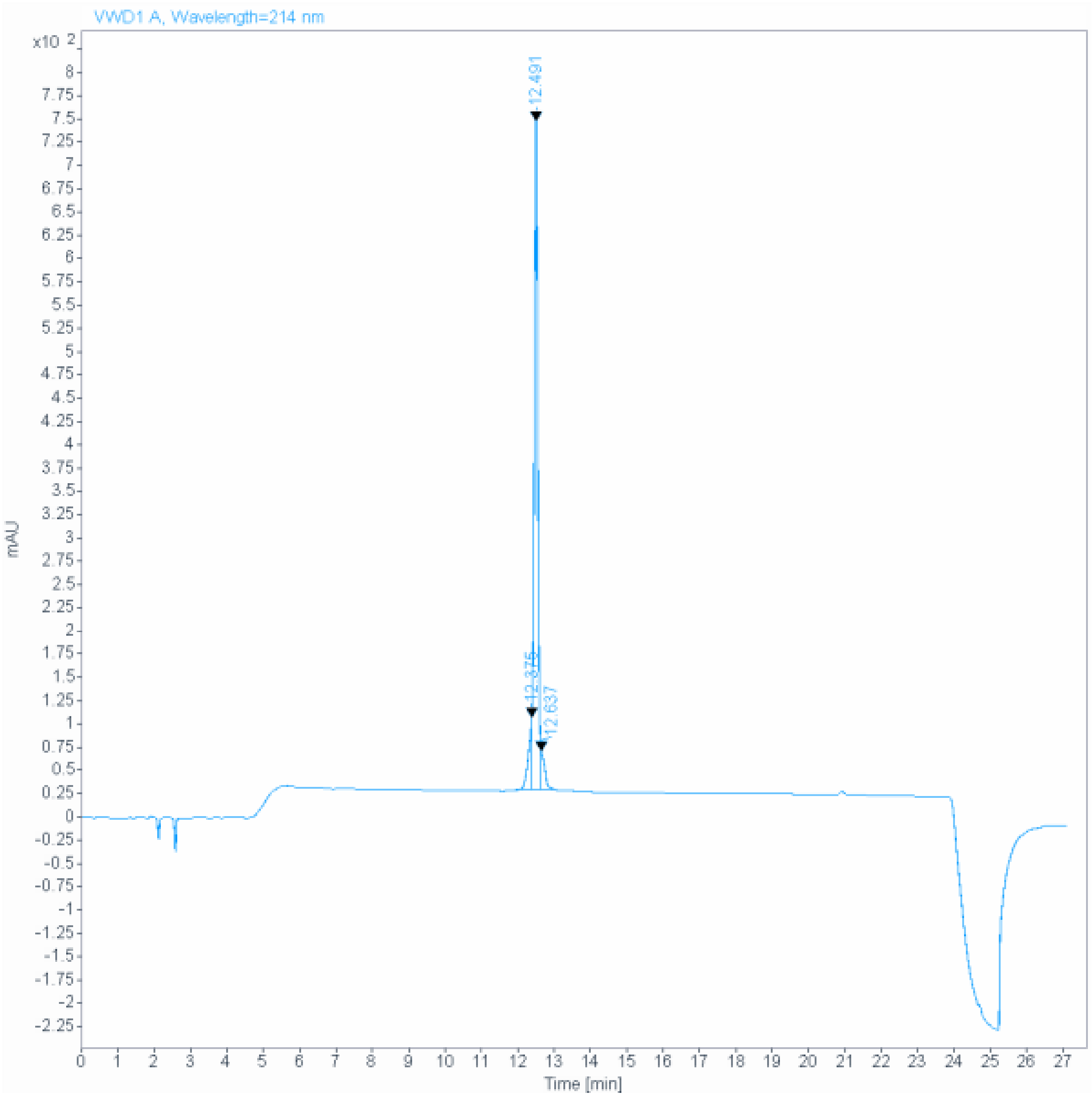

**Signal:** VWD1 A, Wavelength=214 nm

| RT [min] | Type | Width [min] | Area | Height | Area% | Name |
|---|---|---|---|---|---|---|
| 12.375 | MF | 0.1044 | 490.5663 | 78.2872 | 8.5694 | |
| 12.491 | FM | 0.1147 | 4946.5508 | 718.8433 | 86.4086 | |
| 12.637 | FM | 0.1135 | 287.4871 | 42.2170 | 5.0220 | |
| | | Sum | 5724.6042 | | | |

408

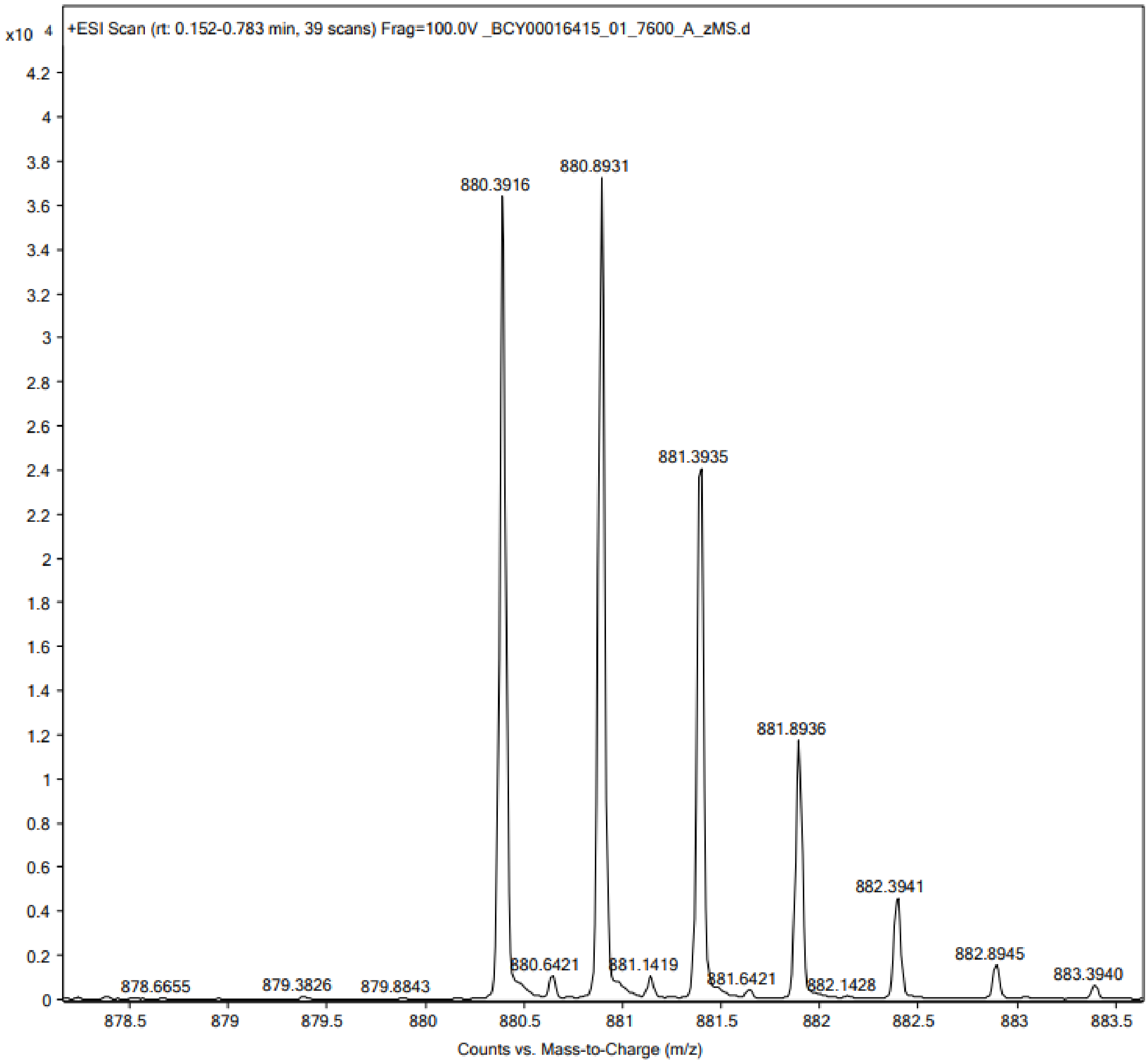
x10 4
+ESI Scan (rt: 0.152-0.783 min, 39 scans) Frag=100.0V _BCY00016415_01_7600_A_zMS.d
880.3916
880.8931
881.3935
881.8936
882.3941
882.8945
883.3940
880.6421
881.1419
881.6421
882.1428
878.6655
879.3826
879.8843
Counts vs. Mass-to-Charge (m/z)

409

### Peptide 68

**Data file:** D:\Chemstation\1\Data\Peptides_facon\7000-7999
_BCY00016416_01_7601_A_zHPLC 15-47-49.D

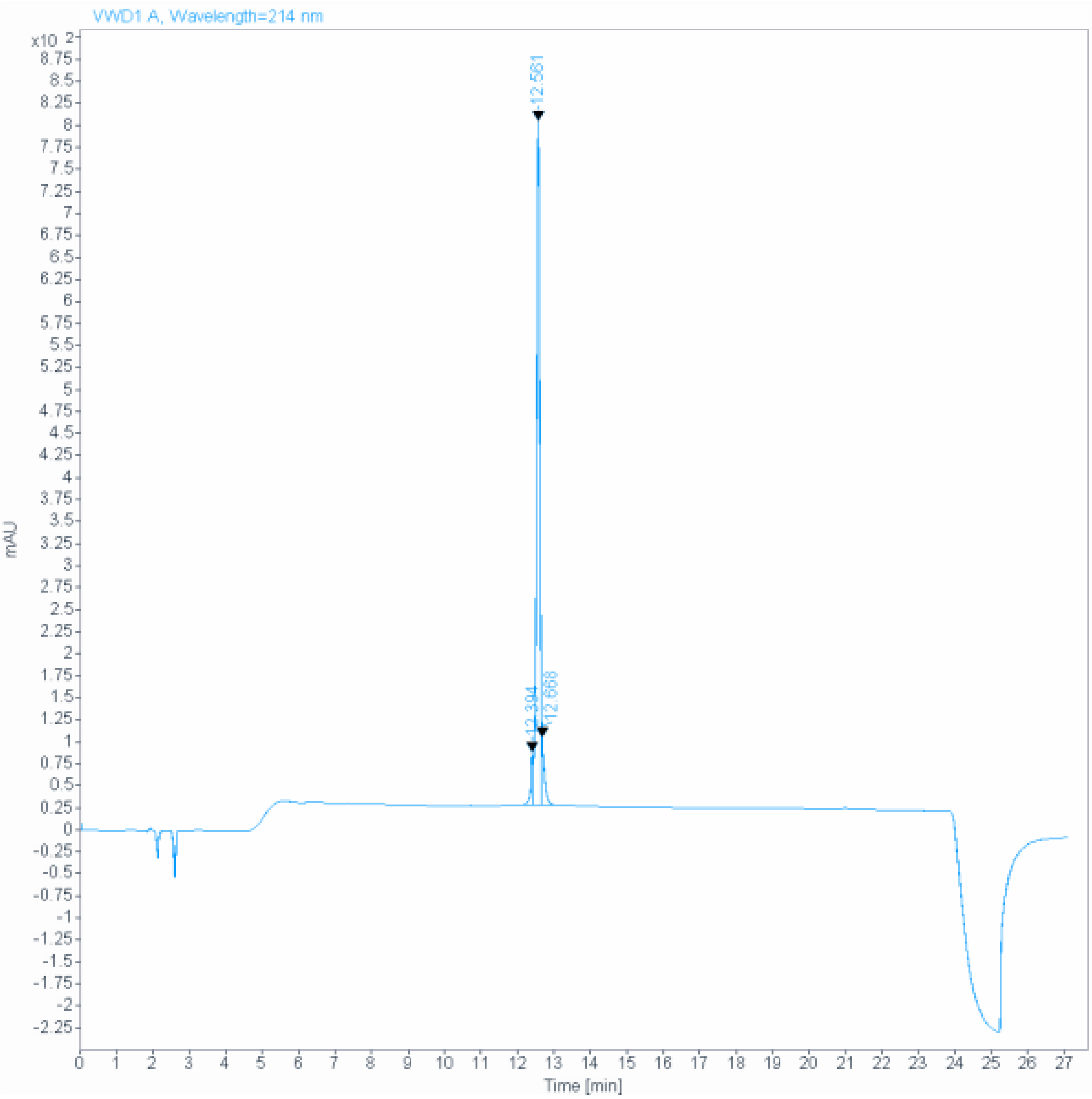

**Signal:** VWD1 A, Wavelength=214 nm

| RT [min] | Type | Width [min] | Area | Height | Area% | Name |
|---|---|---|---|---|---|---|
| 12.394 | MF | 0.0699 | 255.5601 | 60.8998 | 4.5475 | |
| 12.561 | MF | 0.1080 | 5041.1660 | 777.6755 | 89.7044 | |
| 12.668 | FM | 0.0696 | 323.0262 | 77.3246 | 5.7481 | |
| | | Sum | 5619.7524 | | | |

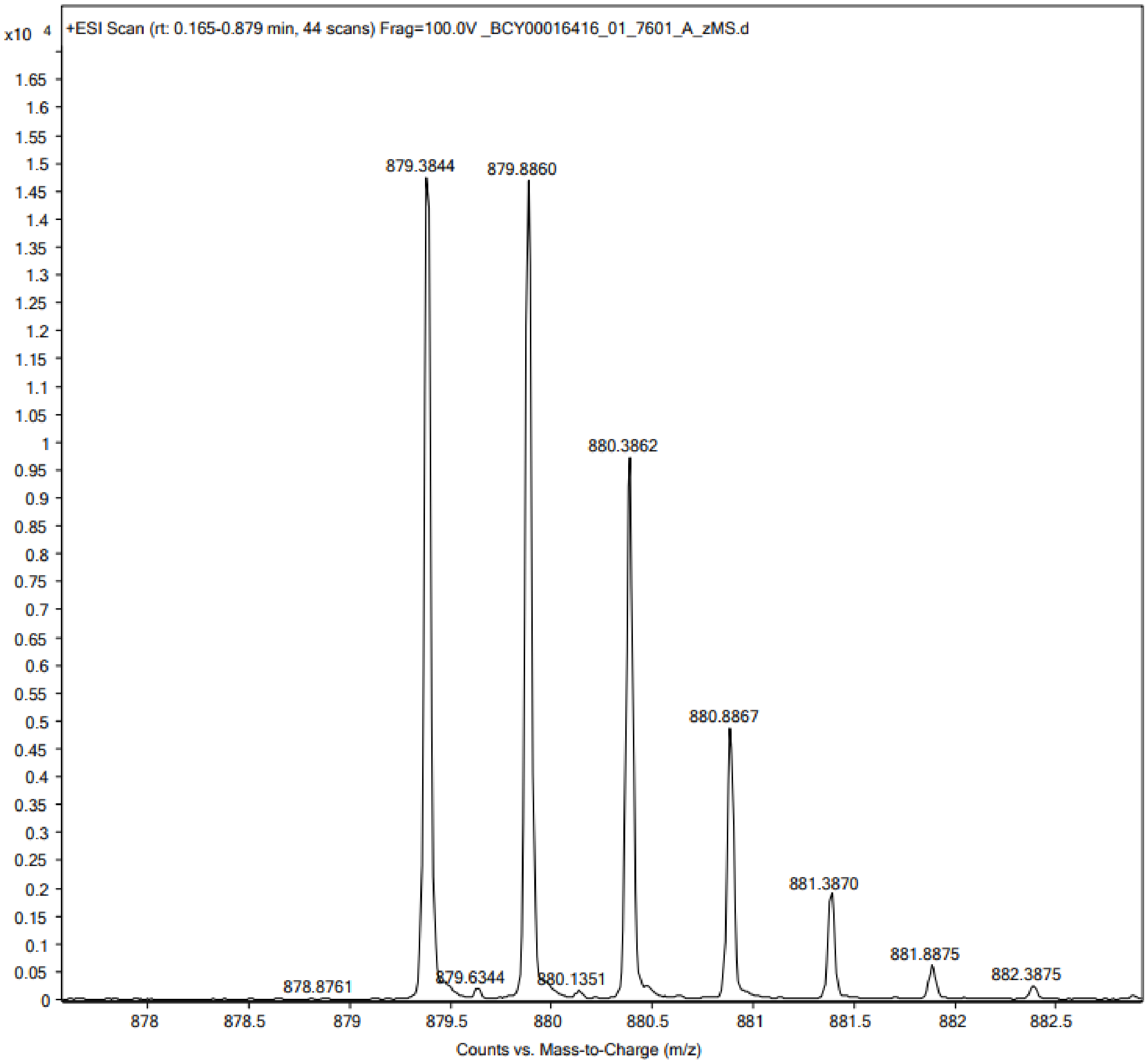

x10 4
+ESI Scan (rt: 0.165-0.879 min, 44 scans) Frag=100.0V _BCY00016416_01_7601_A_zMS.d
879.3844
879.8860
880.3862
880.8867
881.3870
881.8875
882.3875
878.8761
879.6344
880.1351
Counts vs. Mass-to-Charge (m/z)

412

#### Peptide 69

| | |
|---|---|
| Sample name: | (101-08-00) D-Ala7 (TATA) QC |
| Instrument: | 1260_2 |
| Injection date: | 12/12/2019 3:28:14 PM |
| Acq. method: | 0595B_AB_POROSHELL120_15.5MIN.M |

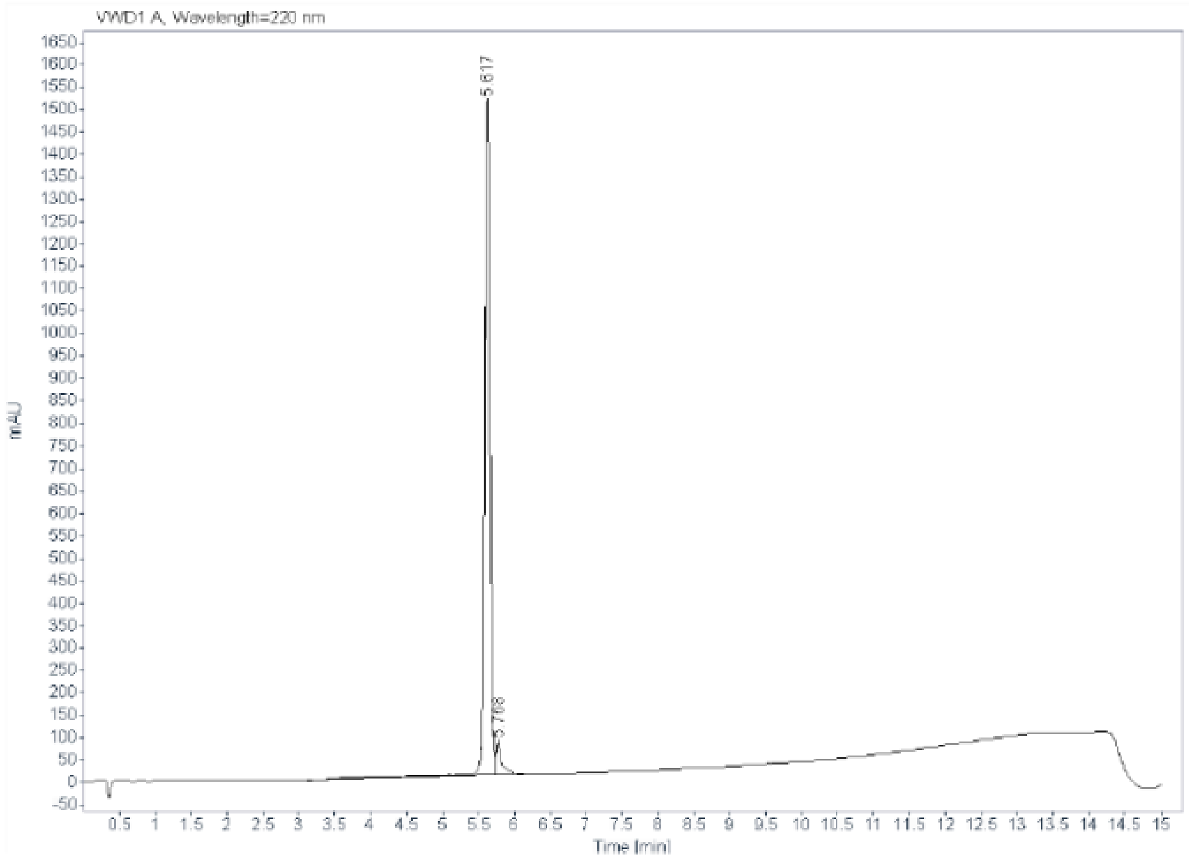

**Signal:** VWD1 A, Wavelength=220 nm

| RT [min] | Type | Width [min] | Area | Height | Area% |
|---|---|---|---|---|---|
| 5.617 | VV R | 0.0816 | 8202.7715 | 1505.3533 | 95.1574 |
| 5.768 | VB | 0.0800 | 417.4467 | 73.7483 | 4.8426 |
| | | Sum | 8620.2182 | | |

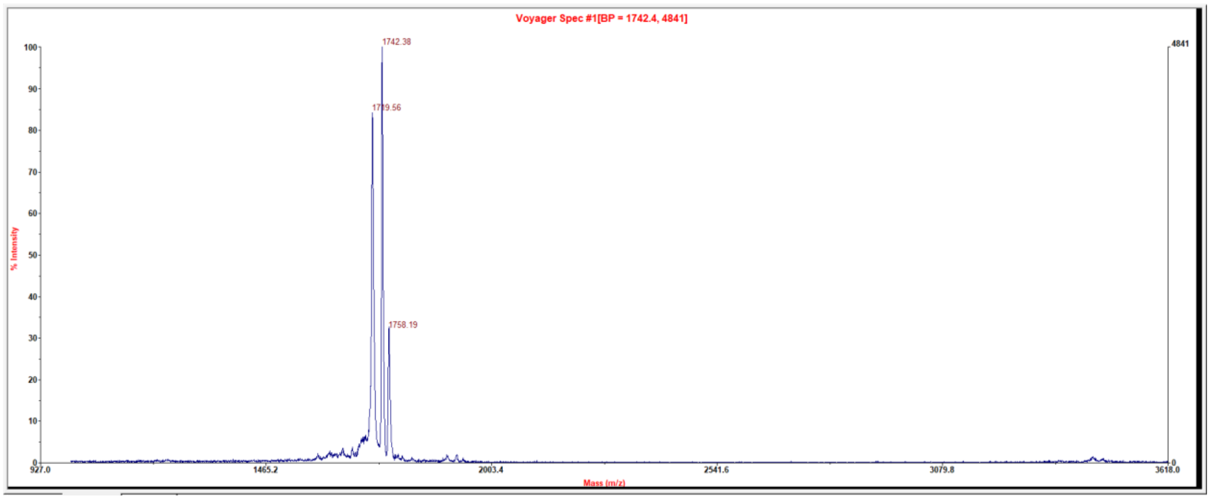

### Peptide 70

| | |
|---|---|
| Sample name: | A-(101-08-00) HArg4 |
| Instrument: | 1260_1 |
| Injection date: | 5/21/2021 1:13:27 PM |
| Acq. method: | 0595B_AB_KxC18_QC _20min.M |

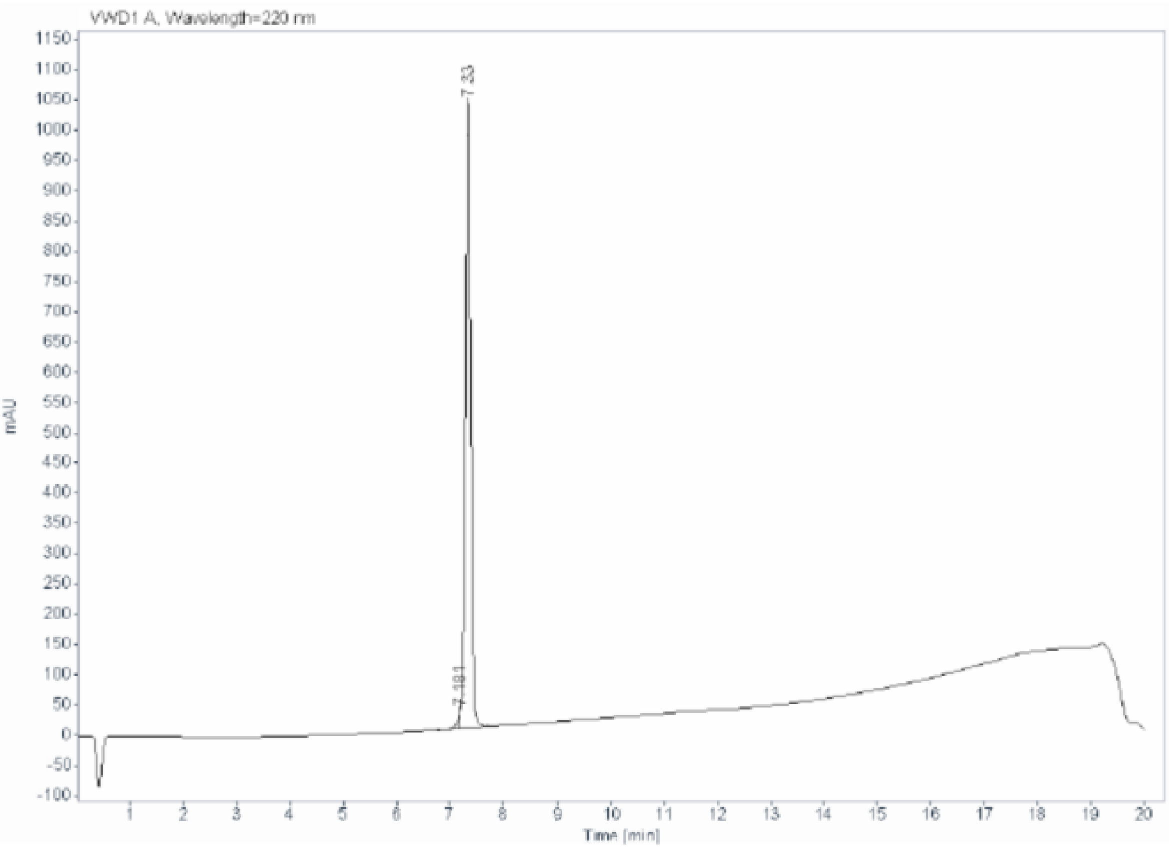

Signal: VWD1 A, Wavelength=220 nm

| RT [min] | Type | Width [min] | Area | Height | Area% |
|---|---|---|---|---|---|
| 7.181 | MF | 0.0577 | 105.6076 | 30.5107 | 1.4226 |
| 7.330 | FM | 0.1173 | 7317.7363 | 1039.9047 | 98.5774 |
| | | Sum | 7423.3439 | | |

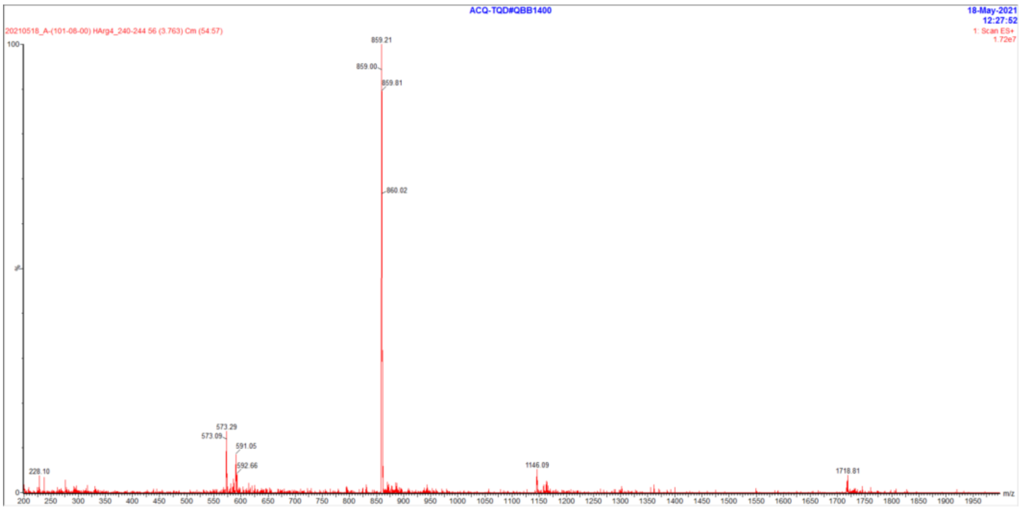

### Peptide 71

**Sample name:** (101-08-00)-A HArg4
**Instrument:** 1260_1
**Injection date:** 5/21/2021 10:00:45 AM
**Acq. method:** 0595B_AB_KxC18_QC _20min.M

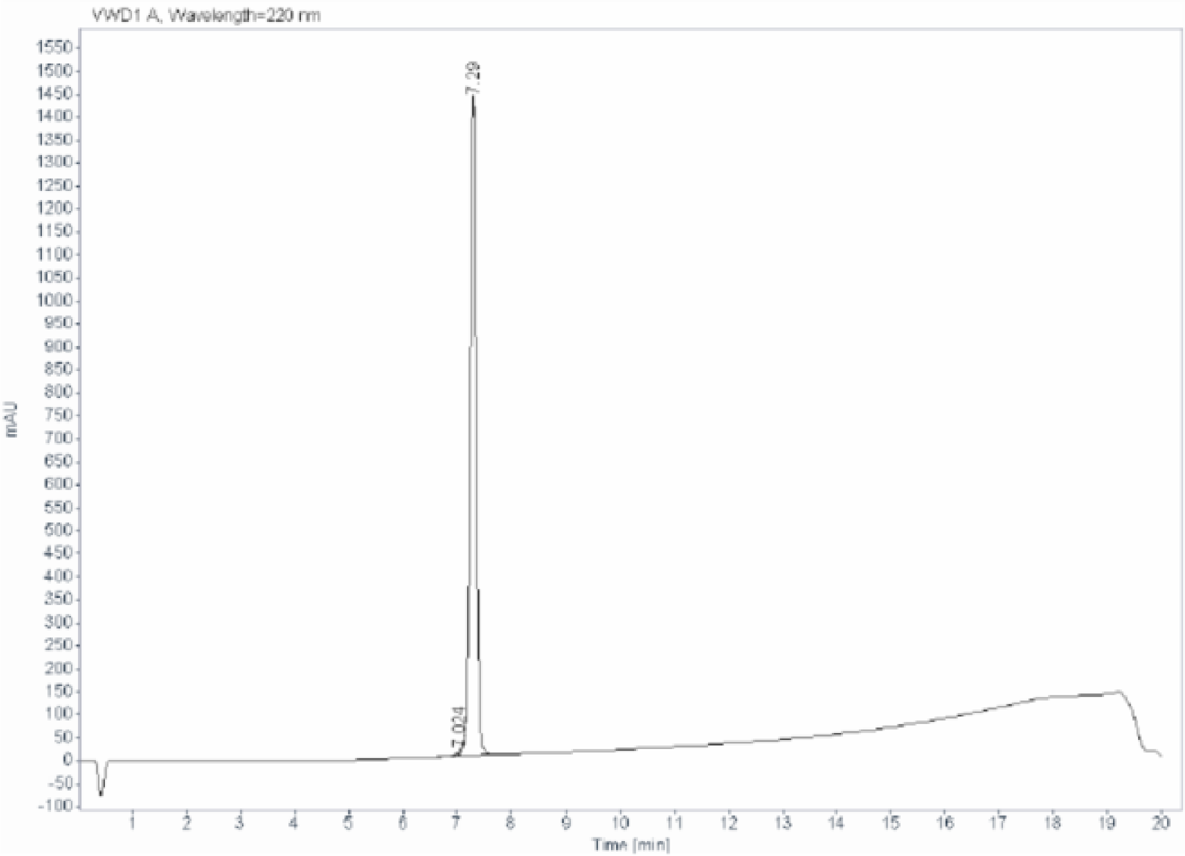

**Signal:** VWD1 A, Wavelength=220 nm

| RT [min] | Type | Width [min] | Area | Height | Area% |
|---|---|---|---|---|---|
| 7.024 | BV E | 0.1226 | 76.7479 | 8.5584 | 0.6741 |
| 7.290 | VB R | 0.1223 | 11309.2490 | 1430.0098 | 99.3259 |
| | | Sum | 11385.9969 | | |

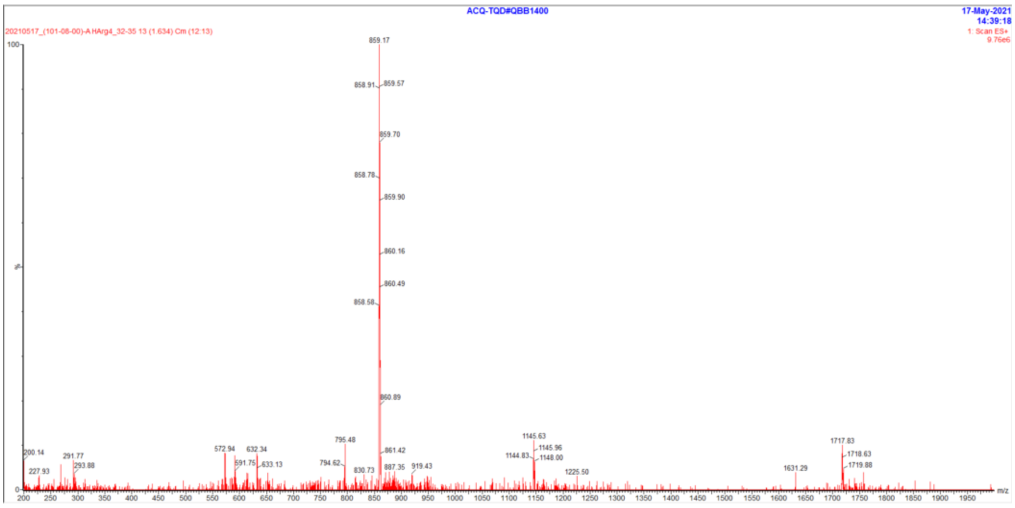

#### Peptide 72

| | |
|---|---|
| Sample name: | (101-08-00) HArg4 |
| Instrument: | 1260_1 |
| Injection date: | 5/21/2021 11:05:35 AM |
| Acq. method: | 0595B_AB_KxC18_QC _20min.M |

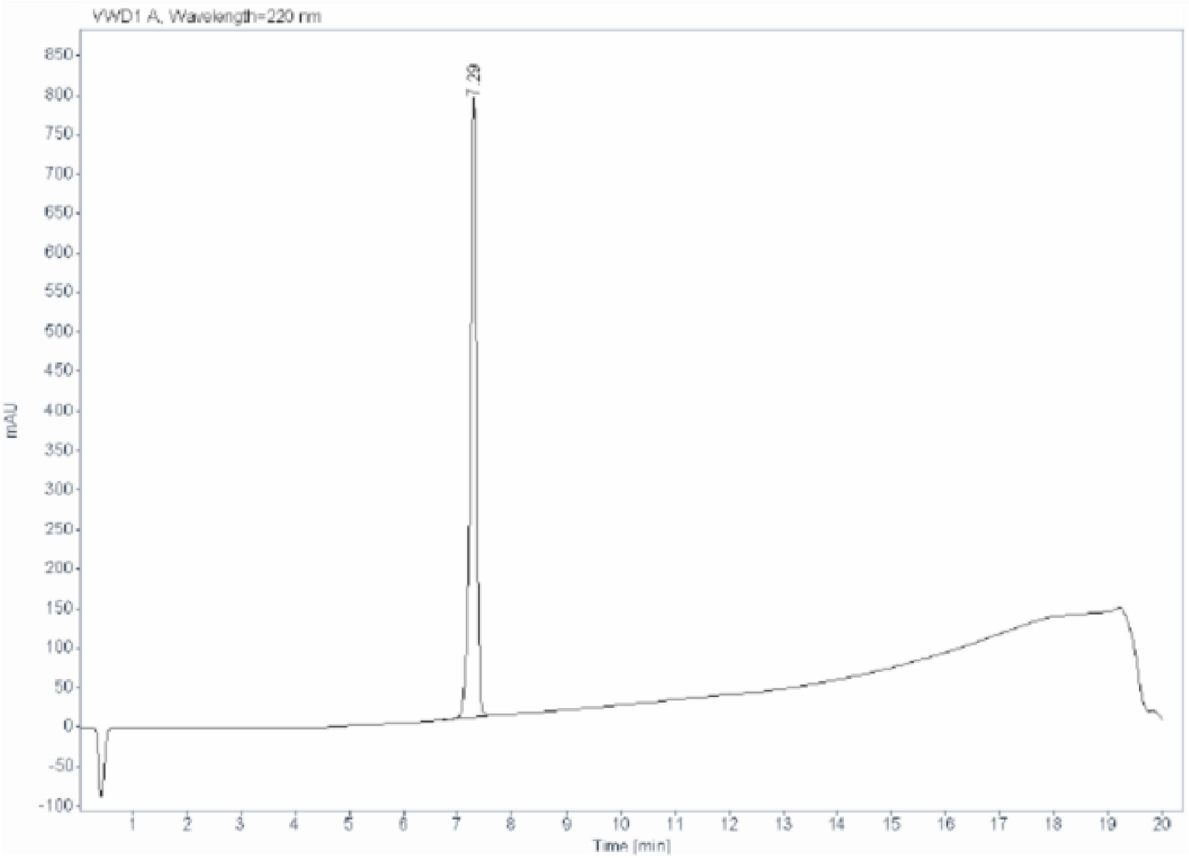

Signal: VWD1 A, Wavelength=220 nm

| RT [min] | Type | Width [min] | Area | Height | Area% |
|---|---|---|---|---|---|
| 7.290 | BB | 0.1211 | 6378.9600 | 782.3913 | 100.0000 |
| | | Sum | 6378.9600 | | |

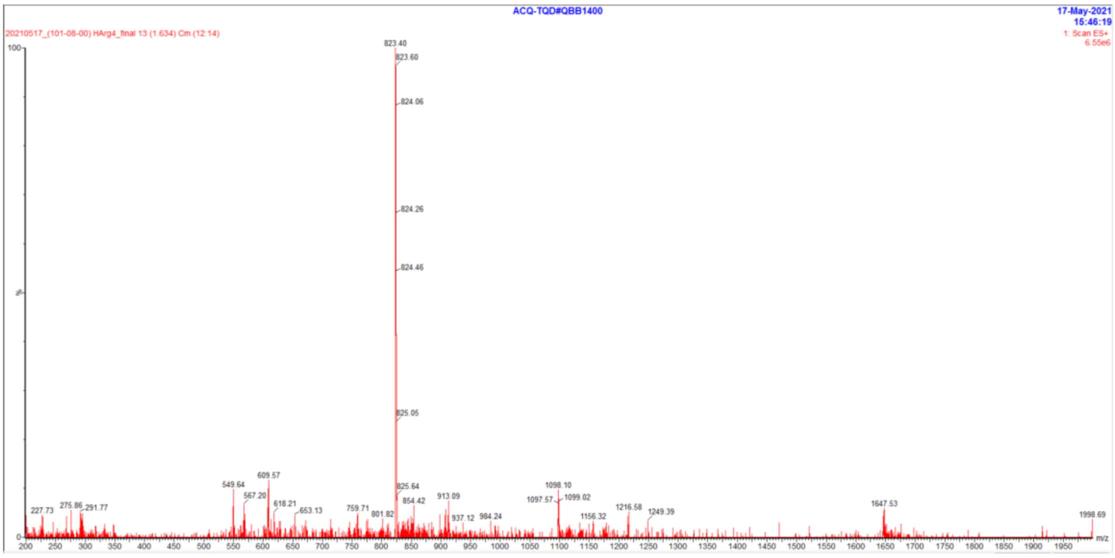

### Peptide 73

| | |
|---|---|
| Sample name: | Ac-A-(101-08-00)(TATA)-A HArg4 |
| Instrument: | 1260_2 |
| Injection date: | 2/23/2021 1:57:49 PM |
| Acq. method: | 0595B_AB_KxC18_QC _20min.M |

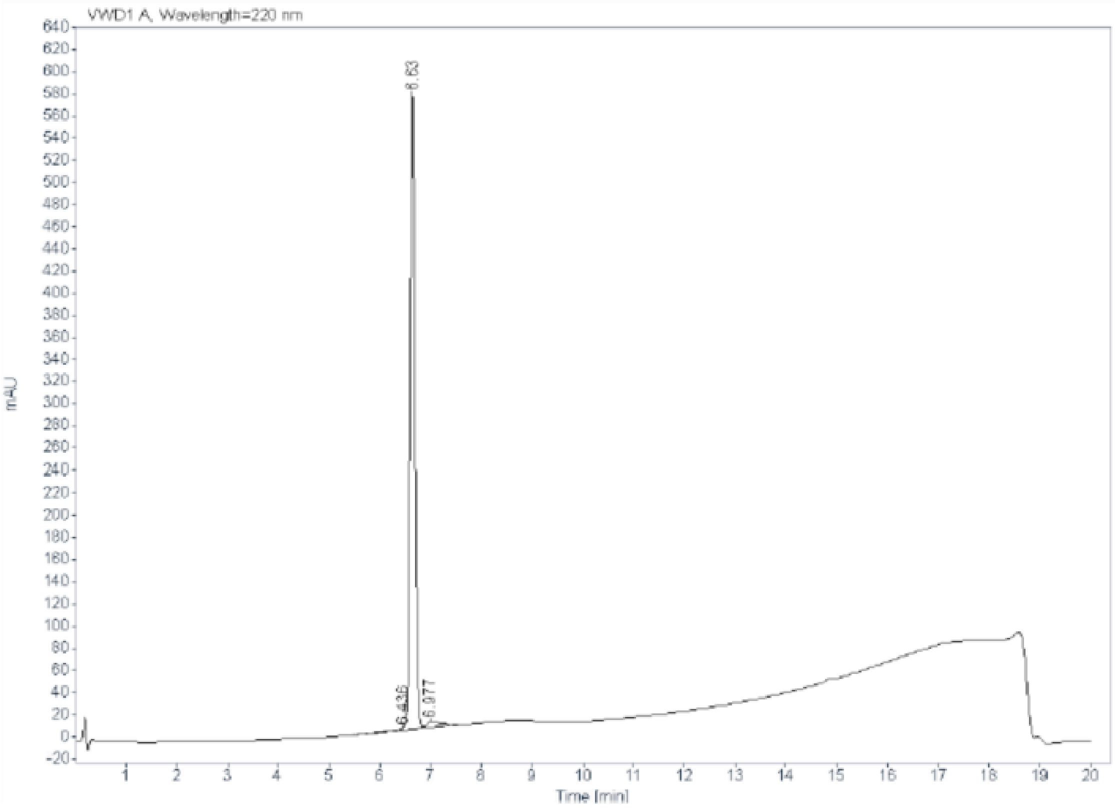

Signal: VWD1 A, Wavelength=220 nm

| RT [min] | Type | Width [min] | Area | Height | Area% |
|---|---|---|---|---|---|
| 6.436 | MF | 0.3005 | 33.3237 | 1.8482 | 0.8637 |
| 6.630 | FM | 0.1079 | 3719.9207 | 574.5839 | 96.4149 |
| 6.977 | FM | 0.3036 | 104.9992 | 5.7642 | 2.7214 |
| | | Sum | 3858.2436 | | |

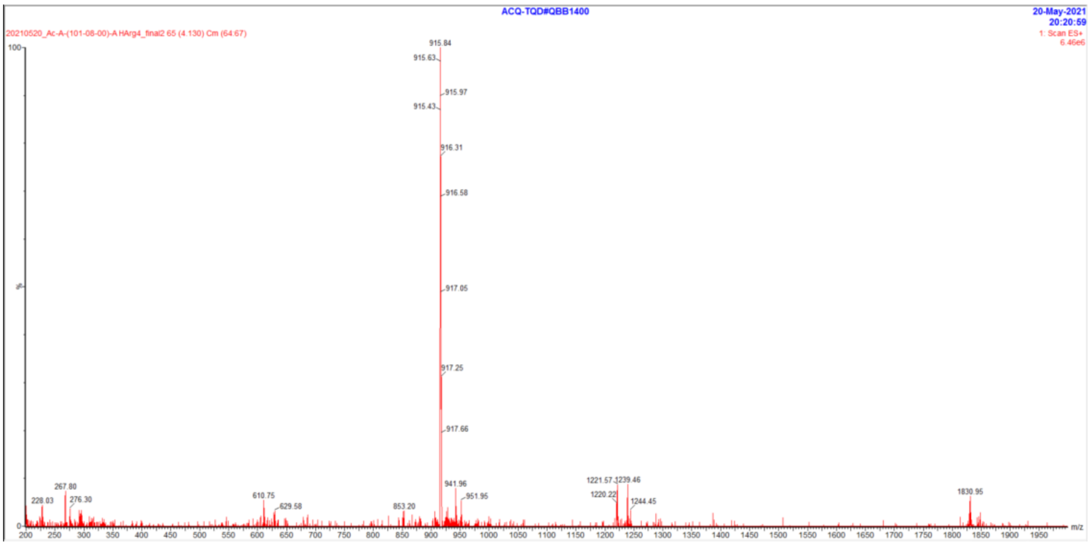

### Peptide 74

| | |
|---|---|
| Sample name: | Ac-(101-08-00)-A HArg4 |
| Instrument: | 1260_1 |
| Injection date: | 5/20/2021 10:39:30 AM |
| Acq. method: | 0595B_AB_KxC18_QC_20min.M |

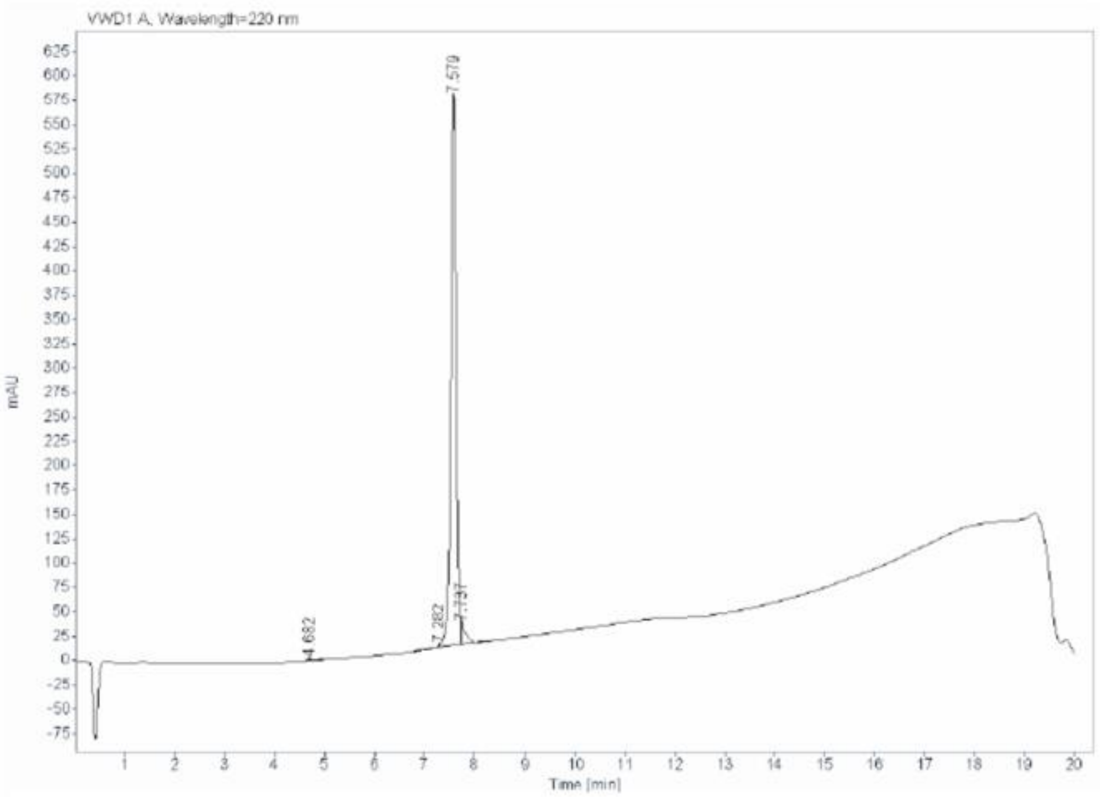

Signal: VWD1 A, Wavelength=220 nm

| RT [min] | Type | Width [min] | Area | Height | Area% |
|---|---|---|---|---|---|
| 4.682 | BB | 0.1267 | 12.5235 | 1.5108 | 0.2884 |
| 7.282 | MF | 0.1087 | 13.9476 | 2.1379 | 0.3212 |
| 7.579 | FM | 0.1236 | 4195.7515 | 565.9112 | 96.6260 |
| 7.737 | FM | 0.0961 | 120.0347 | 20.8225 | 2.7643 |
| | | Sum | 4342.2573 | | |

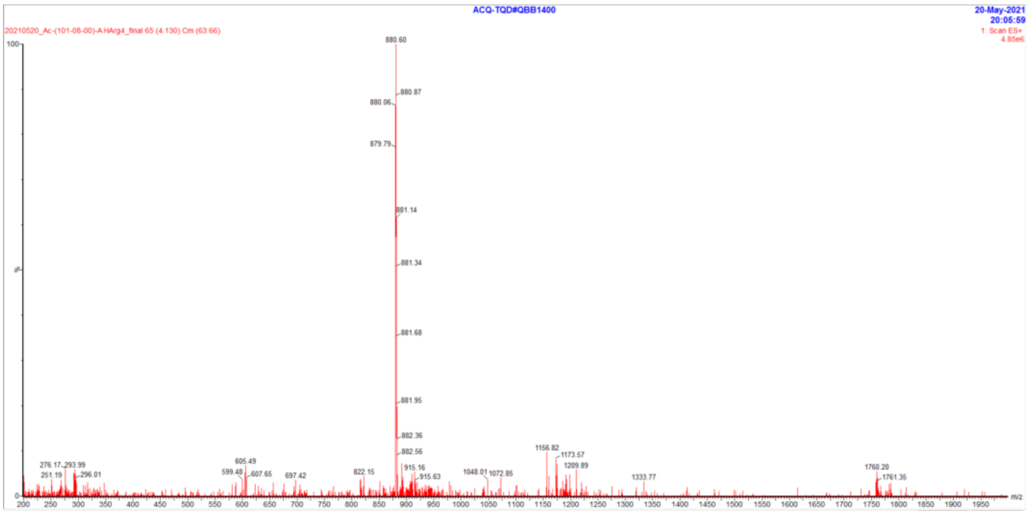

### Peptide 75

**Sample name:** Ac-A-(101-08-00)(TATA)-A HArg4 6ClTrp
**Instrument:** 1260_2
**Injection date:** 2/23/2021 3:26:25 PM
**Acq. method:** 0595B_AB_KxC18_QC_20min.M

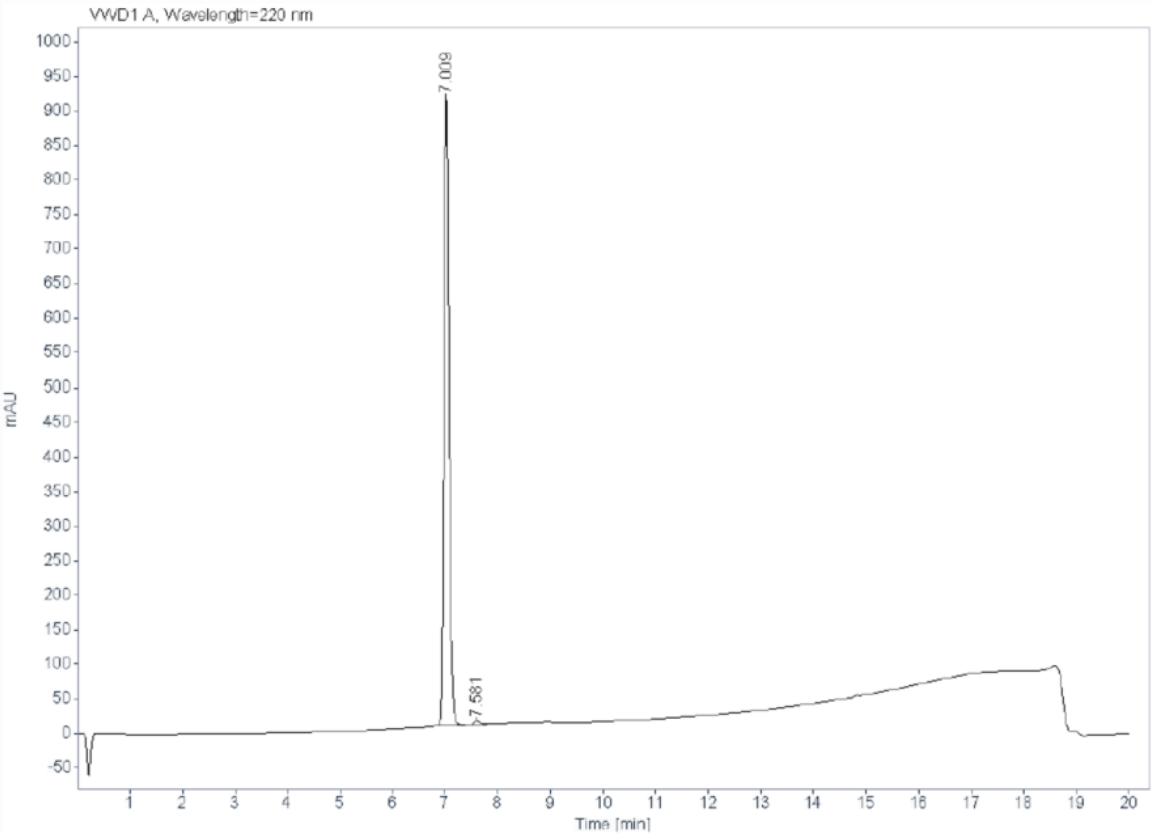

**Signal:** VWD1 A, Wavelength=220 nm

| RT [min] | Type | Width [min] | Area | Height | Area% |
|---|---|---|---|---|---|
| 7.009 | FM | 0.1086 | 5949.2568 | 912.7531 | 98.7811 |
| 7.581 | FM | 0.1850 | 73.4126 | 6.6154 | 1.2189 |
| | | Sum | 6022.6694 | | |

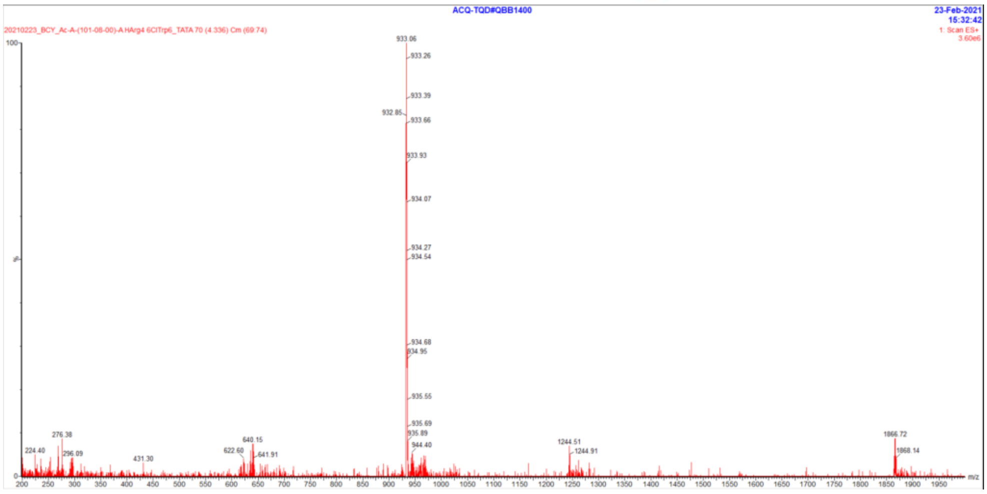

### Peptide 76

**Sample name:** (101-08-00)(TATA)[D-Form] QC
**Instrument:** 1260_2

**Injection date:** 11/11/2019 4:18:26 PM
**Acq. method:** 0595B_AB_Poroshell1
20_15.5min.M

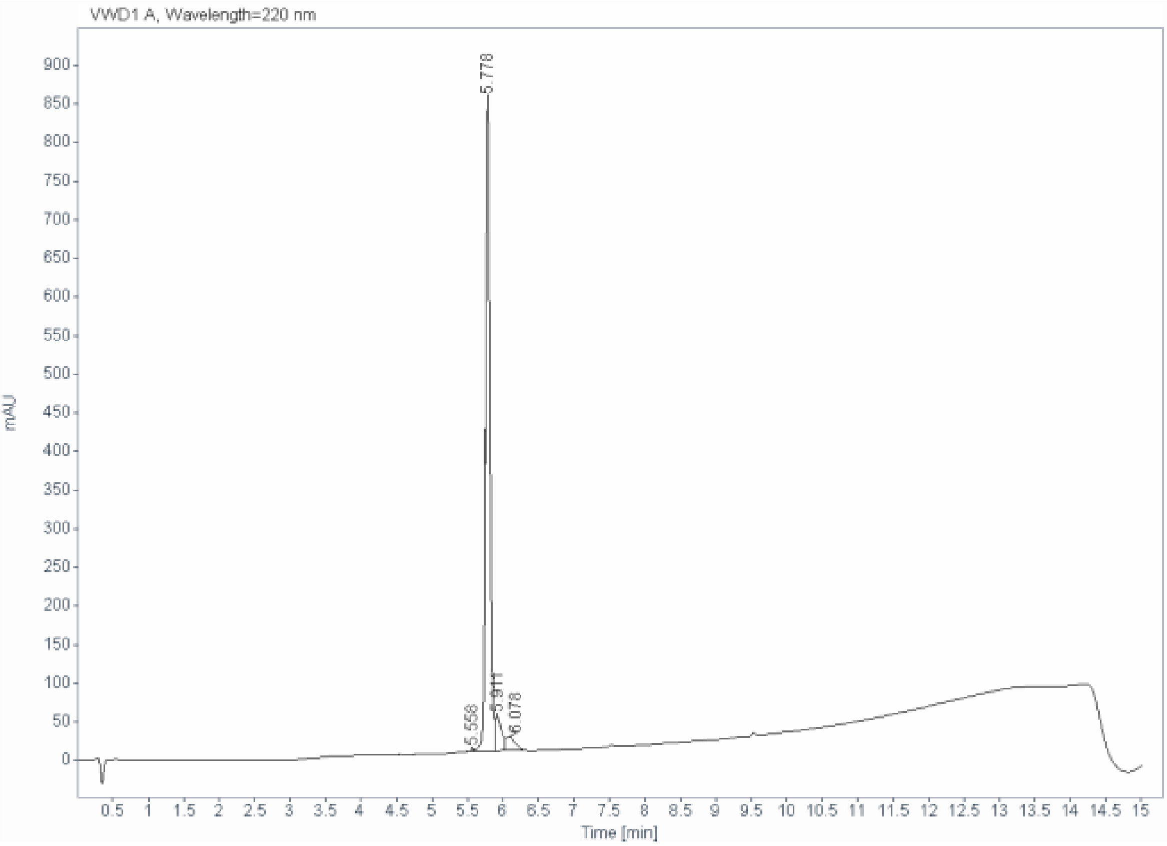

**Signal:** VWD1 A, Wavelength=220 nm

| RT [min] | Type | Width [min] | Area | Height | Area% |
|---|---|---|---|---|---|
| 5.558 | BV | 0.0504 | 10.5694 | 3.1286 | 0.2553 |
| 5.778 | VV | 0.0667 | 3681.5608 | 847.9104 | 88.9329 |
| 5.911 | VV | 0.0845 | 283.0704 | 45.5661 | 6.8379 |
| 6.078 | VB | 0.1286 | 164.5071 | 18.6974 | 3.9739 |
| | | Sum | 4139.7077 | | |

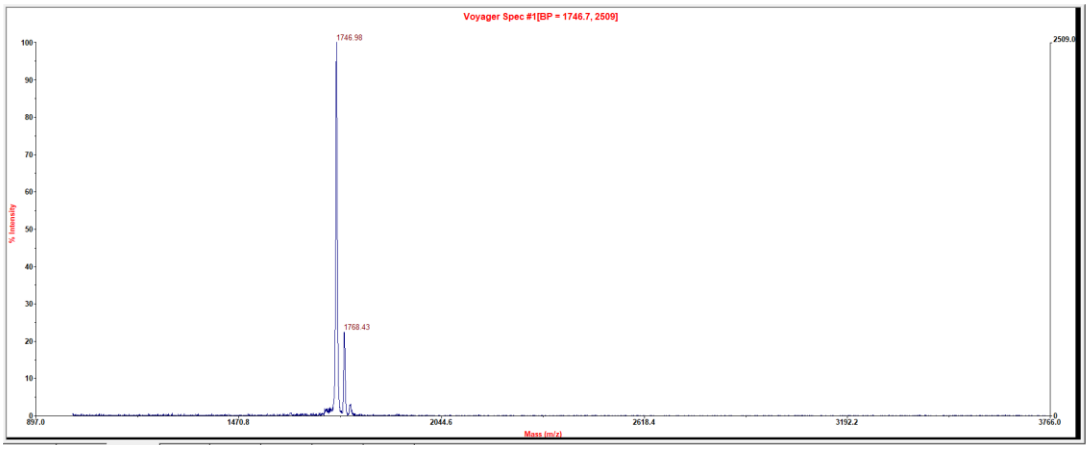

#### Conjugate 1

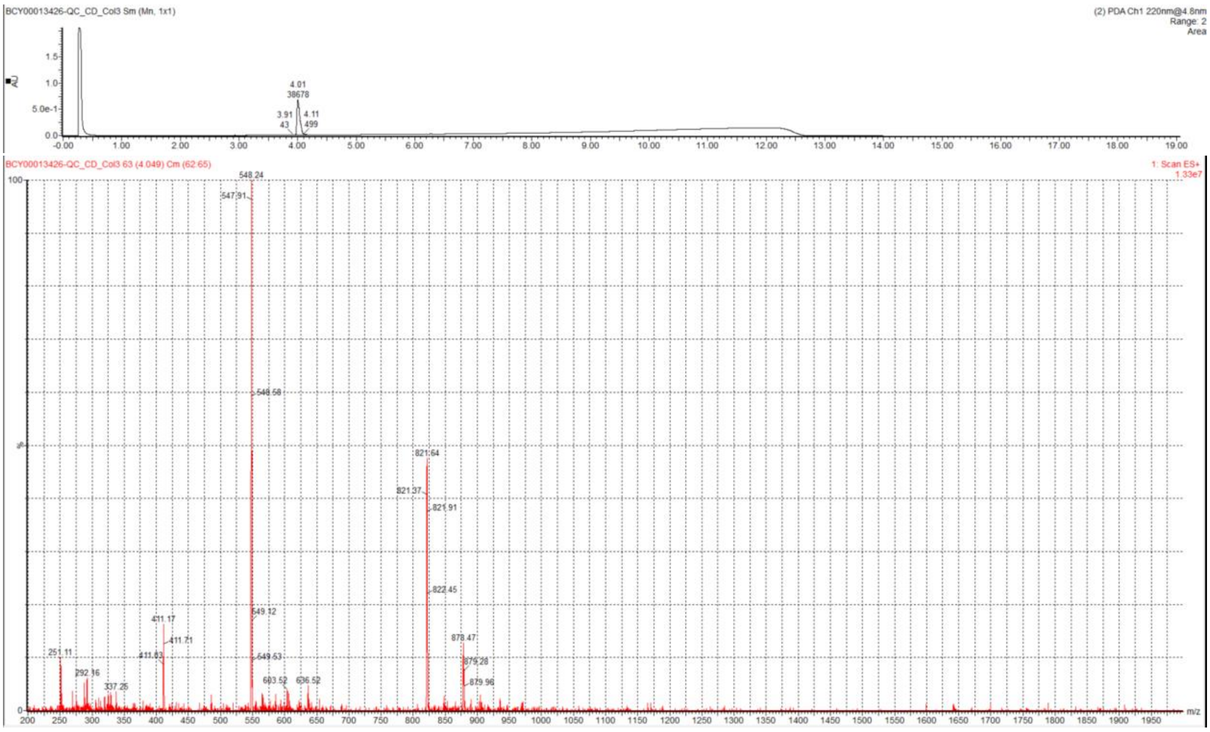

#### Conjugate 2

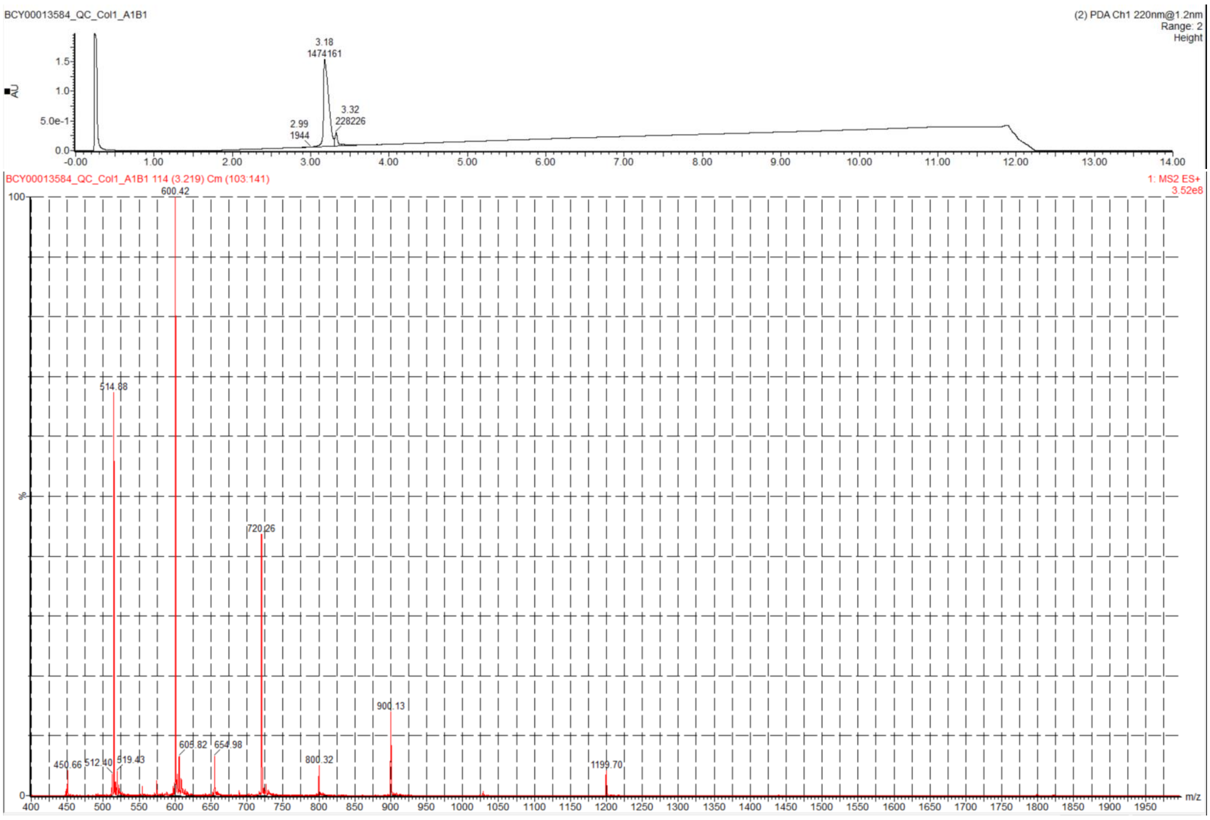

### Conjugate 3

| | | | |
|---|---|---|---|
| **Data file:** | ES19551-238-P1B.dx | | |
| **Sequence Name:** | 2021-10-15 14-26-12+08-00 | **Project Name:** | Installation HPLC |
| **Sample name:** | ES19551-238-P1B | **Operator:** | SYSTEM |
| **Instrument:** | LCMS-QC | **Injection date:** | 2021-10-15 23:41:29+08:00 |
| **Inj. volume:** | 8.000 | **Location:** | P1-B9 |
| **Acq. method:** | 20-50-20min-P1waste.amx | **Colum:** | Gemini-NX C18 5um 110A 150mm4.6 |
| **Processing method:** | *GC_LC Area Percent_DefaultMethod.pmx | **Mobile phase:** | A:0.1%TFAin H2O B:0.1%TFAin ACN |

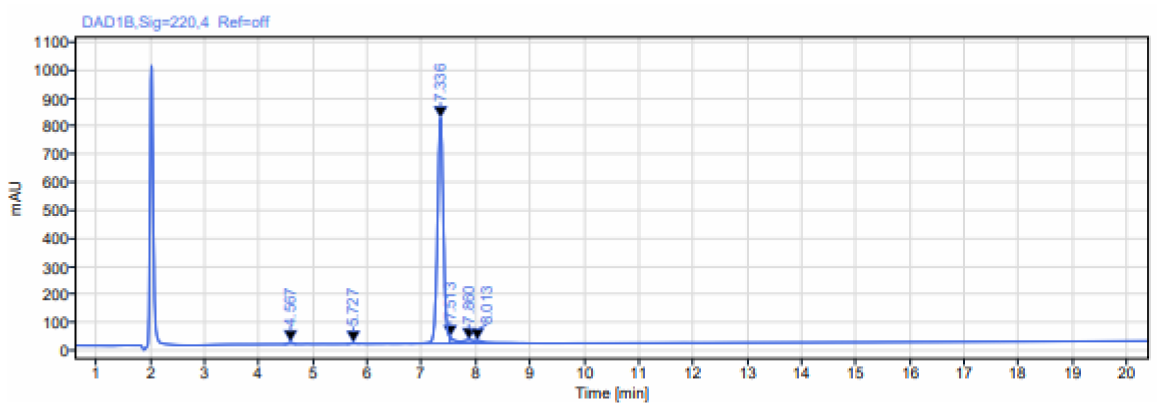

**Signal:** DAD1B,Sig=220,4 Ref=off

| RT [min] | Type | Width [min] | Area | Height | Area% | Name |
|---|---|---|---|---|---|---|
| 4.567 | MM m | 0.24 | 38.11 | 6.87 | 0.58 | |
| 5.727 | MM m | 0.26 | 17.01 | 2.99 | 0.26 | |
| 7.336 | BM m | 0.74 | 6107.46 | 802.41 | 92.86 | |
| 7.513 | MV m | 0.19 | 128.50 | 21.92 | 1.95 | |
| 7.860 | VV | 0.26 | 151.00 | 12.11 | 2.30 | |
| 8.013 | VB | 0.81 | 135.02 | 9.53 | 2.05 | |
| | | **Sum** | **6577.10** | | | |

| | | | |
|---|---|---|---|
| **Sequence Name:** | 2021-10-15 11-53-51+08-00 | | |
| **Data file:** | ES19551-238-P1A.dx | **Project Name:** | Installation |
| **Sample name:** | ES19551-238-P1A | **Operator:** | SYSTEM |
| **Instrument:** | LCMS-QC | **Acquired on:** | 2021-10-15 15:31:35+08:00 |
| **Inj. volume:** | 10.000 | **Location:** | P1-B9 |
| **Acq. method:** | 10-80-3min-1.5.amx | | |
| **Processing method:** | *MS_DefaultMethod.pmx | | |

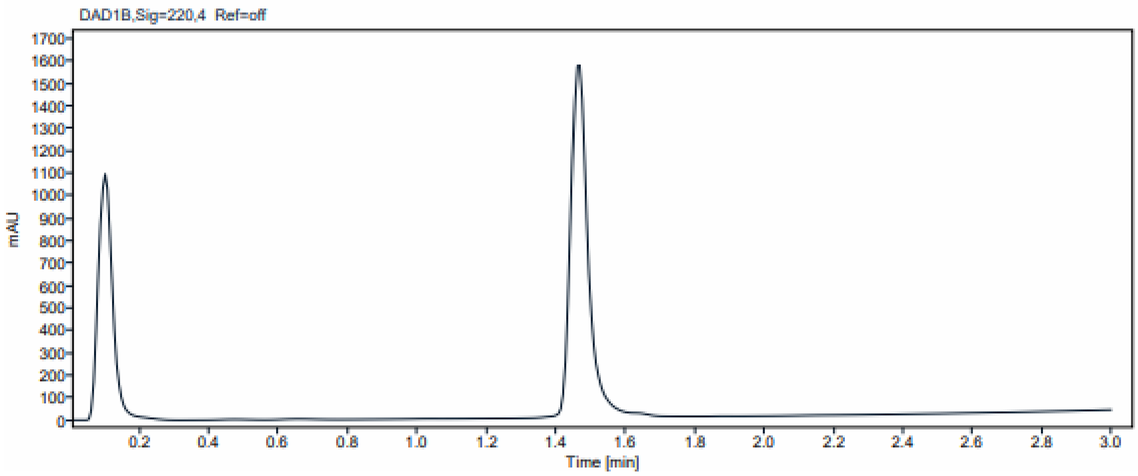

#### Peak Results (Area Percent at least 1%)

| RT (min) | Signal Description | Width (min) | Area | Height | Area% |
|---|---|---|---|---|---|
| 1.470 | MS1 +TIC SCAN ESI Frag=135V Gain=1.0 | 0.148 | 1947137.1 | 623043.0 | 100.00 |
| | **Sum MS1 +TIC SCAN ESI Frag=135V Gain=1.0** | | 1947137.1 | | |

450

**Signal Name** MS1 +TIC SCAN ESI Frag=135V Gain=1.0

**Peak Retention Time** 1.470

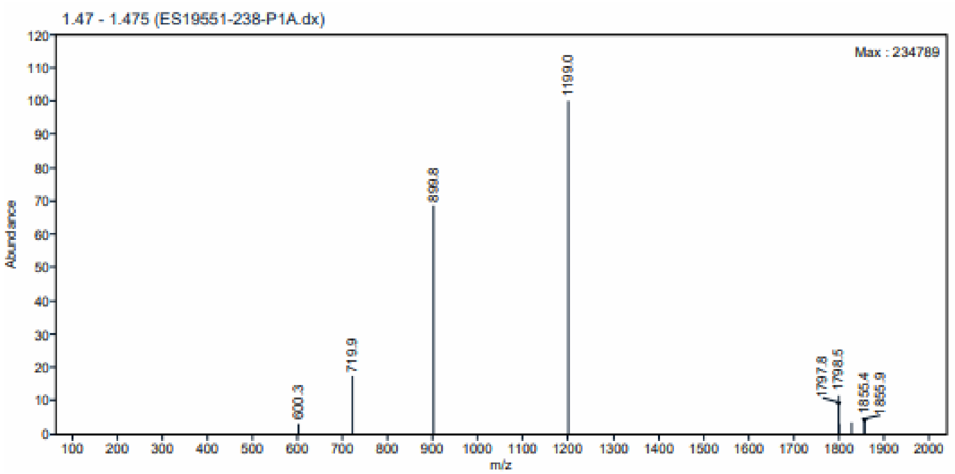

**MS Peak Table**

| m/z | Abundance | Abundance % |
|---|---|---|
| 1199.0 | 234789 | 100.00 |
| 899.8 | 160915 | 68.54 |
| 719.9 | 40389 | 17.20 |
| 1798.5 | 27198 | 11.58 |
| 1797.8 | 19702 | 8.39 |

#### Conjugate 4

Filename : C:\CHEM32\1\DATA\20210116\20210116 2021-01-16 17-07-34\ES12190-1331-P1B.D
Compound ID : ES12190-1331-P1B
Sample ID : ES12190-1331-P1B
Mobile Phase : A: 0.1%TFA in H2O B: 0.1%TFA in ACN
Flow : 1.0ml/min
Column : Gemini-NX C18 5um 110A 150*4.6mm
Instrument : Agilent 1200 HPLC-BE(1-614)

==========================================================================

Injection Date : Sat, 16. Jan. 2021 Location : Vial 46
Injection Time : 7:21:43 PM Inj. Vol. : 10.0 ul
Acq Method : C:\Chem32\1\DATA\20210116\20210116 2021-01-16 17-07-34\G20-50_20+3MIN.M

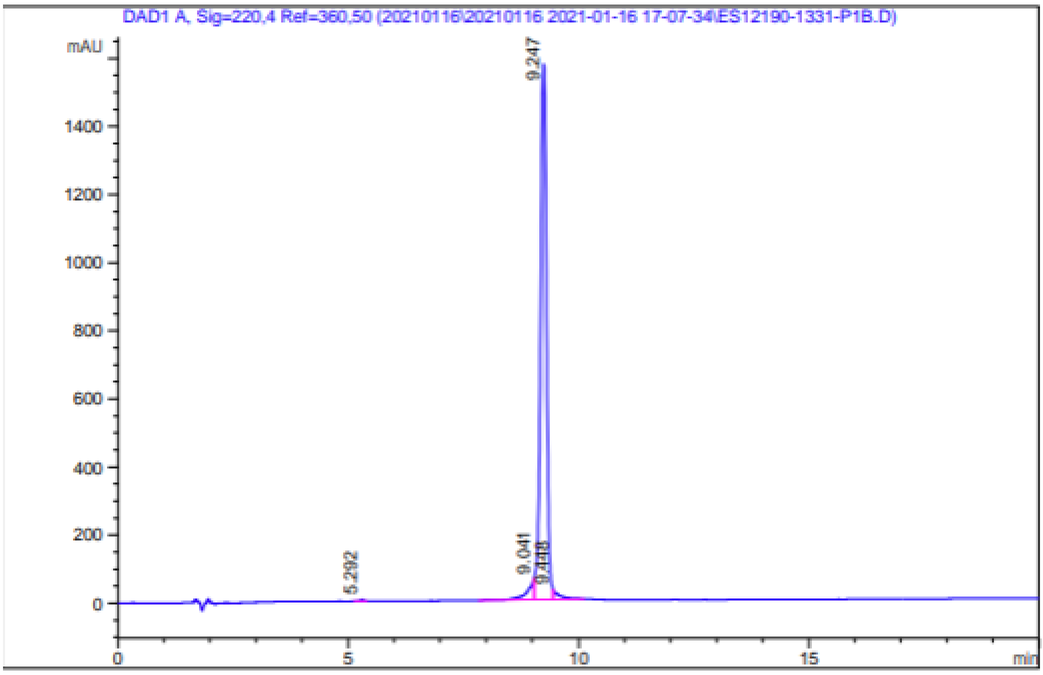

==========================================================================

Area Percent Report

==========================================================================

Signal->:DAD1 A, Sig=220,4 Ref=360,50

| Peak # | RT [min] | Height | Height % | Width [min] | Area | Area % |
|---|---|---|---|---|---|---|
| 1 | 5.292 | 2.924 | 0.175 | 0.091 | 16.865 | 0.119 |
| 2 | 9.041 | 62.039 | 3.719 | 0.163 | 605.734 | 4.260 |
| 3 | 9.247 | 1573.396 | 94.323 | 0.141 | 13329.167 | 93.742 |
| 4 | 9.448 | 29.742 | 1.783 | 0.150 | 267.210 | 1.879 |

| | | | |
|---|---|---|---|
| **Data Filename** | ES12190-1331-P1A.d | **Sample Name** | ES12190-1331-P1A |
| **Sample Type** | Sample | **Position** | P1-B8 |
| **Instrument Name** | Instrument 1 | **User Name** | |
| **Acq Method** | 10-80AB_2MIN-400.m | **Acquired Time** | 1/15/2021 9:53:12 AM |
| **IRM Calibration Status** | Not Applicable | **DA Method** | Default.m |
| **Comment** | | | |

**Sample Group** **Info.**

**Acquisition SW Version** 6400 Series Triple Quadrupole B.06.00 (B6025.4 [illegible])

#### User Chromatograms

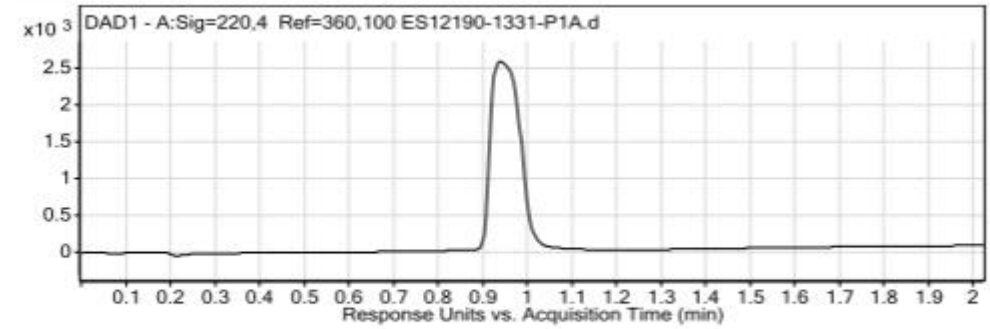

#### User Spectra

| Fragmentor Voltage | Collision Energy | Ionization Mode |
|---|---|---|
| 120 | 0 | ESI |

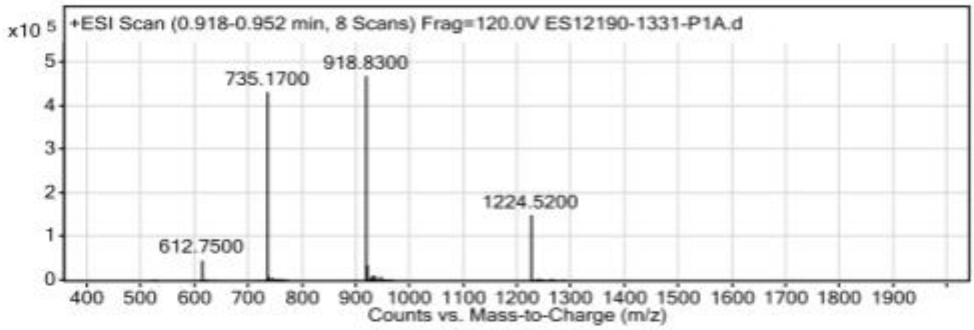

**Peak List**

| *m/z* | z | Abund |
|---|---|---|
| 735 | | 53249.06 |
| 735.17 | | 433253.16 |
| 735.32 | | 98805.48 |
| 918.43 | | 166771.44 |
| 918.67 | | 331814 |
| 918.83 | | 470390.44 |
| 1224.23 | 2 | 111092.84 |
| 1224.4 | 2 | 91019.02 |
| 1224.52 | | 150640.5 |
| 1225.06 | | 51635.01 |

### Conjugate 5

| | | | |
|---|---|---|---|
| **Data file:** | ES19551-443-P1B.dx | | |
| **Sequence Name:** | 2022-03-11 13-49-08+08-00 | **Project Name:** | Installation HPLC |
| **Sample name:** | ES19551-443-P1B | **Operator:** | SYSTEM |
| **Instrument:** | LCMS-QC | **Injection date:** | 2022-03-12 04:03:59+08:00 |
| **Inj. volume:** | 8.000 | **Location:** | P1-F1 |
| **Acq. method:** | 20-50-20min-P1waste.amx | **Colum:** | Gemini-NX C18 5um 110A 150mm4.6 |
| **Processing method:** | *GC_LC Area Percent_DefaultMethod.pmx | **Mobile phase:** | A:0.1%TFAin H2O B:0.1%TFAin ACN |

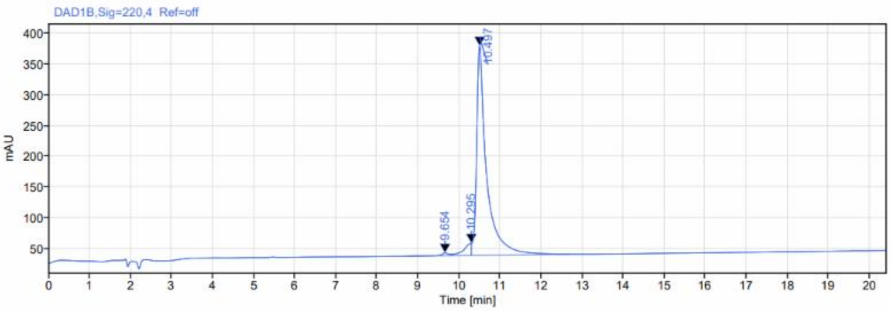

**Signal:** DAD1B,Sig=220,4 Ref=off

| RT [min] | Type | Width [min] | Area | Height | Area% | Name |
|---|---|---|---|---|---|---|
| 9.654 | BV | 1.12 | 61.56 | 4.50 | 1.02 | |
| 10.295 | VM m | 0.49 | 250.04 | 19.89 | 4.13 | |
| 10.497 | MB m | 2.41 | 5737.41 | 339.94 | 94.85 | |
| | | **Sum** | **6049.01** | | | |

| | | | |
|---|---|---|---|
| **Sequence Name:** | 2022-03-11 11-38-37+08-00 | | |
| **Data file:** | ES19551-443-P1A.dx | **Project Name:** | Installation |
| **Sample name:** | ES19551-443-P1A | **Operator:** | SYSTEM |
| **Instrument:** | LCMS-QC | **Acquired on:** | 2022-03-11 14:55:49+08:00 |
| **Inj. volume:** | 10.000 | **Location:** | P1-F1 |
| **Acq. method:** | 10-80-2min-1.5_P2.amx | | |
| **Processing method:** | *MS_DefaultMethod.pmx | | |

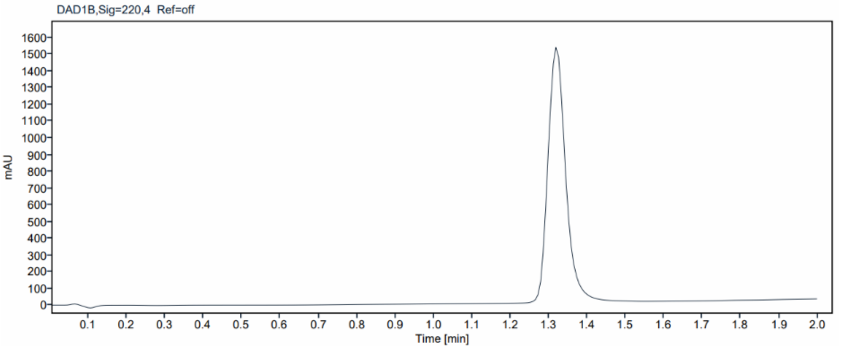

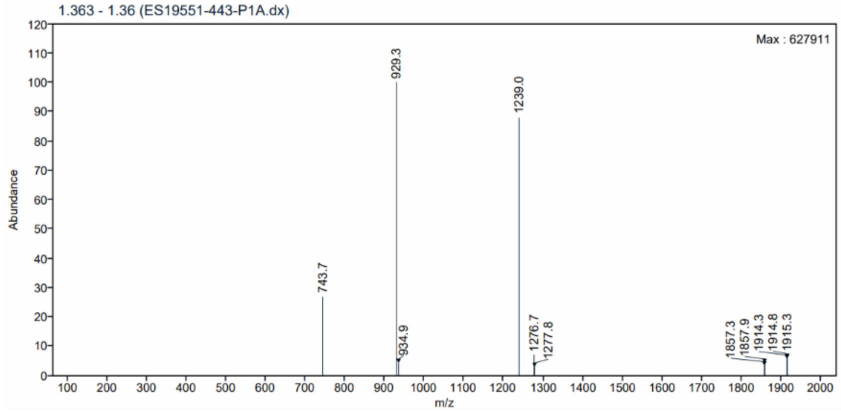

### Conjugate 6

| | | | |
|---|---|---|---|
| **Data file:** | ES19551-187-P1B.dx | | |
| **Sequence Name:** | 2021-09-15 16-15-13+08-00 | **Project Name:** | Installation HPLC |
| **Sample name:** | ES19551-187-P1B | **Operator:** | SYSTEM |
| **Instrument:** | LCMS-QC | **Injection date:** | 2021-09-16 07:40:14+08:00 |
| **Inj. volume:** | 3.000 | **Location:** | P2-D4 |
| **Acq. method:** | 20-50-20min-P1waste.amx | **Colum:** | Gemini-NX C18 5um 110A 150mm4.6 |
| **Processing method:** | *GC_LC Area Percent_DefaultMethod.pmx | **Mobile phase:** | A:0.1%TFAin H2O B:0.1%TFAin ACN |

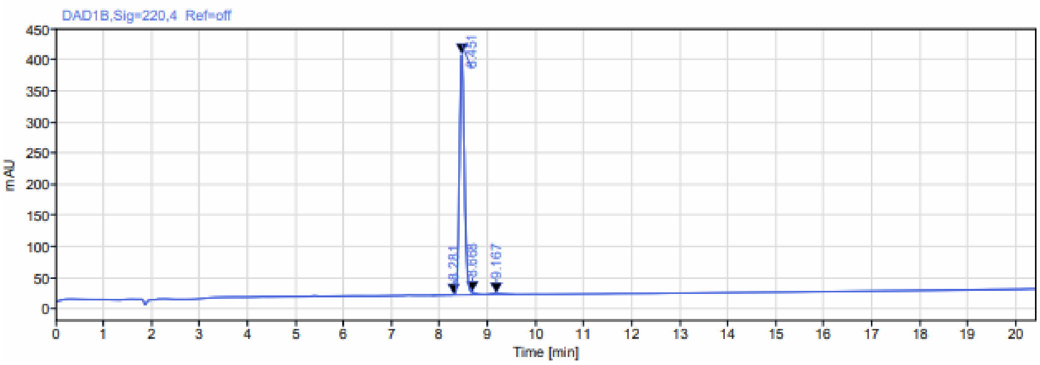

**Signal:** DAD1B,Sig=220,4 Ref=off

| RT [min] | Type | Width [min] | Area | Height | Area% | Name |
|---|---|---|---|---|---|---|
| 8.281 | BM m | 0.46 | 7.99 | 0.72 | 0.28 | |
| 8.451 | MM m | 0.38 | 2778.23 | 386.87 | 97.53 | |
| 8.668 | MV m | 0.25 | 26.34 | 4.44 | 0.92 | |
| 9.167 | VB | 0.61 | 35.99 | 1.96 | 1.26 | |
| | | **Sum** | **2848.54** | | | |

**Signal Name** MS1 +TIC SCAN ESI Frag=135V Gain=1.0

**Peak Retention Time** 1.556

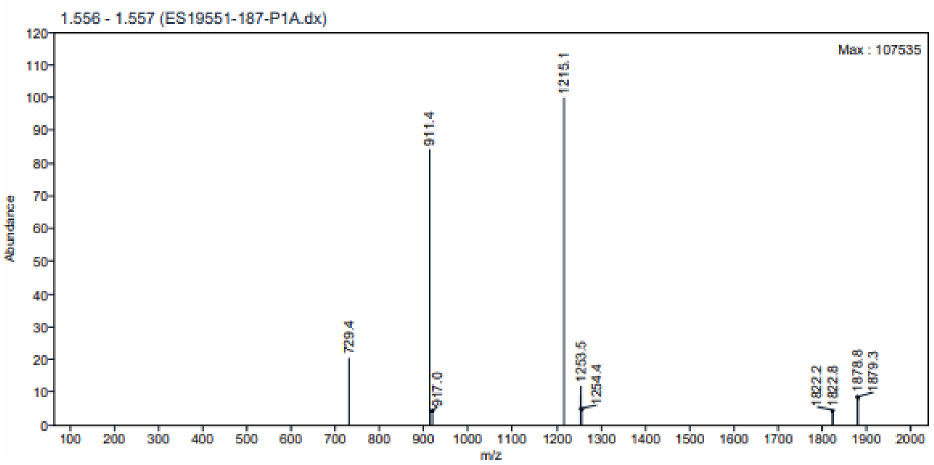

**MS Peak Table**

| m/z | Abundance | Abundance % |
|---|---|---|
| 1215.1 | 107535 | 100.00 |
| 911.4 | 90406 | 84.07 |
| 729.4 | 21925 | 20.39 |
| 1253.5 | 12725 | 11.83 |
| 1252.7 | 9832 | 9.14 |
| 1878.8 | 9693 | 9.01 |
| 1879.3 | 8156 | 7.58 |

#### Conjugate 7

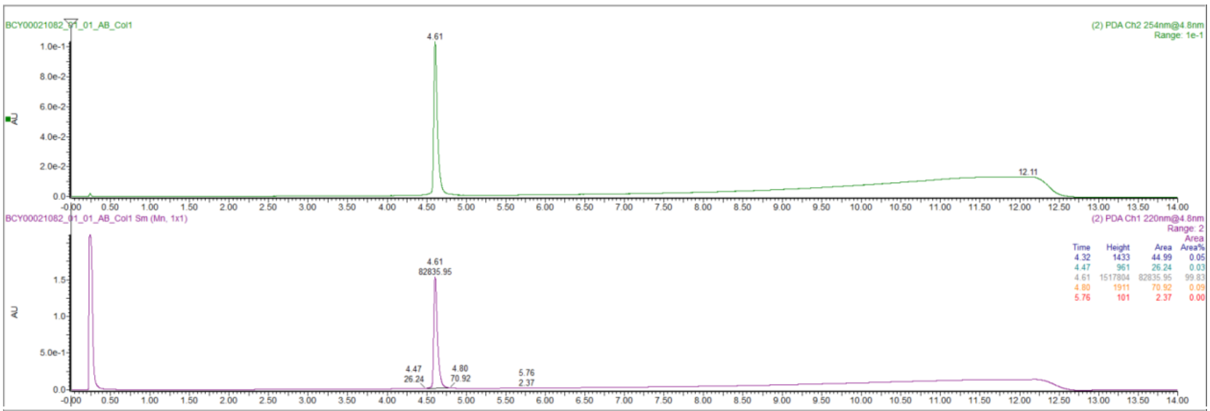
BCY00021082_01_01_AB_Col1
(2) PDA Ch2 254nm@4.8nm
Range: 1e-1
4.61
12.11
BCY00021082_01_01_AB_Col1 Sm (Mn, 1x1)
(2) PDA Ch1 220nm@4.8nm
Range: 2
AU
4.61
82835.95
4.47
26.24
4.80
70.92
5.76
2.37
Time Height Area Area%
4.32 1433 44.99 0.05
4.47 961 26.24 0.03
4.61 1517804 82835.95 99.83
4.80 1911 70.92 0.09
5.76 101 2.37 0.00

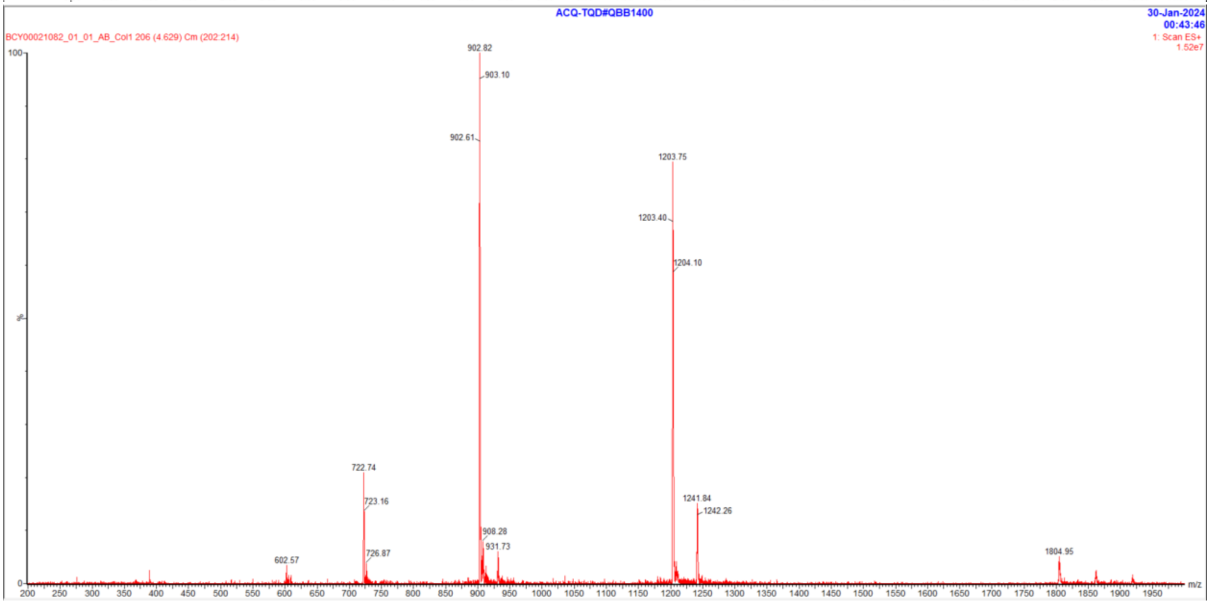
ACQ-TQD#QBB1400
30-Jan-2024
00:43:46
BCY00021082_01_01_AB_Col1 206 (4.629) Cm (202:214)
1: Scan ES+
1.52e7
902.82
903.10
902.61
1203.75
1203.40
1204.10
722.74
723.16
726.87
602.57
908.28
931.73
1241.84
1242.26
1804.95
m/z

#### Conjugate 8

**Sample name:** (101-08-00)-K[N3]K-DF-18563 RP QC
**Instrument:** 1260_1
**Injection date:** 11/4/2019 12:27:21 PM
**Acq. method:** 0595B_AB_Poroshell1 20_15.5min.M

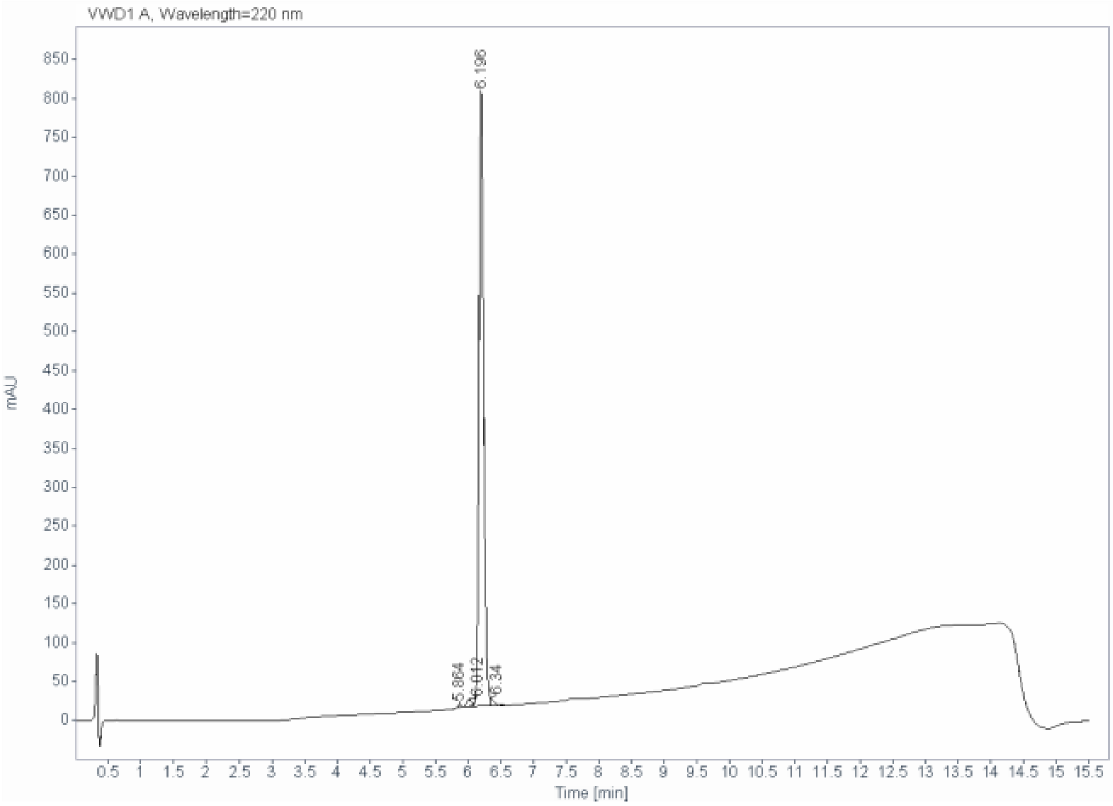

**Signal:** VWD1 A, Wavelength=220 nm

| RT [min] | Type | Width [min] | Area | Height | Area% |
|---|---|---|---|---|---|
| 5.864 | BB | 0.0605 | 17.6040 | 4.6144 | 0.4390 |
| 6.012 | BV E | 0.0828 | 51.9744 | 9.3539 | 1.2962 |
| 6.196 | VF R | 0.0756 | 3902.7305 | 790.5502 | 97.3337 |
| 6.340 | VB | 0.0601 | 37.3299 | 10.3479 | 0.9310 |
| | | Sum | 4009.6388 | | |

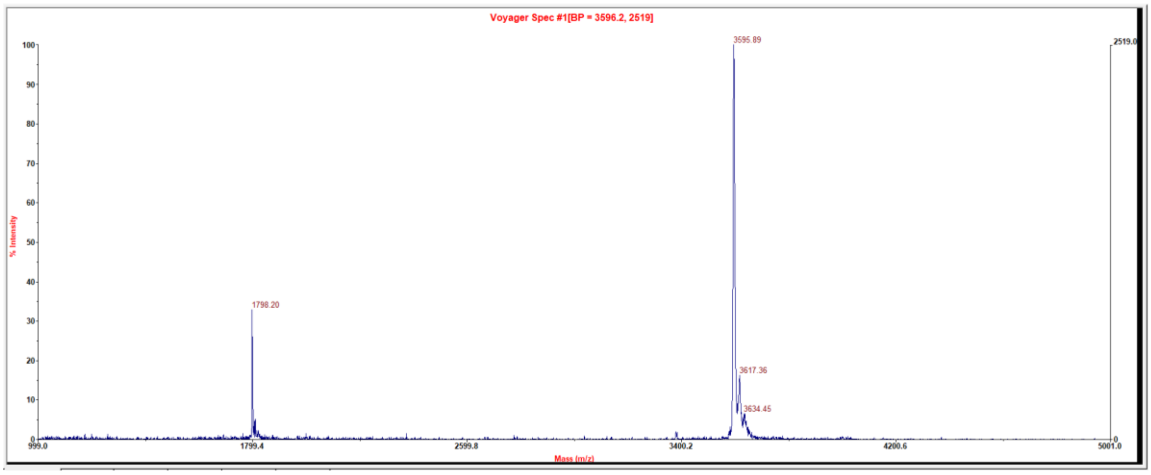

#### Tracer 1

| | |
|---|---|
| **Sample name:** | Fl-G-Sar5-(101-08-00) #2 |
| **Instrument:** | 1260_2 |
| **Injection date:** | 10/26/2020 4:48:46 PM |
| **Acq. method:** | 0595B_AB_KxC18_QC _20min.M |

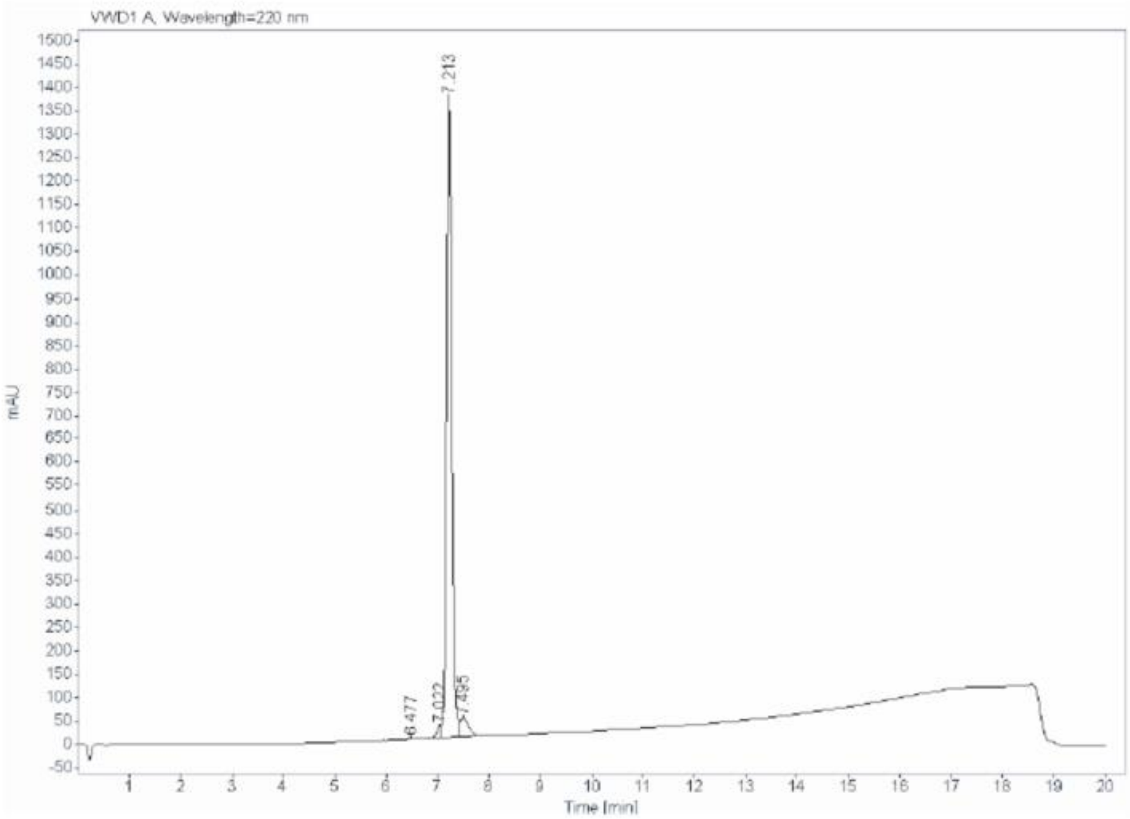

**Signal:** VWD1 A, Wavelength=220 nm

| RT [min] | Type | Width [min] | Area | Height | Area% |
|---|---|---|---|---|---|
| 6.477 | VV | 0.0752 | 7.8869 | 1.6659 | 0.0784 |
| 7.022 | VV F | 0.1011 | 170.6320 | 24.5470 | 1.6972 |
| 7.213 | VV | 0.1043 | 9408.2314 | 1367.5016 | 93.5769 |
| 7.495 | VB | 0.1672 | 467.2570 | 40.3145 | 4.6475 |
| | | Sum | 10054.0073 | | |

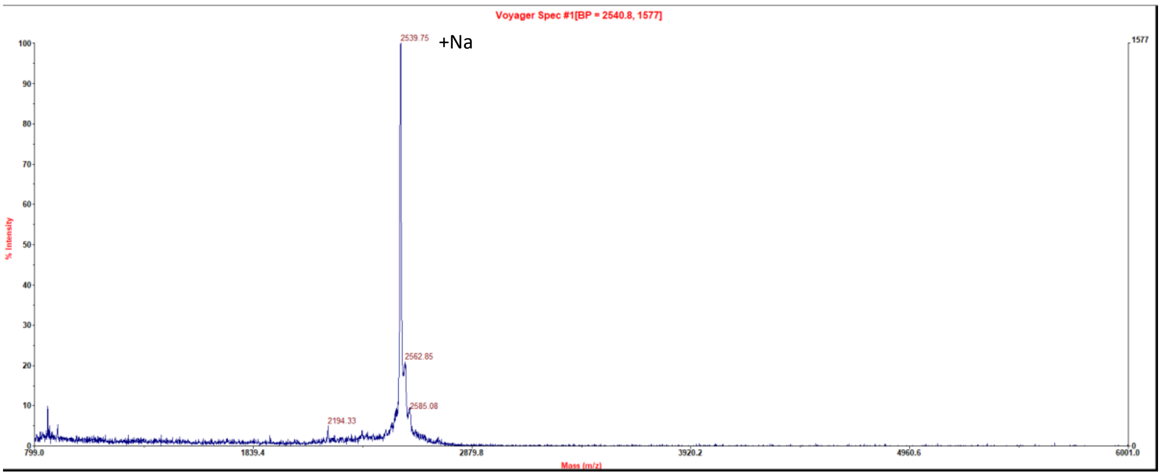

#### Tracer 2

| | |
|---|---|
| Sample name: | 101-10-00-Sar6-K(Fl)(TATA) QC |
| Instrument: | 1260_2 |
| Injection date: | 6/27/2019 3:00:40 PM |
| Acq. method: | 0595B_15.5min.M |

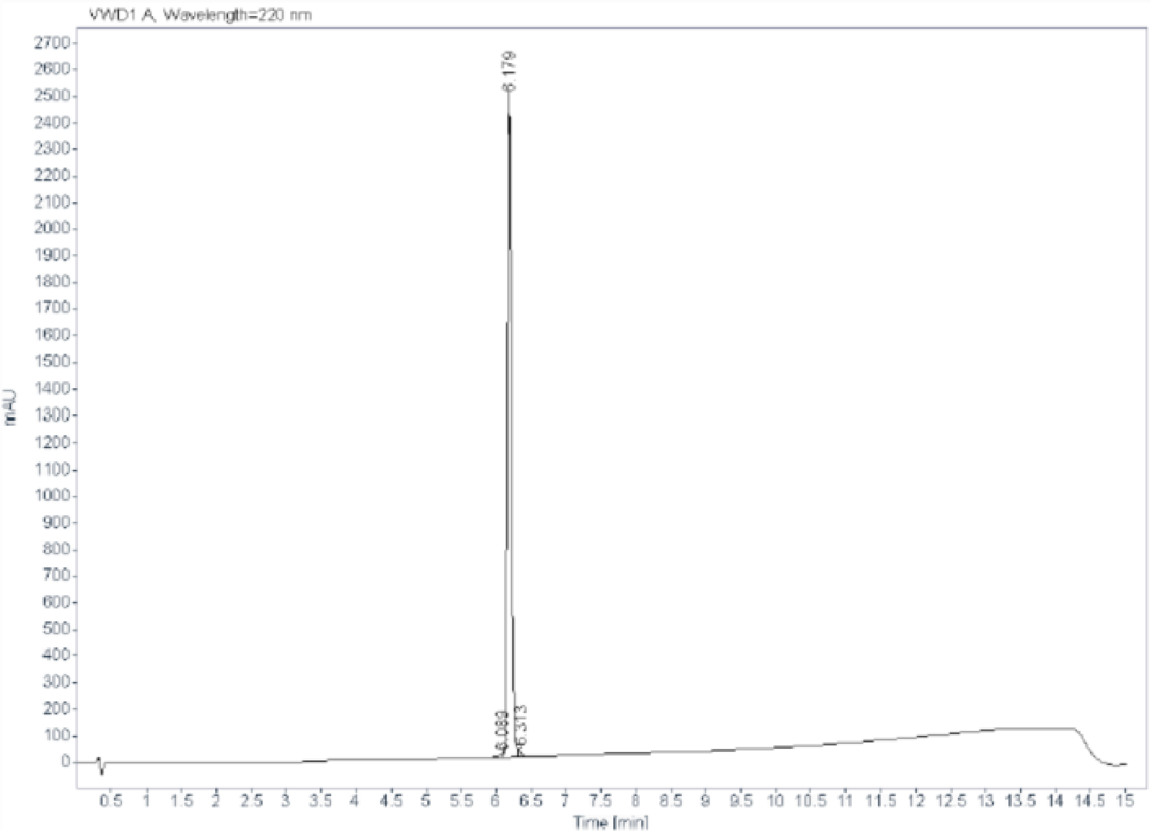

**Signal:** VWD1 A, Wavelength=220 nm

| RT [min] | Type | Width [min] | Area | Height | Area% |
|---|---|---|---|---|---|
| 6.089 | MF | 0.1085 | 69.8961 | 10.7321 | 0.6991 |
| 6.179 | FM | 0.0660 | 9854.9658 | 2488.1157 | 98.5732 |
| 6.313 | FM | 0.0434 | 72.7495 | 27.9667 | 0.7277 |
| | | Sum | 9997.6114 | | |

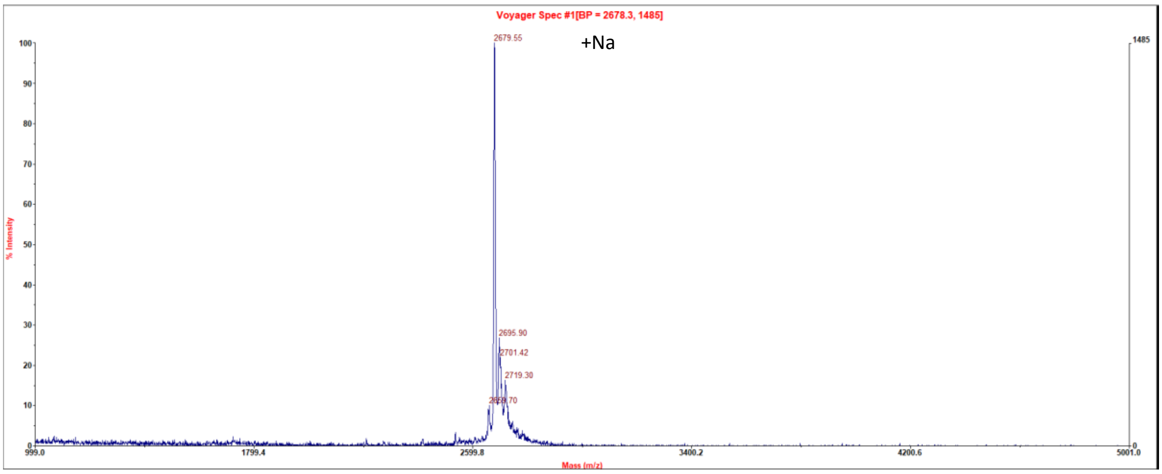
